# Using optimal control to understand complex metabolic pathways

**DOI:** 10.1101/2020.05.07.082198

**Authors:** Nikolaos Tsiantis, Julio R. Banga

## Abstract

**Background:** We revisit the idea of explaining and predicting dynamics in biochemical pathways from first-principles. A promising approach is to exploit optimality principles that can be justified from an evolutionary perspective. In the context of the cell, several previous studies have explained the dynamics of simple metabolic pathways exploiting optimality principles in combination with dynamic models, i.e. using an optimal control framework. For example, dynamics of gene expression in small metabolic models can be explained assuming that cells have developed optimal adaptation strategies. Most of these works have considered rather simplified representations, such as small linear pathways, or reduced networks with a single branching point.

**Results:** Here we consider the extension of this approach to more realistic scenarios, i.e. biochemical pathways of arbitrary size and structure. We first show that exploiting optimality principles for these networks poses great challenges due to the complexity of the associated optimal control problems. Second, in order to surmount such challenges, we present a computational framework based on multicriteria optimal control which has been designed with scalability and efficiency in mind, extending several recent methods. This framework includes mechanisms to avoid common pitfalls, such as local optima, unstable solutions or excessive computation time. We illustrate its performance with several case studies considering the central carbon metabolism of *S. cerevisiae* and *B. subtilis*. In particular, we consider metabolic dynamics during nutrient shift experiments.

**Conclusions:** We show how multi-objective optimal control can be used to predict temporal profiles of enzyme activation and metabolite concentrations in complex metabolic pathways. Further, we show how the multicriteria approach allows us to consider general cost/benefit trade-offs that have been likely favored by evolution. In this study we have considered metabolic pathways, but this computational framework can also be applied to analyze the dynamics of other complex pathways, such as signal transduction networks.

## Background

This paper is based on two main concepts: (i) genetic and biochemical dynamics are key to understand biological function, and (ii) optimality hypotheses enable predictions in biology. We start by briefly reviewing the relevant previous literature, with emphasis on early works. Then, we discuss the integration of these two main ideas in an optimal control framework which can then be used to analyze and understand complex biochemical pathways. We finally illustrate this approach with case studies related with metabolic networks.

### Dynamics

Mathematical modelling allows us to understand complex biological systems [1–3]. Wolkenhauer and Mesarovic [4] argue that the central dogma of systems biology is that system dynamics give rise to the functioning and function of cells. In this view, the language of dynamical systems is used to represent mechanisms at different levels (metabolic, signalling and gene expression) in order to describe the observed biochemical and biological phenomena. This mechanistic dynamic modelling is usually carried out using systems of coupled ordinary differential equations [5, 6]. Dynamical systems theory has a long history in physiology [2, 7, 8], and is receiving increasing attention in molecular systems biology [9–17].

The dynamic behaviour of biological systems is often explained in terms of feedback regulation mechanisms [4, 18]. Feedback is also a pillar in control theory, and in this context it is important to remember that the early concepts of feedback control were inspired by the study of regulation in biosystems [2]. Not surprisingly, a number of researchers have suggested that molecular systems biology can greatly benefit from the powerful methods of modern systems and control theory [1, 19–24]. Similar arguments have been made for synthetic biology [25–29].

### Optimality

Can biology be predictive? It has been argued that the systems approach to biology will enable us to predict biological outcomes despite the complexity of the organism under study [30–32]. According to Sutherland [33], optimality is the only approach biology has for making predictions from first principles.

The optimality hypothesis as an underlying principle in living matter has a long history. It was already used in the early 1900s to explain structural and functional organization in physiology [34, 35]. A few decades later, Rosen [36] published what seems to be the first monograph dedicated to optimality in biology, reviewing a number of applications. In this book, Rosen started highlighting the successful use of optimality principles in physics (where they are usually called variational principles), and their interrelationships with other disciplines including biology, mathematics and the social sciences (especially with economics). Rosen then proceeded to discuss how evolution via natural selection explains the appearance of optimality principles in biology, and how these principles can explain biological systems. These ideas were updated in a later publication [37], noting the two distinct (yet related) roles that optimality can play:

- (i) analytic (in the sense of explanatory), i.e. helping us to understand, or even predict, the way in which a biological system occurs. Many examples can be found in ecology [38] and evolutionary [39, 40] and behavioral biology [41].
- (ii) synthetic (in the sense of design and/or decision making), where it helps us to decide how to optimally manipulate, change or even build a bioprocess or biosystem. Examples can be found in biomedicine [42], metabolic engineering [43–45] and synthetic biology [45, 46].

### Optimality in cellular systems

In the remainder of this paper, we will focus on the explanatory role of optimality in the context of cellular systems. Savageu [47] and Heinrich and collaborators[48–51] developed many of the early theoretical applications of optimality principles in this domain, mostly to analyze metabolic networks and their regulation. Other notable examples of optimization studies in the context of biochemical pathways can be found in [43–46, 52–62] and references cited therein. Chapters specifically devoted to the interrelation between evolution and optimality in the context of biochemical pathways can be found in [63, 64]. From all these studies, two key ideas can be distilled: (i) evolution via natural selection can be understood as a fitness optimization process; (ii) therefore fitness optimality should allow us to understand, explain and even predict the evolution of the design of biochemical networks provided we can characterize their function(s) and how such function(s) impact on the organism fitness.

Heinrich et al [50] noted that the optimality hypothesis finds support in the fact that perturbations (e.g. mutations) or changes in the structure of enzymes usually leads to worse functioning of the metabolism. These authors also reviewed the cost functions most frequently considered in metabolic networks, concluding that they were mostly related with fluxes, concentrations of intermediates, transition times and thermodynamic efficiencies.

In this context, it is worth remembering that any optimization problem involves at least two elements: the objective (or cost) function, i.e. the criterion being maximized or minimized, and the decision variables, i.e. the degrees of freedom of the system which can vary to seek the optimal cost. Additionally, in most realistic situations the optimization problem also needs to incorporate constraints, i.e. relationships describing what is feasible or acceptable (i.e. requirements and fundamental limitations). Both in biology and physics, the set of constraints include physicochemical laws (e.g. conservation of mass, thermodynamics). But a main difference between physics and biology is the presence of additional functional constraints in the latter [1]. That is, biological systems evolve to fulfill functions, with insufficient performance leading to extinction.

### Optimality and dynamics: optimal control theory

Most of the above mentioned early studies of optimality in biology considered stationary (i.e. steady state) systems. However, as remarked at the beginning of this paper, biological function is closely linked to dynamics. Thus we now focus on the question of explaining and/or predicting dynamics (i.e. function) exploiting optimality principles.

The optimization of dynamical systems is studied by optimal control theory [65, 66]. Optimal control considers the optimization of a dynamic system, that is, one seeks the optimal time-dependent input (control) to minimize or maximize a certain performance index (cost function). Optimal control is sometimes also called dynamic optimization when the system is open loop. The basic elements of an optimal control problem are the performance index (cost function to be optimized), the control variables (time-varying decision variables), and the constraints (which can be inequalities or equalities, dynamic or static, and can be active during all or part of the time horizon considered). Typically, equality dynamic constraints constitute the time-varying model of the system under consideration.

Rosen was probably the first to recognize the importance of optimal control as an unifying framework that could bring together important areas of theoretical biology. Although in his works [36, 37] he outlined the potential of optimal control theory for biological research, he did not present any illustrative example. Successful applications of optimal control in biology and biomedicine started to appear in the 1970*s*, with notable impetus in the case of optimal decision making in biomedical engineering (e.g. drug scheduling [67]). Around the same time, explanatory uses of optimal control in biology started to appear (an introductory book discussing several early examples can be found in [68]).

In recent years, optimal control has been increasingly used to explain cellular phenomena (e.g. [69–71]). In the case of metabolic systems and their regulation, Klipp et al [72] were the first to predict temporal gene expression profiles in a metabolic pathway assuming optimal function under a constraint limiting total enzyme amount. They used an optimization approach similar to control parameterization methods in numerical optimal control. Notably, these researchers considered a simplified model of the central metabolism of yeast undergoing a diauxic shift, and they were able to predict time-dependent enzyme profiles which agreed well with experimental gene expression data [73]. Further, considering a simple linear pathway model, they found a wave-like enzyme activation profile which agreed with previous observations of gene expression during the cell cycle [74, 75], indicating that for this pathway topology, genes involved in a certain function are activated when such function is needed. Interestingly, these results were later experimentally confirmed by Alon and co-workers [76], who named this kind of activation profile “just-in-time” (a term originated in industrial manufacturing to describe a methodology aimed at reducing production times and inventory sizes). As noted by Ewald et al [77], this timing pattern had also been previously observed in the flagellum assembly pathway [78]. Similar activity motifs were also found in the timing in transcriptional control of the yeast metabolic network [79].

There are several important messages in the pioneering work of Klipp et al [72]: (i) it is possible to predict the dynamics of biological pathways without knowing the underlying regulatory mechanisms, confirming the concepts proposed by Heinrich and collaborators a decade earlier [50]; (ii) the timing of gene expression allows the cell metabolism to optimally adapt to varying external conditions; (iii) the optimal operation of the pathway needs to take into account constraints that arise from physico-chemical limitations of the cell (e.g. total enzyme concentration is limited by protein synthesis capacity); (iv) for the case studies considered, the authors considered different (and single) objective functions, which encapsulate the optimality hypothesis.

The work of Klipp et al [72] spurred the application of optimal control to characterize cellular dynamics related with metabolism [80–96]. Ewald et al [77] have recently reviewed many of these studies, illustrating how dynamic optimization is a powerful approach that can be used to decipher the activation and regulation of metabolism. Furthermore, Ewald et al [77] have also reviewed other important types of applications, including dynamic resource allocation in cells (as studied by e.g. [91]), the genomic organization of metabolic pathways (e.g. [85]), the development of effective treatments against pathogens (e.g. [93, 97]), and other manifold applications in metabolic engineering and synthetic biology.

### Cellular trade-offs and multicriteria optimality

Trade-offs are ubiquitous in evolutionary biology: very often one trait can not improve without a decrease in another, as already noted by Darwin [98]. Numerous works have studied cellular trade-offs considering e.g. the design of microbial metabolism [99, 100] and strategies for resource allocation, storage and growth [101–104]. Similarly, trade-offs between economy and effectiveness have been found in biological regulatory systems [105]. Analyses of different trade-offs have been used to uncover design principles in cell signalling networks [106–108].

A mathematical framework that combines in a natural way the concepts of optimality and trade-offs is multicriteria optimization, where one seeks the best (optimal) trade-offs (the so called Pareto optimal set) that correspond to the simultaneous optimization of several objectives. The concept of Pareto optimality was originally developed in the field of economics, but it was already suggested for applications to ecology in the early 1970*s* [109]. In the case of biochemical networks, multicriteria optimality was discussed by Heinrich and Schuster in the 1990*s* for metabolic steady-state conditions [49, 63, 110]. During the 2000*s*, the approach gained traction: El-Samad et al [111] studied the Pareto optimality of gene regulatory network associated with the heat shock response in bacteria; multi-objective approaches were used to perform metabolic network optimization [112, 113] and flux balance analysis [114–116]. More recently, Alon and co-workers applied Pareto optimality to explain evolutionary trade-offs [117] and biological homeostasis systems [105]. Higuera et al [118] analyzed optimal trade-offs for the allosteric regulation of enzymes in a simple model of a metabolic substrate-cycle. Different optimal tradeoffs in molecular networks capable of regulation and adaptation have also been studied in the context of synthetic biology [119–123]. Many other applications of multicriteria optimality in biology are reviewed in [124–126].

Considering the prediction of dynamics in biochemical pathways, to the best of our knowledge the study of de Hijas-Liste et al [86] was the first to apply a multicriteria dynamic optimization framework. These authors proposed multi-objective formulations for several metabolic case studies, showing how this framework provided biologically meaningful results in terms of the best trade-offs between conflicting objectives.

Both in single and multicriteria formulations there is an underlying key question that needs to be addressed: the selection of the objective functions. In other words, we need to formulate objective functions which encapsulate the fitness and the trade-offs of the biological system under study. Heinrich and Schuster [63] suggested following heuristic arguments and checking their validity by comparing the predictions derived from the associated optimality principle with experimental observations. This study-hypothesize-test approach has been the one followed by the vast majority of the works cited above. Recently, we proposed an alternative based on an inverse optimal control formulation [127] that aims to find the optimality criteria that, given a dynamic model, can explain a set of given dynamic (time series) measurements. In other words, inverse optimal control can be used to systematically infer optimality principles in complex pathways from measurements and a prior dynamic model.

### Challenges

As discussed above, optimal control can provide important insights in the domain of systems biology. So far, existing studies have made use of rather simple dynamic models (see the review of Ewald et al.[77]). In many situations, using a simple model might be in fact a totally satisfactory approach: an excellent illustration is the study by Giordano et al [91], where optimal control of coarse-grained dynamic models was used to explain resource allocation strategies in microbial physiology.

However, for many applications, more complex models need to be used. Can optimal control be applied at these larger scales? The answer to this question ultimately relies on two key issues: (i) the availability of the detailed (time-series) measurements need to build these more complex dynamic models, and (ii) the existence of numerical optimal control methods that can solve these problems in a reliable and efficient way. Regarding (i), Ewald et al.[77] mention recent advances in experimental techniques (like large-scale quantitative proteomic data) that should make it possible to apply these optimality principles at more complex molecular levels. In contrast, regarding (ii), the situation is currently not so clear. Despite the many significant advances during the last decades, reliably solving nonlinear optimal control problems can be very challenging, even for relatively small problems. This is true even in areas with a long optimal control tradition, such as aerospace [128].

In the case of biochemical pathways, the application of optimal control faces important challenges and pitfalls, including: complicated control profiles with many switching points and singular arcs, multimodality (local solutions), path constraints, possible discontinuous dynamics, and scaling issues. More details regarding these issues are given below.

### Our objectives and approach

Here, our goal is to provide a multi-objective optimal control approach that can be applied to dynamic models of complex biochemical pathways, surmounting the above mentioned challenges. In particular, we present a computational workflow that is reliable (robust), avoiding convergence to local solutions, but efficient enough (in terms of reasonable computation times) to handle realistic network topologies and arbitrary nonlinear kinetics. We will show how this workflow is capable of scaling up well with network size, and how it can handle multi-criteria formulations.

We illustrate the performance of this workflow considering three different case studies, based on dynamic models of the central carbon metabolism of *S. cerevisiae* and *B. subtilis*. In particular, we use our workflow to explain metabolic dynamics during nutrient shift experiments.

## Methods

### Optimal control problem: general formulation

We consider dynamic systems described by nonlinear ordinary differential equations (ODEs). The problem of *optimal control* (OCP) consists of computing the optimal decision variables (also called time-varying inputs, or controls) and time-invariant parameters that minimize (or maximize) a given cost functional (performance index), subject to the set of ODEs and possibly algebraic inequality constraints. Mathematically, the OCP is usually stated as follows:

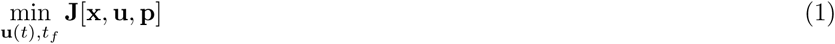

Subject to:

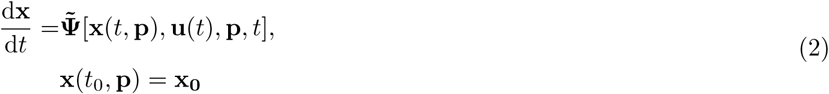

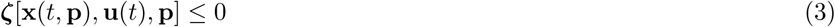

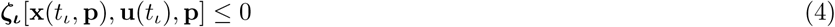

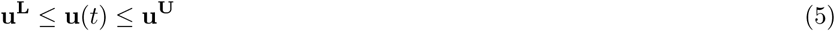

where **J**[**x, u, p**] is the objective functional (sometimes called performance index, or cost functional), encapsulating the optimality criteria; **u**(*t*) are the time-dependent control variables which must be computed in order to minimize (or maximize) the objective functional (**J**[**x, u, p**]). The problem is subject to constraints, including the dynamics of the system described by Eqns. (2), i.e. the set of ordinary differential equations and their corresponding initial values (**x**(*t*_0_)), forming the so-called initial value problem (IVP); inequality (***ζ***) path constraints are encoded in equations (3), representing inequalities relationships that must be enforced during the time horizon considered (e.g. total enzyme capacity, critical thresholds for specific concentrations, etc.). In some cases we also need to consider inequality (***ζ***_***ι***_) time-point constraints, encoded in Eqns. (4). Finally, (**u**^*U*^, **u**^*L*^) are the upper and lower bounds for the control variables, as stated in Eqns (5).

In the case of a multicriteria formulation, the cost functional **J**[**x, u, p**] is a set of objective functions corresponding to the *N* different criteria considered:

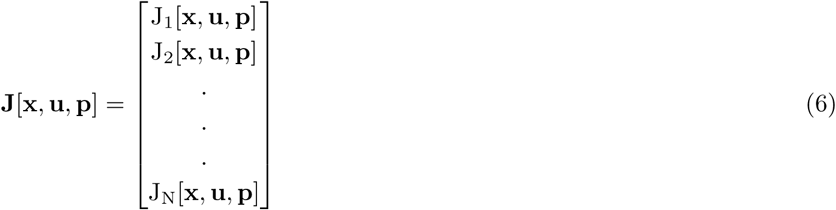

where, in its general form, each objective functional *J*_*i*_ in this set (*i* ∈ [1, *N*]) consists of a Mayer 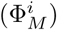 and a Lagrange 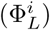 term:

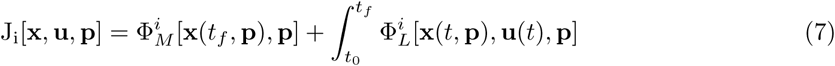

We assume the same general form for the single-objective case.

#### Multicriteria optimal control

In multicriteria optimization, the objectives are usually in conflict (improving one damages the others), so the optimal solution is not unique, but a set of optimal trade-offs (i.e. the set of the best compromises, which is also called the Pareto set). Many methods have been developed for multicriteria optimal control, and they are usually classified in three categories [129]: scalarization techniques, continuation methods and set-oriented approaches. Here we use the *E*-constraint scalarization method, which transforms the original multiobjective problem into a finite set of single-objective optimal control problems.

The *ϵ*-constraint method proceeds by solving a single objective problem with respect to one of the objectives *J*_*a*_ while treating the rest of the objectives *J*_*i*_ as algebraic constraints:

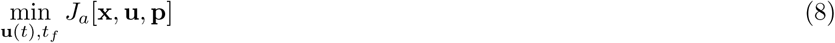

Subject to:

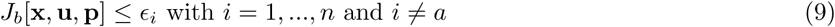

The above problem will also be subject to the rest of the differential and algebraic constraints considered in the original multicriteria formulation. Therefore the Pareto set is obtained by solving a set of single objective problem for different values of *ϵ*_*i*_.

### Numerical solution of nonlinear optimal control problems

Methods for the numerical solution of nonlinear optimal control problems can be classified under three categories: dynamic programming, indirect and direct approaches (Figure 1). Dynamic programming [130, 131] suffers from the so-called *curse of dimensionality*, so the latter two are the most promising strategies for realistic problems. Indirect approaches were historically the first developed and are based on the transformation of the original optimal control problem into a multi-point boundary value problem using Pontryagin’s necessary conditions [65, 66]. Direct methods transform the optimal control problem into a nonlinear programming problem (NLP). They are based on the discretization of either the control, known as the sequential strategy, or both the control and the states, known as the simultaneous strategy.

**Figure 1:**
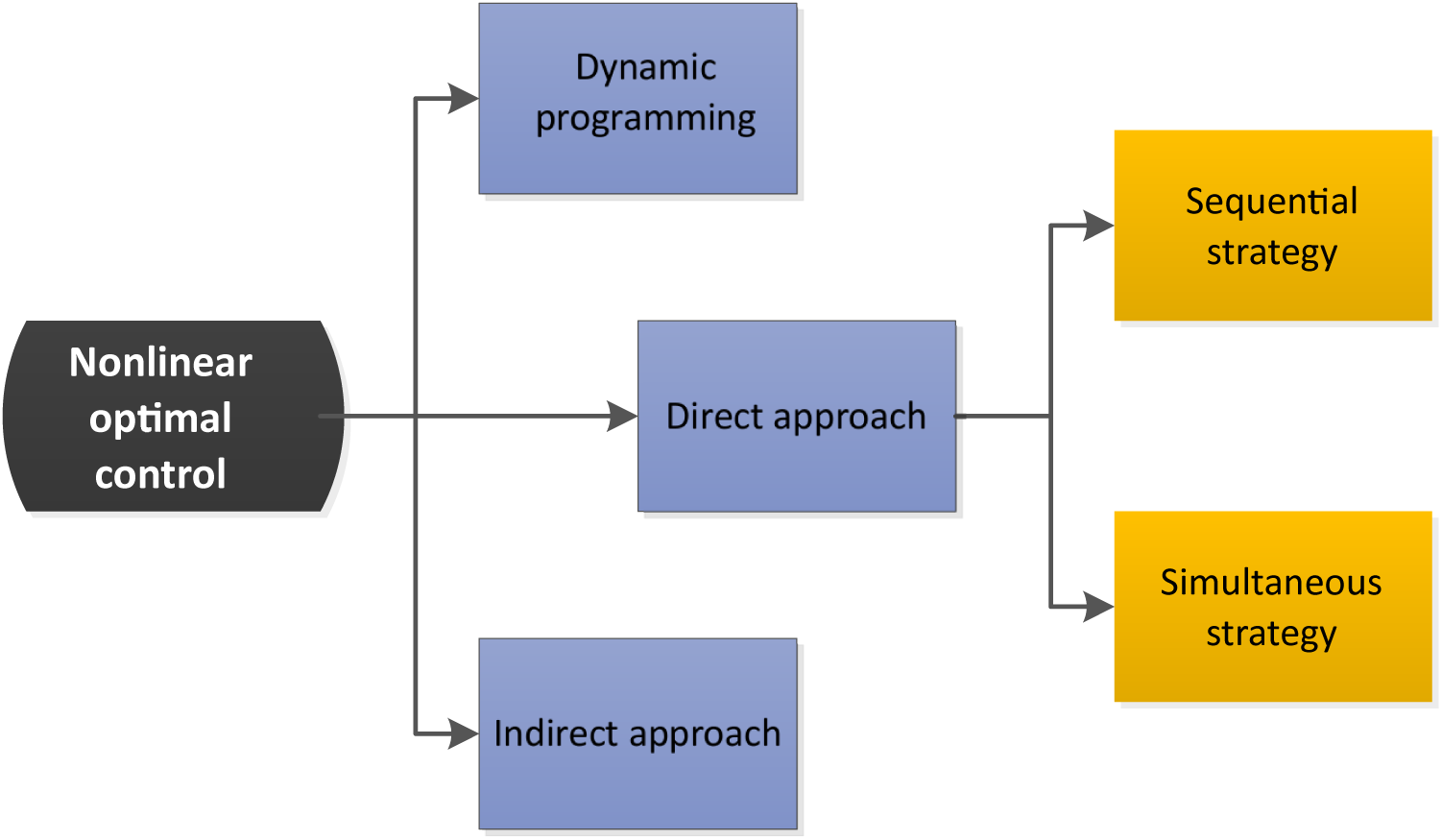
Classification of solution strategies for nonlinear optimal control problems. Figure adapted from [137].

#### Sequential strategy (control vector parameterization)

In the sequential strategy[132–134], also known as control vector parametrization, the controls are approximated by piecewise functions, usually by a low order polynomial, the coefficients of which are the decision variables. Thus, the problem is transformed into an outer non-linear programming (NLP) problem with an inner initial value problem (IVP) where the dynamic system is integrated for each evaluation of the cost function.

The control vector parameterization (CVP) approach proceeds by dividing the time horizon into a number of elements (*ρ*). The control variables (*j* = 1 *… n*_*u*_) are then approximated within each interval (*i* = 1 *… ρ*) by means of some basis functions, usually low order Lagrange polynomials [135], as follows:

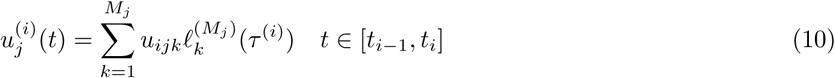

where *τ* is the normalized time in each element *i*:

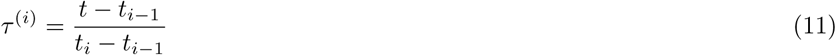

and *M*_*j*_ the order of the Lagrange polynomial (*£*). In this work we will consider *M*_*j*_ = 1 or *M*_*j*_ = 2, i.e. piecewise constant or piecewise linear approximations of the controls.

Using the above discretization, the controls can be expressed as functions of a new set of time invariant parameters corresponding to the polynomial coefficients (**w**). Therefore the original infinite dimensional problem is transformed into a non-linear programming problem with dynamic (the model) and algebraic constraints, and where the decision variables correspond to the unknown parameters in *θ* and **w**. In other words, this strategy gives rise to an outer NLP problem with an embedded inner initial value problem, i.e. the dynamics must be integrated for each evaluation of the cost function and the algebraic constraints.

#### Simultaneous strategy (complete discretization)

In the simultaneous strategy[136–138], also known as complete parameterization approach, both states and controls are discretized by dividing the whole time domain into small intervals. This is done either using multiple shooting, where similarly to the sequential approach, the problem is integrated separately in each interval and linked with the rest through equality constraints, or by a collocation approach, where the solution of the dynamic system is being coupled with the optimization problem.

The direct collocation approach is probably the most well known complete discretization apparch. In this method the solution of the infinite dimensional problem is transcribed into a non-linear programming problem discretizing both the states and the controls [138] by means of low-order polynomial approximations. The integration is usually approximated by a K-stage Runge-Kutta scheme. Different Runge-Kutta schemes use polynomials of different order (*K* +1) to approximate the system’s solution in each integration step (*i*) with stepsize *h*_*i*_:

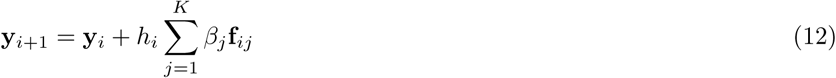

with:

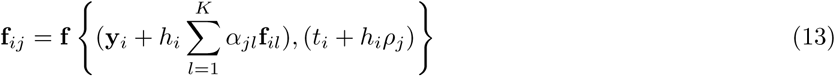

where **f** is the right hand side of the ODEs while *β*_*j*_, *α*_*jl*_, *ρ*_*j*_ are order-dependant known constants. Thus, this method transforms the original infinite dimensional problem into a large NLP problem which, in contrast to the CVP method above, does not require the integration of the dynamic system during the iterative solution of the NLP.

### Effect of constraints on the optimal solution

Constraints play a key role in mathematical modeling in biology. From the computational optimization point of view, constraints limit the solution space, adding further complexity to already complex mathematical formulations. However, from the biological point of view, constraints add information to the model, making it more realistic.

In general, constraints can have physical and biological meaning [63]. For example, thermodynamic feasibility is often neglected in metabolic models although it can provide information on both directionality of reactions and range of parameters [139]. Additionally, information on the physico-chemical properties and the stoichiometry of the network under consideration can greatly restrict the solution space.

Apart from physical constraints, biological limitations can be of even higher importance and complexity [37]. These limitations are not only complex to express but also hard to get estimates for. Considering variations of the critical values of certain metabolites or proteins, necessary for the survival of the cell, can have a quite significant effect on the optimal solution. Other relevant constraints can be related to the maximum protein production rate, the total enzyme burden or the total protein investment [48, 140].

In this context, it would be interesting to analyze the interplay between constraints in biological models with the optimal trade-offs (Pareto set). In some cases, constraints can act as the operating principles we are trying to identify and understand [141]. It would also be interesting to study (i) the impact of constraints not only on the current behavior of a biological network, and (ii) their role on how the network has evolved [119, 142].

In mathematical optimization, once an optimal solution is found, it is always interesting to analyze the role of the different constraints on such solution. In constrained optimization in economics, especially in linear programming, the concept of shadow price is used to quantify how much the optimal value of the objective function changes by relaxing a constraint [143, 144]. In optimal control the equivalent of the shadow price is the adjoint variable [68]. The adjoint variables ***λ***(*t*) (also called *costate variables*) can be estimated and provided along with the results as part of the solution of the optimal control problem. However, an interpretation of their significance is not necessarily intuitive. As mentioned previously, in nonlinear optimal control theory and while using direct methods, the problem is formulated as a constrained NLP optimization problem where both algebraic and differential equations are constraining the search space. In such a problem, the Lagrange multipliers represent what the cost would be if those constraints were violated. Additionally the Lagrange multipliers with respect to the state variables are discrete approximation of the adjoint variables, the approximation be depending on the transcription method of choice. Consequently, in nonlinear optimal control the adjoint variables can be used to explain the possible variation on the cost function associated with an incremental change in the states [68].

Using direct optimization methods the general optimal control problem is transcribed into a NLP problem. The original performance index functional is transcribed into the corresponding NLP cost function *F* (**y**), dependent on *n* variables **y**. This cost has to be minimized subject to the *m* ≤ *n* constraints:

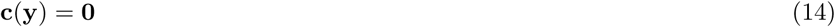

The Lagrangian is defined as:

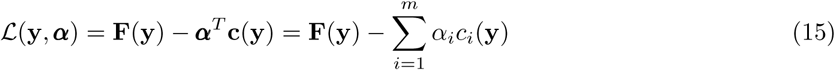

with ***α*** corresponding to the Lagrange multipliers.

The original infinite-dimensional optimal control problem has been transformed to a discretized (finite-dimensional) NLP problem, where the controls **u**(*t*) need to be computed by approximation in order to minimize the cost function:

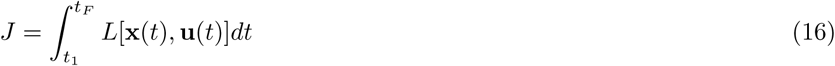

subject to the differential-algebraic equation (DAE) constraints:

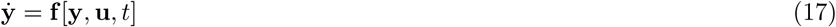

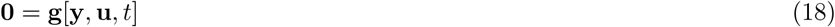

The NLP performance index is formed as follows:

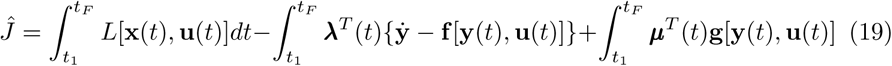

Note that in (16) and (19) *L* is the integrand:

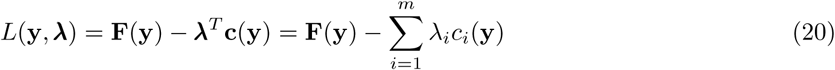

where ***λ*** are the adjoint variables.

The difference between the notation used for the Lagrangian (ℒ) with respect to the Lagrange multipliers (***α***) and the Lagrangian (*L*) with respect to the adjoint variables (***λ***) depends on how the continuous variables are approximated. In particular, the relationship between the Lagrange multipliers and the adjoint variables is dependent on the transcription method. Generally, it has been shown that the Lagrange multipliers estimate the adjoint variables in the limit of *ρ* → ∞, where *ρ* is the number of time intervals in which the time horizon has been discretized [138].

### Challenges and pitfalls in numerical optimal control

Using direct methods to solve optimal control problems involving nonlinear models frequently result in multimodal problems, i.e. there are multiple local solutions and at least a global one. This is a consequence of the presence of nonlinear dynamics and path constraints. As a result, local optimization methods will usually converge to bad local solutions [45, 145]. Often, researchers resort to the use of a multi-start strategy, i.e. initializing local methods from multiple points in the search space. However, this approach becomes inefficient for problems of realistic size [86].

In theory, deterministic global optimization methods can surmount these difficulties and find the global optimum with guarantees, but currently their computational cost does not scale up well with problem size [146]. Purely stochastic global optimization can also be used, but they are rather inefficient, requiring many evaluations of the cost function, resulting in large computation times that become prohibitive if refined solutions are sought. Alternatively, hybrid global-local optimization strategies combine the advantages of global and local methods, showing good performance and robustness for many realistic problems of small and medium size [86, 147]. However, these hybrid strategies also become too computationally expensive if we seek refined solutions of large optimal control problems.

In addition to multimodality, other numerical and computational issues in optimal control include complicated control profiles with multiple and very sensitive switching points, discontinuous dynamics, path constraints and singular arcs [86, 148].

### Our combined strategy for numerical optimal control

Although several surveys of numerical methods and software for optimal control have been published during the last decade [149–151], these reviews are neither very recent nor very exhaustive. More importantly, there is a lack of studies comparing different approaches in a fair way and using a set of well defined benchmark problems. Therefore, choosing the current best numerical method and software to solve the class of problems considered here becomes a daunting task, especially for non-experienced users. To make things worse, using many currently available software packages we might get solutions that look reasonable but which are, however, artifacts or bad local solutions.

Therefore, here we present a robust workflow and guidelines to avoid, as much as possible, the many challenges and pitfalls that are common in nonlinear optimal control. We have designed an strategy with three key ideas in mind, i.e. it should be able to: (i) run without good initial guesses, (ii) avoid convergence to local solutions, (iii) approximate complicated control profiles with good accuracy and reasonable computation cost, (iv) scale up well in terms of control and state variables. Further, the approach is able to handle multiobjective optimal control problems by transforming them into a set of nonlinear optimal control problems, as depicted in Figure 2.

**Figure 2:**
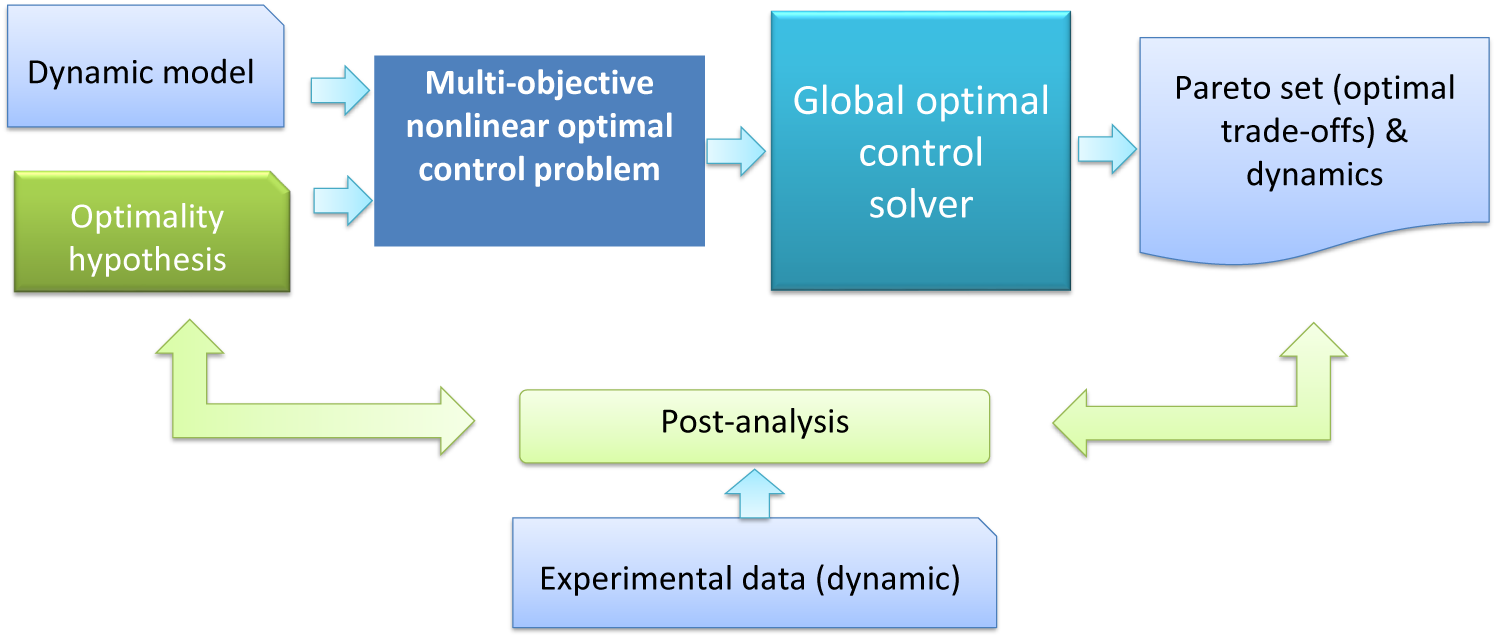
General workow of our approach

In order to meet these requirements, we tested a number of different options and finally arrived to the following two-phase strategy named `AMIGO2_DO+ICLOCS:`

- first phase using `AMIGO2_DO`, a hybrid stochastic-deterministic method based on control vector parameterization
- second phase using `ICLOCS`, a simultaneous (complete discretization) fast local method

The main justification for the first phase is the handling of the multimodality issues using a robust approach that has the additional benefit of not requiring good initial guesses. We have used the latest version of the `AMIGO2_DO` solver, a sequential optimal control solver included in the `AMIGO2` toolbox [152]. Although this solver allows the user to select different combinations of global and local methods, we have found that the enhanced scatter search (eSS) metaheuristic [153] provided the best performance. The control vector parametrization strategy implemented in this solver is easier to apply to arbitrary nonlinear dynamics, results in smaller optimization problems and relies on well tested initial value solvers.

For the second phase we used `ICLOCS` [154], a very efficient and fast optimal control solver based on complete discretization and deterministic local optimization methods. Since the second phase is initialized in a near-global solution provided by the first phase, its local optimization character is not an issue. And this second phase allows a very good aproximation of the control profiles due to the use of the complete discretization approach. The simultaneous strategy leads to larger optimization problems but has the advantage of avoiding the repeated integration of the system dynamics at each iteration, so it can approximate highly discretized control profiles very efficiently.

Choosing the right settings for these solvers is of key importance, but this task can be a real challenge for non-expert users. Here we provide the best settings that we found for the class of problems considered.

Recommended settings for the `AMIGO2_DO` phase:

- control discretization options: an initial piecewise-linear discretization of the control using 5 − 10 elements proved successful. When used alone, the `AMIGO2_DO` mesh-refinement options can be used to approximate better difficult controls. When used as part of `AMIGO2_DO+ICLOCS`, we found more efficient to refine the controls during the `ICLOCS` phase
- optimization solver: best results were obtained using the `eSS` (enhanced scatter search) global solver [153] with `FSQP` [155] as local solver
- initial value problem solver: `CVODES` [156], with relative and absolute integration tolerances settings of at least 1.0*E* − 7. It is important that the optimization criteria tolerance is at least two orders of magnitude larger then the IVP integration tolerance. Otherwise, the integration numerical noise introduces non-smoothness making the optimization solver to converge less efficiently or fail.

Recommended settings for the `ICLOCS` phase:

- initial guesses: a good initial guesses for the unknown parameters and state trajectories is key to improve convergence. When used in the hybrid approach, the solution obtained by `AMIGO2_DO` was used as the initial guess for `ICLOCS`.
- transcription method: trapezoidal
- derivative generation: analytic. Providing analytic expressions for the gradient of the cost function, Jacobian of the constraints and Hessian of the Lagrangian was found extremely mportant. Generating this information can be very cumbersome for non-trivial problems, so we have coded an auxiliary function that automates it using symbolic manipulation
- NLP solver: IPOPT [157]

### Multiplicity of solutions

Some optimal control problems can have non-unique solutions, i.e. different control trajectories corresponding to the same cost function value. This is an often overlooked issue that, although mathematically correct, might have unforseen consequences when trying to explain or predict biological behaviour.

In a strict mathematical sense, such an issue is related with the system’s structure. For example, a dynamic system with highly correlated controls that can compensate their impact on the cost function might result in multiple solutions with different controls, yet identical cost function. In particular, in optimal control theory, a very well known issue is the existence of a singular arc. In a singular arc, the controls appear linearly in the system’s Hamiltonian and therefore can not be uniquely identified.

However, from a computational point of view, practical multiplicity of solutions can also be defined. When solving a non-linear optimal control problem one can obtain solutions of different control trajectories that correspond to very similar or, even, tolerance-wise same cost function values. Such a finding implies that the cost function is practically insensitive to at least one of the controls in at least a certain interval of the time horizon.

In this work we investigate the non-uniqueness of solutions from a practical and numerical point of view considering the ensemble of solutions found in the close vicinity of the global solution (typically using a threshold of 0.5%). We also illustrate how to distill valuable information about the role of constraints using the Lagrange multipliers.

## Results and discussion

Here we illustrate and evaluate our new two-phase approach, `AMIGO2_DO+ICLOCS`, by solving three challenging case studies:

- LPN3B: a three step linear pathway with transition time and accumulation of intermediates as objectives
- SC: the central carbon metabolism of *Saccharomyces cerevisiae* during diauxic shift, which was originally formulated as a single-objective problem in [72], and subsequently considered as a multi-objective problem by de Hijas-Liste et al[86]
- BSUB: the central carbon metabolism in *Bacillus subtilis* during a nutrient shift, extended and adapted from a single-objective formulation described in Töpfer[158].

For each case study, we present and discuss the results obtained with our combination strategy, `AMIGO2_DO+ICLOCS`. In order to illustrate its advantages, we also provide a comparison with the separate solvers, `AMIGO2_DO` and `msICLOCS` (that is, a multistart of `ICLOCS`, which was only evaluated in this way because single runs would likely result in local solutions). Software source code with the implementations of the above case studies for these solvers is available at https://doi.org/10.5281/zenodo.3793620.

### Three-step linear pathway (LPN3B)

This case study is a relatively simple yet non-trivial problem presented in [86], extending the formulation of Bartl et al[81]. The problem represents a simple linear pathway of enzymatic reactions, described by mass action kinetic, where a substrate *S*_1_ is converted to a final product *S*_4_ in three steps (two intermediate metabolites *S*_2−3_). Here we consider a multiobjective formulation with two objectives: minimization of transition time and minimization of the intermediates accumulation. The transition time is taken as the necessary time for a given amount of product to be reached. Note that the substrate is considered to be in abundance, so its concentration remains constant. The network representation is given in Figure 3. The mathematical formulation of the multiobjective optimal control problem is:

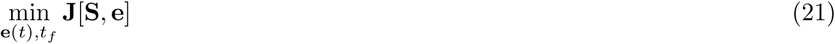

**Figure 3:**
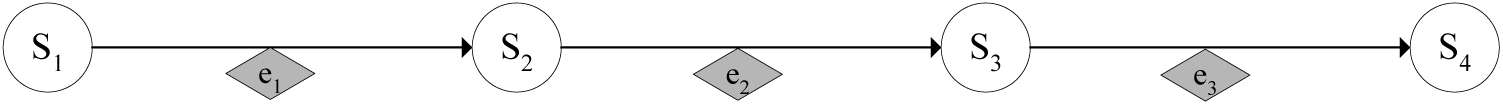
Network representation for the LPN3B case study.

Where:

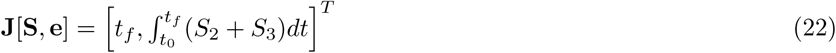

Subject to the system dynamics:

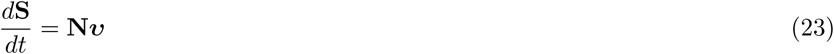

Where the states’ vector is:

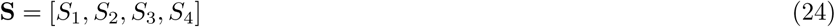

While the stoichiometric matrix **N** is:

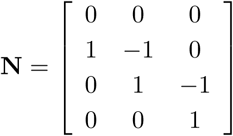

The kinetics are described by:

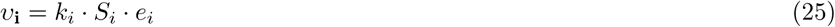

With the following end-point constraint:

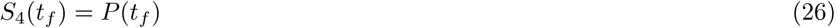

and path constraint:

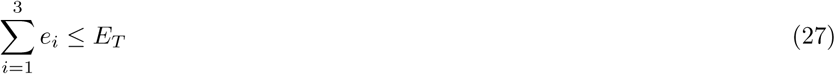

with: *E*_*T*_ = 1 M, *S*_1_(*t*_0_) = 1 M, *S*_*i*_(*t*_0_) = 0 for *i* = 2, 3, 4, *k*_1−3_ = 1 and *P* (*t*_*f*_) = 0.9 M. The total amount of enzyme concentrations is bounded by the path constraint (27), representing the assumption that cells can only allocate a limited amount of enzymes to a given pathway, and also supported by the concentration limitations arising from molecular crowding [72, 81]. The end-point equality constraint (26) assigns the product target value that defines the transition time. The path constraint (27), representing the biological limitation for total enzyme concentration, makes this problem quite difficult to solve, and in fact several existing optimal control software packages tried in [86] failed to converge, or converged to local solutions.

Here we computed the Pareto front of this multi-objective optimal control with the three different numerical strategies described above. The results are summarized in the Pareto front presented in Figure 4. The optimal controls corresponding to the extreme points (A and C) and the knee point (B) of the Pareto front in Figure 4 are shown in Figure 5. This comparison allows the visualization of the impact on the optimal control policies of the different trade-offs between the criteria considered. The path constraint (27) was found to be active throughout the whole time-horizon for all the solutions in the Pareto set.

**Figure 4:**
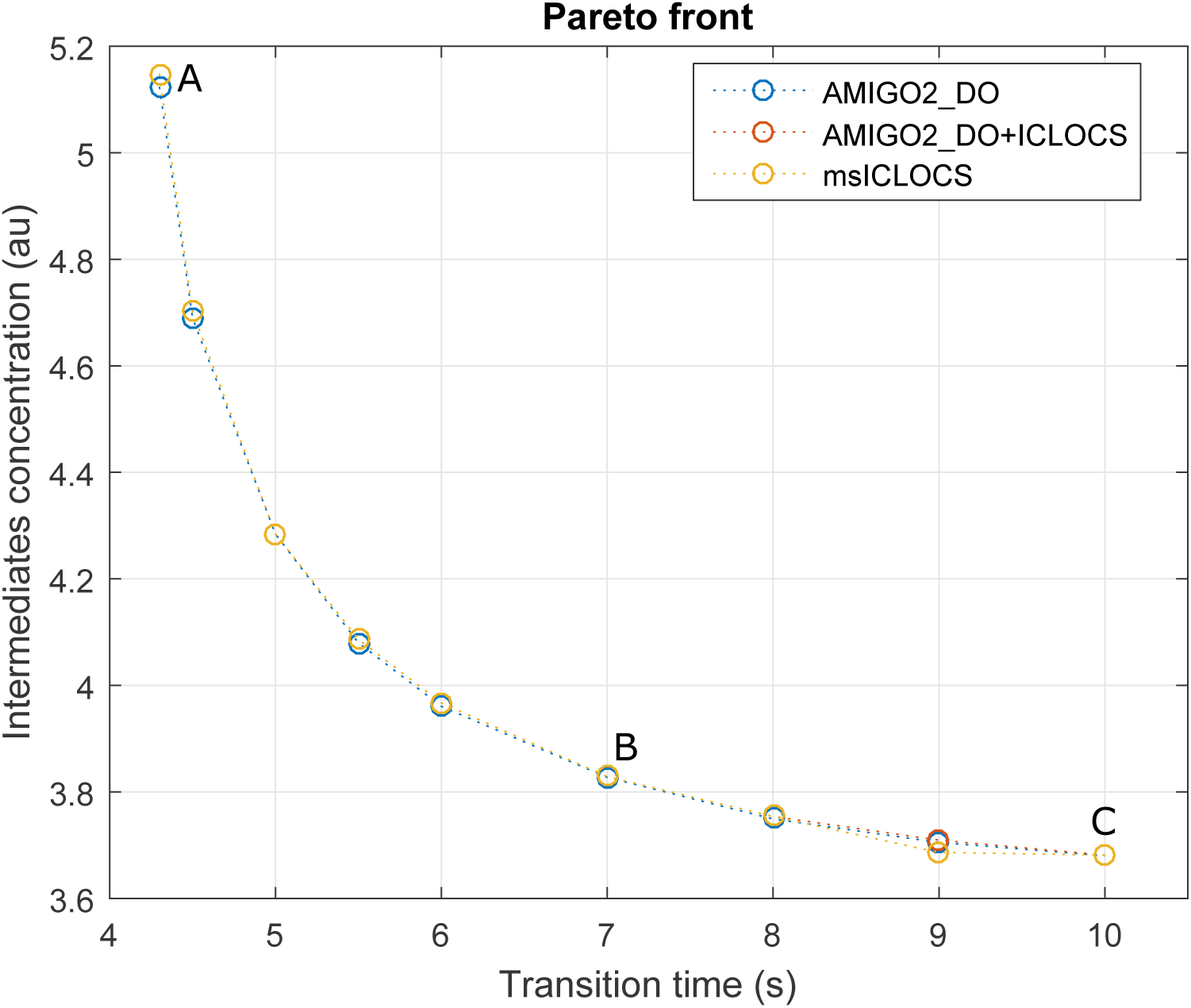
Case study LPN3B: Pareto front computed with three different approaches.

**Figure 5:**
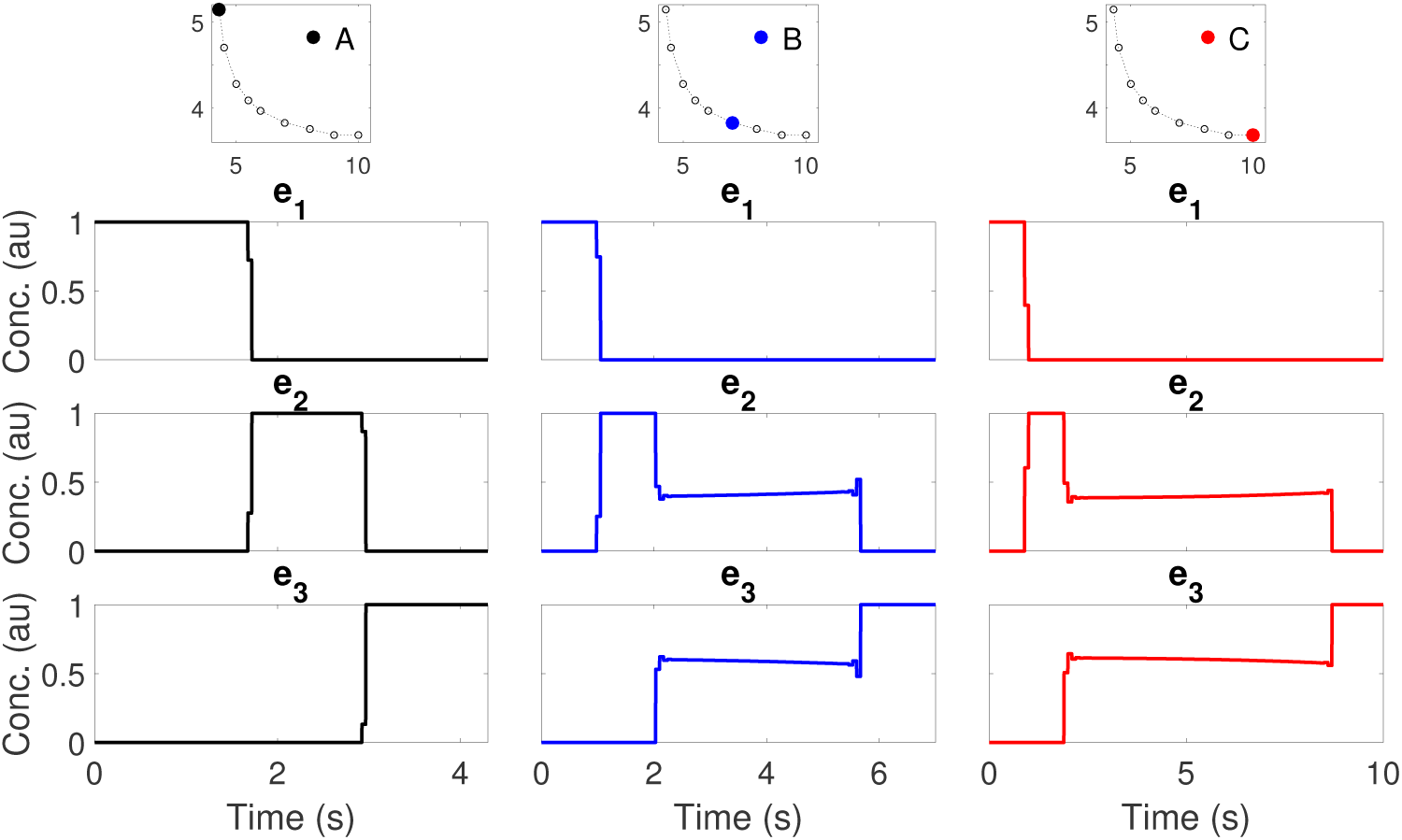
Case study LPN3B: Optimal controls for different points (A, B and C) on the Pareto front. The solutions presented here were computed with a multistart of `ICLOCS (msICLOCS)`.

These results are in close agreement with the best Pareto set obtained in [86]. However it should be noted that, while several of the the solvers used in [86] resulted in bad local solutions, here the three strategies considered (`AMIGO2_DO+ICLOCS, AMIGO2_DO` and `msICLOCS`) converged to essentially the same Pareto set of near-globally optimal solutions.

We further support this claim by providing additional detailed comparisons of the optimal controls and optimal state trajectories computed by the three approaches for several points in the Pareto front (see Additional File 1:LPN3B). It is worth mentioning that the solution of the problem is very sensitive to the switching times in the optimal controls. The different strategies considered managed to find the optimal switchng times either by using a control discretization with elements of varying size (`AMIGO2_DO`) or by using a very refined control mesh (`msICLOCS`).

In Additional File 1:LPN3B, we also provide a detailed computational comparison of the three strategies. From these results, we conclude that the fine-tuned `msICLOCS` was the best numerical strategy for this case study, providing very good results with very modest computational costs (around 20 s on a standard PC). Fine-tuning `msICLOCS` was critical, providing great benefits with respect to the default settings: it accelerated convergence by at least one order of magnitude, and it eliminated convergence to local solutions. The `AMIGO2_DO` solver and the combination strategy `AMIGO2_DO+ICLOCS` also managed to provide very good final solutions, but with a larger computational cost. The convergence curves of the three approaches are also given in the additional file, showing the superiority of `msICLOCS` for this problem.

Finally, we checked the possible existence of near-optimal solutions with significantly different control profiles. This was done by performing an analysis of solution multiplicity and sensitivity of the cost function with respect to control profiles from 100 runs of `AMIGO2_DO+ICLOCS`. Details are given in Additional File 1:LPN3B, showing how the optimal controls for those near-optimal solutions differ significantly from the global one, loosing the clear sequential wave-like behavior of the enzyme activation. These results indicate that, in order to obtain a clear cut characterization of the optimal enzyme activation, one needs to enforce convergence to the close vicinity of the global solution for this type of problems.

### Central metabolism of *Saccharomyces cerevisiae* during diauxic shift (SC)

This problem, formulated in [86], considers a simplified network of the central carbon metabolism of *Saccharomyces cerevisiae*, accounting for the pathways of upper and lower glycolysis, TCA cycle and respiratory chain, as depicted in Figure 6. It is modeled as a mass-action kinetic metabolic model representing glucose, triose phosphates, pyruvate and ethanol by states *X*_1−4_.

**Figure 6:**
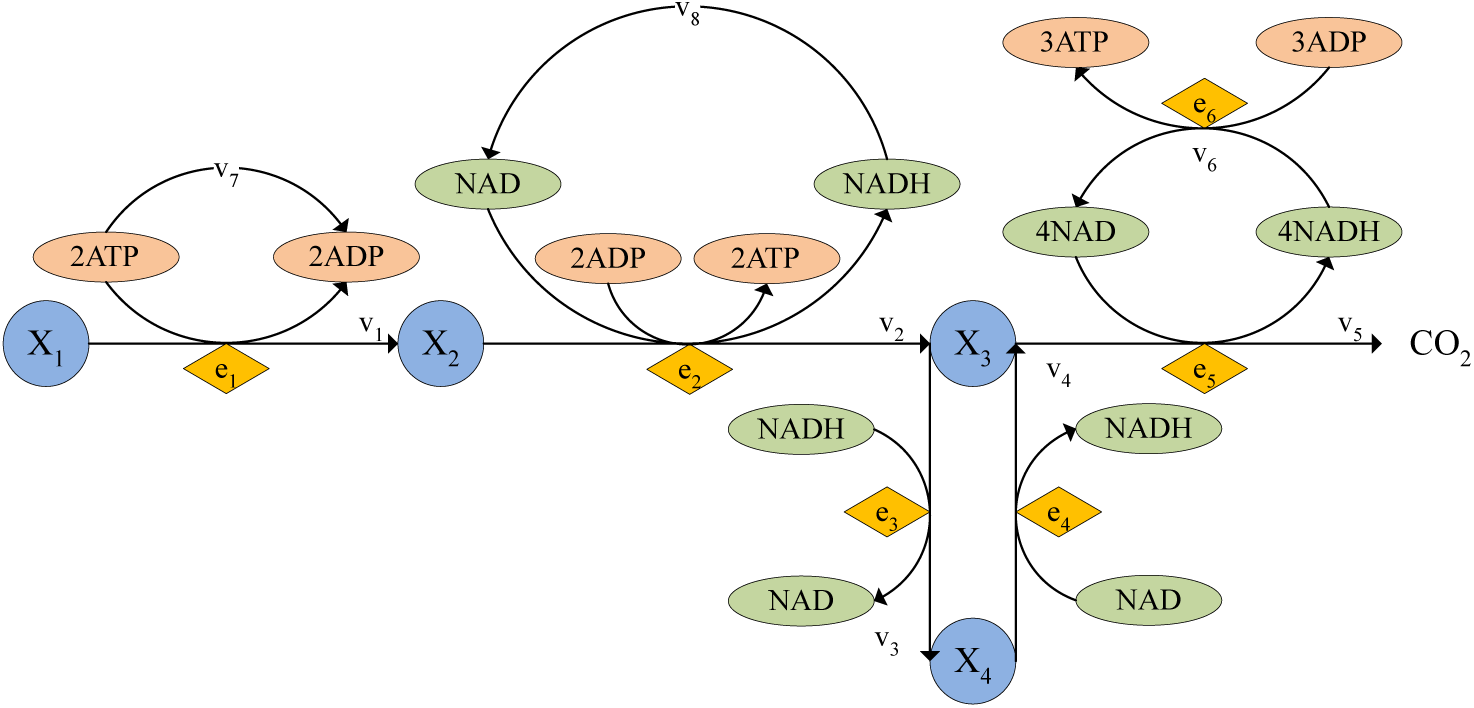
Network representation of the SC case study

The scenario examined is a diauxic shift under glucose depletion, where the main metabolic route changes dynamically through enzyme activation from glycolysis to aerobic utilization of ethanol deposits. This strategy of the yeast cell can be interpreted as extending its survival time as much as possible using ethanol as an alternative substrate. The survival of the cell is modeled taking into account critical lower bound values for NADH and ATP concentrations, implemented as path contraints, Eqns. ((32) - (33)), in the optimization formulation.

If we formulate an optimal control problem for this scenario with a single objective, the maximization of the survival time, as done in [86], the corresponding optimal control involves an unrealistically large amount of enzymes. Therefore, it makes more sense to formulate a multicriteria optimal control problem with two objectives: maximization of the survival and minimization of the protein investment (enzyme synthesis effort). In this way, we obtain a good range of the cost/benefit trade-offs which can be subsequently used to explain the observed biological behavior.

The mathematical formulation is given as follows:

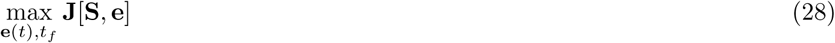

Where:

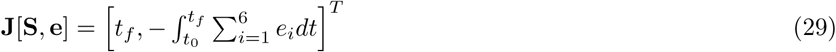

Subject to:

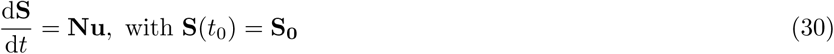

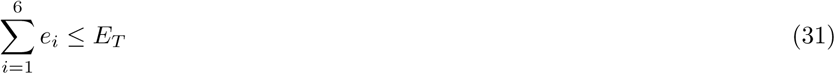

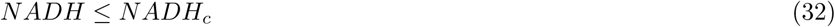

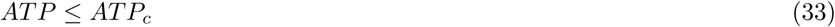

Where the states’ vector is:

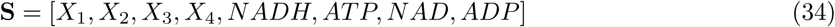

The stoichiometric matrix along with the reactions’ kinetics corresponds to (35) and (36) respectively and are given by:

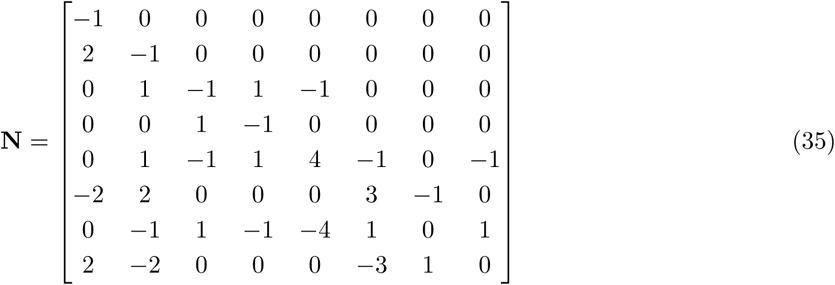

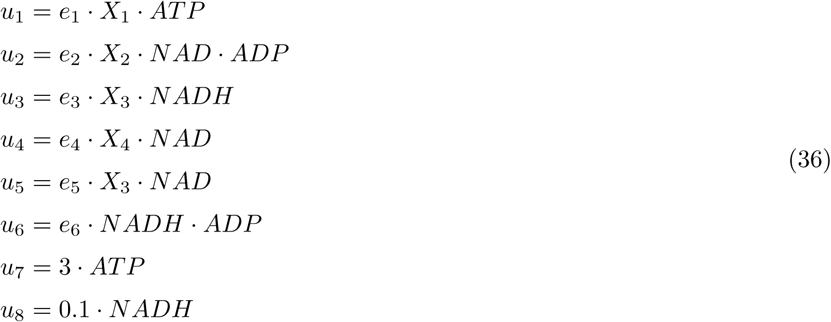

With the following initial values:

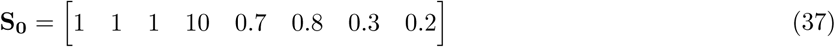

All concentrations are arbitrary and the time is given in hours. A path constraint on the total amount of enzymes at any time is implemented in Eqn (31) where *E*_*T*_ is fixed at 11.5. Eqns (32) and (33) are path constraints representing the critical values of NADH and ATP for cell survival.

We solved the above multicriteria optimal control problem with the three approaches, obtaining essentially the same Pareto front with all of them, as depicted in Figure 7. More detailed results are given in Additional File 2:SC. These results are also in close agreement with those reported in [86], not only in terms of the Pareto front but also in terms of a qualitative comparison with available transcriptomics data (see Additional File 2:SC), illustrating how optimal control can explaim the observed biological behavior.

**Figure 7:**
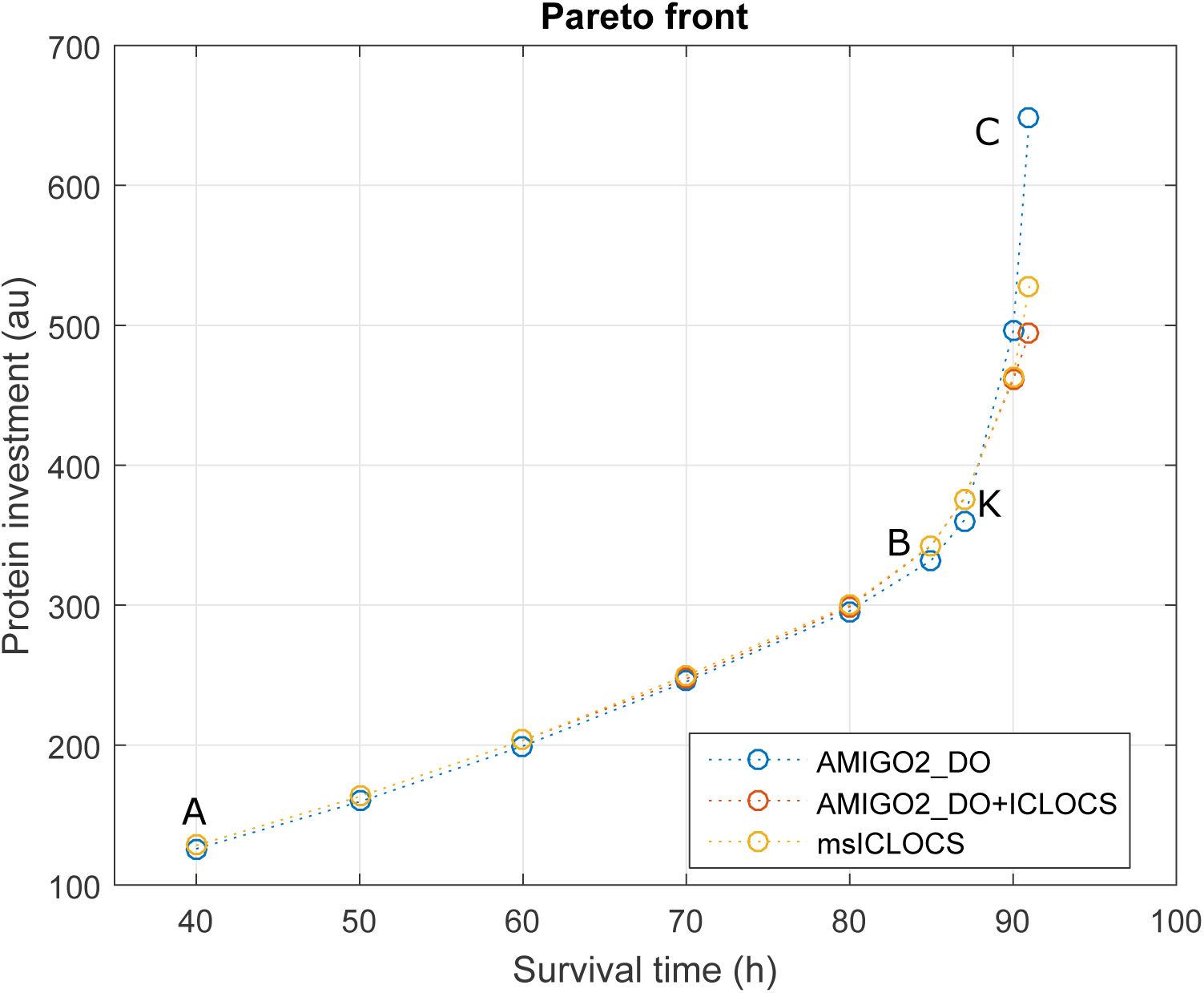
Case study SC: Pareto front computed with the three different approaches. Point A and C correspond to the extreme points of the three approaches while B and K are in the vicinity of the knee point.

In contrast with the previous case study, we found that `AMIGO2_DO+ICLOCS` was the best numerical strategy for this problem, arriving to the best solution in only 5 min of computation using a standard PC, and avoiding convergence to local solutions. Interestingly, `msICLOCS` converged to local solutions or crashed rather frequently, even with the use of fine-tuning. In Figure 8 we present a comparison of the frequency histograms of the solutions reached by these two strategies for one of the points in the Pareto. Further computational detail are given in Additional File 2:SC.

**Figure 8:**
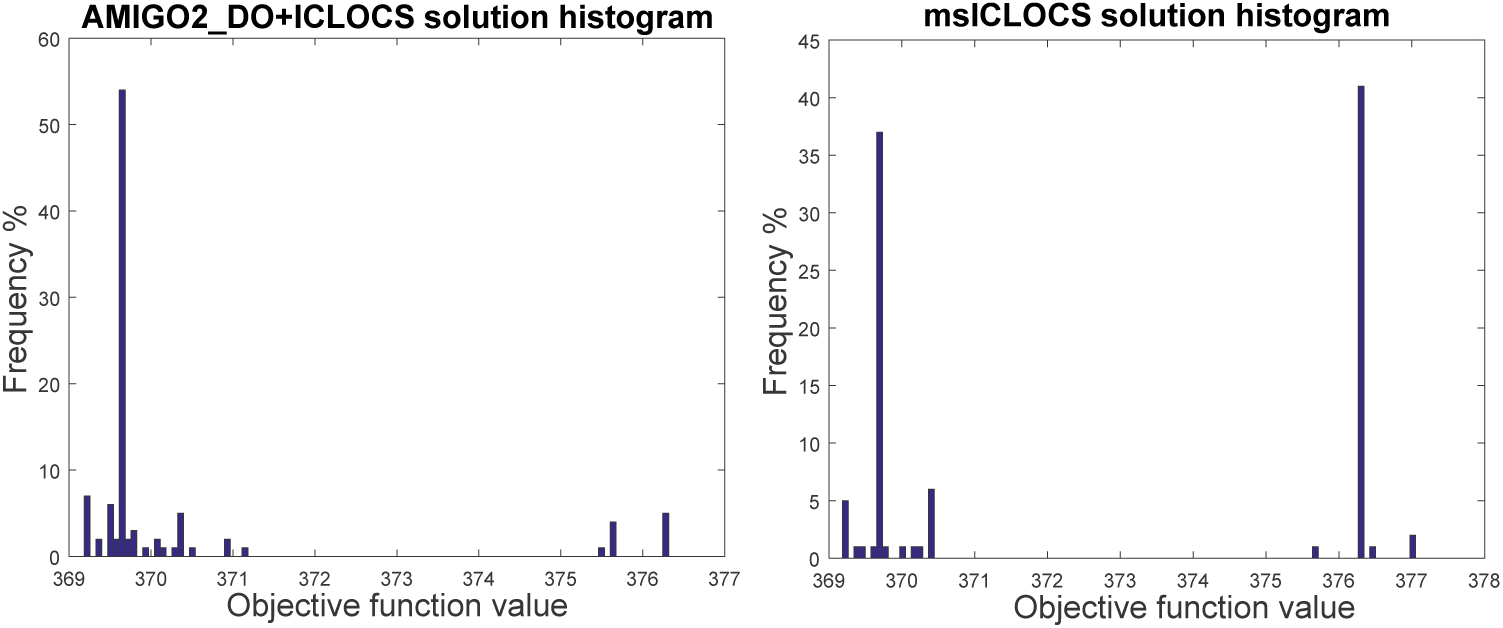
Case study SC: comparison of the solution histograms corresponding to the strategies `AMIGO2_DO+ICLOCS` (on the left) and `msICLOCS` (on the right), for point K on the Pareto front shown in Figure 7.

We also performed an analysis of solution multiplicity and sensitivity of the cost function with respect to the controls by analyzing 100 runs of the hybrid approach. Considering the solutions within 0.5% of the best, most of the controls (time-dependent enzyme concentrations) present a considerable spread (details can be found in Additional File 2:SC). In the case of the states, the spread is also significant, although in most cases the qualitative behaviour is retained. In any case, these results indicate the need of using rather tight convergence criteria to ensure a consistent quantitative computation of the optimal controld and state dynamics involved.

This case study includes two path constraints representing the critical values of ATPc and NADHc which are needed to ensure the survival of the cell. We performed a numerical analysis to evaluate the impact of these path constraints on the optimal solution. In Figure 9 we present a brief overview of the results corresponding to the analysis of the critical ATP value. Subfigure I shows the different Pareto fronts obtained for several values of this constraint, illustrating its very significant impact. In subfigure II, the corresponding state dynamics can be found. It can be seen that although the survival times changed considerable, most of the state dynamics retained the same qualitative shape, including the initial ethanol production phase. Interestingly, although the ATP values are almost always active at the critical value, confirming its high sensitivity, this is not always the case for the NADH dynamics. This was further studied by performing a similar analysis for the NADHc values, obtaining results which confirmed its less significant impact on the optimal solutions (details in Additional File 2:SC).

**Figure 9:**
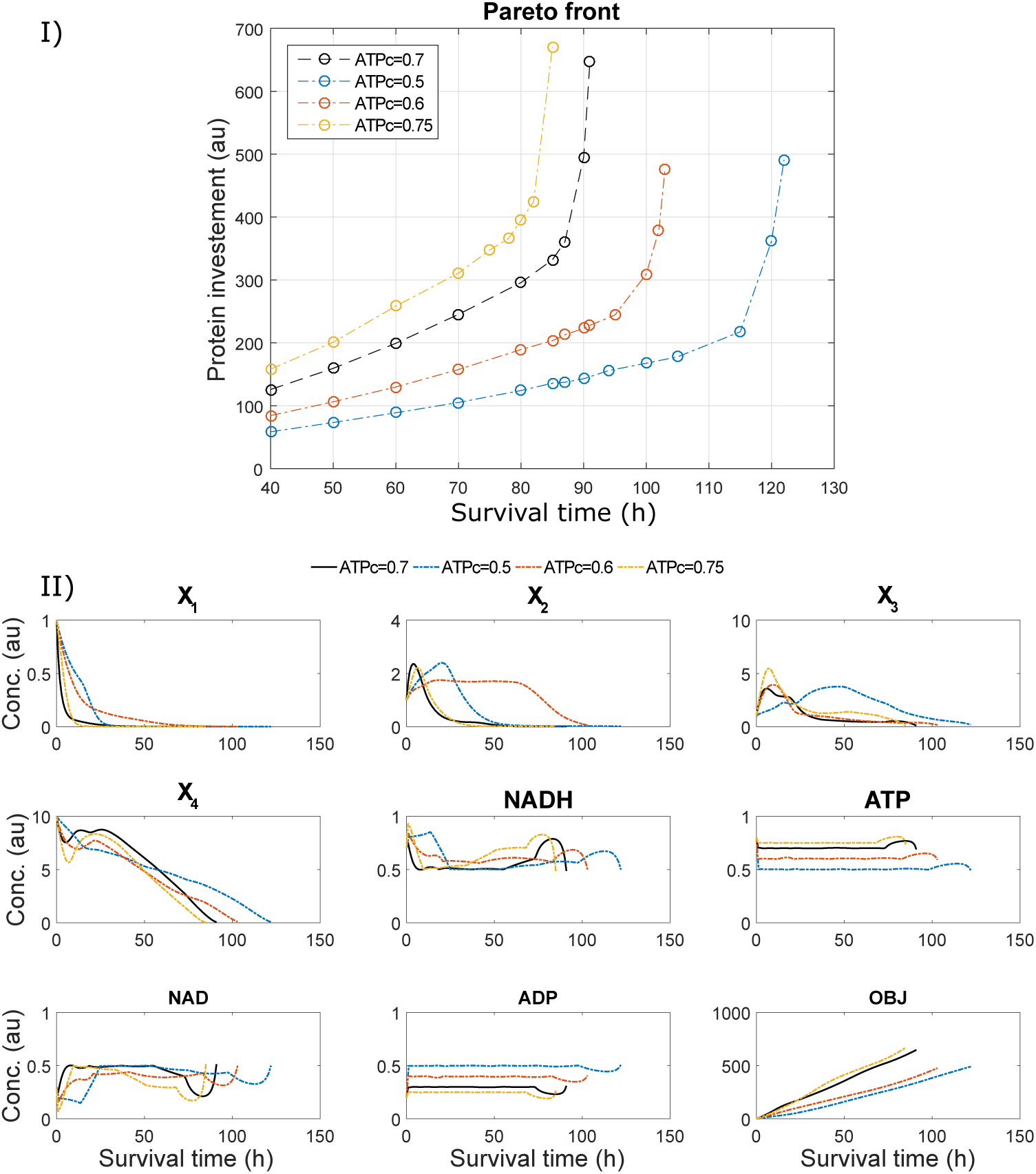
Case study SC: an illustration of the impact of the ATP critical value path constraint on the Pareto front previously shown in Figure 7 is presented in subfigure I. Subfigure II shows the impact of this path constraint on the optimal state trajectories corresponding to the extreme point of maximum *t*_*f*_.

Next we performed an analysis of the adjoint variables (in practice, the Lagrange multipliers) to get further insights regarding the local sensitivity of the performance index with respect to the constraints. That is, the adjoint variables for specific constraints or bounds indicate the local sensitivity of the solution with respect to their incremental change. Detailed results for the Lagrange multipliers are presented in the Additional file 2:SC, confirming large values for ATPc, and reletively important ones for NADHc, in agreement with the results shown in Figure 9 and discussed above. It should be noted that this concept becomes also interesting to evaluate the role of uncertainty and/or variability of the constraints (both prominent in biological modelling). For example, are the dynamics predicted by optimal control robust with respect to variability in critical constaints? Or, which are the most important constraints in terms of impact on the optimal cost?

### Central carbon metabolism in *Bacillus subtilis* during a change of substrates in the environment (BSUB)

This case study is based on a simplified kinetic model of the central carbon metabolism of *B.subtilis* taken from [158]. The model considers important pathways such as upper and lower glycolysis, TCA cycle, glyconeogenesis, overflow metabolism and biomass production. The network representation is given in Figure 10.

**Figure 10:**
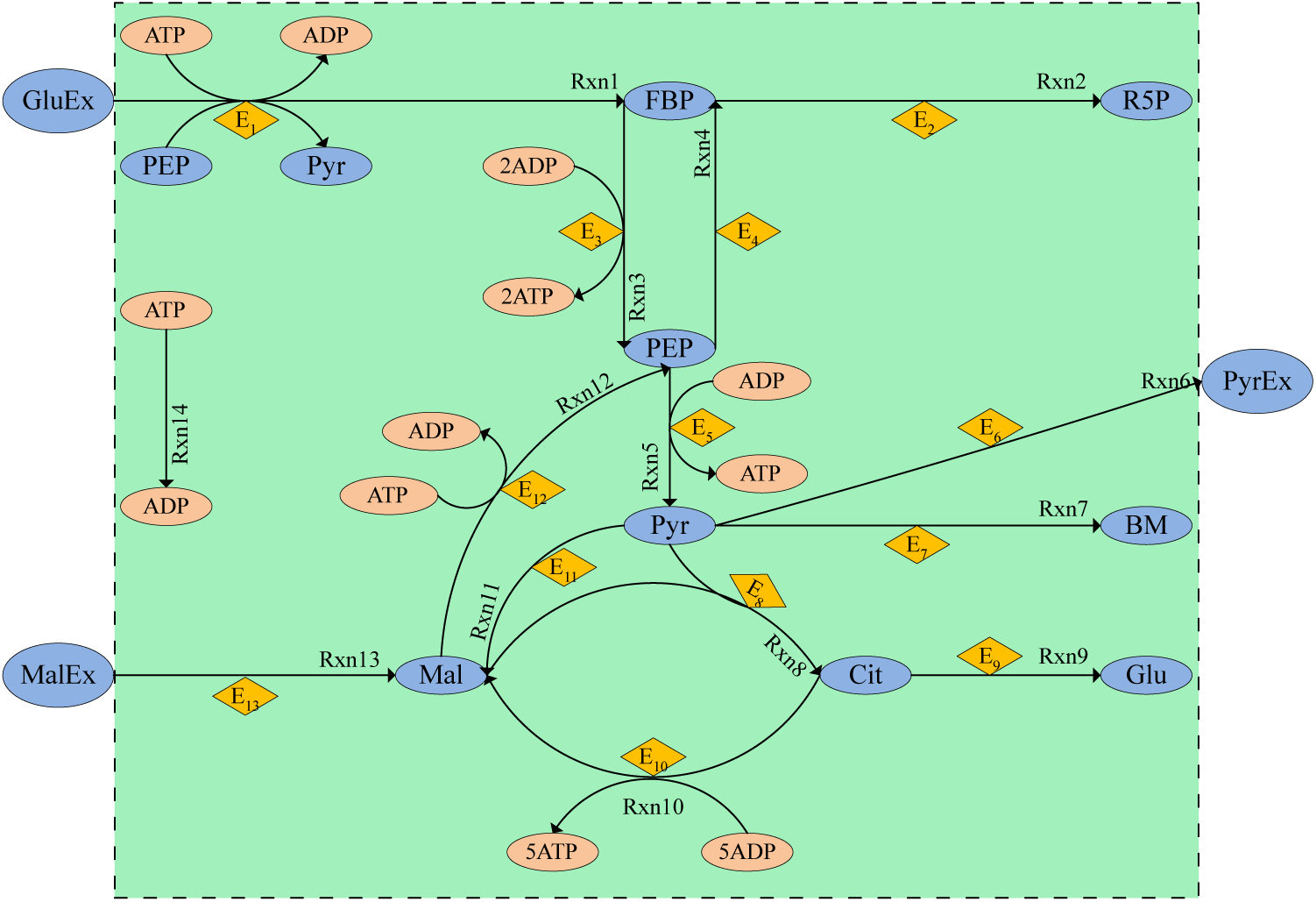
Network representation of case study BSUB

Here, the objectives considered are the maximization of the overall ATP production and the minimization of the protein investment (enzyme synthesis). We aim to find the cost/benefit trade-offs, where the benefit is computed as the integral of ATP levels along the time horizon, and the cost is given by the integral of the total enzyme concentrations for the whole time horizon.

The corresponding multicriteria optimal control formulation is as follows:

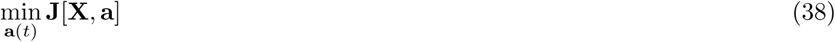

Where:

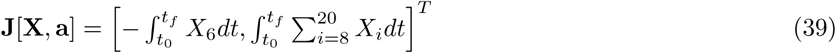

Subject to:

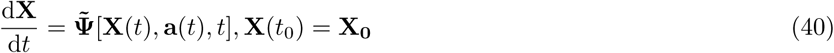

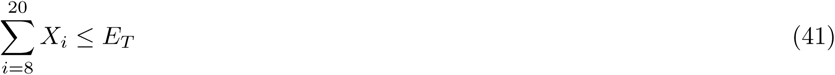

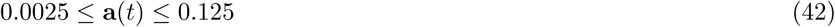

Where the right hand side of the differential equations corresponds to:

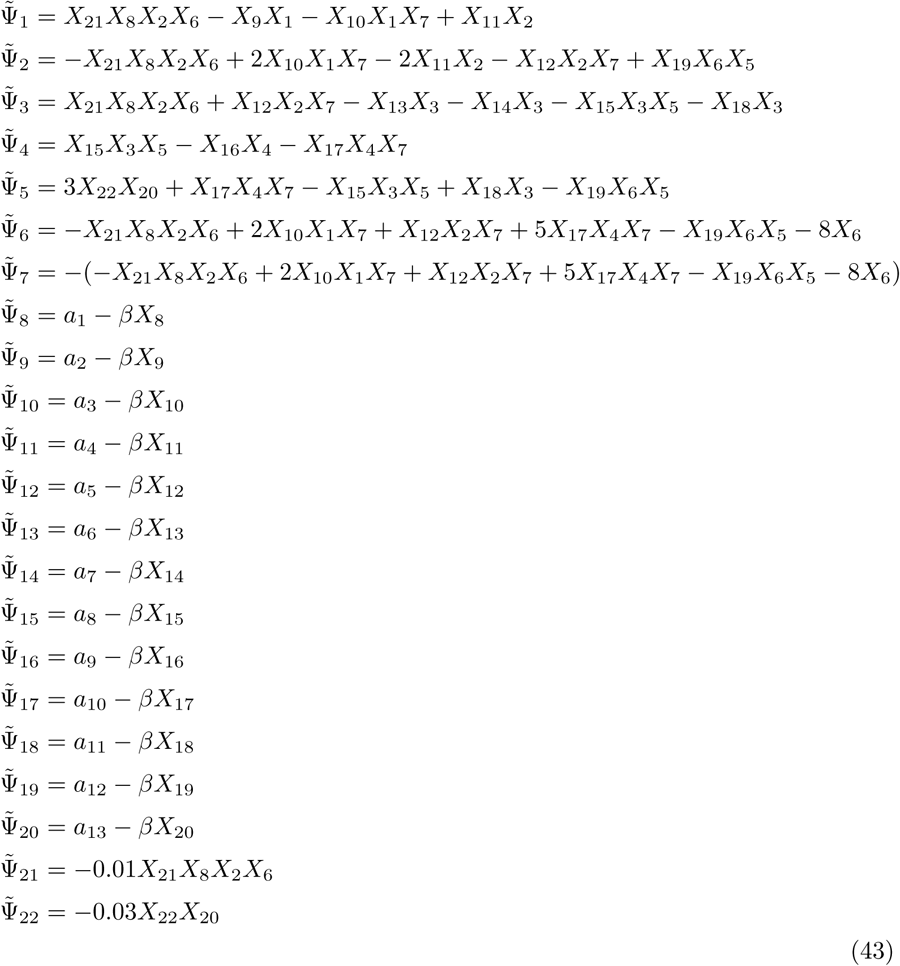

With the vector of states corresponding to the following metabolites, enzymes and substrates as presented in the network representation:

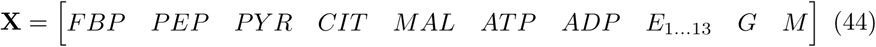

The terms 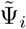 are the right hand sides of the kinetics of the different reactions *i* = 1..22. Reaction 1 coresponds to the glucose uptake coupled with the conversion of phospho-enol-pyruvate (PEP) to pyruvate (Pyr). Reactions 3, 4 and 5 represent glycolysis, yielding 4 ATP per FBP conversion to 2 Pyr. Reactions 2, 6 and 7 correspond to gluconeogenesis, excretion of pyruvate through fermentation and biomass production respectively. Reactions 8 and 10 describe the respiratory pathway of the TCA cycle yielding 5 ATP per Pyr. Reaction 9 describes glutamate (Glu) production to be used in amino acid synthesis. Malate uptake from the environment is described by reaction 13. Malate can be also created from Pyr (reaction 11) and then converted to PEP (reaction 12). Enzyme dynamics are taken into account using a simple linear dynamic expression (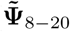 in (43)) with a rate constant of degradation (*β* = 0.25) and production rates (**a**) as the decision variables (controls) of the optimal control formulation. The total enzyme capacity is implemented in Eqn.(41) with a bound of 6.5 (arbitrary units).

In this case study we are considering two dynamic scenarios. In the first scenario (named G-M) we consider a culture growing in an environment where glucose is the only substrate. At a certain time-point we add malate and we observe the dynamic behavior. The second scenario (M-G) is vice-versa. That is, the culture is initially growing on malate, before instantly adding glucose as a substrate and then observe the dynamics. The first phase of both scenarios is computed starting from the same initial values and reaching a steady state for the consumption of the substrate available.

The initial conditions for scenario G-M are:

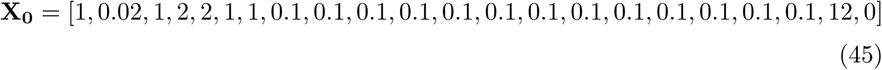

The initial conditions for scenario M-G are:

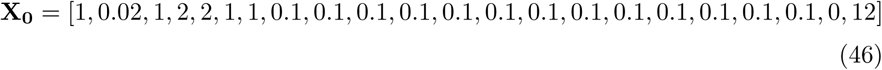

Considering both scenarios, we solved the corresponding multicriteria optimal control problems using the three strategies, the hybrid `AMIGO2_DO+ICLOCS` and the separate solvers, `AMIGO2_DO` and `msICLOCS`. All of them produced almost identical Pareto fronts for each scenario. Figure 11 (I) shows the Pareto front corresponding to scenario G-M. The corresponding optimal state trajectories for point B in the Pareto front are presented in Figure 11 (II), illustrating how the three strategies converged to essentially the same near-global optimal solution. Similar results were found for scenario M-G. Detailed results can be found in Additional File 3:Bsub.

**Figure 11:**
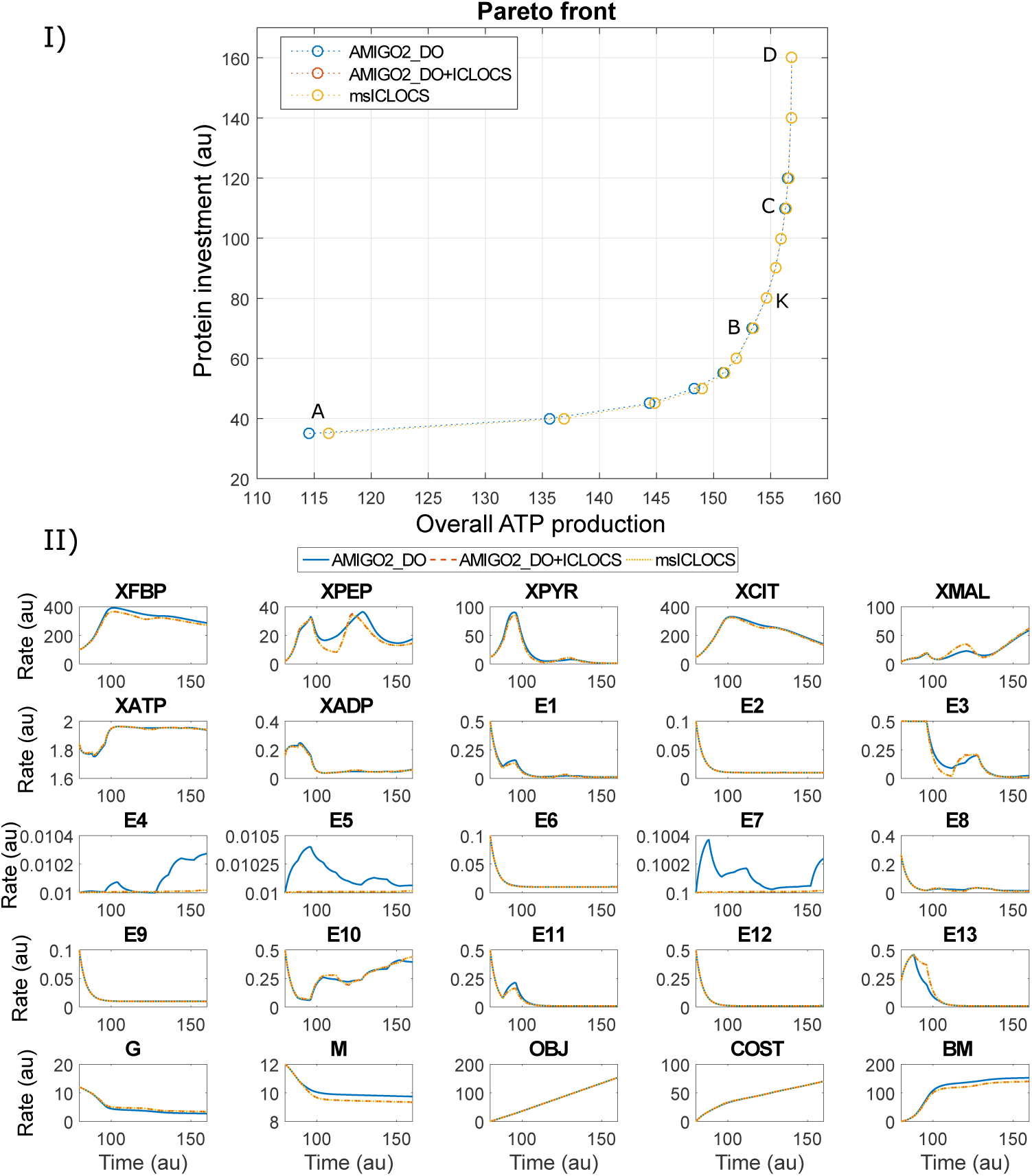
Case study BSUB: subfigure I shows the Pareto front obtained with the three different strategies. Subfigure II compares the corresponding optimal state trajectories for point B of the Pareto (subfigure I).

Even though the three methods arrived to essentially the same multicriteria optimal control Pareto sets, `msICLOCS` was the fastest, converging to very good results in very modest computation times (typically less than 150 s per `ICLOCS` run on a standard PC when fine tuning was used). As in the previous case studies, fine-tuning `ICLOCS` was found to be critical to improve its convergence speed and to avoid convergence to local solutions. In Additional File 3:Bsub we provide detailed results including the effect of fine-tuning on the performance and robustness of `msICLOCS`, and comparisons with the other strategies.

Finally, we performed an analysis possible solution multiplicity and sensitivity of the cost function with respect to the controls. We were not able to identify multiplicity of solutions as shown in the respective figures in Additional File 3:Bsub.

### Overall summary of results

The above results show that the novel combined strategy `AMIGO2_DO+ICLOCS` was the best in terms of robustness, solving these three challenging case studies without any issues, while requiring a very reasonable computational cost for all of them. Although the multi-start strategy `msICLOCS` was faster in two of the problems, it often failed when attempting the solve the SC case study.

The performance of the `AMIGO2_DO+ICLOCS` strategy also compares very favorably with the previous results of the different optimal control methods considered by de Hijas-Liste et al [86] for the LNP3B and SC cases. Our new strategy allows a significant reduction of computation times (more than 50% for the SC case). Further, `AMIGO2_DO+ICLOCS` has also shown good scalability, solving without issues problems such as BSUB, which has many controls that require fine discretizations and which could not be handled by the previous methods considered in [86]. Finally, our approach provides adjoint information that is particularly useful to asses the role of the different constraints in the optimal solutions.

## Conclusions

Here we have considered the exploitation of optimality principles to explain dynamics in biochemical networks in the absence of complete prior knowledge about the regulatory mechanisms. After carefully reviewing the previous literature, we have presented a multicriteria optimal control framework to generate biologically meaningful predictions for rather complex metabolic pathways. This strategy requires, as a starting point, a calibrated dynamic model of the pathway and the set of cost functions to be considered in the multicriteria formulation. Next, the approach uses a Pareto optimality hypothesis to compute the dynamic effects of regulation in a metabolic pathway. Such computation is performed without the need of making any assumption about the pathway regulation.

We have also discussed the many possible challenges and pitfalls involved in the solution of these nonlinear multicriteria optimal control problems. In order to surmount these difficulties, here we have considered several numerical strategies with the aim of finding a method suitable to handle realistic networks of arbitrary topology. We have carefully evaluated their performance with several challenging case studies considering the central carbon metabolism of *S. cerevisiae* and *B. subtilis*. Our results indicate that the two-phase strategy `AMIGO2_DO+ICLOCS` presented excellent scalability and a good compromise between efficiency and robustness. We also show how this framework allows us to explore the interplay and trade-offs between different biological costs functions and constraints in biological systems.

Our vision is that multicriteria optimal control can play a major role in fully exploiting the explanatory and predictive power of kinetic models in computational systems biology. In future work, we will illustrate its usefulness for the analysis of cell signaling pathways, and for the solution of metabolic engineering problems considering kinetic models.

## Supporting information

Additional file 1 - case LPN3B

Additional file 2 - case SC

Additional file 3 - case BSUB

## Authors’ contributions

JRB conceived, planned and coordinated the study. NT implemented the methods and carried out all the computations. NT and JRB analysed the results. JRB drafted the manuscript. NT contributed to the drafting of the methods and results. All authors read and approved the final manuscript.

## Acknowledgements

Preliminary results from this study were presented at the 6^th^ IFAC Conference on Foundations of Systems Biology in Engineering (FOSBE 2016) in Magdeburg, Germany.

## Tables

### Additional Files

Additional file 1 — Solution details for case study LPN3B

Additional file 2 — Solution details for case study SC

Additional file 3 — Solution details for case study BSUB

