## Additional file 1 - case LPN3B for "Using optimal control to understand complex metabolic pathways"

### Additional file 1: Solution details for case study LPN3B

### Supplementary information

In Figure S1 we show that all three different approaches practically resulted the same Pareto front. In Figures S3-S8 we show in detail that the solutions with the three approaches (AMIGO2_DO, AMIGO2_DO+ICLOCS, msICLOCS) match for all points A,B and C respectively, by comparing both the resulted optimal controls and the optimal state trajectories which correspond to them.

Additionally, we show a computational analysis for one of the points of the Pareto front (point B). First we show the overall performance of msICLOCS in Figures S9-S14 by providing the convergence curves, the solution histograms and histograms of the CPU requirements in this multistart scheme. We provide evidence of the critical role fine-tuning can play in the approach's performance even in a small and non complex case study like this one.

This can be easily visualized by comparing Figures S9,S11,S13 which show the results without any fine-tuning with the respective Figures S10,S12,S14. In both cases the initialization of msICLOCS was performed with the exact same random 100 points. The only difference is that in the second case (Figures S10,S12,S14) the system dynamics (state trajectories) corresponding to these 100 random points were also provided as part of the initial guess. By doing so we observe a significant effect on the solver's effectiveness (decrease in local solutions convergence) and convergence speed (decrease in the CPU requirements). This is of course due to the minimization of the initial guess' constraint violation.

In addition to that we show the performance of the hybrid approach by showing how different set-ups of the sequential AMIGO2_DO+ICLOCS approach perform (Figure S15). Also, the respective Figures S16-S18 illustrate how AMIGO2_DO on its own is outperformed by the other two approaches.

Finally, we performed an analysis of sensitivity of the controls in order to evaluate the uniqueness of the obtained solution and identify any possible problems. To do this we run AMIGO2_DO+ICLOCS 100 times (from random points) for point B of the Pareto front. We intentionally tried to use a hybrid set-up that is too fast, yet no issues were identified. In Figures S19-S25 the results are reported. We did not observe any insensitivity in the controls/states nor any abnormal results in the Lagrange multipliers.

To further visualize the sensitivity of the controls, in Figure S26 we made a comparison between the global optimal solution and two local ones (obtained with msICLOCS) which can also be seen in more detail in Figures S27,S28.

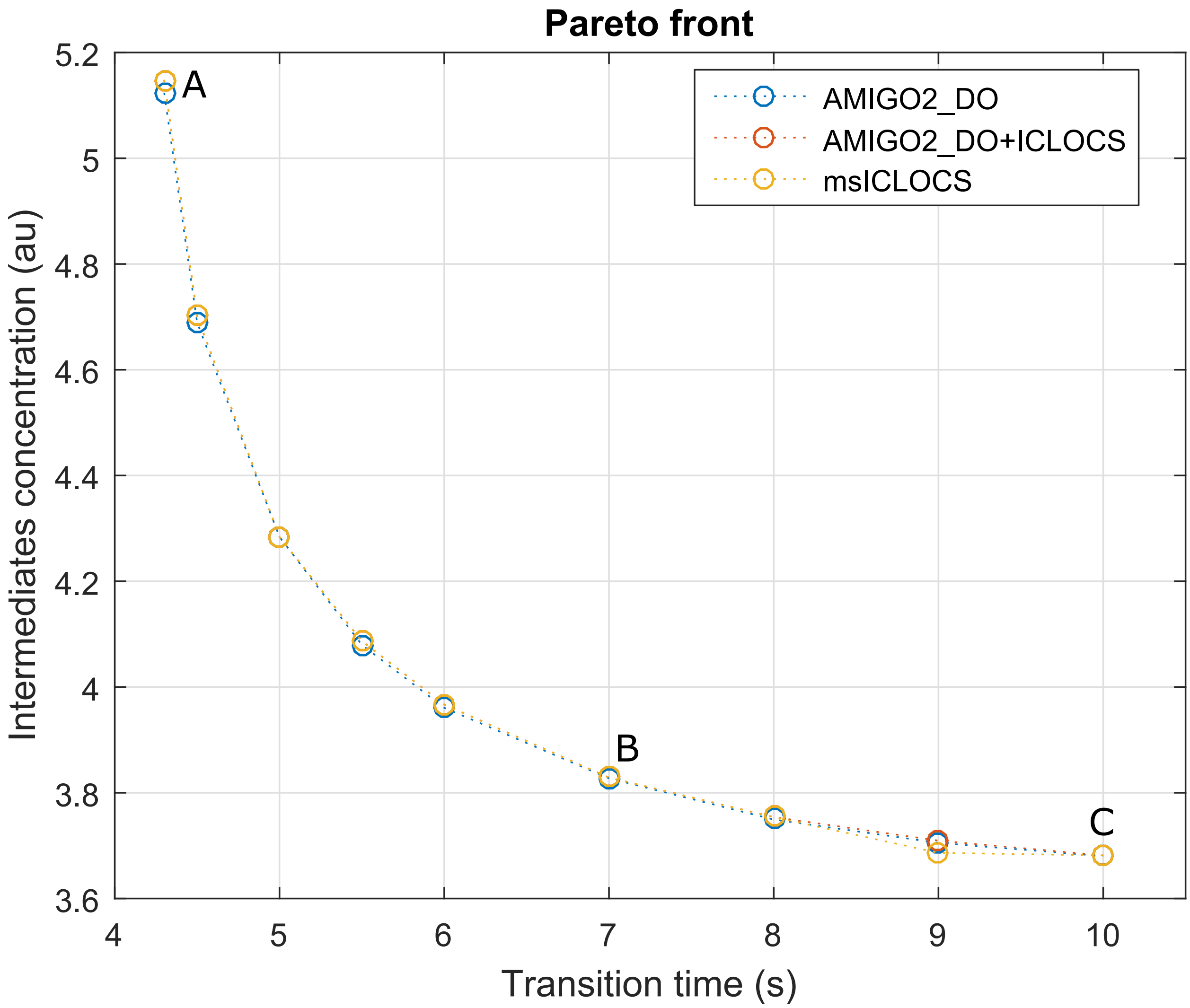


Figure S1: LPN3B: Pareto front computed with the three different approaches.

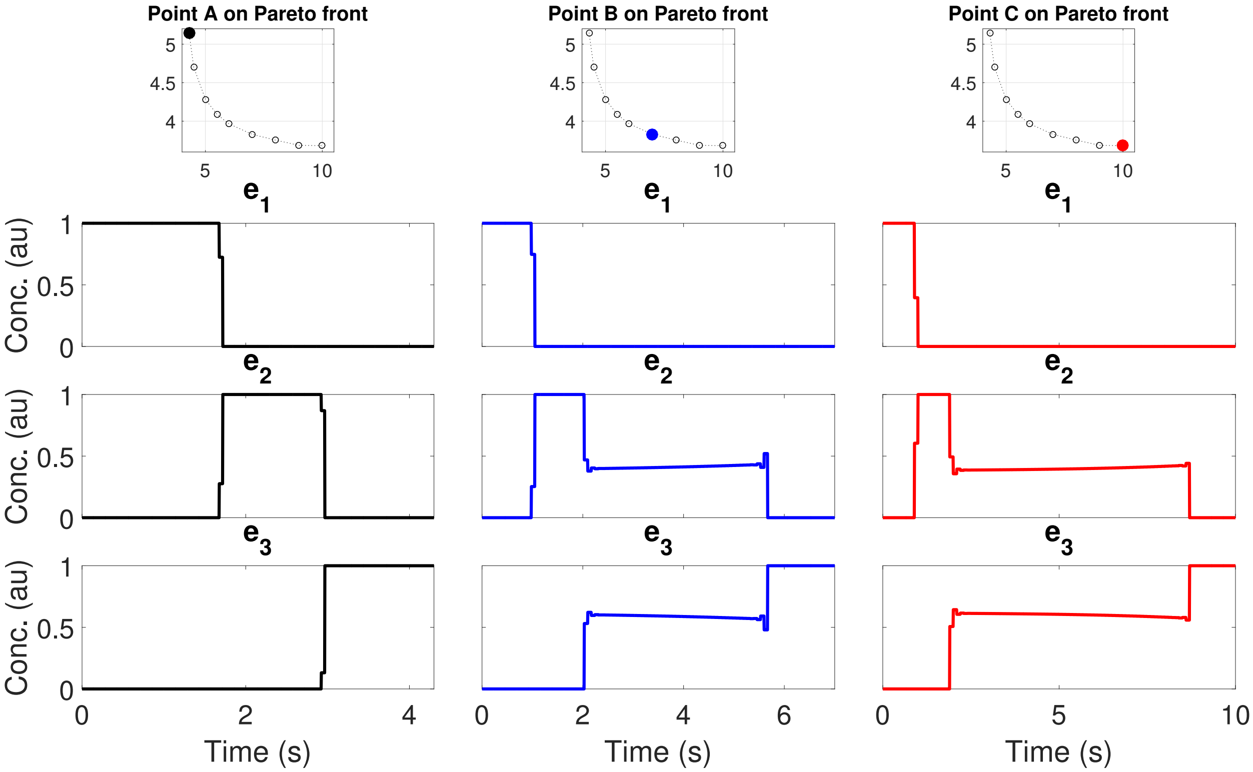


Figure S2: LPN3B: Optimal controls computed with `msICLOCS` for points A, B and C in S1.

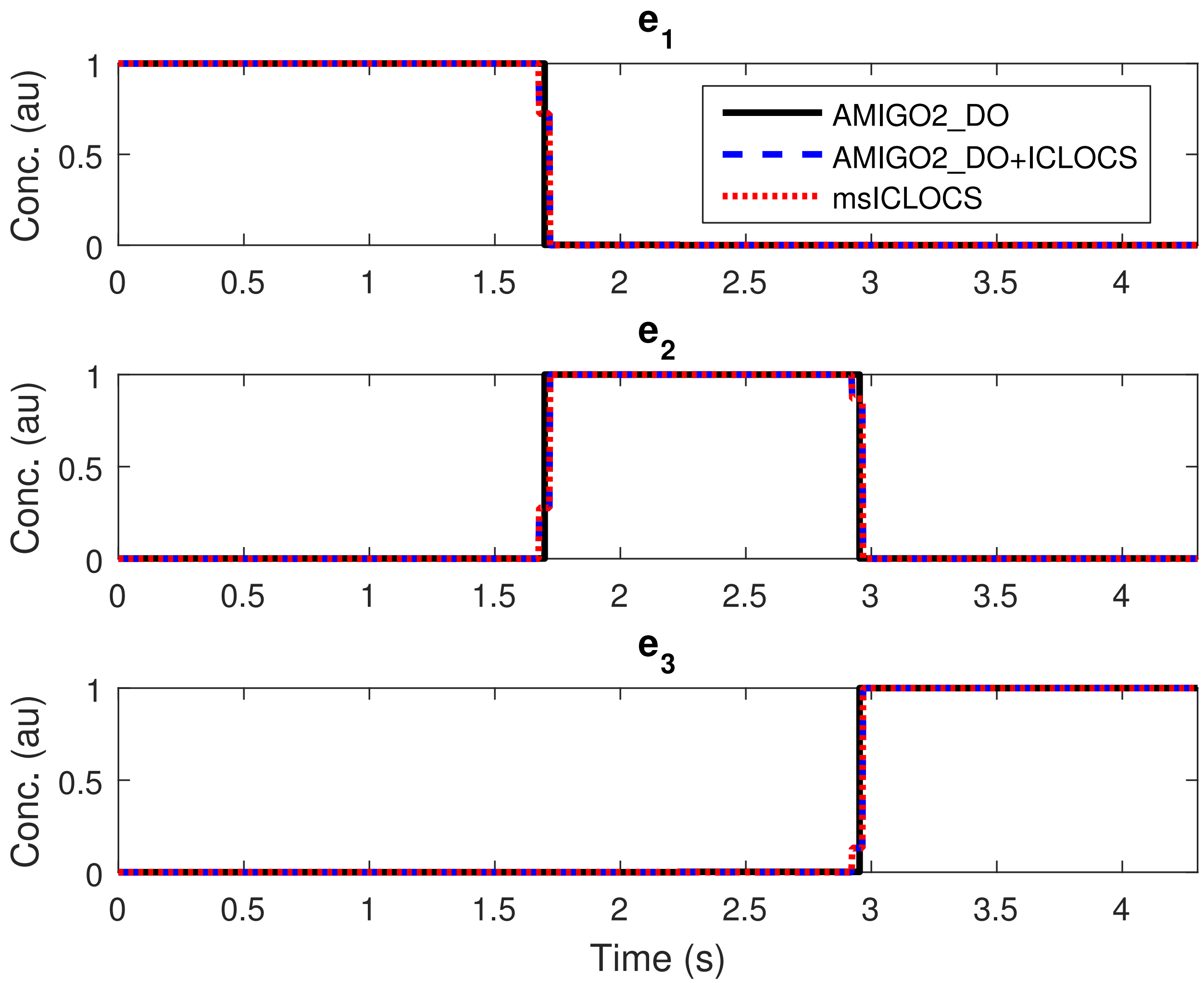


Figure S3: LPN3B: Comparison of the optimal control solutions for point A on the Pareto front. Each method compared is represented by a line of different color and style. Solid black line for `AMIGO2_DO` 4PWCv, dashed blue line for the hybrid of `AMIGO2_DO+ICLOCS` 100PWC and dotted red line for `msICLOCS` 100PWC.

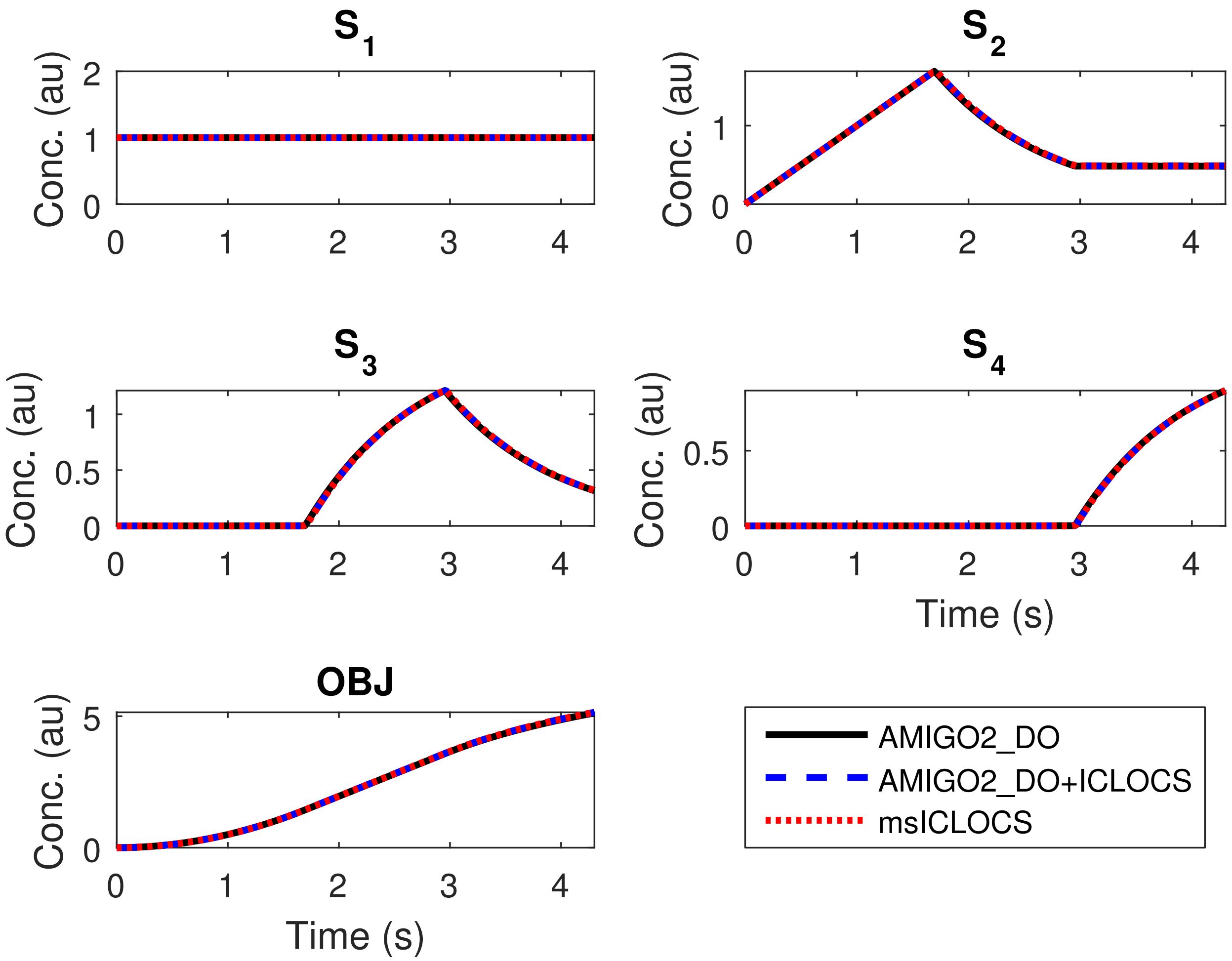


Figure S4: LPN3B: Comparison of the optimal state trajectories for point A on the Pareto front. Each method compared is represented by a line of different color and style. Solid black line for `AMIGO2_DO` 4PWCv, dashed blue line for the hybrid of `AMIGO2_DO+ICLOCS` 100PWC and dotted red line for `msICLOCS` 100PWC.

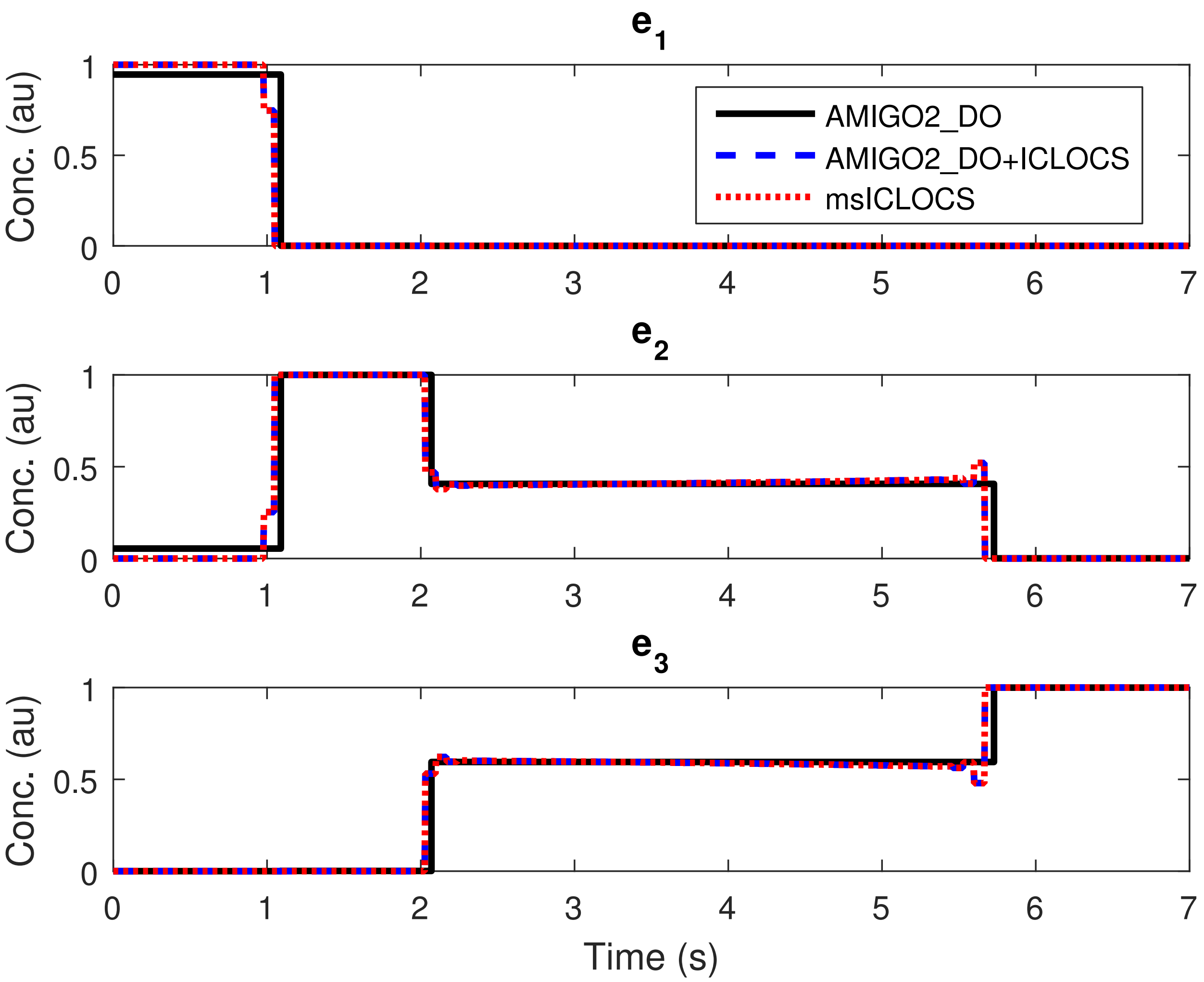


Figure S5: LPN3B: Comparison of the optimal control solutions for point B on the Pareto front. Each method compared is represented by a line of different color and style. Solid black line for AMIGO2_DO 4PWCv, dashed blue line for the hybrid of AMIGO2_DO+ICLOCS 100PWC and dotted red line for msICLOCS 100PWC.

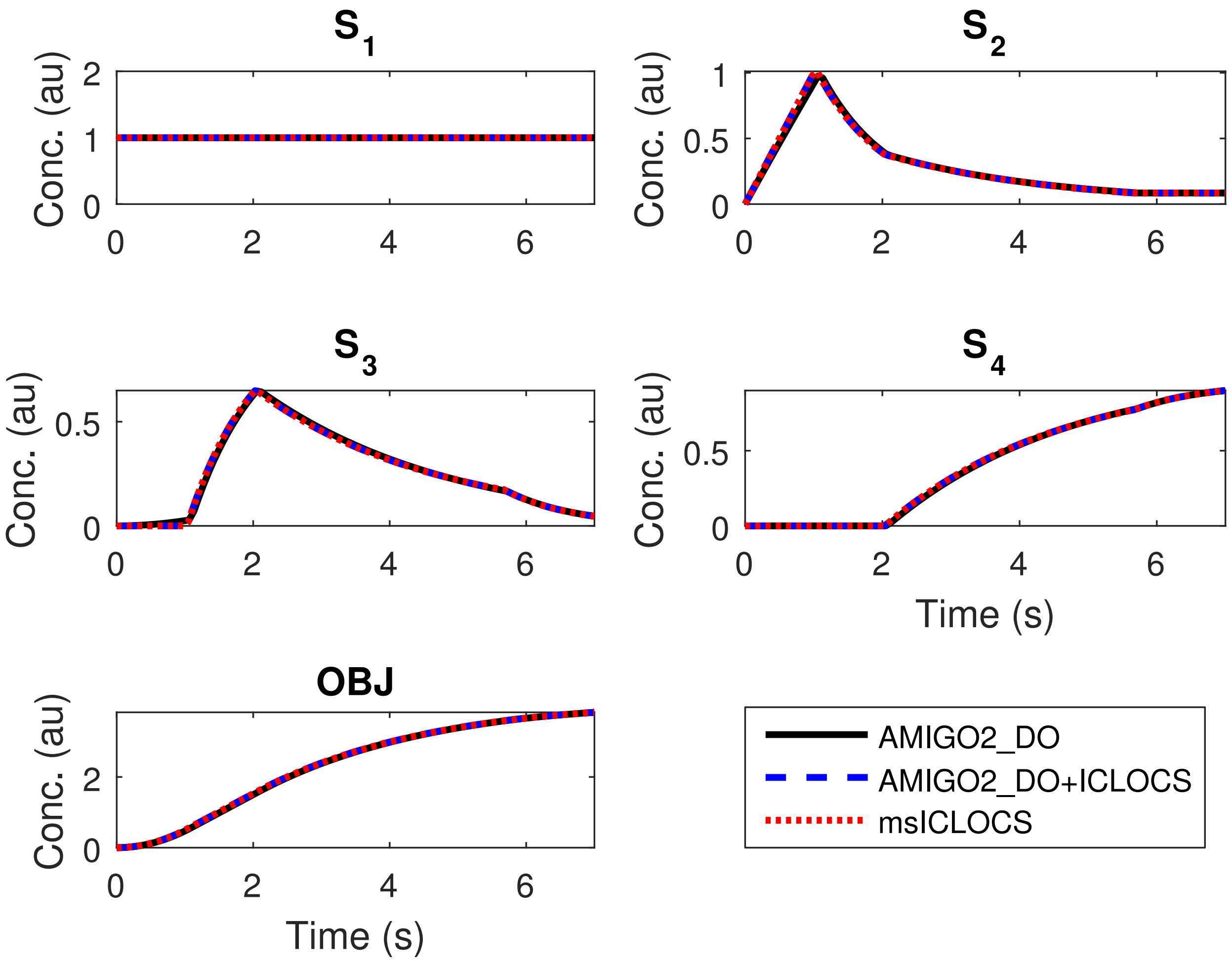


Figure S6: LPN3B: Comparison of the optimal state trajectories for point B on the Pareto front. Each method compared is represented by a line of different color and style. Solid black line for `AMIGO2_DO` 4PWCv, dashed blue line for the hybrid of `AMIGO2_DO+ICLOCS` 100PWC and dotted red line for `msICLOCS` 100PWC.

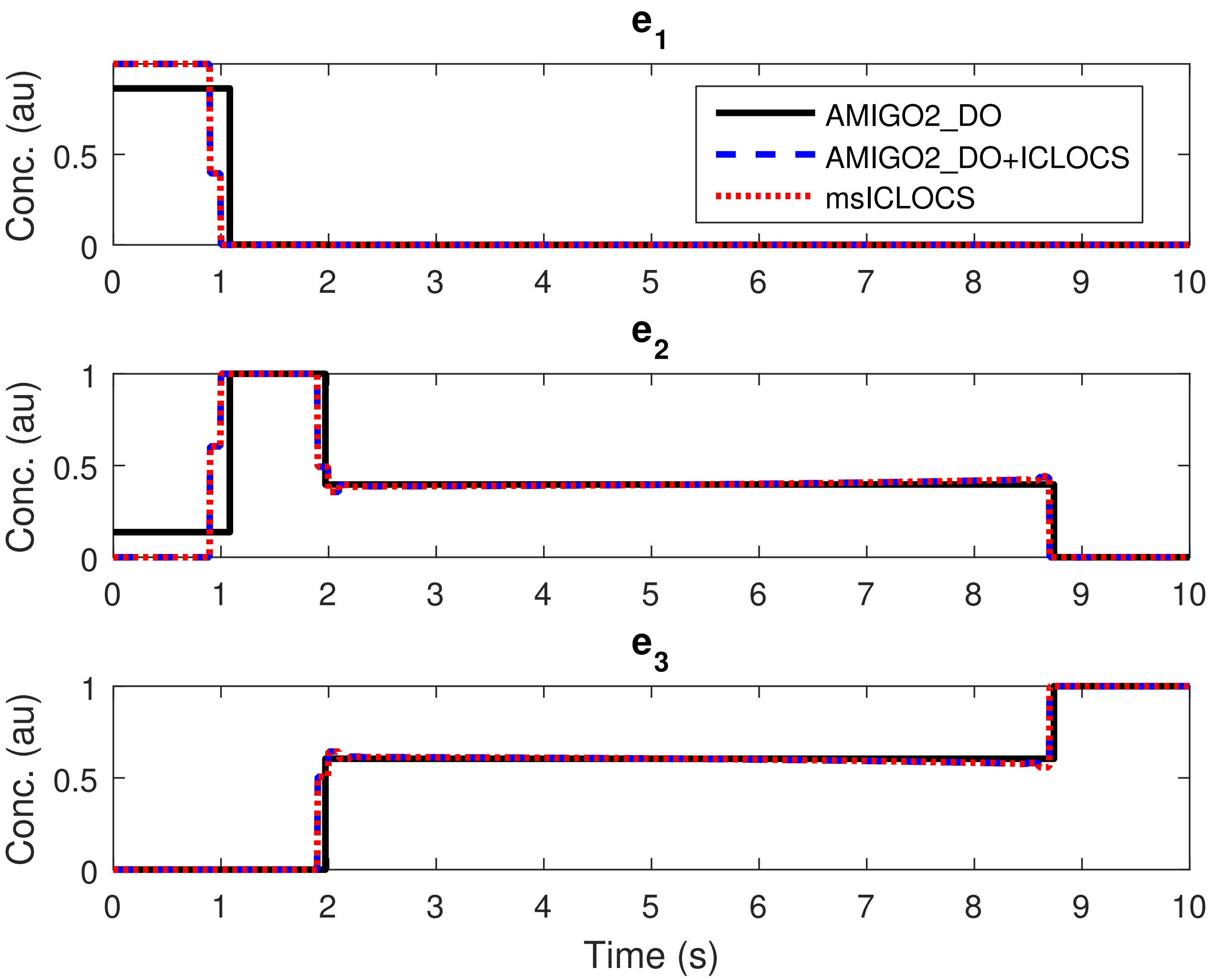


Figure S7: LPN3B: Comparison of the optimal control solutions for point C on the Pareto front. Each method compared is represented by a line of different color and style. Solid black line for `AMIGO2_DO` 4PWCv, dashed blue line for the hybrid of `AMIGO2_DO+ICLOCS` 100PWC and dotted red line for `msICLOCS` 100PWC.

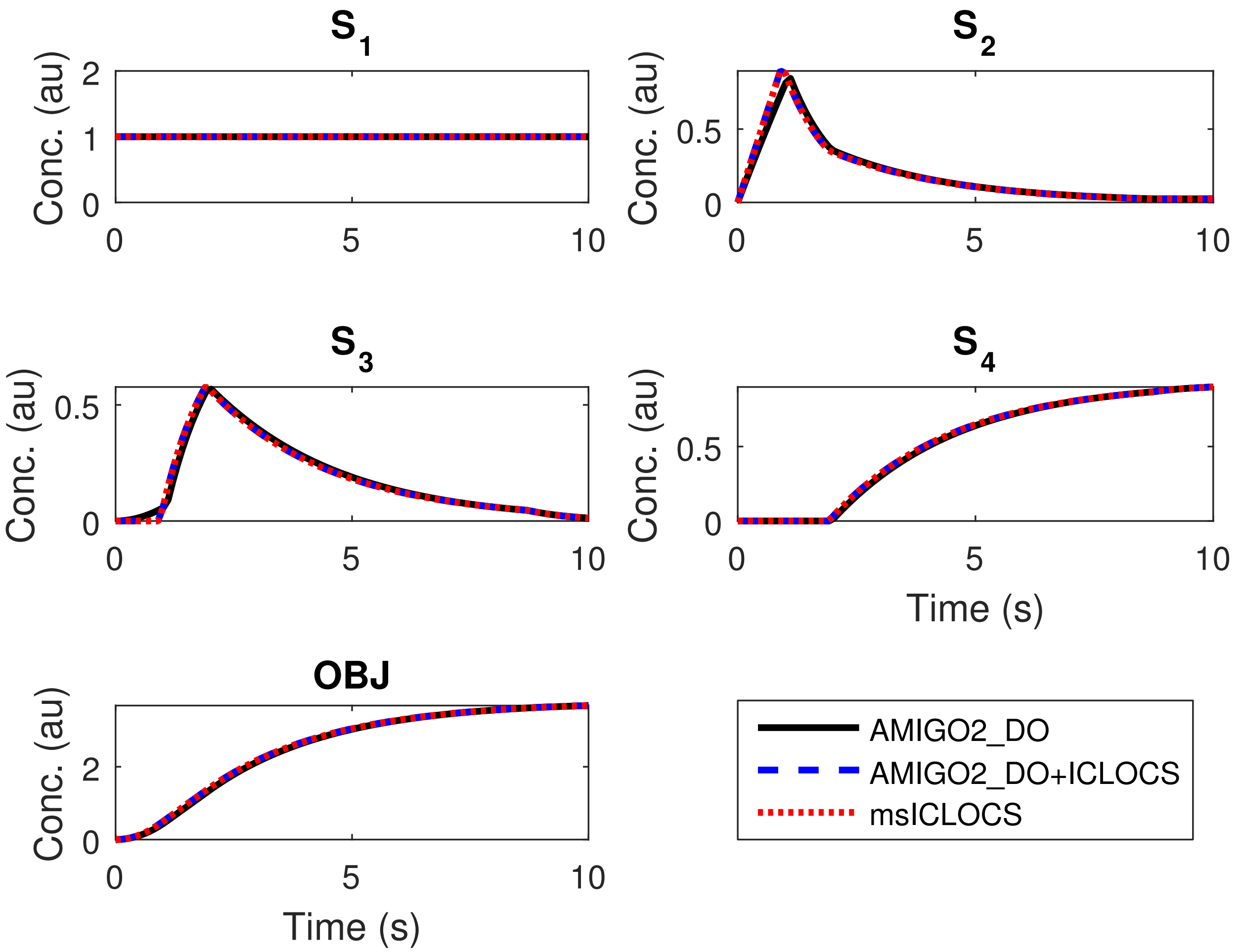


Figure S8: LPN3B: Comparison of the optimal state trajectories for point C on the Pareto front. Each method compared is represented by a line of different color and style. Solid black line for `AMIGO2_DO` 4PWCv, dashed blue line for the hybrid of `AMIGO2_DO+ICLOCS` 100PWC and dotted red line for `msICLOCS` 100PWC.

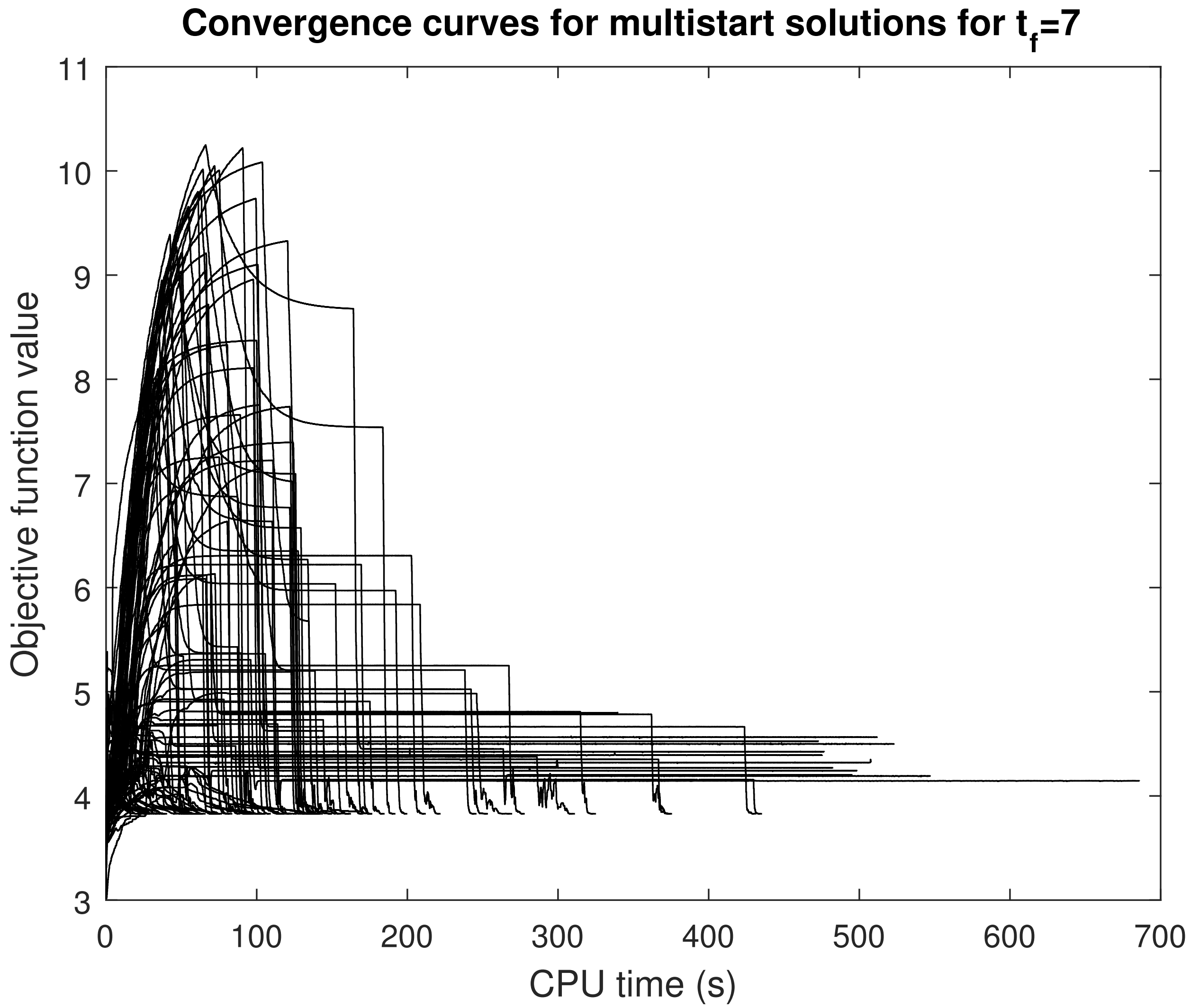


Figure S9: LPN3B: Convergence curves of a multistart (random initialization from 100 points) with ICLOCS 100PWC for point B on the Pareto front. Here the dynamics corresponding to the random initial point (100PWC controls) were not simulated and were not provided to ICLOCS as part of the initial guess. The initial guess provided was only the random controls.

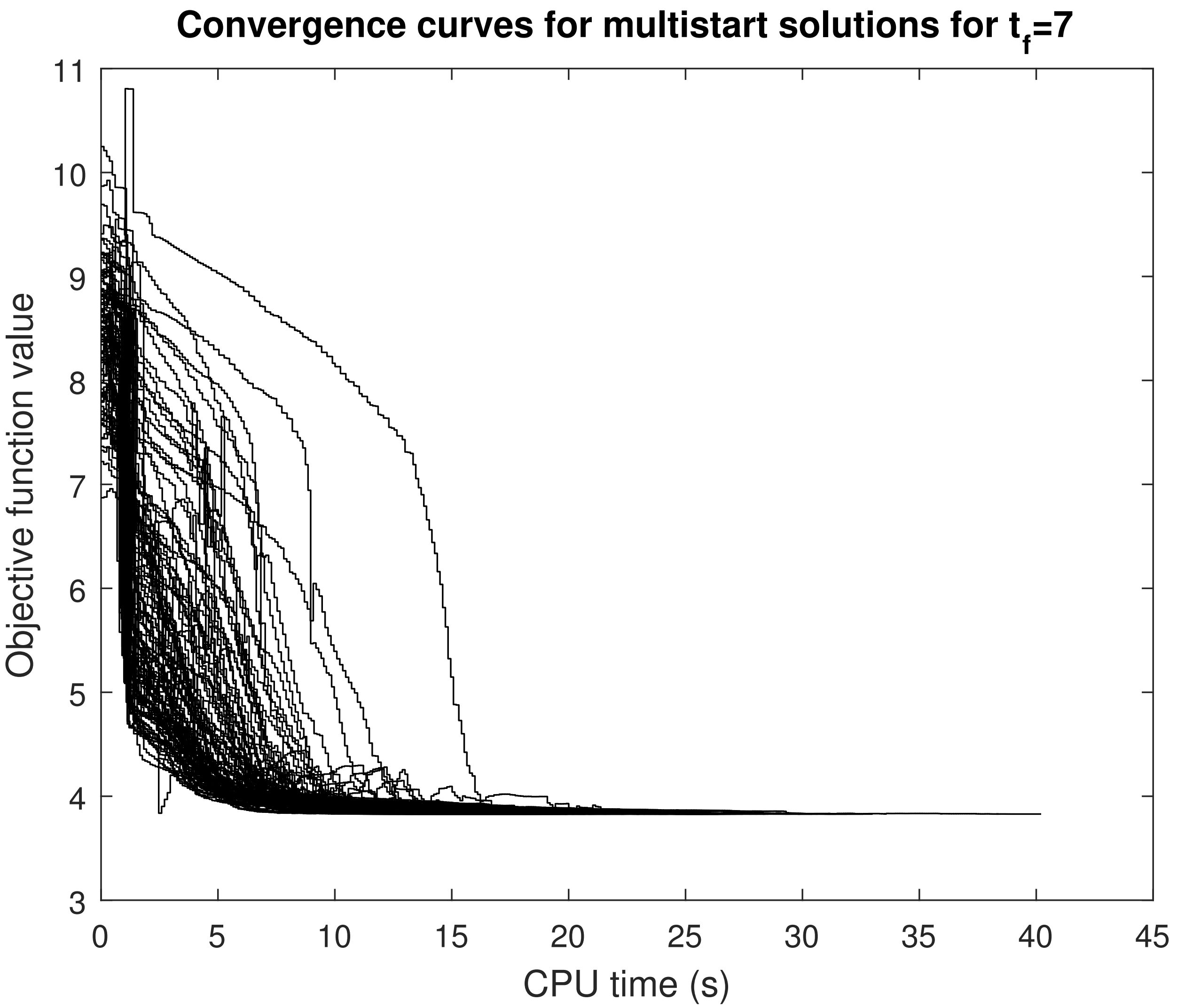


Figure S10: LPN3B: Convergence curves of a multistart (random initialization from 100 points) with `ICLOCS` 100PWC for point B on the Pareto front. However, in contrast with Figure S9, here the dynamics corresponding to the random initial point (100PWC controls) have been simulated and provided to `ICLOCS` as part of the initial guess. In this way, the transcribed constraint in the NLP formulation that correspond to the ODEs is not violated in the initial guess, resulting to a huge improvement in the convergence of the solver.

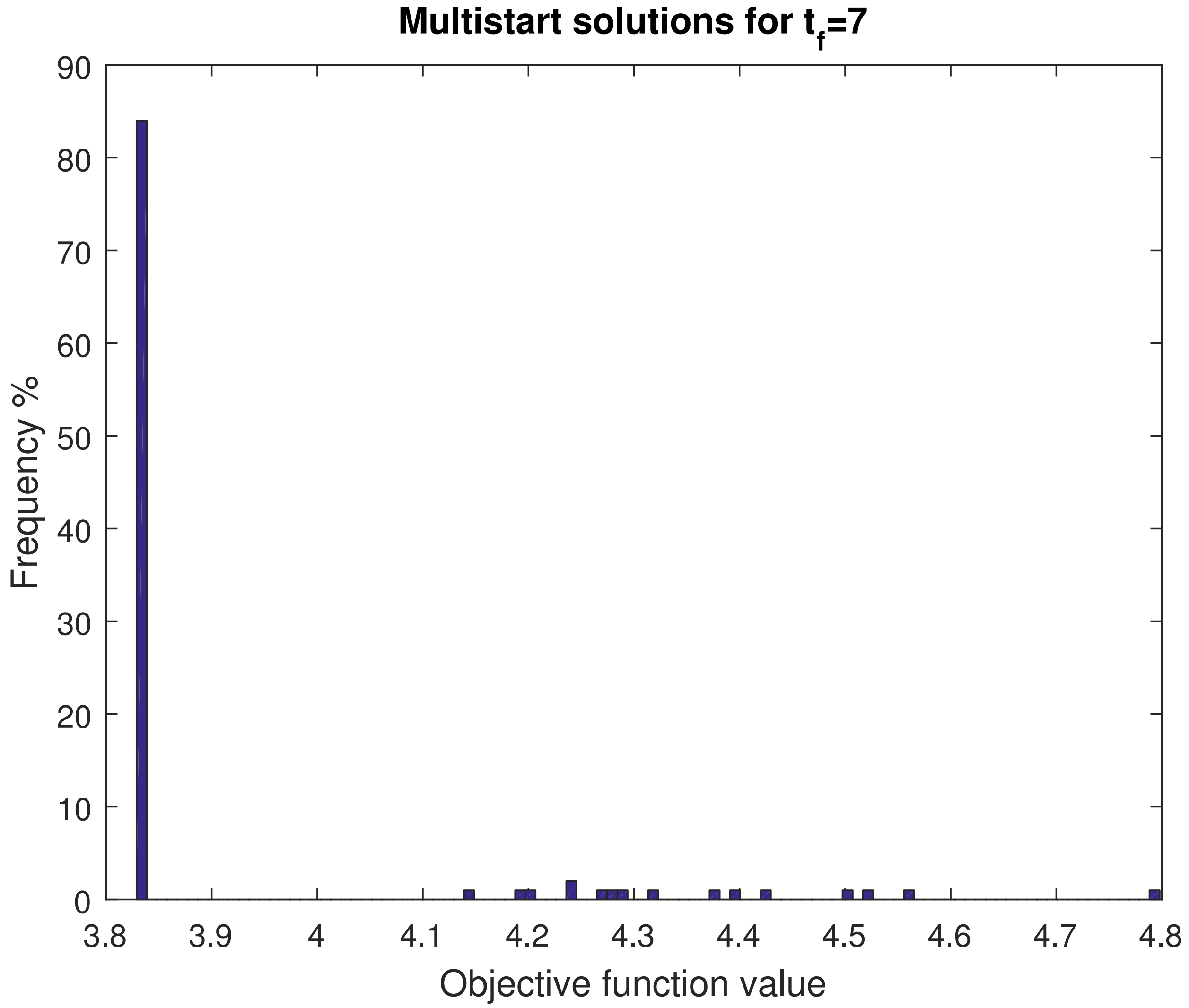


Figure S11: LPN3B: Solution histogram of the `msICLOCS` 100PWC for point B on the Pareto front, that correspond to Figure S9.

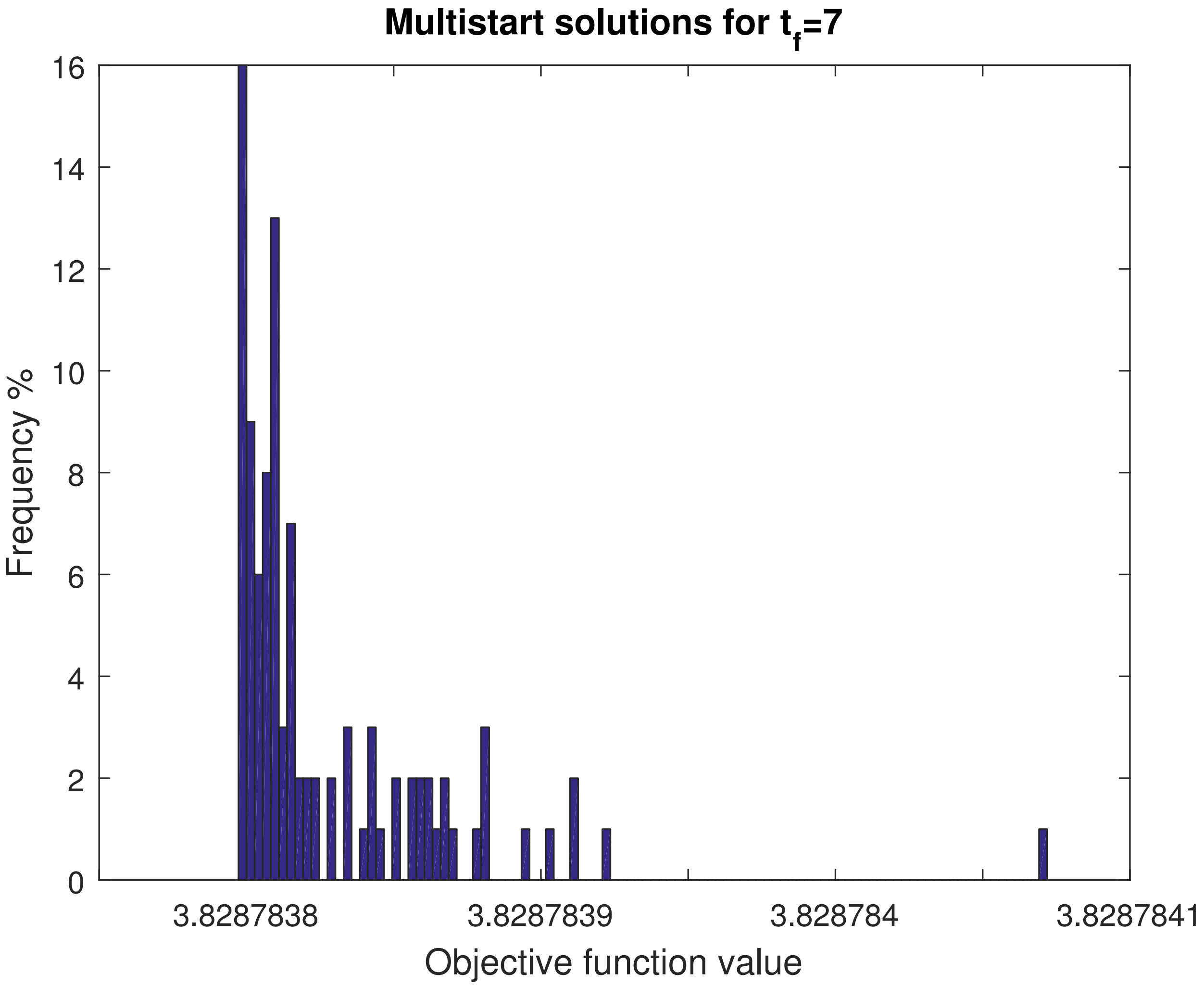


Figure S12: LPN3B: Solution histogram of the `msICLOCS` 100PWC for point B on the Pareto front, that correspond to Figure S10.

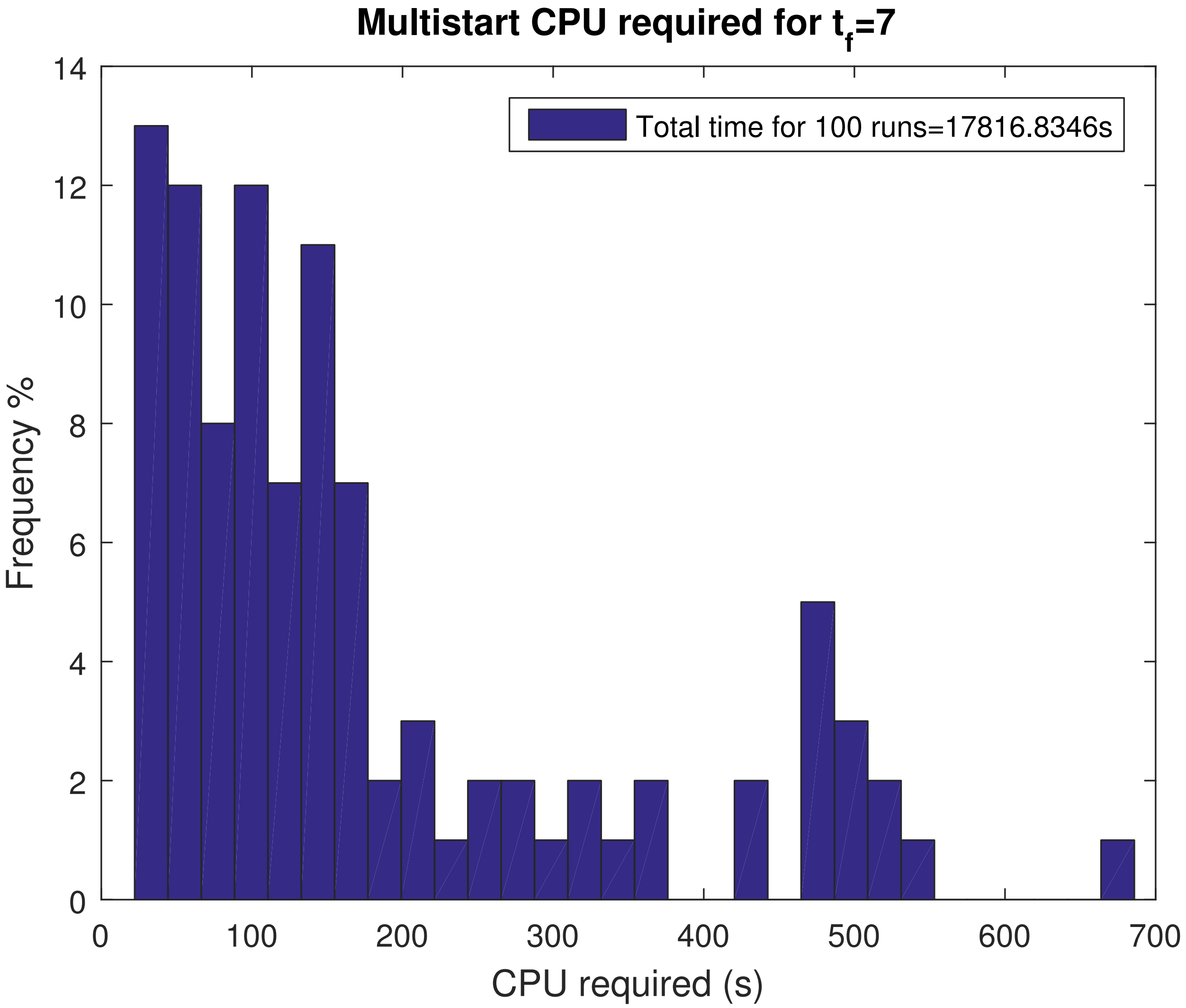


Figure S13: LPN3B: Solution histogram of the computational requirements of the `msICLOCS` 100PWC for point B on the Pareto front, that correspond to Figure S9.

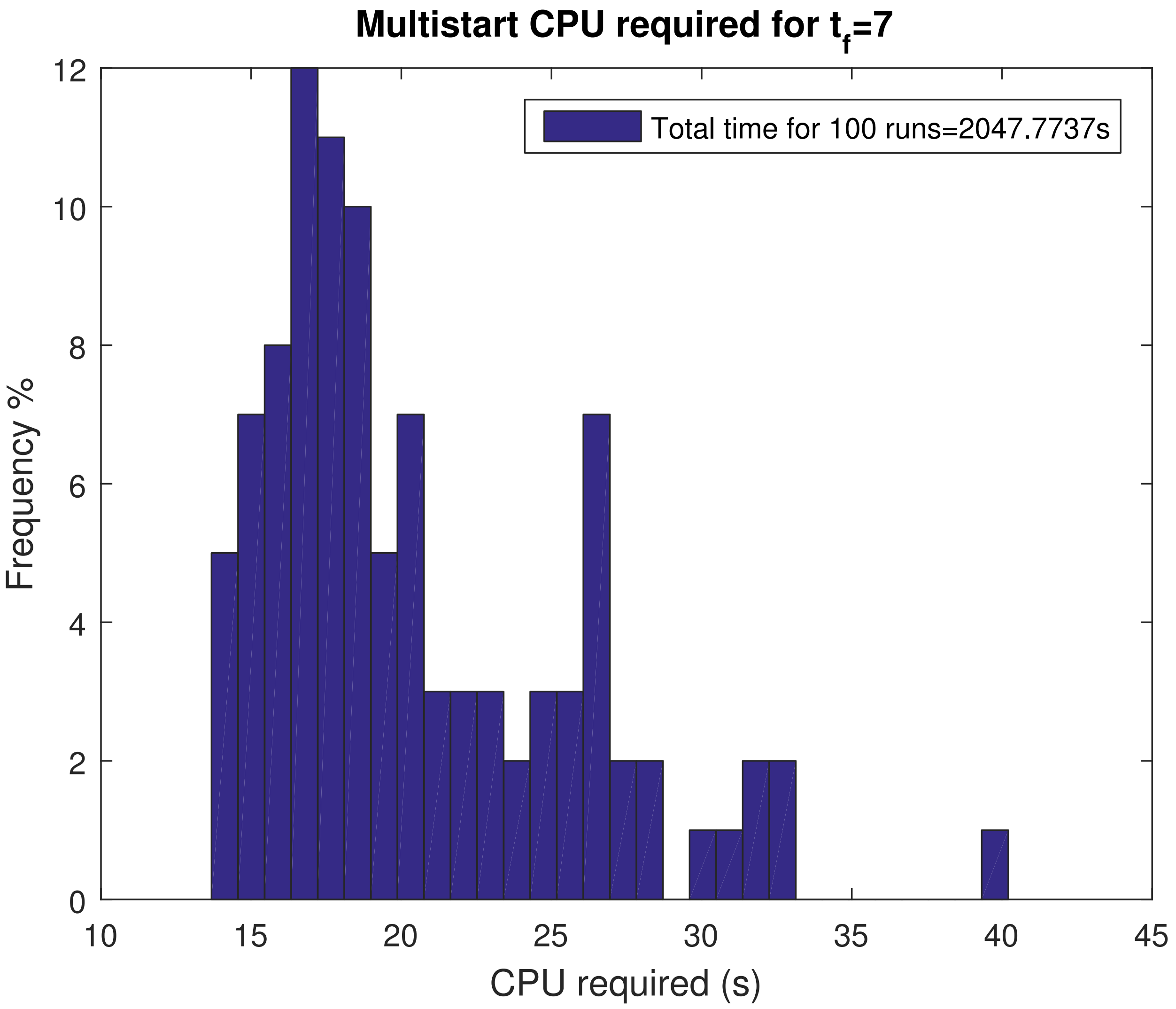


Figure S14: LPN3B: Solution histogram of the computational requirements of the `msICLOCS` 100PWC for point B on the Pareto front, that correspond to Figure S10.

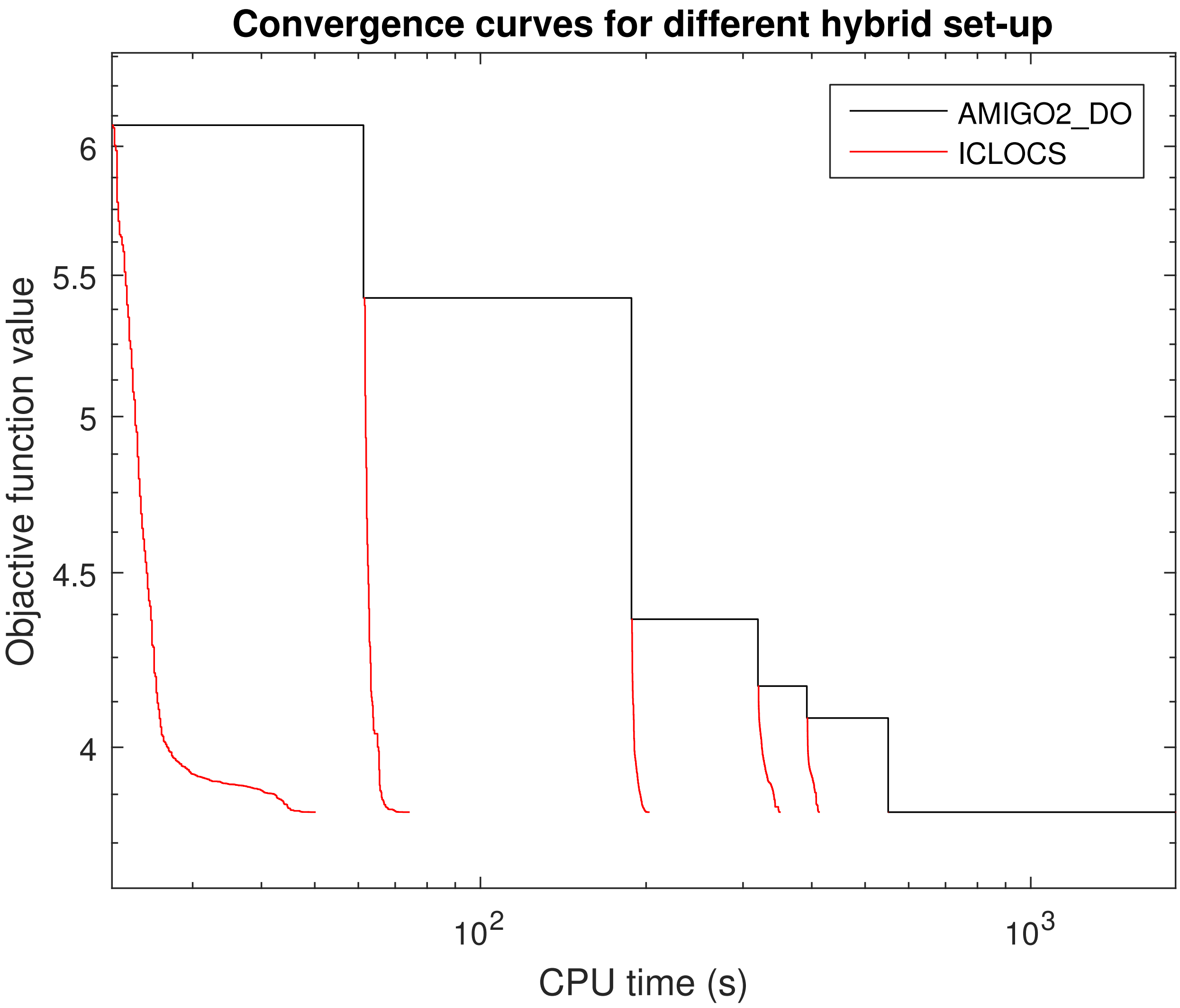


Figure S15: LPN3B: Convergence curves of the hybrid approach for point B on the Pareto front. ICLOCS (100PWC) is initialized from different point of the AMIGO2_DO (4PWCv) convergence.

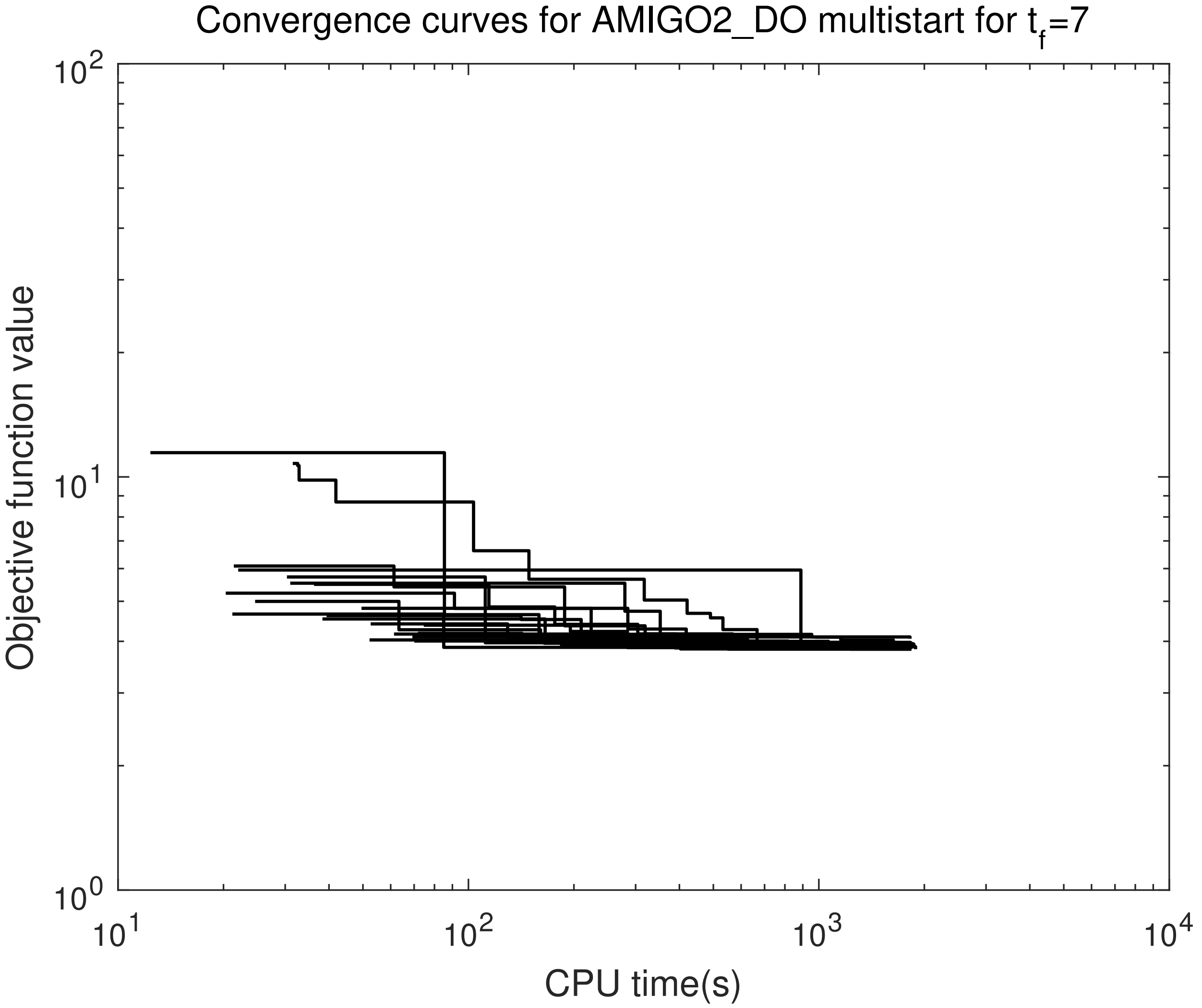


Figure S16: LPN3B: Convergence curves of a multistart of 20 runs with `AMIGO2_DO` (4PWCv) for point B on the Pareto front.

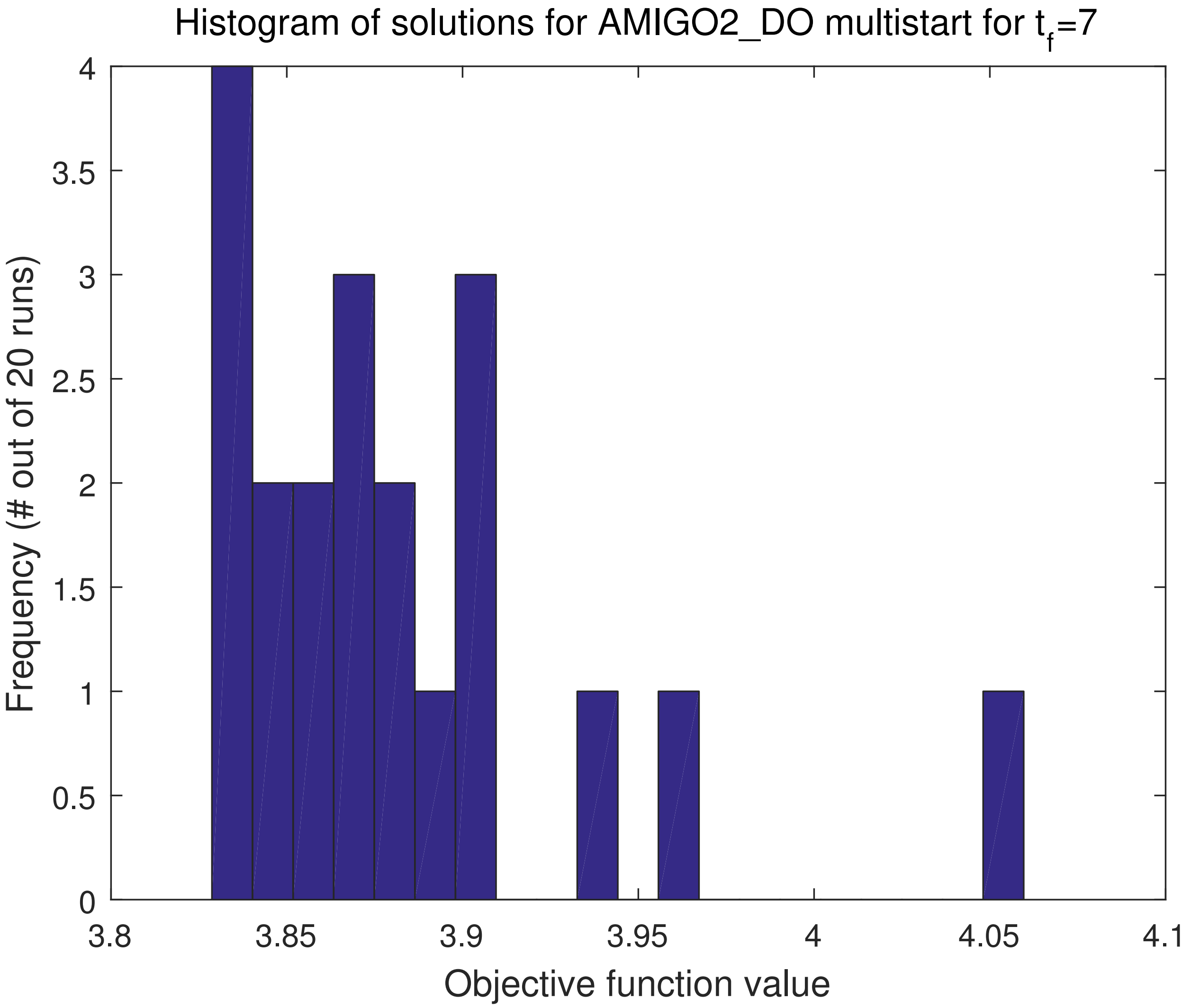


Figure S17: LPN3B: Solution histogram of the multistart of 20 runs with AMIGO2_DO (4PWCv) for point B on the Pareto front.

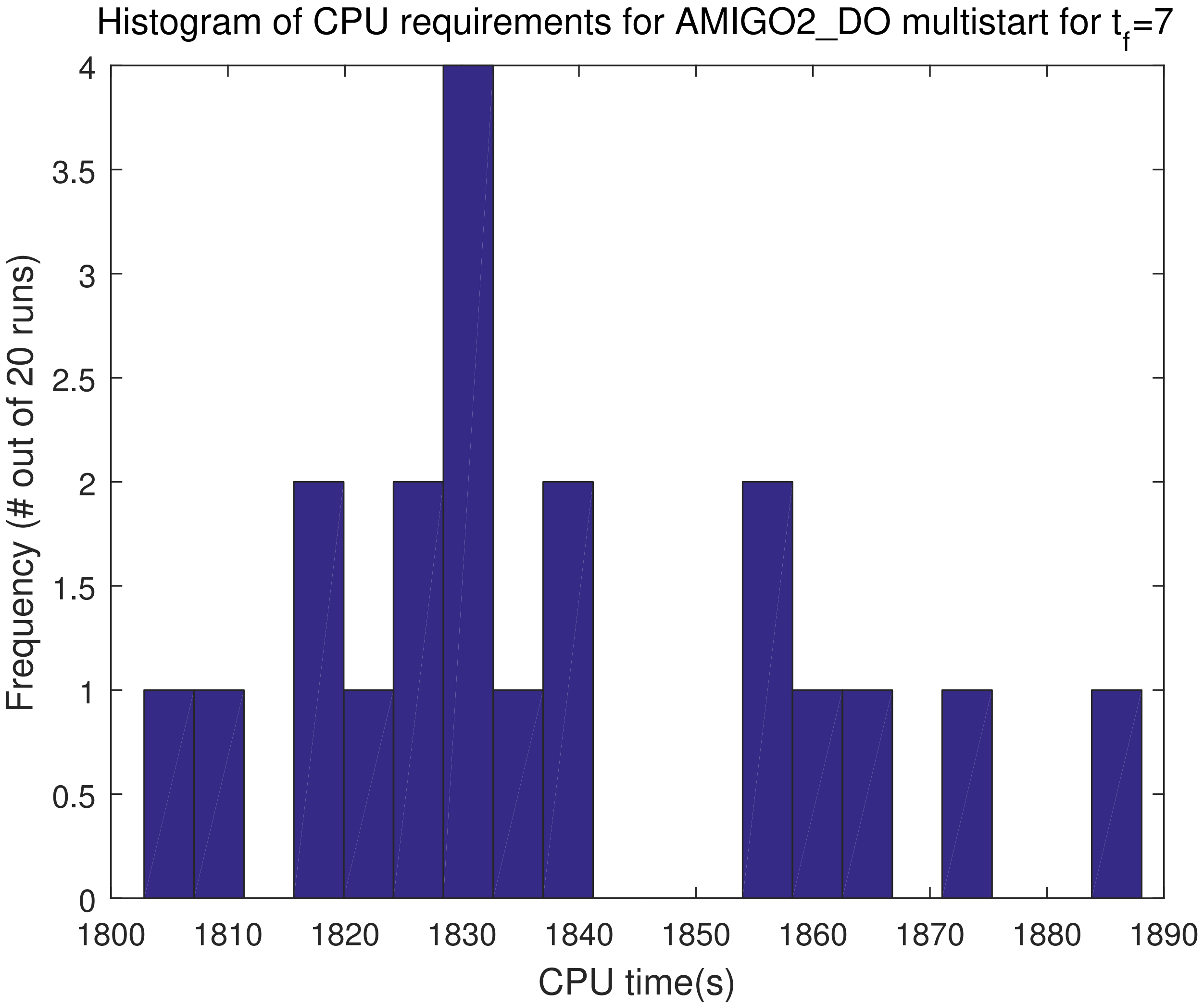


Figure S18: LPN3B: Computational requirements histogram of the multistart of 20 runs with AMIGO2_DO (4PWCv) for point B on the Pareto front.

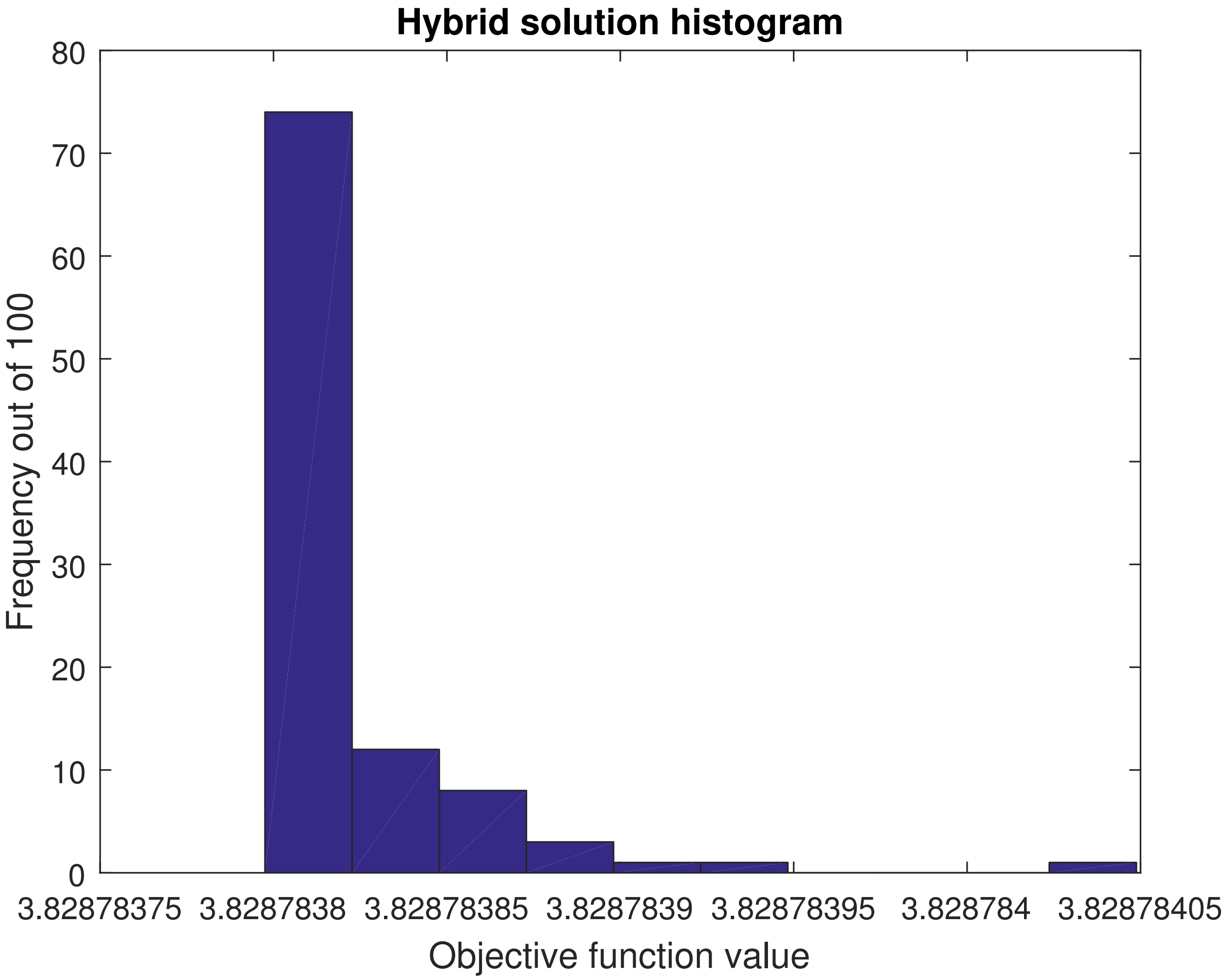


Figure S19: LPN3B: Histogram of the obtained solutions from the 100 `AMIGO2_DO+ICLOCS` runs for point B on the Pareto front. No multiplicity of solutions found.

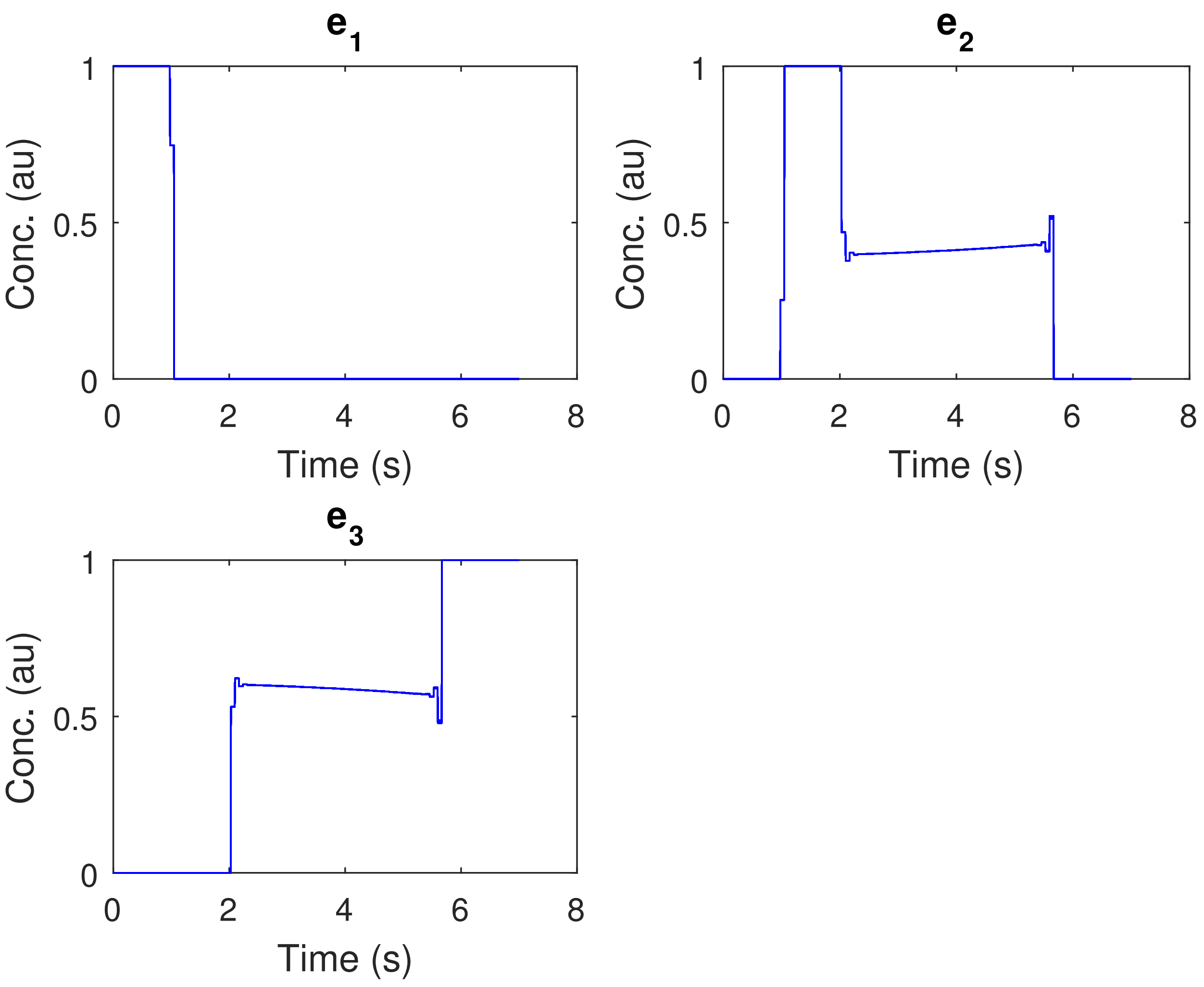


Figure S20: LPN3B: Optimal controls of all 100 AMIGO2_DO+ICLOCS solutions with 100PWC.

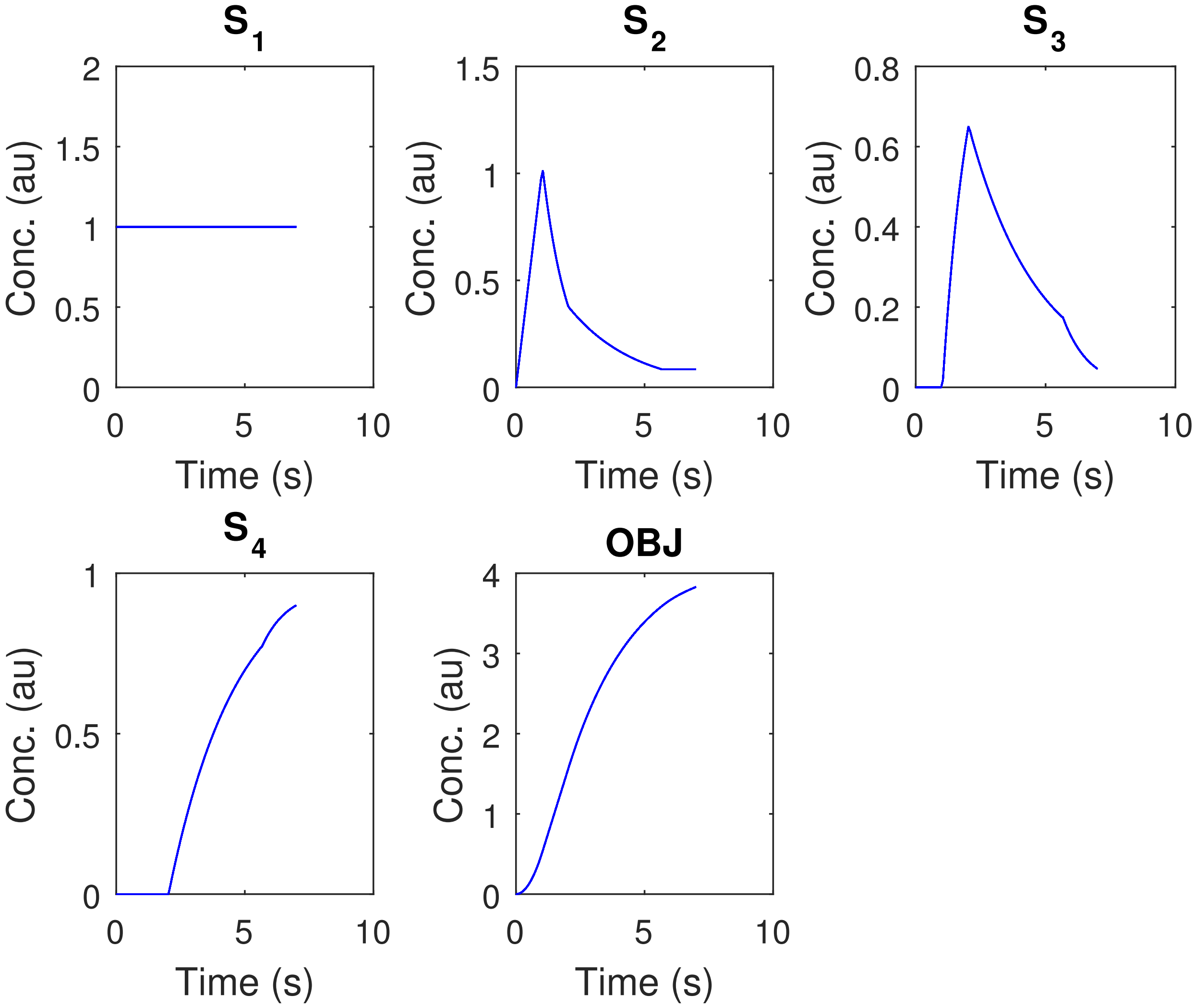


Figure S21: LPN3B: State trajectories of all 100 AMIGO2_DO+ICLOCS solutions with 100PWC.

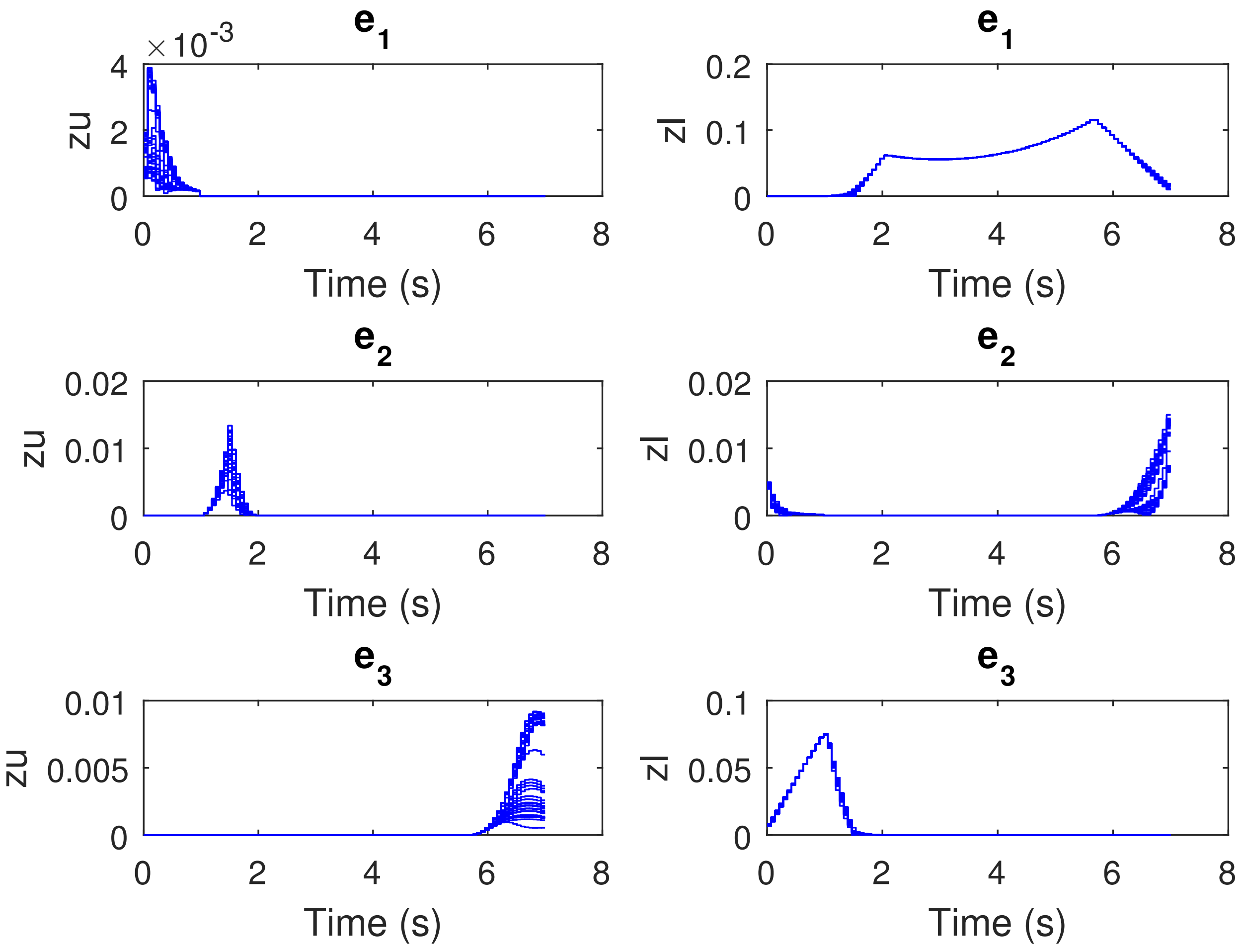


Figure S22: LPN3B: Lagrange multipliers for the controls' upper and lower bounds of all 100 `AMIGO2_DO+ICLOCS` solutions with 100PWC.

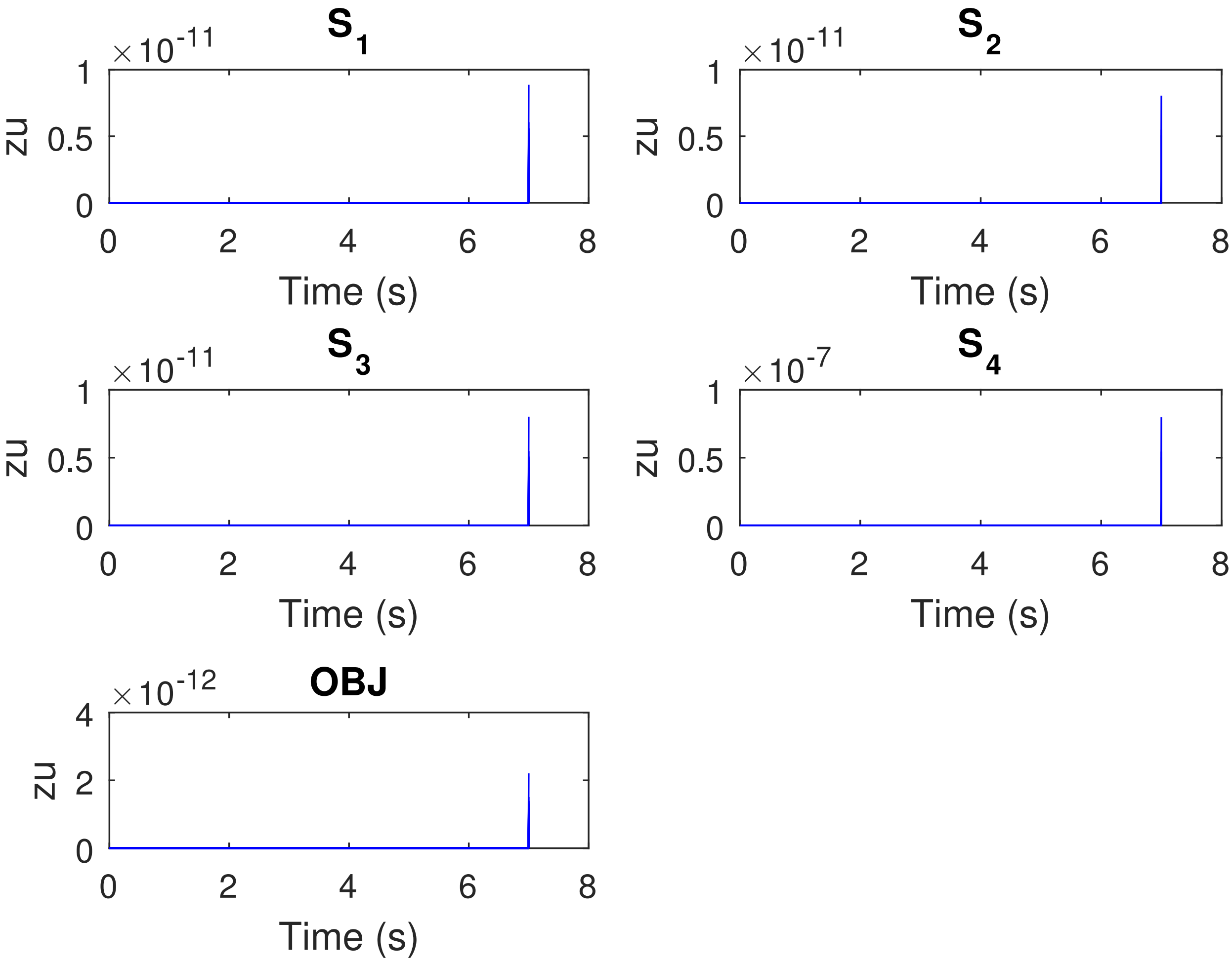


Figure S23: LPN3B: Lagrange multipliers for the states' upper bounds of all 100 AMIGO2_DO+ICLOCS solutions with 100PWC.

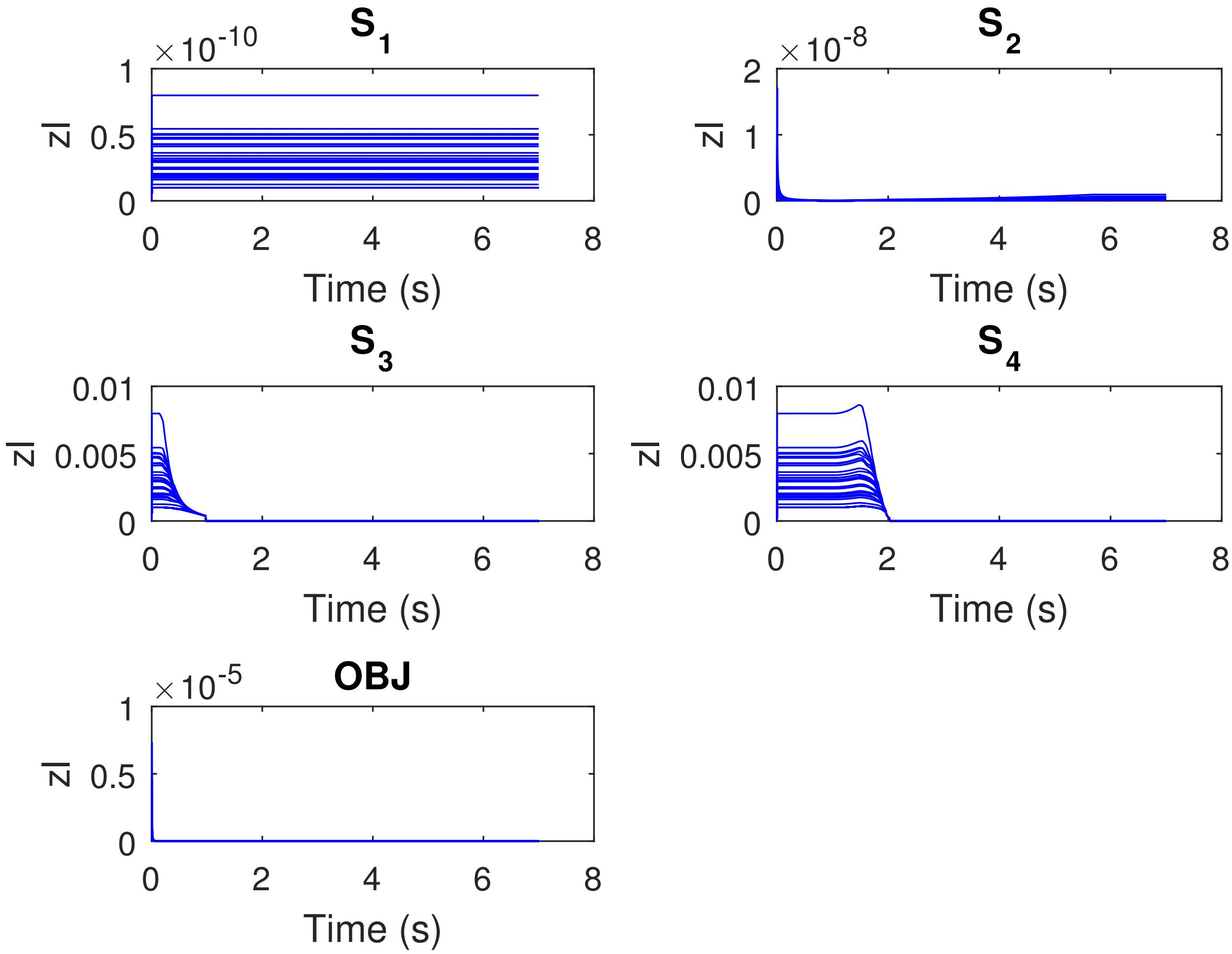


Figure S24: LPN3B: Lagrange multipliers for the states' lower bounds of all 100 `AMIGO2_DO+ICLOCS` solutions with 100PWC.

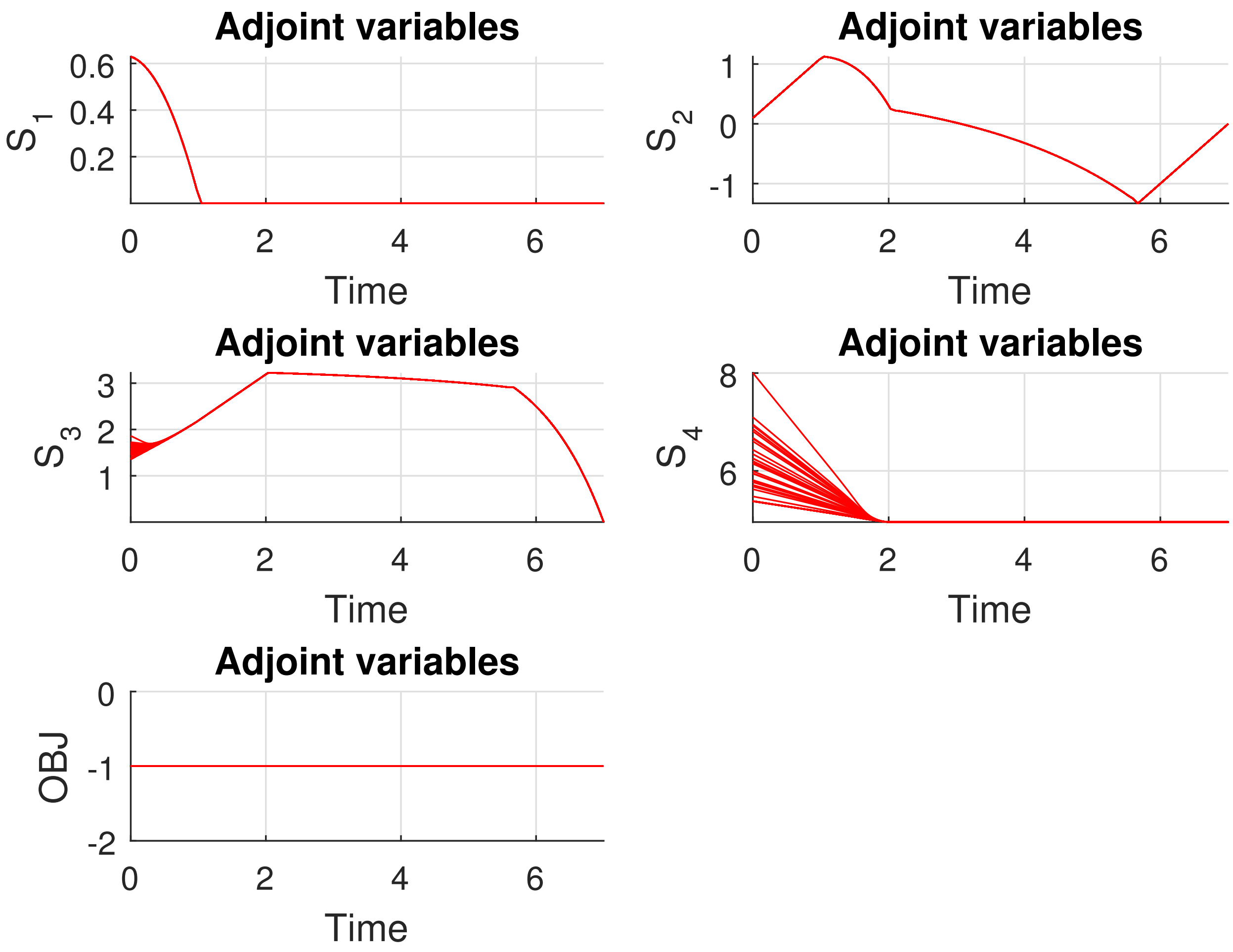


Figure S25: LPN3B: Adjoint variables of all 100 AMIGO2_DO+ICLOCS solutions with 100PWC.

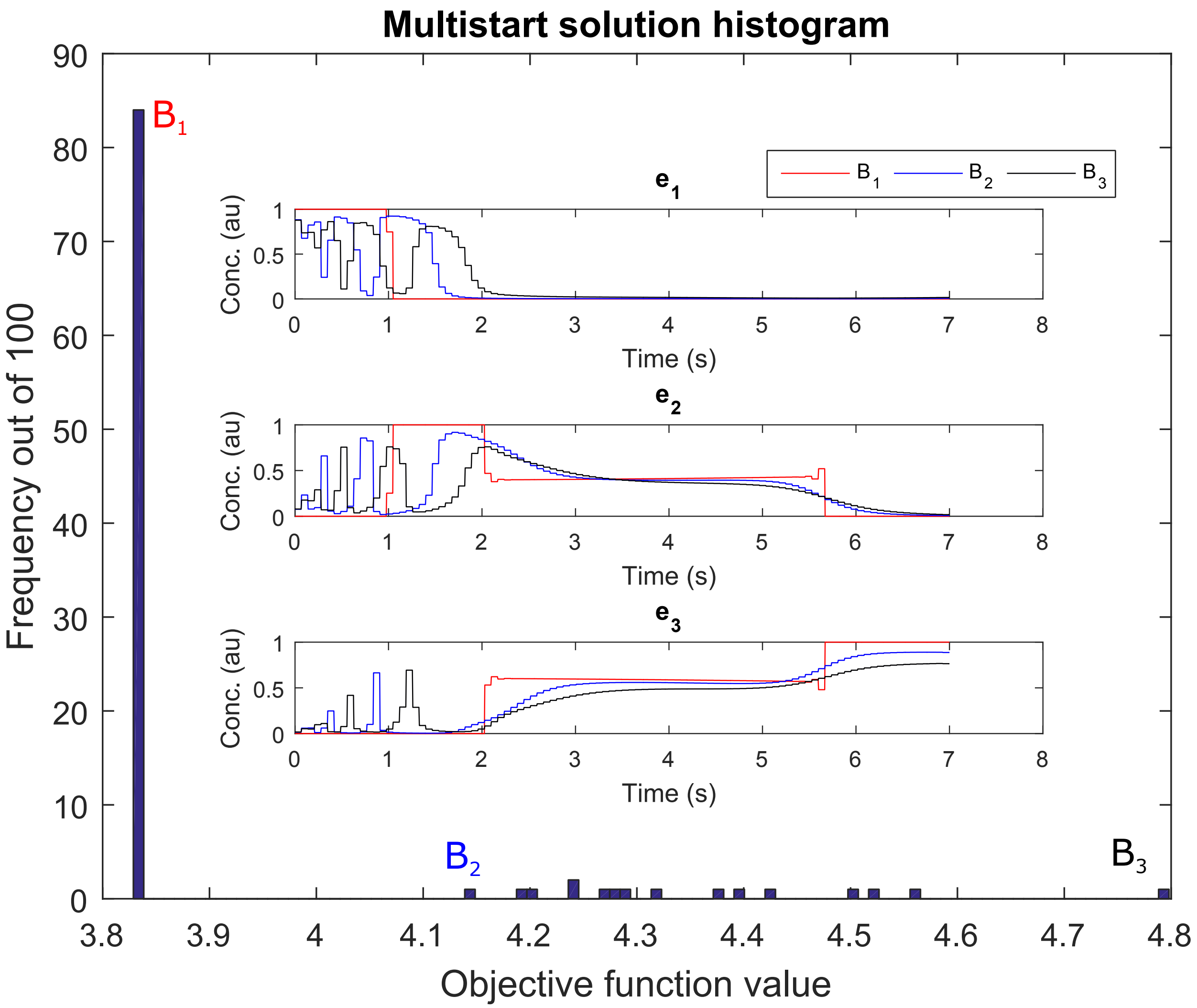


Figure S26: LPN3B: A comparison of the optimal controls that correspond to three solutions taken with the `msICLOCS` 100PWC for point B on the Pareto front, that correspond to Figures S11. The trajectories in pink correspond to the global optimal solution with objective function value around 3.83 while the ones in cyan and blue correspond to two local solutions with objective function values around 4.14 and 4.80 respectively.

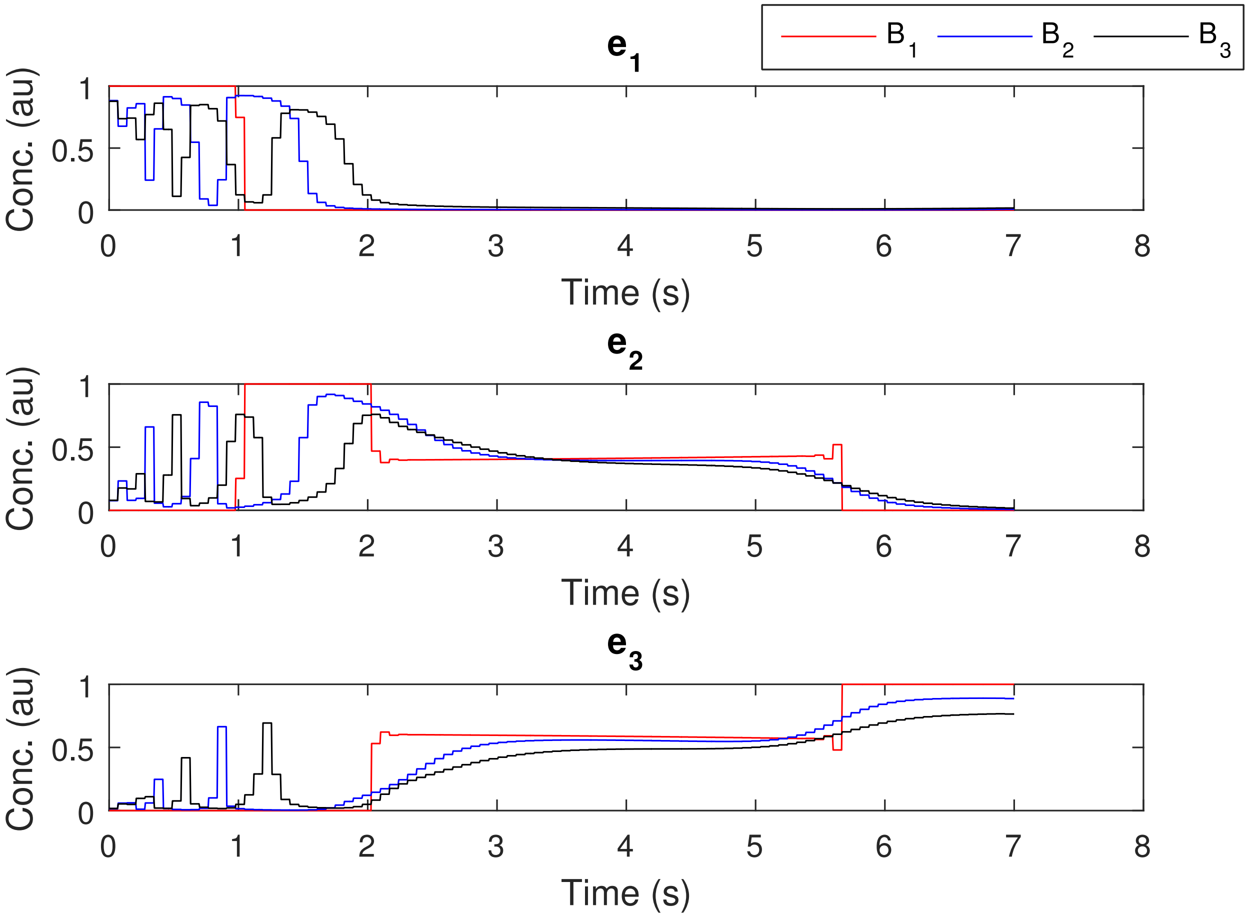


Figure S27: LPN3B: A comparison of the optimal controls that correspond to three solutions taken with the `msICLOCS` 100PWC for point B on the Pareto front, that correspond to Figures S11. The trajectories in pink correspond to the global optimal solution with objective function value around 3.83 while the ones in cyan and blue correspond to two local solutions with objective function values around 4.14 and 4.80 respectively.

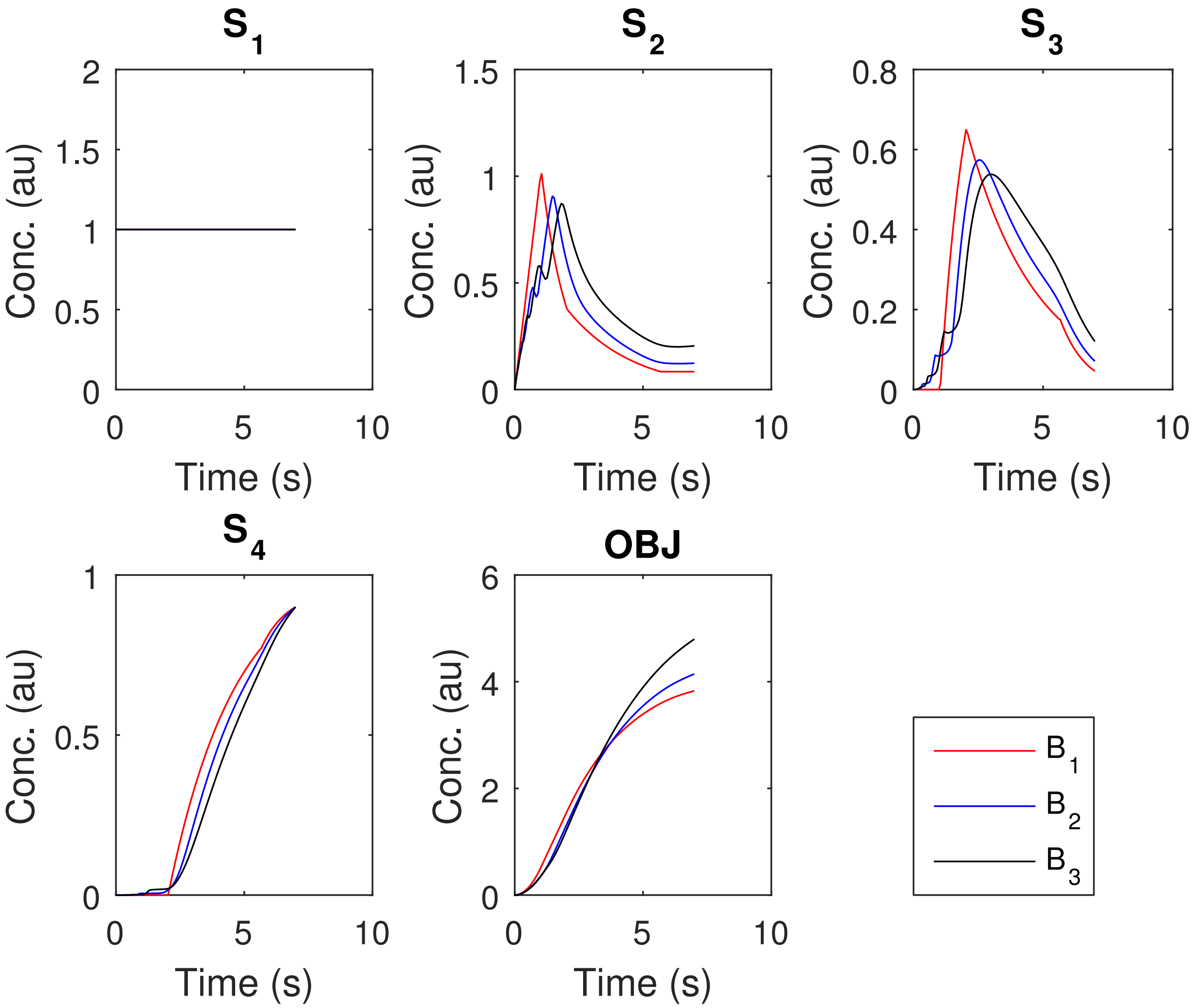


Figure S28: LPN3B: A comparison of the optimal state trajectories that correspond to three solutions taken with the `msICLOCS` 100PWC for point B on the Pareto front, that correspond to Figures S11. The trajectories in pink correspond to the global optimal solution with objective function value around 3.83 while the ones in cyan and blue correspond to two local solutions with objective function values around 4.14 and 4.80 respectively.
