## Additional file 2 - case SC for "Using optimal control to understand complex metabolic pathways"

### Additional file 2: Solution details for case study SC

### Supplementary information

As mentioned in the main text, in this case study the comparison between the three different approaches includes one important factor, the discretization. While in AMIGO2_DO+ICLOCS and msICLOCS a piecewise constant discretization with 10 elements (10PWC) has been used, in AMIGO2_DO the discretization used was 7 piecewise linear elements (7PWL). This is adding value to the comparison amongst them as the similar profiles that all three approaches resulted in, despite this difference in the discretization, provide evidence of successful inference of the true solution which seems to have a very defined shape.

In Figure S1 we show that despite their aforementioned difference, all approaches result to a very similar Pareto front. We further focus on these results by visualizing the optimal controls and the optimal state trajectories that correspond to points A, B and C, on the Pareto front, in Figures S2-S7.

Also, in Figure S8 we show the qualitative comparison with the real experimental transcriptomics data as mentioned in the main text. A similar comparison with the same experimental data has been previously done in the literature [1] and is used to argue that these theoretical results are in agreement with the observed biology.

Regarding the computational analysis, in Figure S9 we can see how the hybrid set-up can affect the approach's performance. We observe that after a certain time threshold the method avoids convergence in local solutions. In this more demanding case study, the hybrid approach is proven to be outperforming both AMIGO2_DO and msICLOCS.

AMIGO2_DO seems to require a lot more CPU time to converge to the near global optimal solution as we can observe in Figures S10-S12 which correspond to a multistart of AMIGO2_DO.

On the contrary msICLOCS presents different issues. Its best results are presented in Figures S13-S15 and have been taken using fine-tuning. However msICLOCS results in many local solutions, as shown in the histogram.

Similarly to LPN3B, we compare its performance with and without providing the system dynamics in the initial guess, in order to illustrate the critical role fine-tuning plays in such a solver. In Figures S16-S18 the results taken without providing the system dynamics in the initial guess are presented. Here we can observe a terrible performance, with a big increase in the local solutions, massive CPU requirements and a few solver crashes.

As we did in the LPN3B case study we analyzed the sensitivity of the controls with respect to the cost function. We did so by analyzing 100 runs of the hybrid were it was intentionally given too little CPU time. The resulting cost, controls and dynamics are presented in Figures S19-S21. Here the results are very interesting, especially from an engineering point of view. In Figure S19, we can observe a large number of distinct solutions within just the 0.5% of the solution range. Those solutions, as the results of a computer algorithm should be considered practically the same. Therefore we analyzed their differences in order to visualize how different their controls and states actually are. This is presented in Figures S22 and S23, where we observe that even though the cost difference is insignificant amongst these solutions their dynamics and controls have significant differences. This is evidence of insensitivity of the controls with respect to the cost function. In Figures S24-S26 we compare the best solution in the histogram ($K_1$) with the most frequent one ($K_2$ which is also found within the 0.5% range) and the worst one ($K_3$) in the histogram.

In Figures S27-S31 the computed Lagrange multipliers and adjoint variables are presented. As explained in the main text, this analysis can provide indications about the influence of some boundary conditions on the cost function and even though this analysis is not as straight forward as the respective analysis in linear programming it can be proven useful as a tool. For instance the orders of magnitude difference in the Lagrange multipliers corresponding to the lower bounds of ATP and NADH provided us with a clear indication that we should investigate the influence of those constraints on the solutions and more specifically whether ATP has more impact or not.

Consequently, we tried to numerically test the influence of the two co-factor critical values on the system dynamics. First we re-calculated the Pareto front after simultaneously changing both path constraints at the same time (see Figure S32). Second we re-caclulated the Pareto front after changing one out of the two path constraints at a time (see Figures S33, S34). Interestingly, we observed that indeed the impact of the critical ATP value was much bigger.

In Figures S35-S38 we present some comparisons of the optimal controls and optimal state trajectories for the same survival time (80h) and for the extreme points of the Pareto fronts corresponding to different critical values. We observe how these different critical values can result to non-intuitive changes in the dynamics and in the temporal activation of the enzymes. Interestingly we also observe that the qualitative strategy depicted in the state dynamics of all extreme points is very similar.

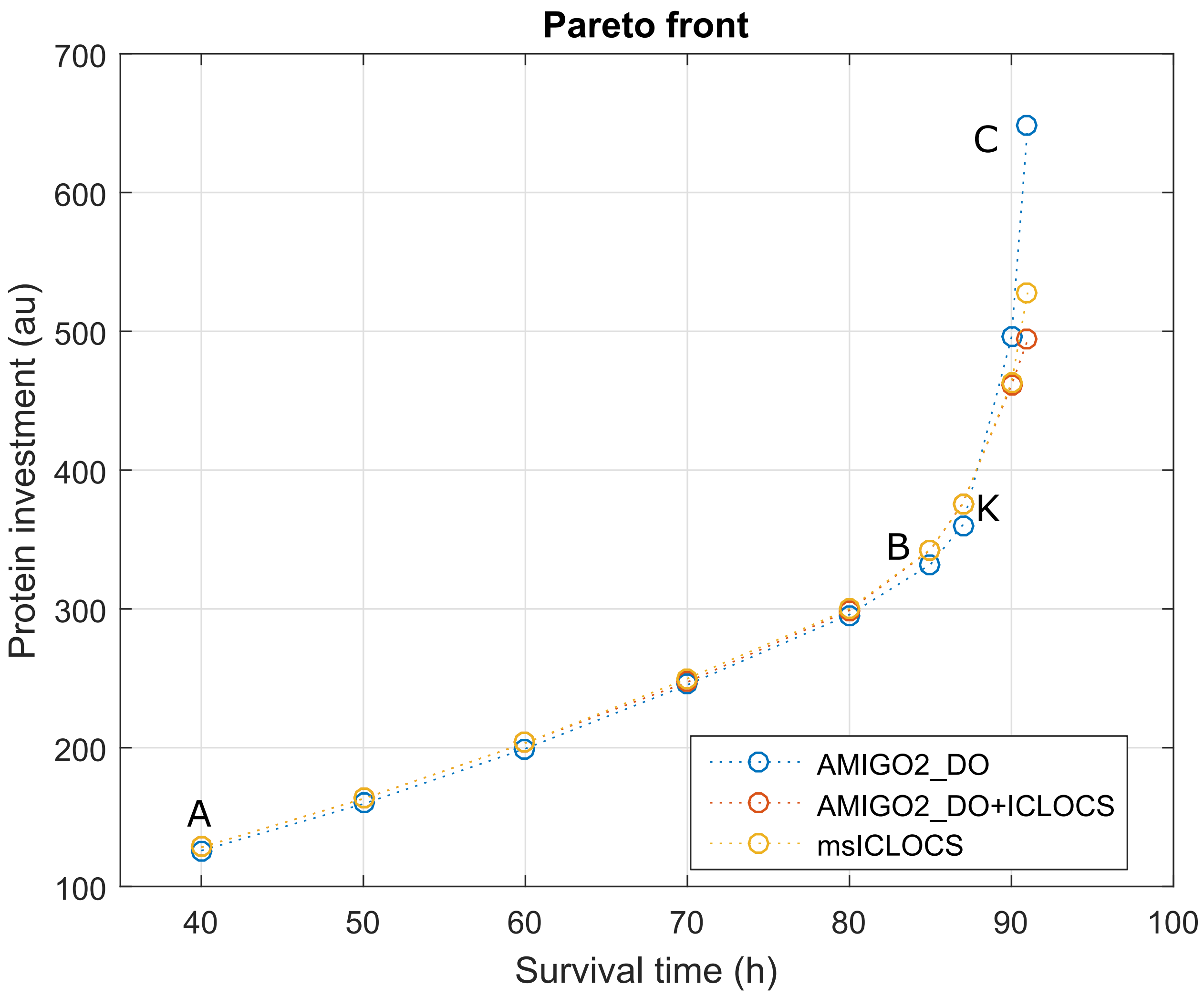


Figure S1: SC: Pareto front obtained with three different approaches, for SC case study. Note that the two extreme points of the Pareto front resulted by each of the individual approaches are named with the letters A and C, as well as the points around the knee of the Pareto front are named B and O. For instance, the letter C corresponds to the solutions taken with the three different approaches for survival time of 91h.

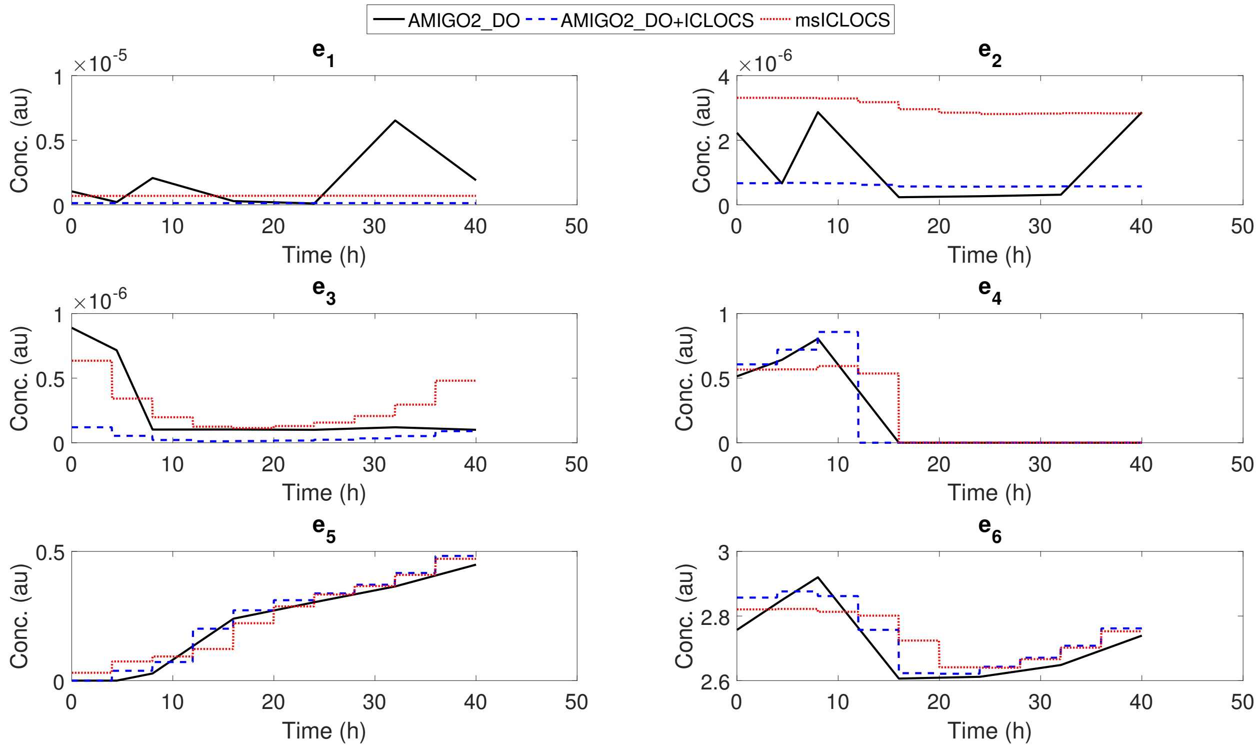


Figure S2: SC: Comparison of the optimal control solutions for point A on the Pareto front. Each method compared is represented by a line of different color and style. Solid black line for `AMIGO2_DO` 6PWL, dashed blue line for the hybrid `AMIGO2_DO+ICLOCS` 10PWC and dotted red line for `msICLOCS` 10PWC.

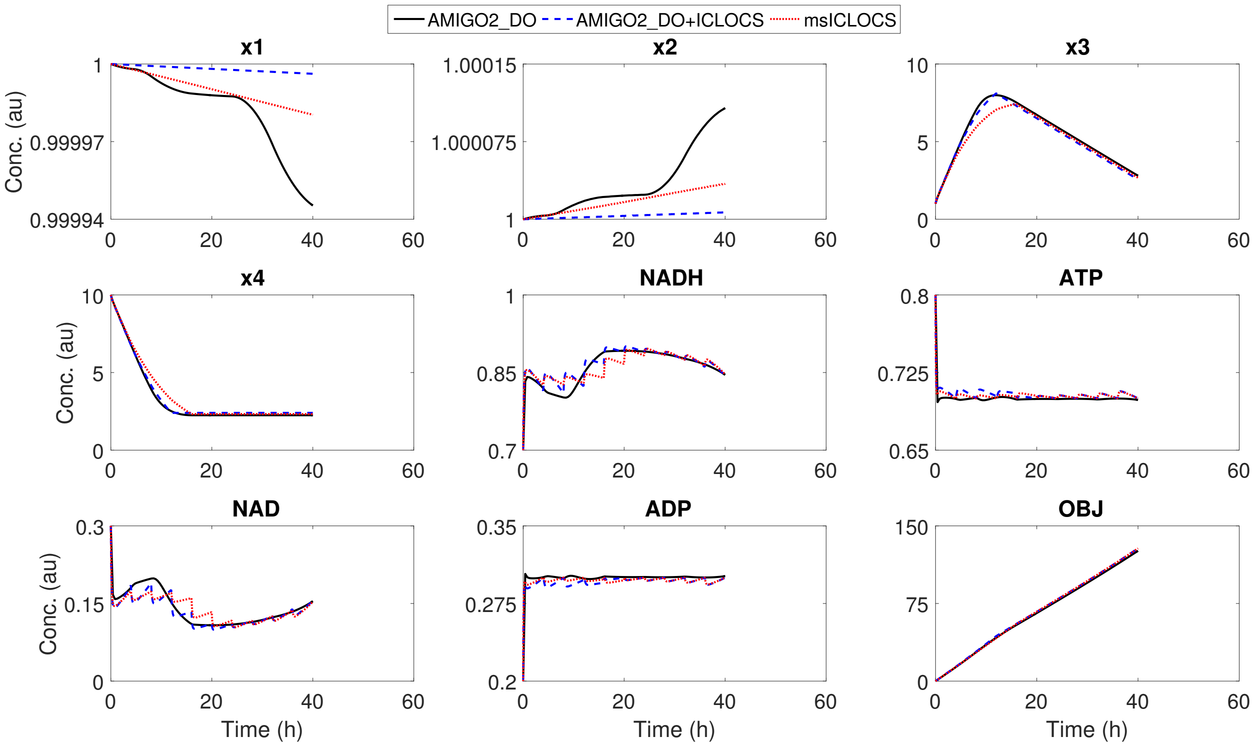


Figure S3: SC: Comparison of the optimal state trajectories solutions for point A on the Pareto front. Each method compared is represented by a line of different color and style. Solid black line for `AMIGO2_DO` 6PWL, dashed blue line for the hybrid `AMIGO2_DO+ICLOCS` 10PWC and dotted red line for `msICLOCS` 10PWC.

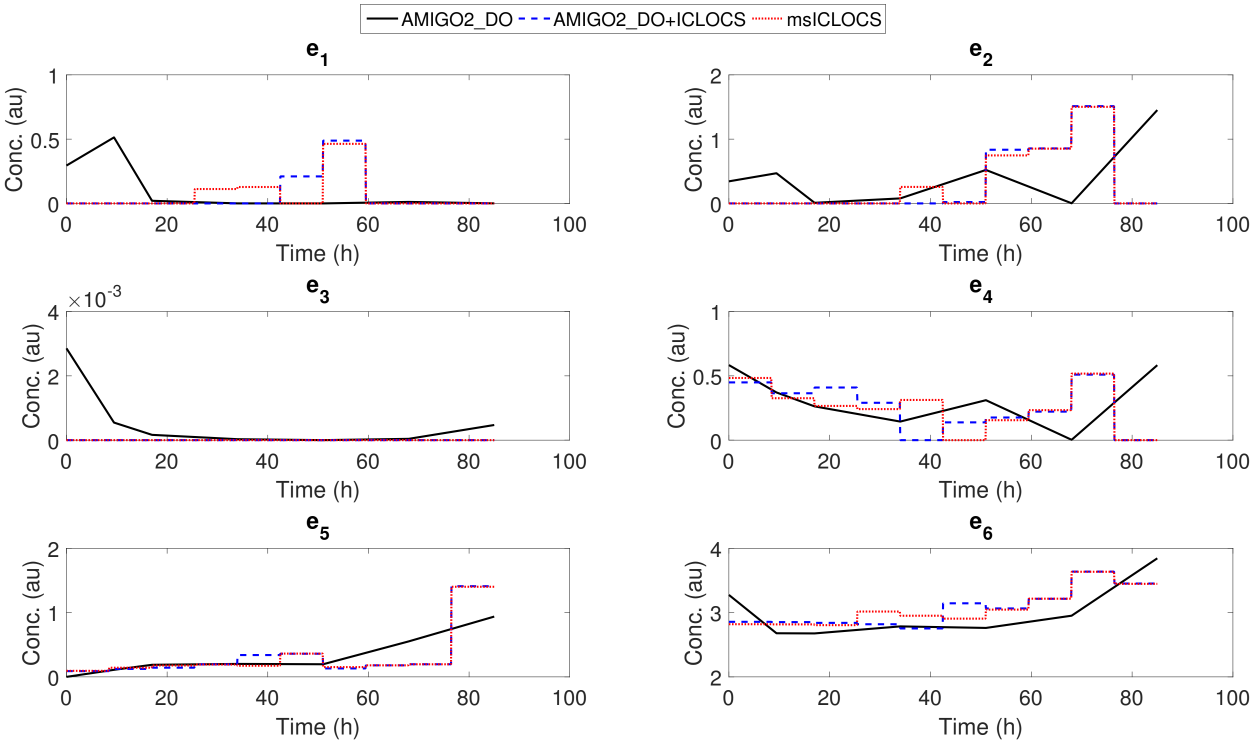


Figure S4: SC: Comparison of the optimal control solutions for point B on the Pareto front. Each method compared is represented by a line of different color and style. Solid black line for `AMIGO2_DO` 6PWL, dashed blue line for the hybrid of `AMIGO2_DO+ICLOCS` 10PWC and dotted red line for `msICLOCS` 10PWC.

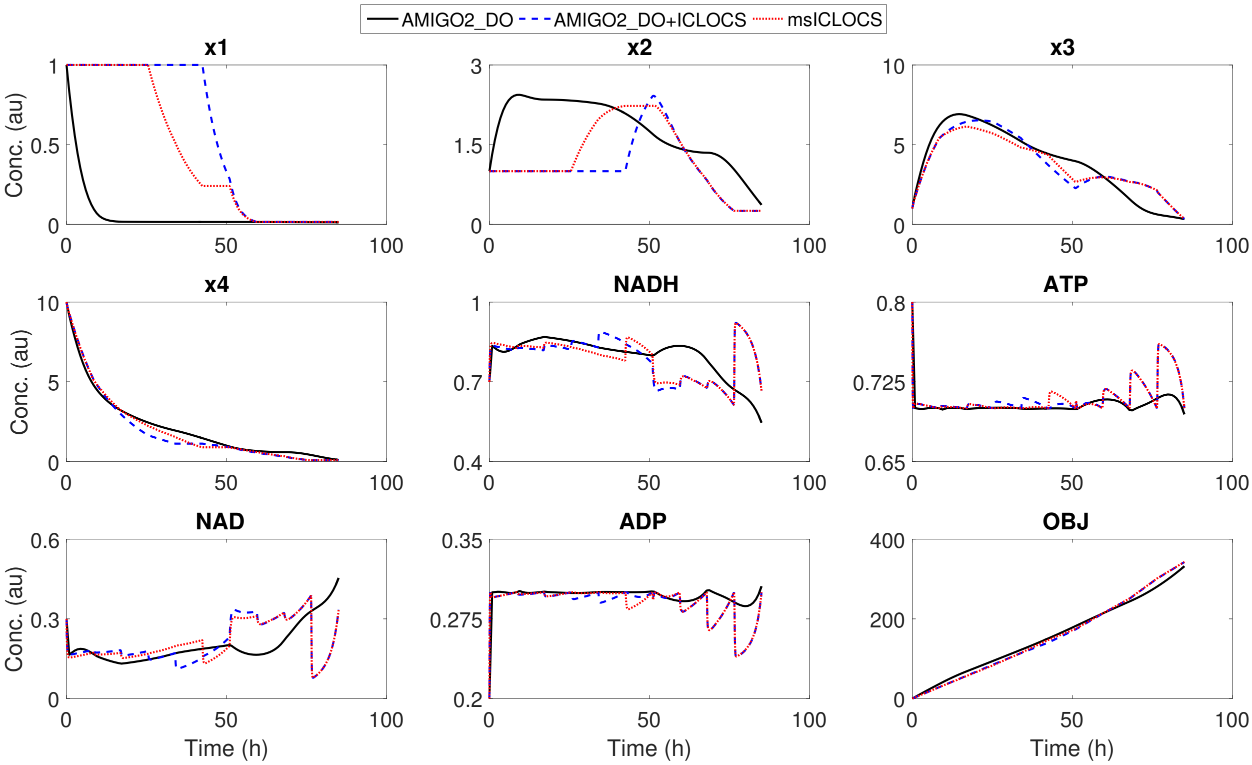


Figure S5: SC: Comparison of the optimal state trajectories solutions for point B on the Pareto front. Each method compared is represented by a line of different color and style. Solid black line for `AMIGO2_DO` 6PWL, dashed blue line for the hybrid `AMIGO2_DO+ICLOCS` 10PWC and dotted red line for `msICLOCS` 10PWC.

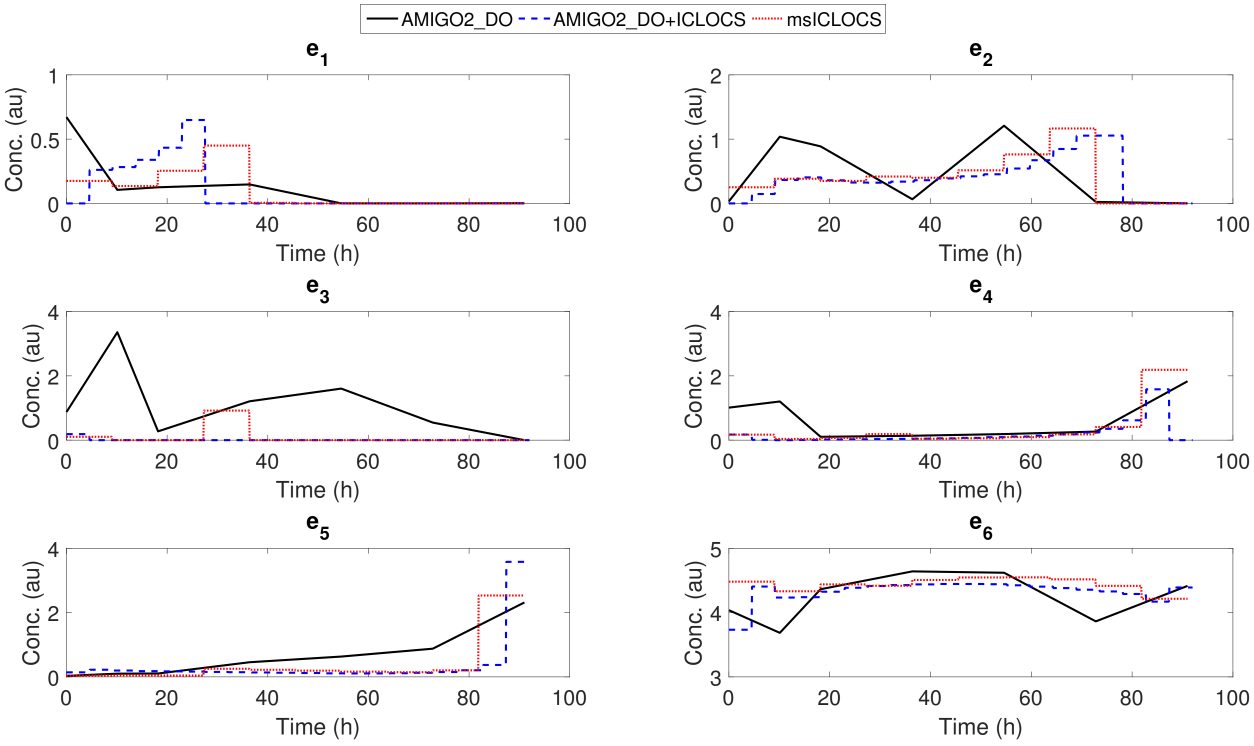


Figure S6: SC: Comparison of the optimal control solutions for point C on the Pareto front. Each method compared is represented by a line of different color and style. Solid black line for `AMIGO2_DO` 6PWL, dashed blue line for the hybrid `AMIGO2_DO+ICLOCS` 10PWC and dotted red line for `msICLOCS` 10PWC.

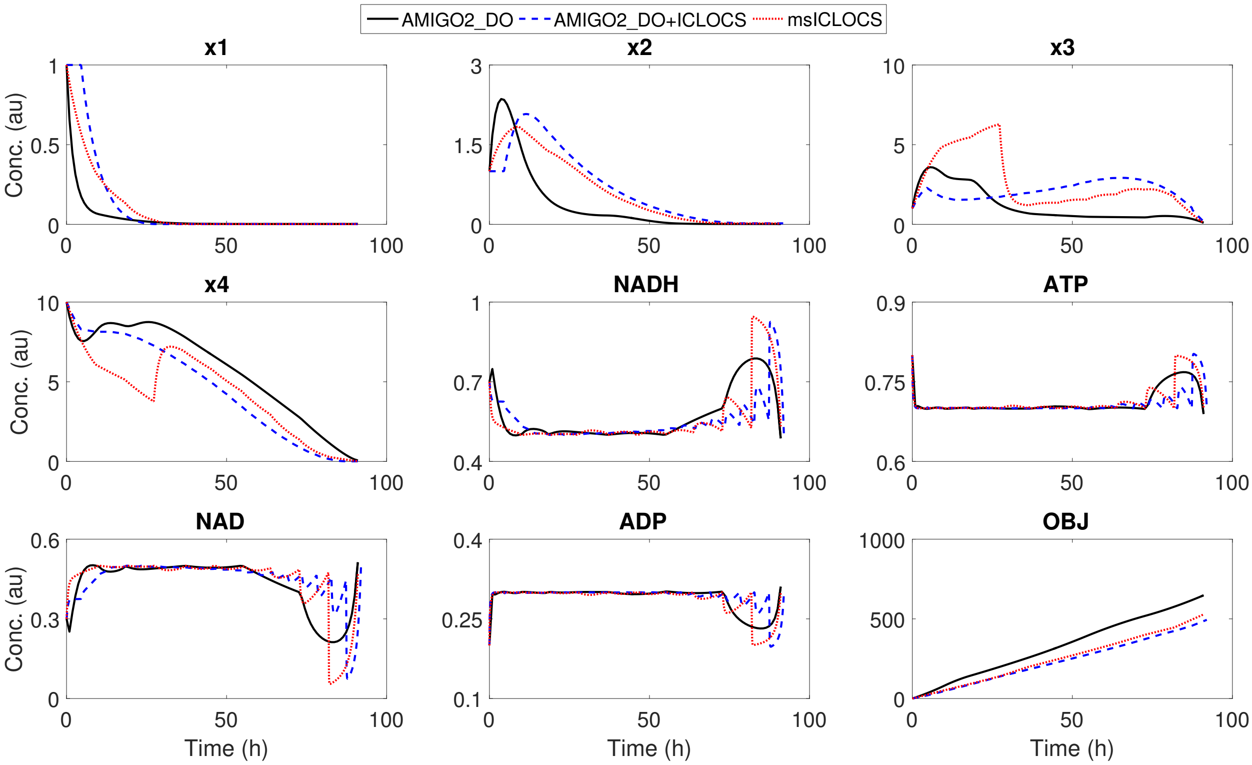


Figure S7: SC: Comparison of the optimal state trajectories solutions for point C on the Pareto front. Each method compared is represented by a line of different color and style. Solid black line for AMIGO2_DO 6PWL, dashed blue line for the hybrid AMIGO2_DO+ICLOCS 10PWC and dotted red line for msICLOCS 10PWC.

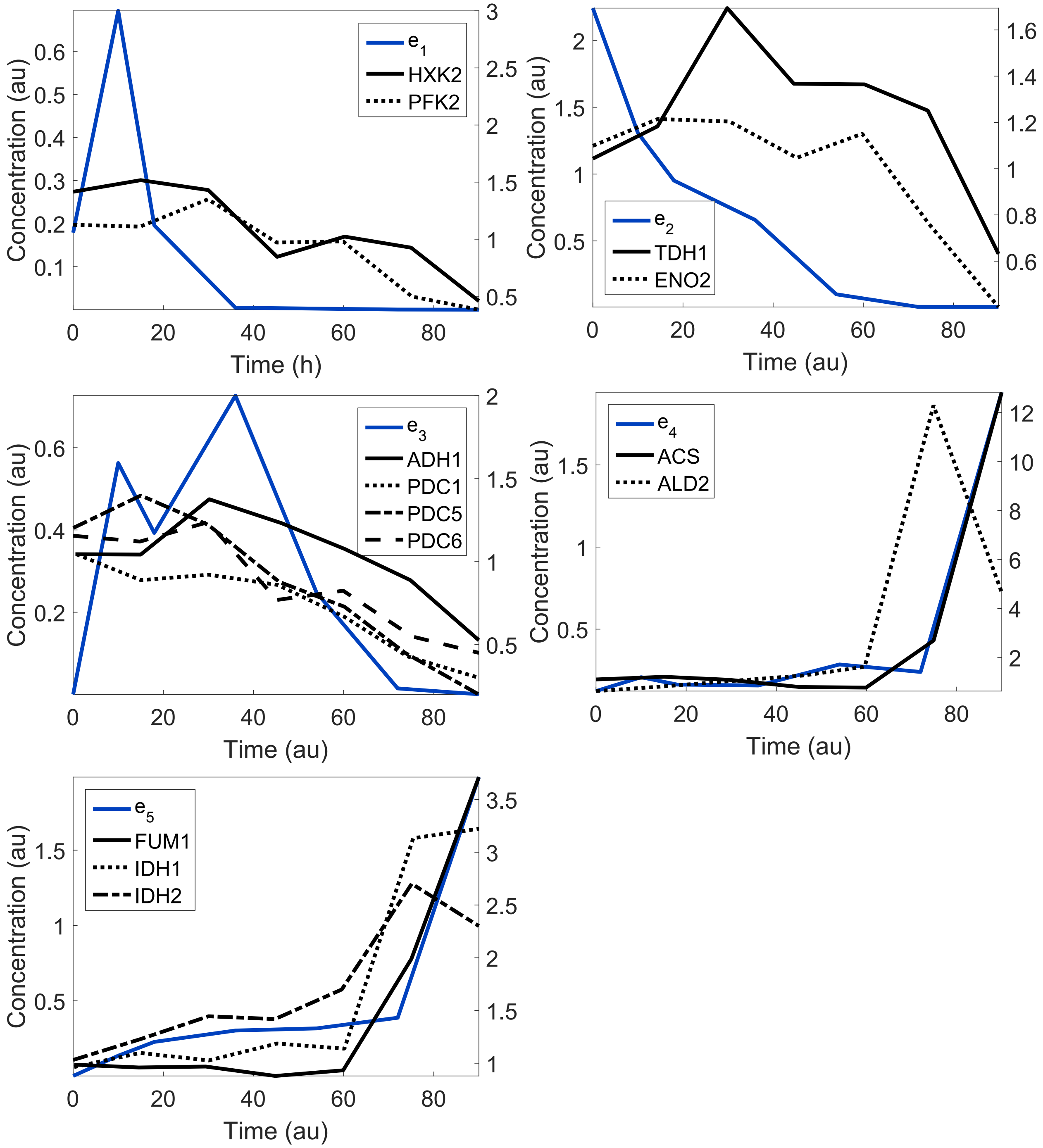


Figure S8: SC: Qualitative comparison of the OCP solution taken with AMIGO2_DO (6PWL) for $t_f = 90$ with real experimental data.

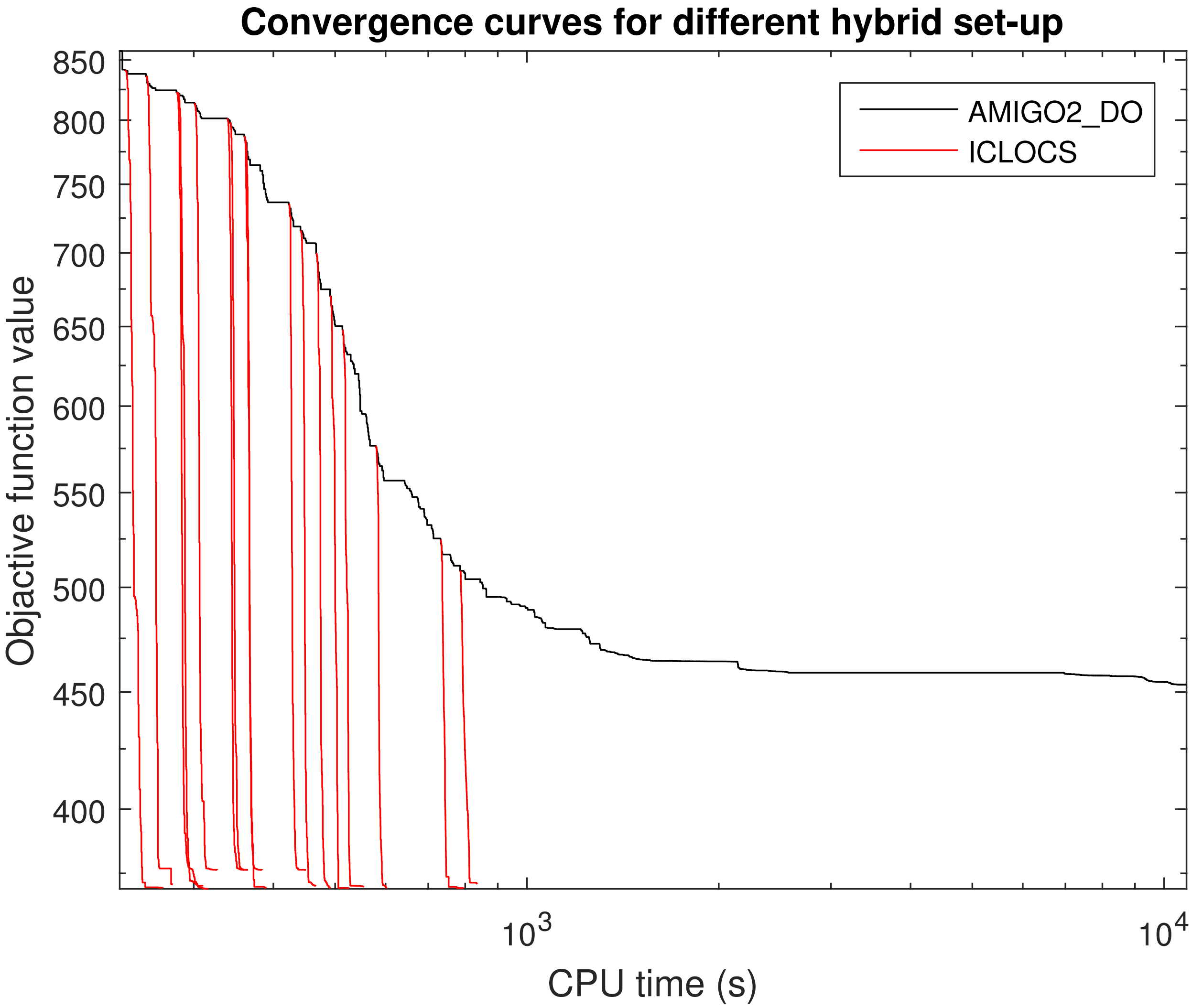


Figure S9: SC: Convergence curves of the hybrid approach for point O on the Pareto front. ICLOCS is initialized from different points of the AMIGO2_DO convergence.

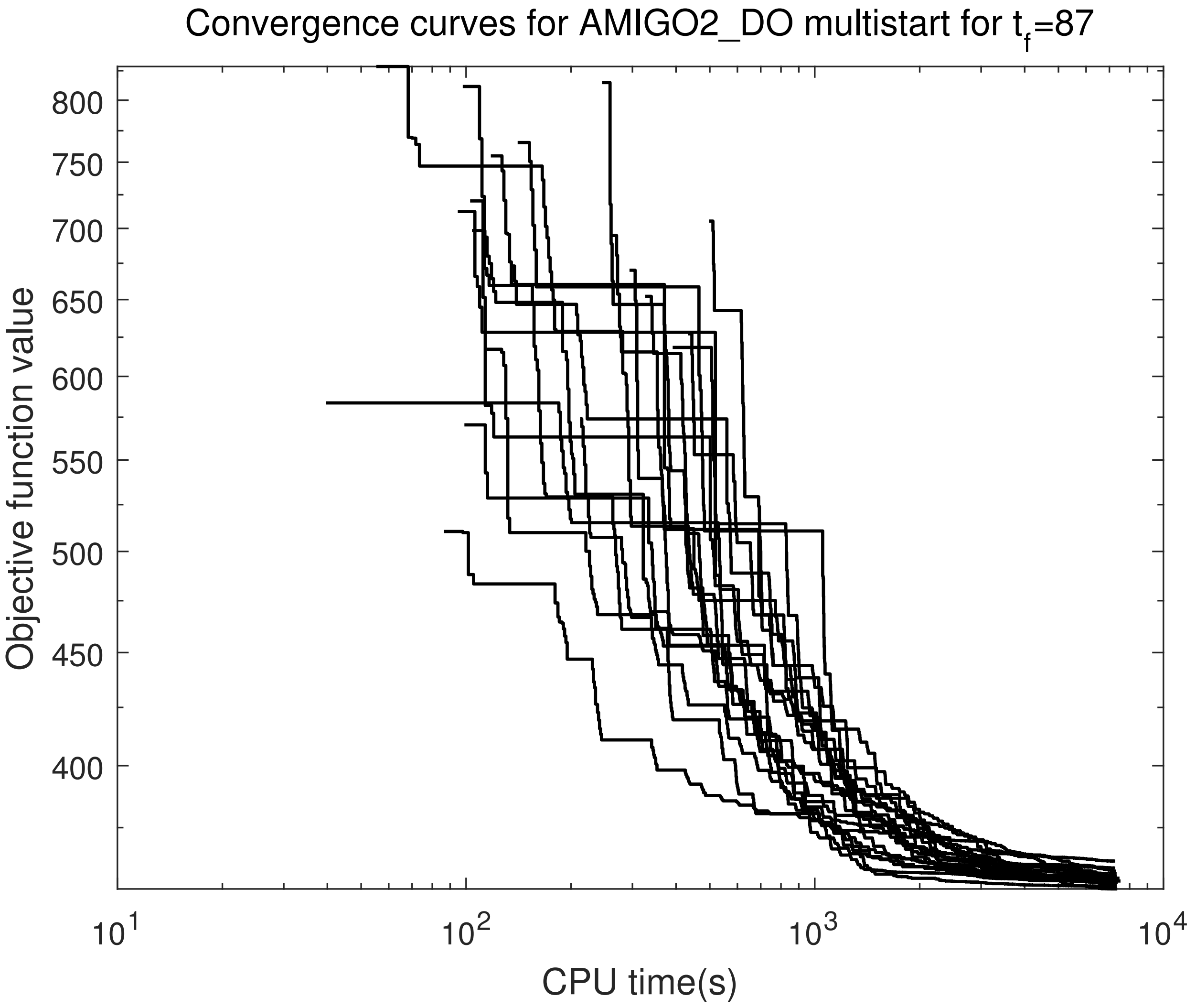


Figure S10: SC: Convergence curves of 20 `AMIGO2_DO` runs for point O on the Pareto front with 6PWL.

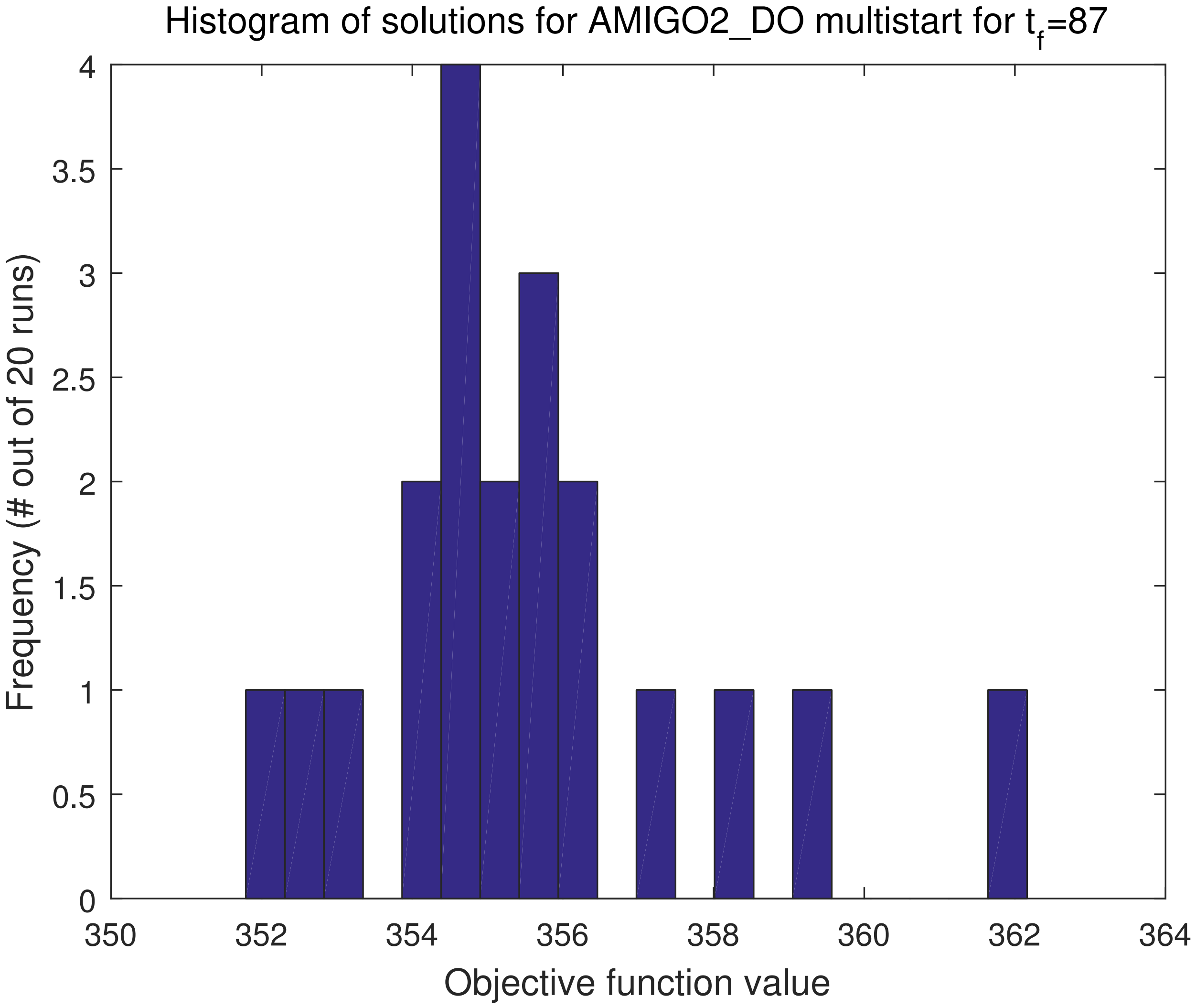


Figure S11: SC: Histogram of solutions from 20 AMIGO2_DO runs for point O on the Pareto front with 6PWL.

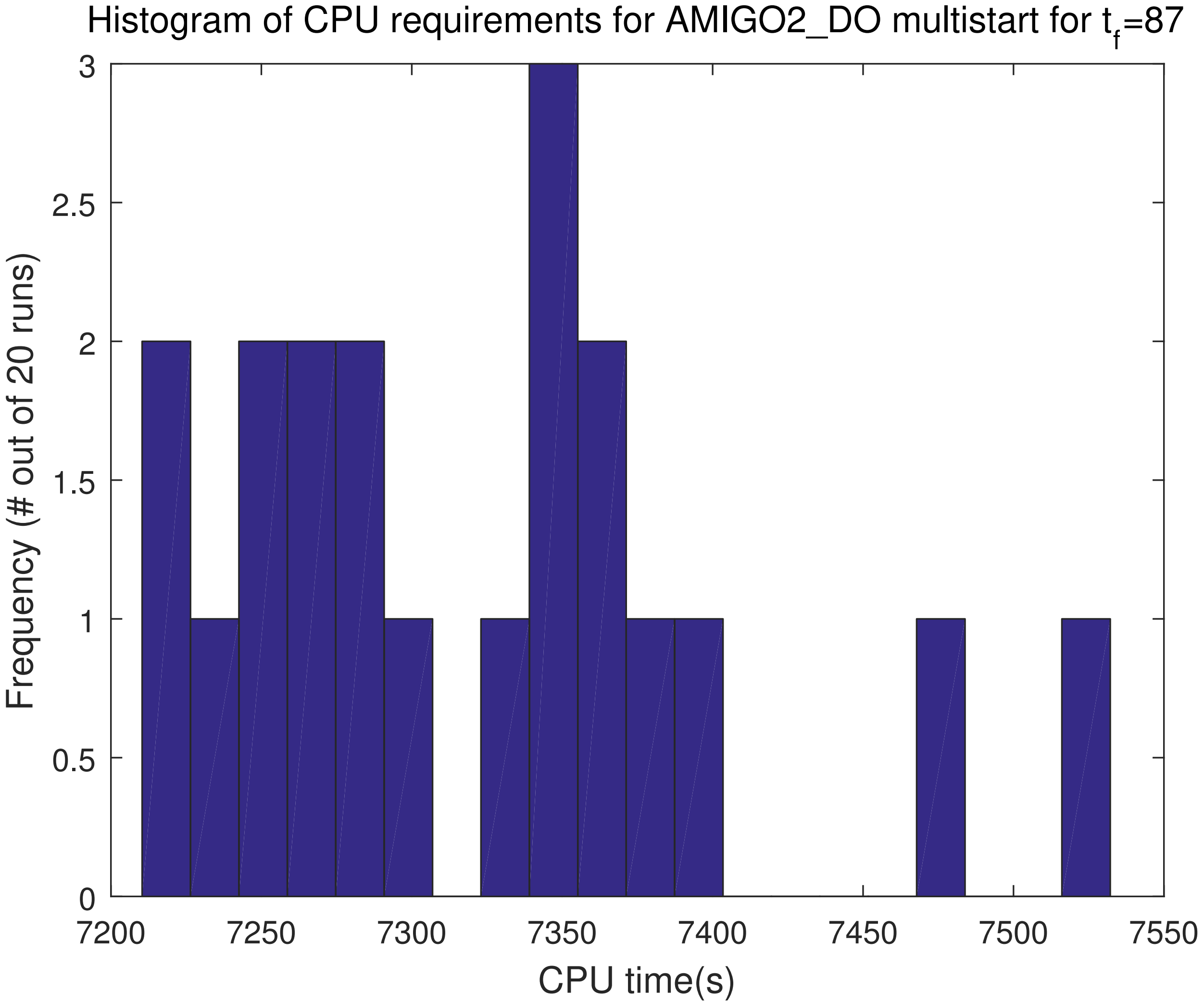


Figure S12: SC: Histogram of CPU requirements from 20 AMIGO2_DO runs for point O on the Pareto front with 6PWL.

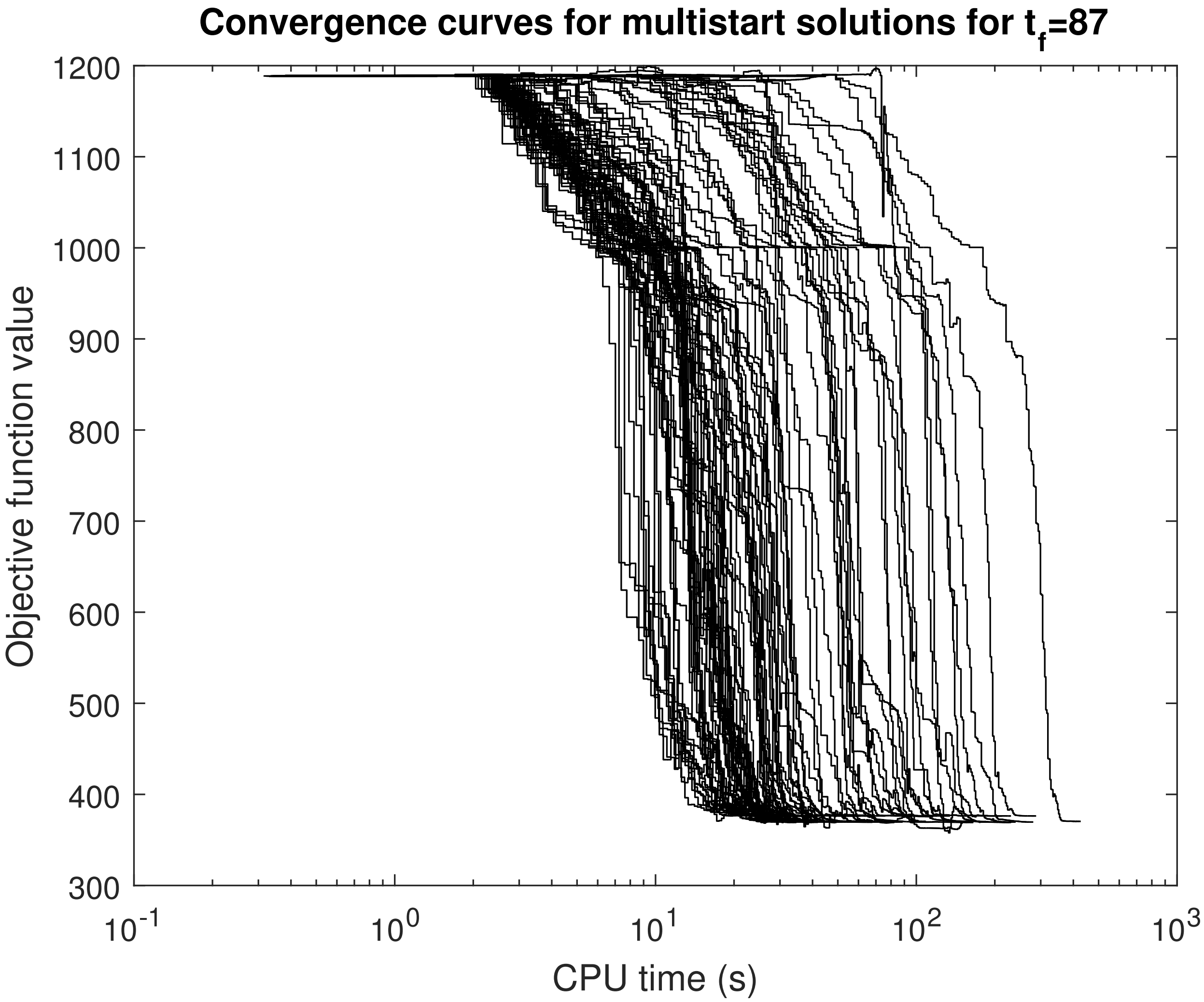


Figure S13: SC: Convergence curves of a multistart (random initialization from 100 points) with `ICLOCS` 10PWC for point O on the Pareto front. However, in contrast with Figure S16, here the dynamics corresponding to the random initial point (10PWC controls) have been simulated and provided to `ICLOCS` as part of the initial guess. In this way, the transcribed constraint in the NLP formulation that correspond to the ODEs is not violated in the initial guess, resulting to a huge improvement in the convergence of the solver.

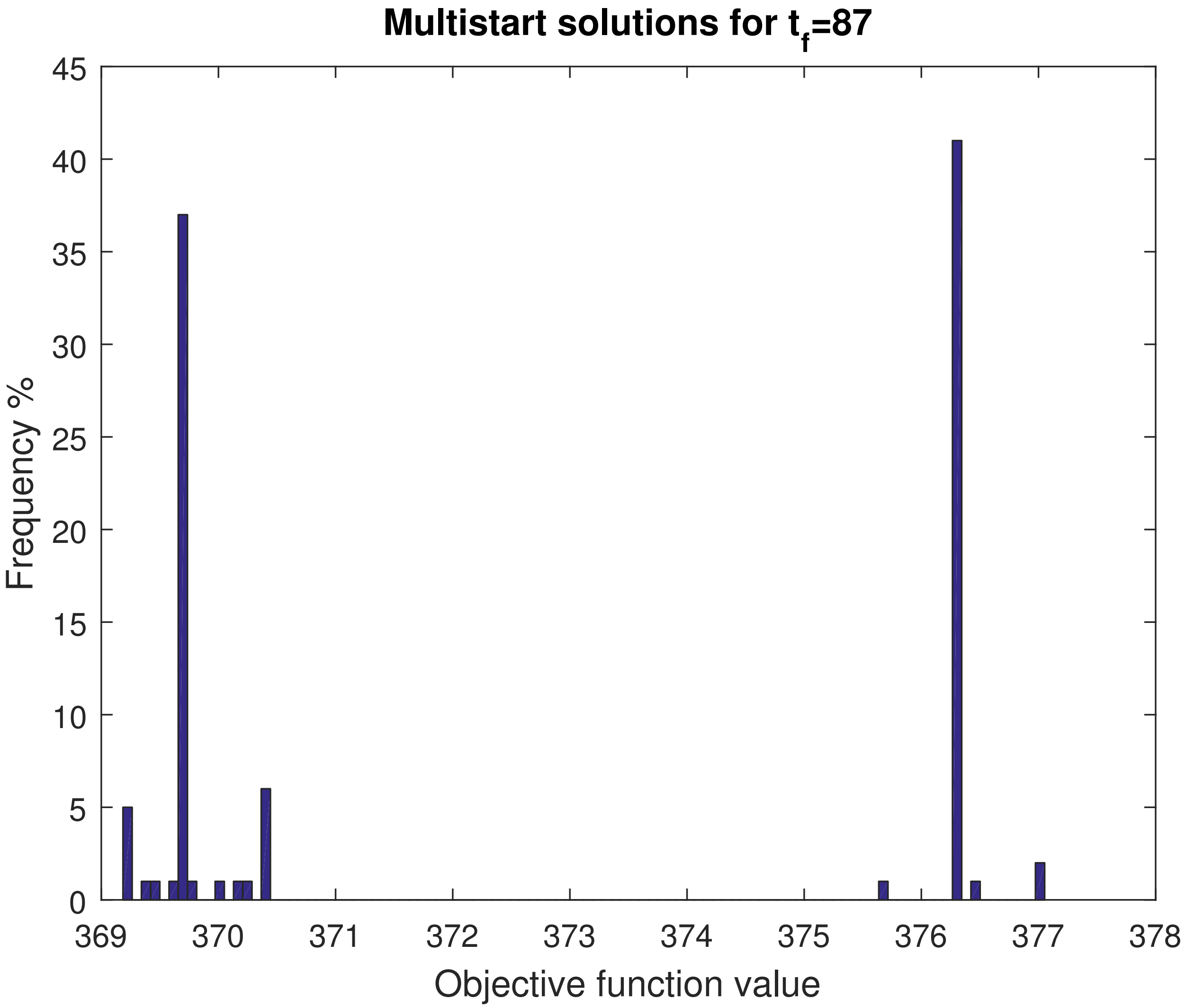


Figure S14: SC: Histogram of the multistart OCP solutions for point O on the Pareto front and a total of 100 `msICLOCS` runs (with 10PWC) initialized randomly in the search space, that correspond to Figure S13.

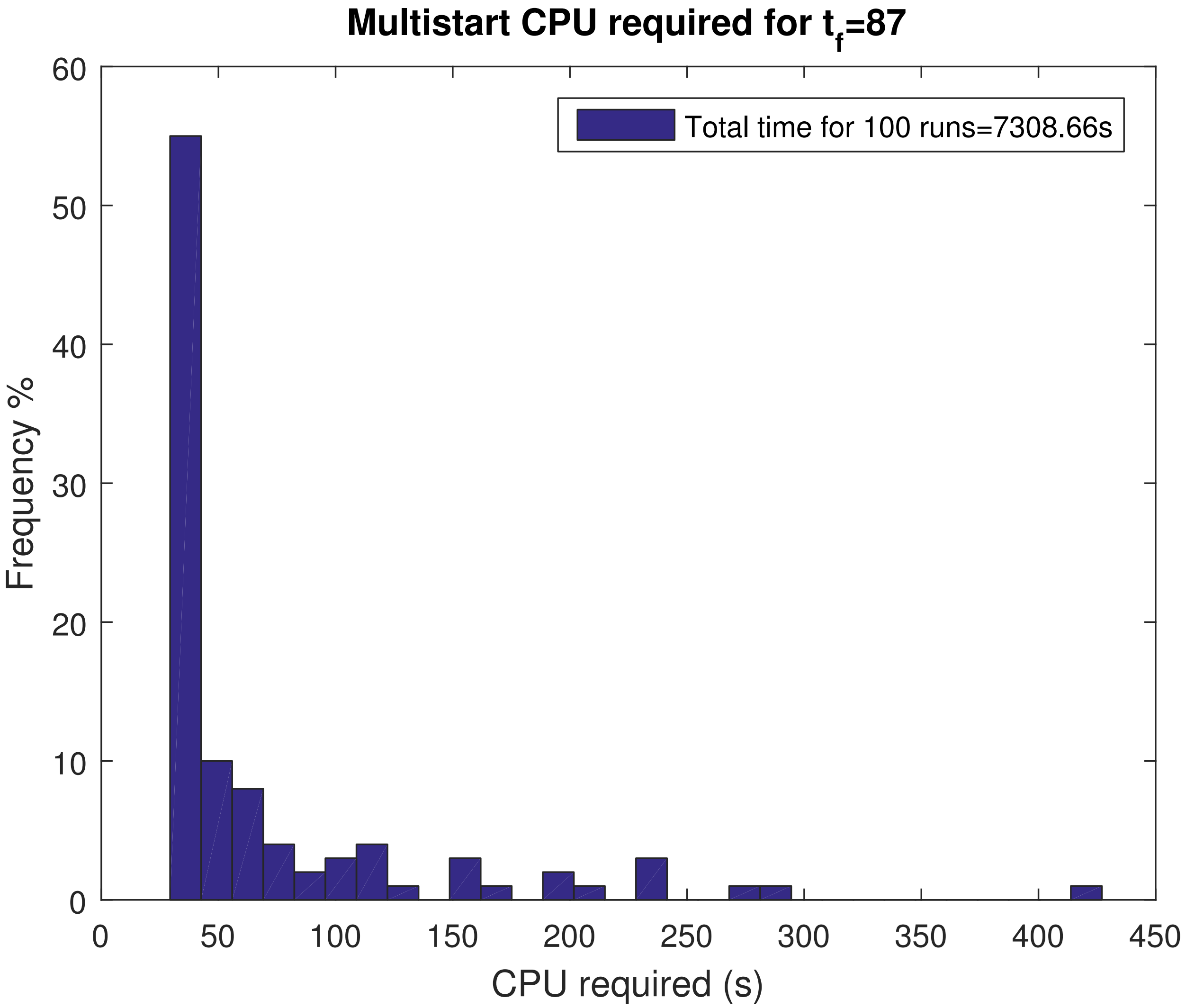


Figure S15: SC: Histogram of the computational time required obtaining the multistart OCP solutions for point O on the Pareto front and a total of 100 `msICLOCS` runs (with 10PWC) initialized randomly in the search space, that correspond to Figure S13.

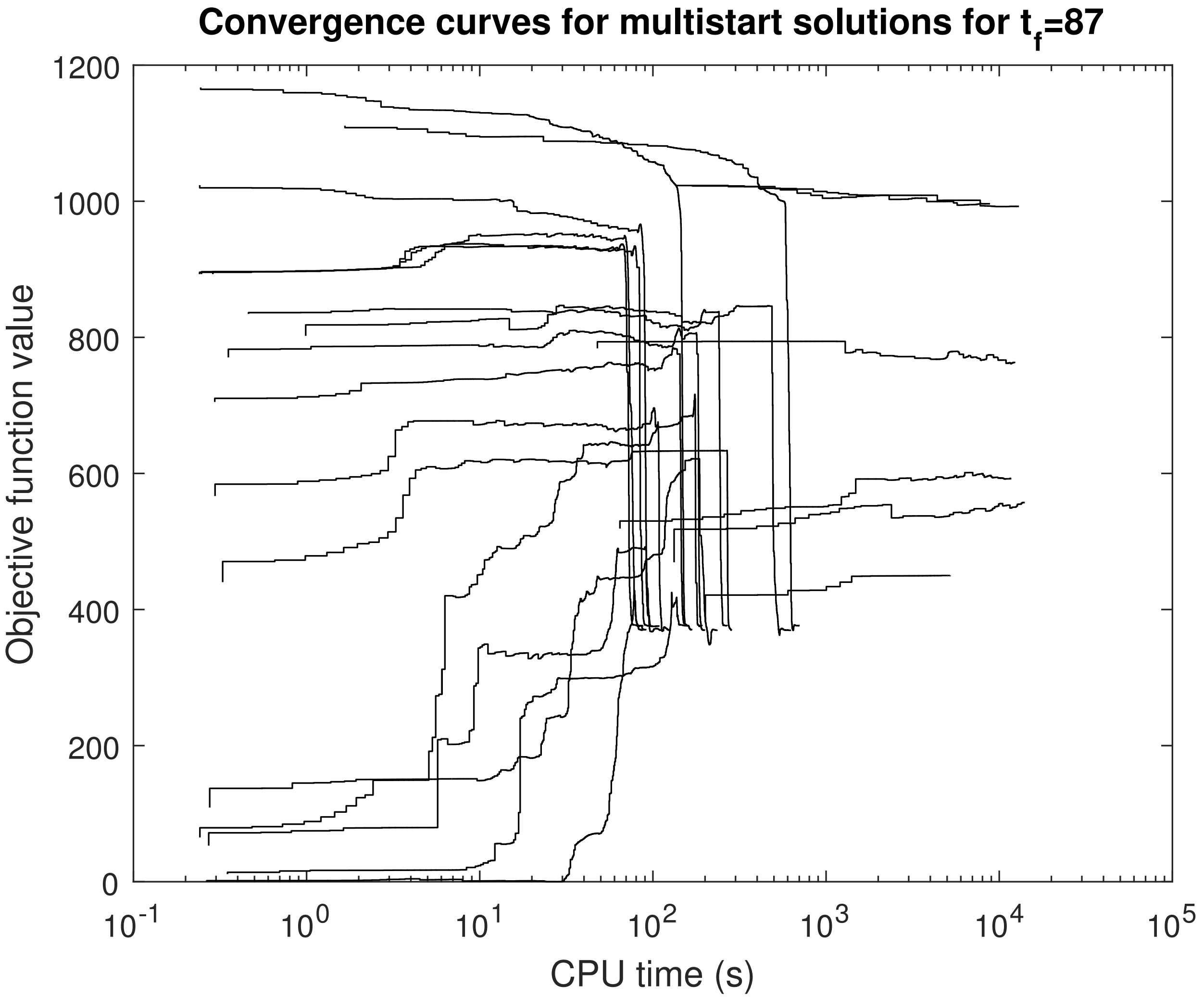


Figure S16: SC: Convergence curves of a multistart (random initialization from 100 points) with `ICLOCS` 10PWC for point O on the Pareto front. Here the dynamics corresponding to the random initial point (10PWC controls) were not simulated and were not provided to `msICLOCS` as part of the initial guess. The initial guess provided was only the 10PWC random controls. Note that out of the 25 runs, two crashed and several others failed to converge.

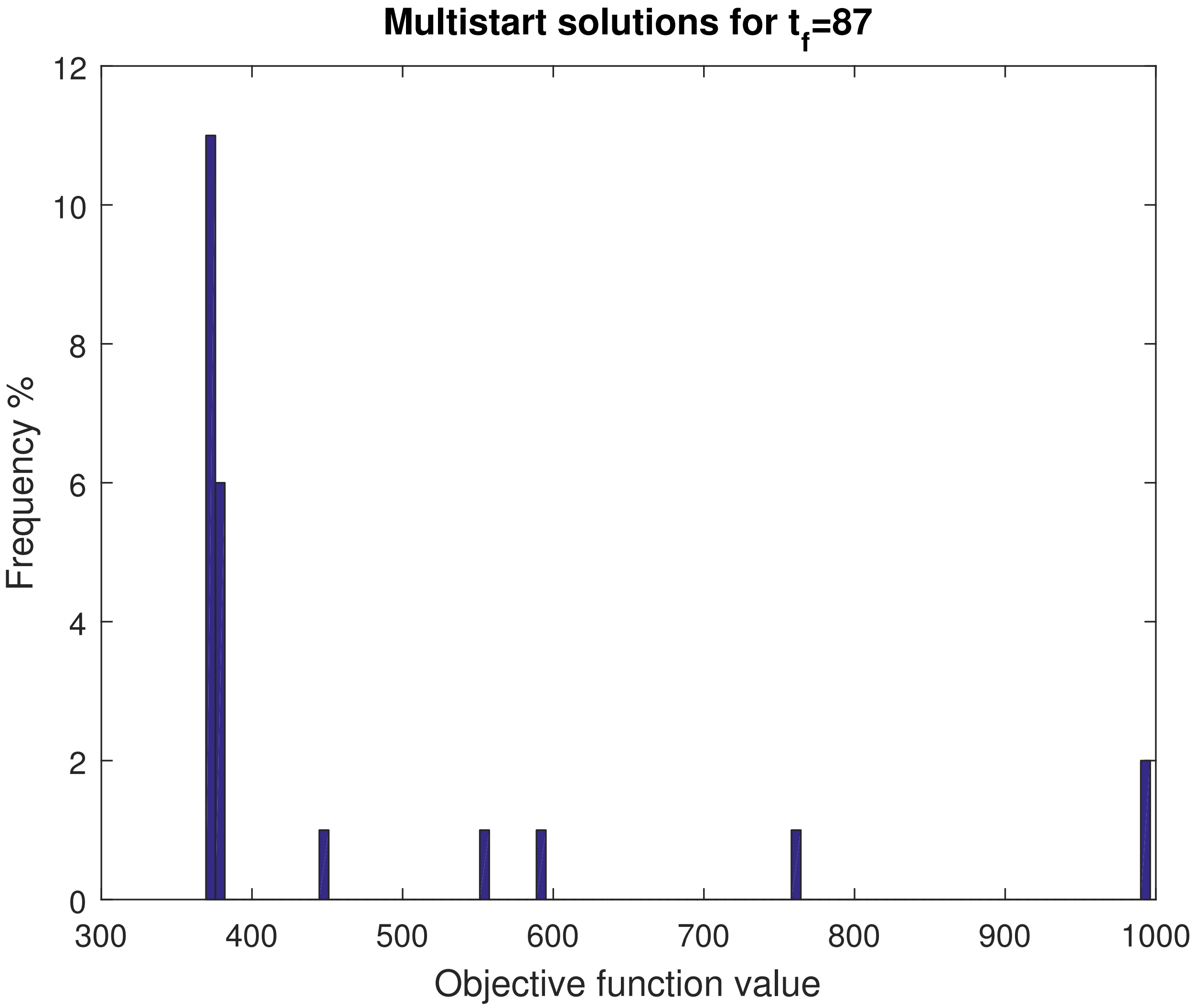


Figure S17: SC: Histogram of the multistart OCP solutions for point O on the Pareto front and a total of 100 `msICLOCS` runs (with 10PWC) initialized randomly in the search space, that correspond to Figure S16. Note that out of the 25 runs, two crashed and several others failed to converge.

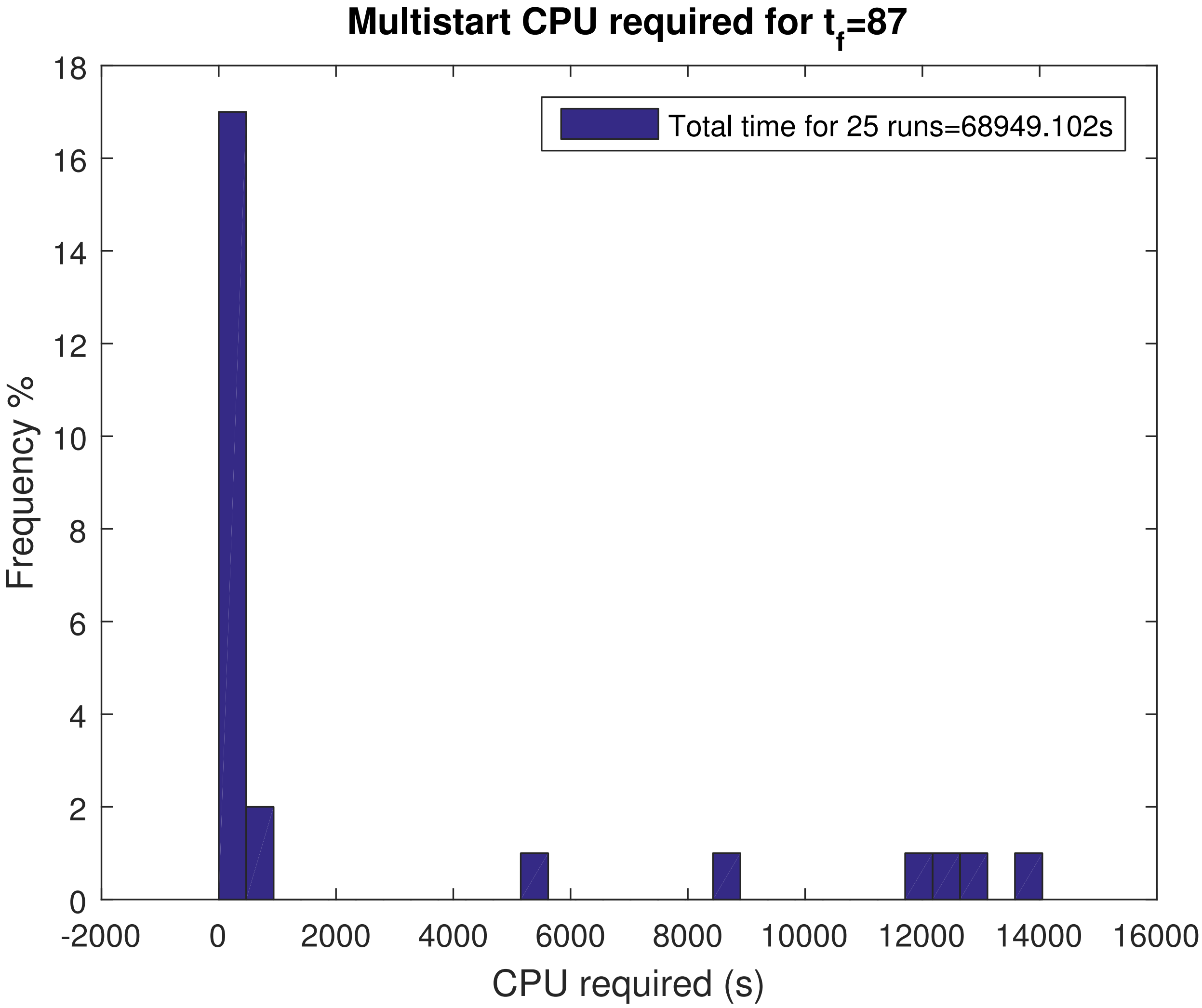


Figure S18: SC: Histogram of the computational time required obtaining the multistart OCP solutions for point O on the Pareto front and a total of 100 `msICLOCS` runs (with 10PWC) initialized randomly in the search space, that correspond to Figure S16. Note that out of the 25 runs, two crashed and several others failed to converge.

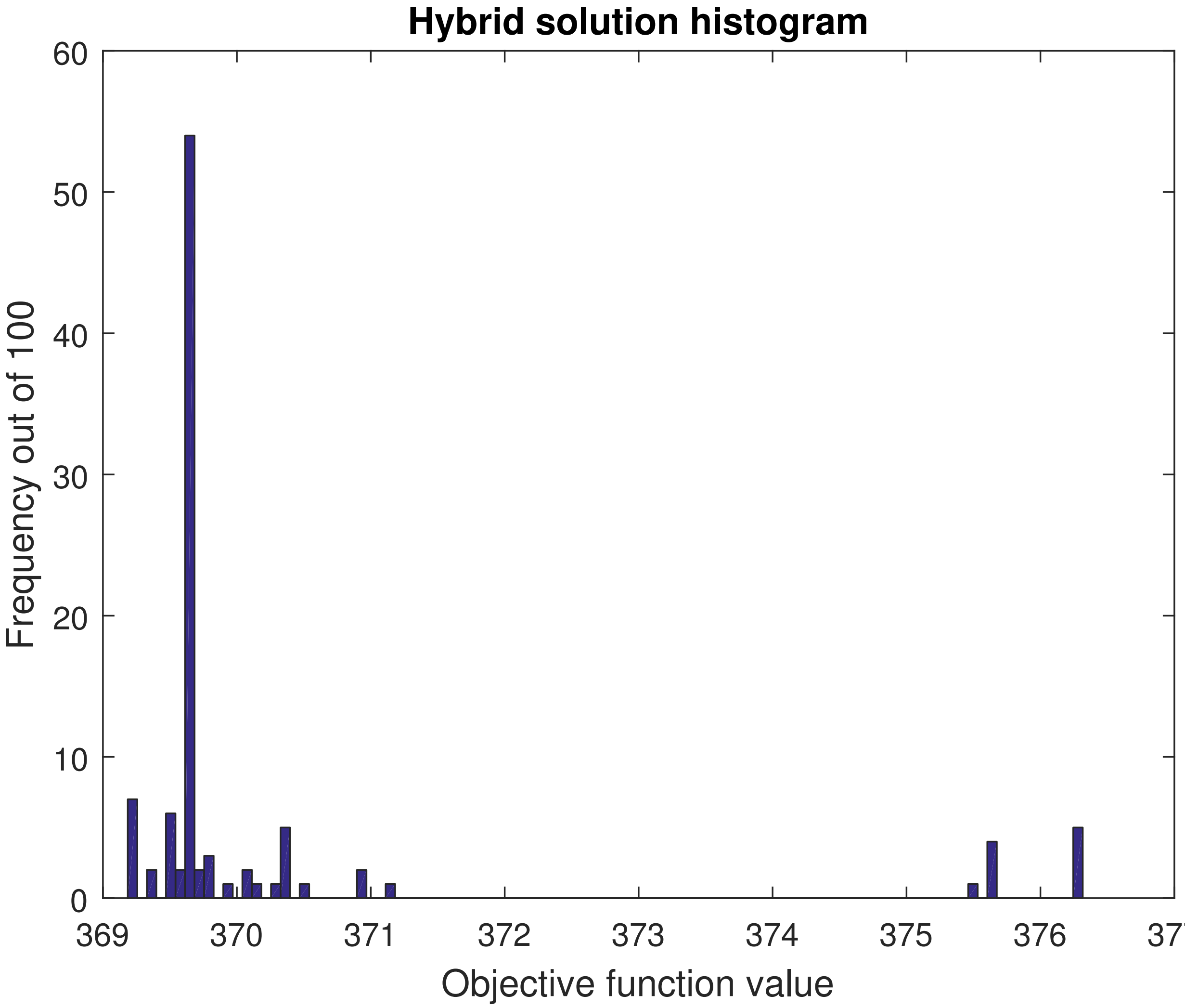


Figure S19: SC: Histogram of the multiple AMIGO2_DO+ICLOCS runs for point O on the Pareto front. Starting from different random initial points and performing a fast hybrid run, we attempt to illustrate the possible practical multiplicity of solutions in this scenario. All final solutions below 371 belong to a 0.5% of the solution range and can be considered practically equivalent.

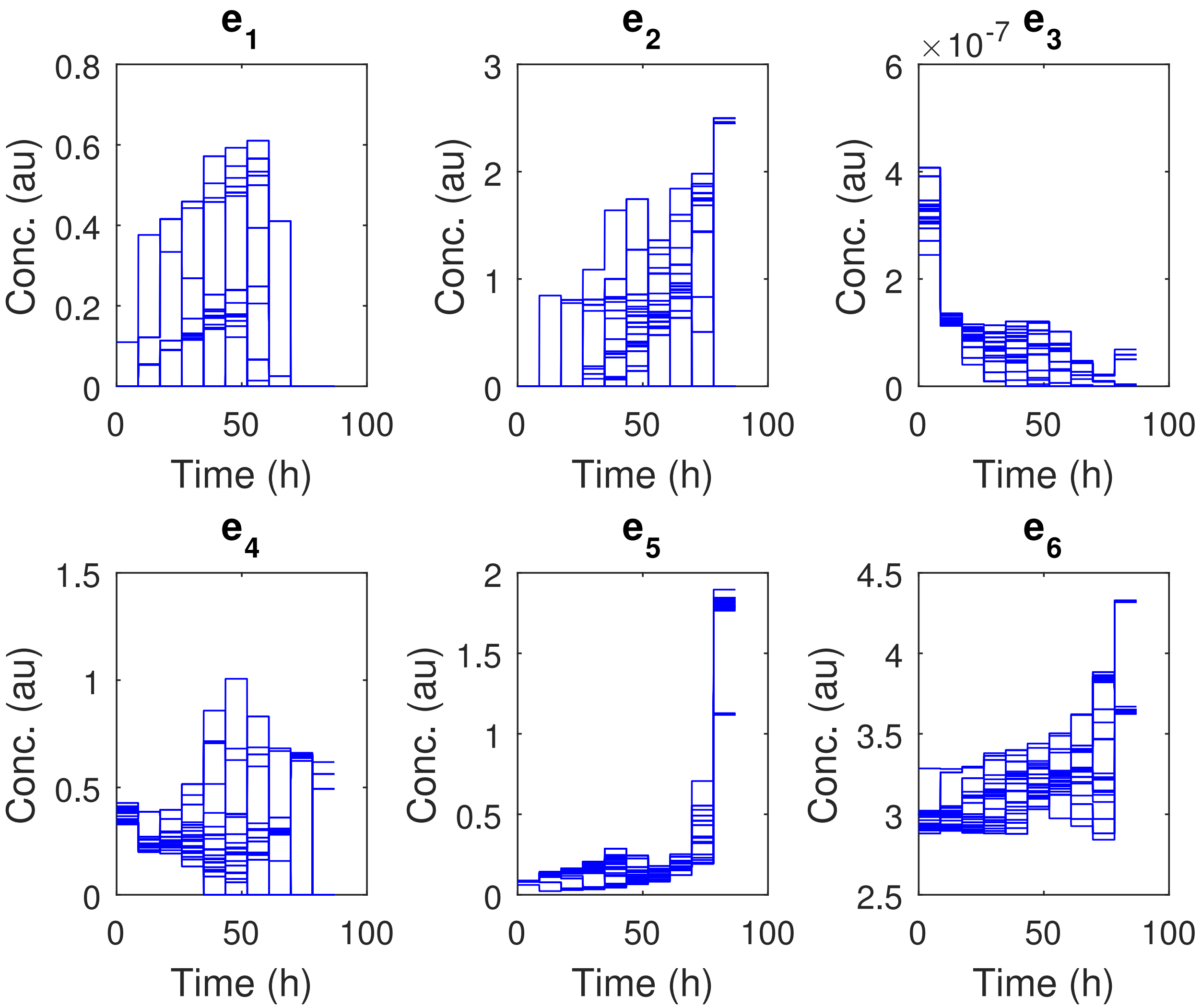


Figure S20: SC: All optimal controls obtained from the 100 `AMIGO2_DO+ICLOCS` runs for point O on the Pareto front that are shown in Figure S19.

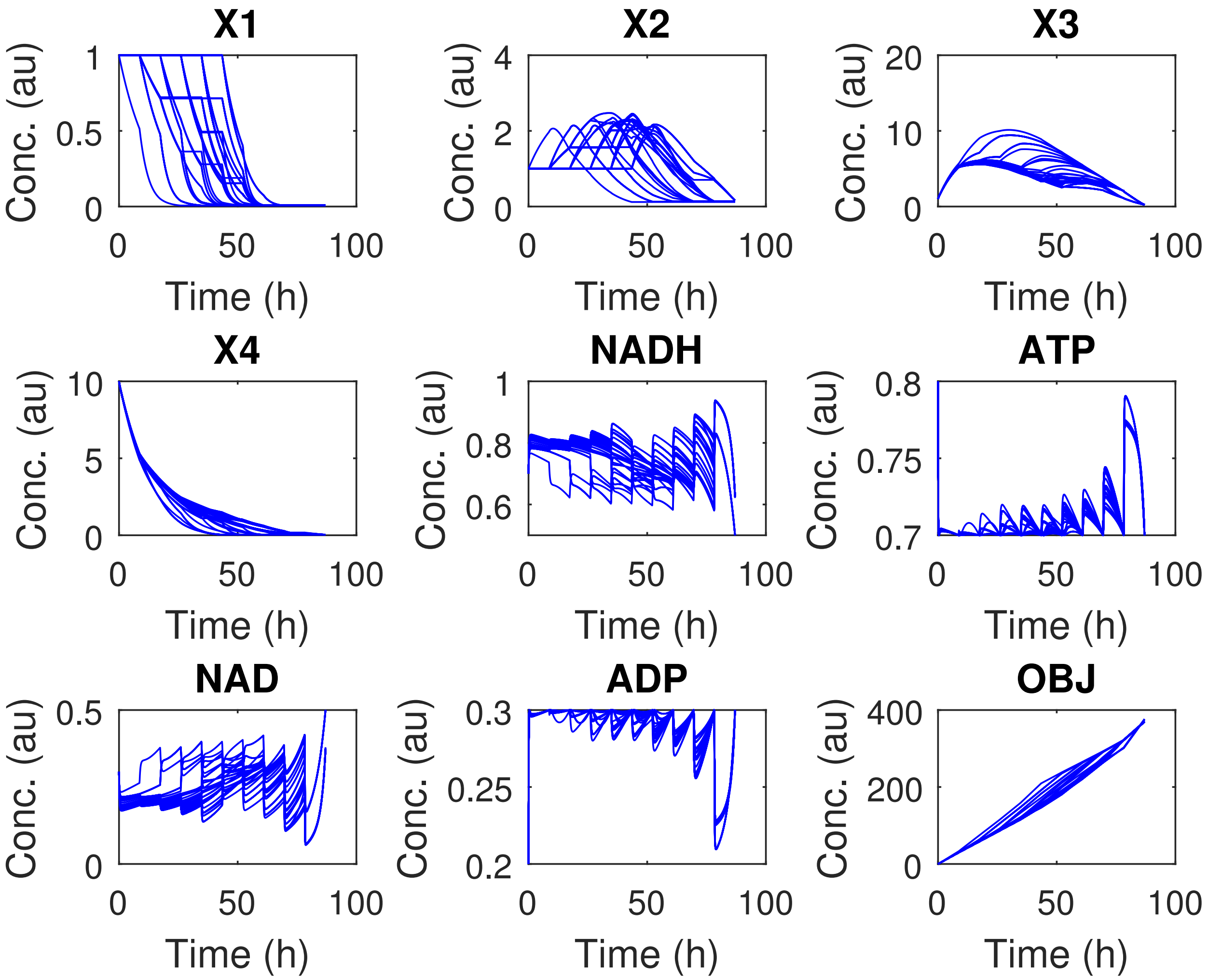


Figure S21: SC: All optimal state trajectories obtained from the 100 `AMIGO2_DO+ICLOCS` runs for point O on the Pareto front that are shown in Figure S19.

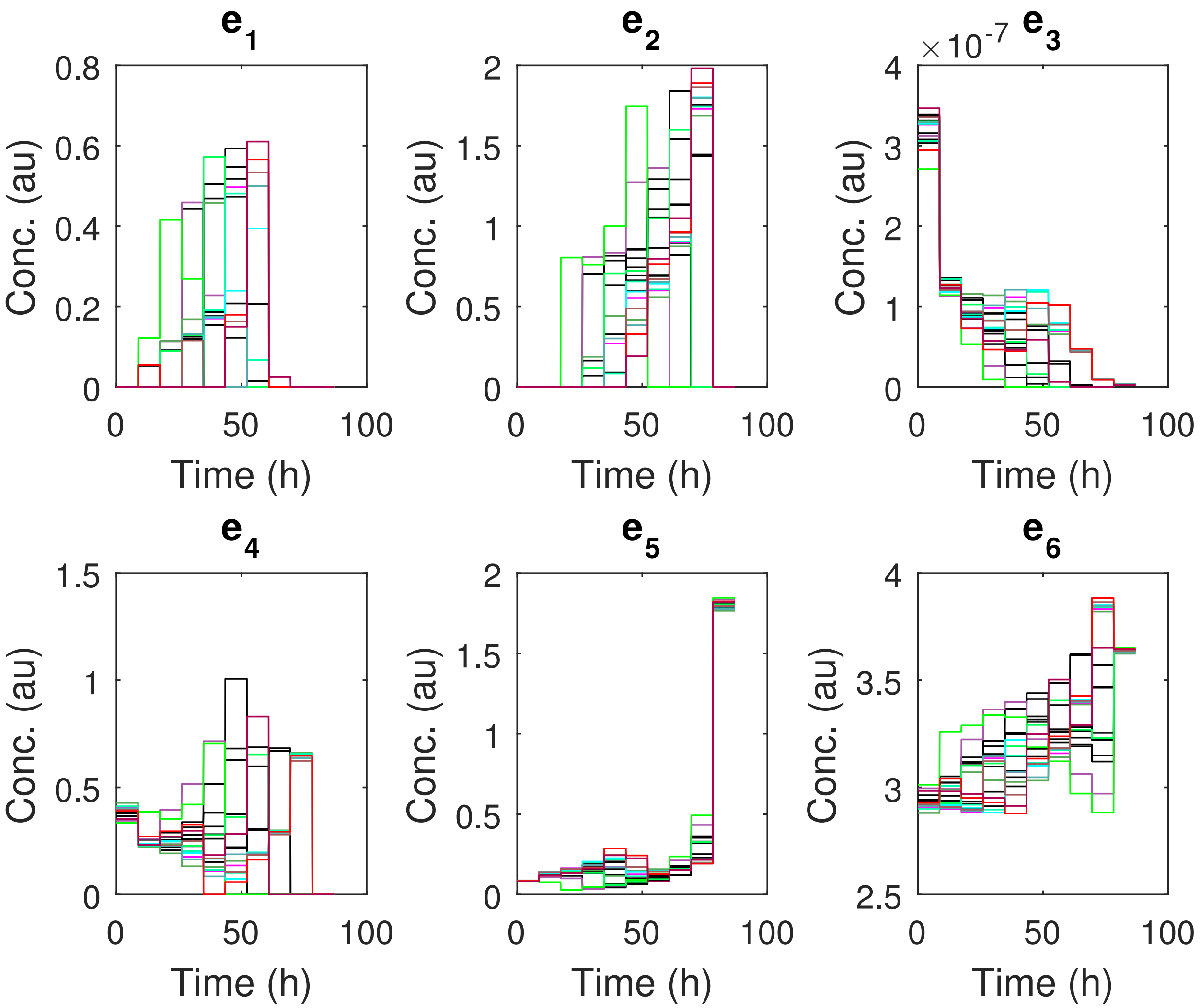


Figure S22: Comparison of the optimal controls for point O on the Pareto front 100 `AMIGO2_DO+ICLOCS` runs, in SC case study. All these solutions are identified in Figure S19 as practically equivalent in terms of the objective function value, falling below the 0.5% threshold ($\leq 371.04$).

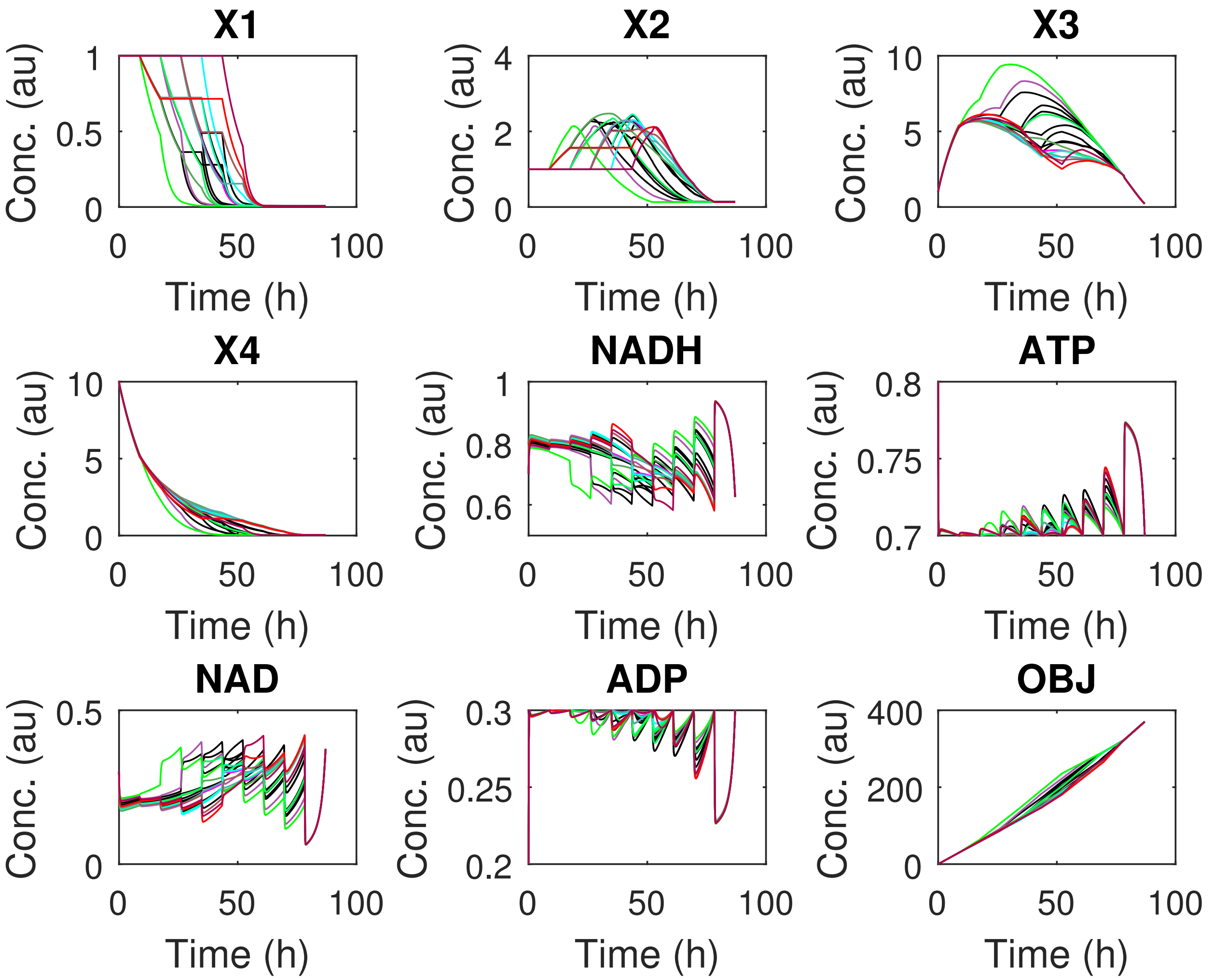


Figure S23: Comparison of the optimal state trajectories for point O on the Pareto front 100 `AMIGO2_DO+ICLOCS` runs, in SC case study. All these solutions are identified in Figure S19 as practically equivalent in terms of the objective function value, falling below the 0.5% threshold ($\leq 371.04$).

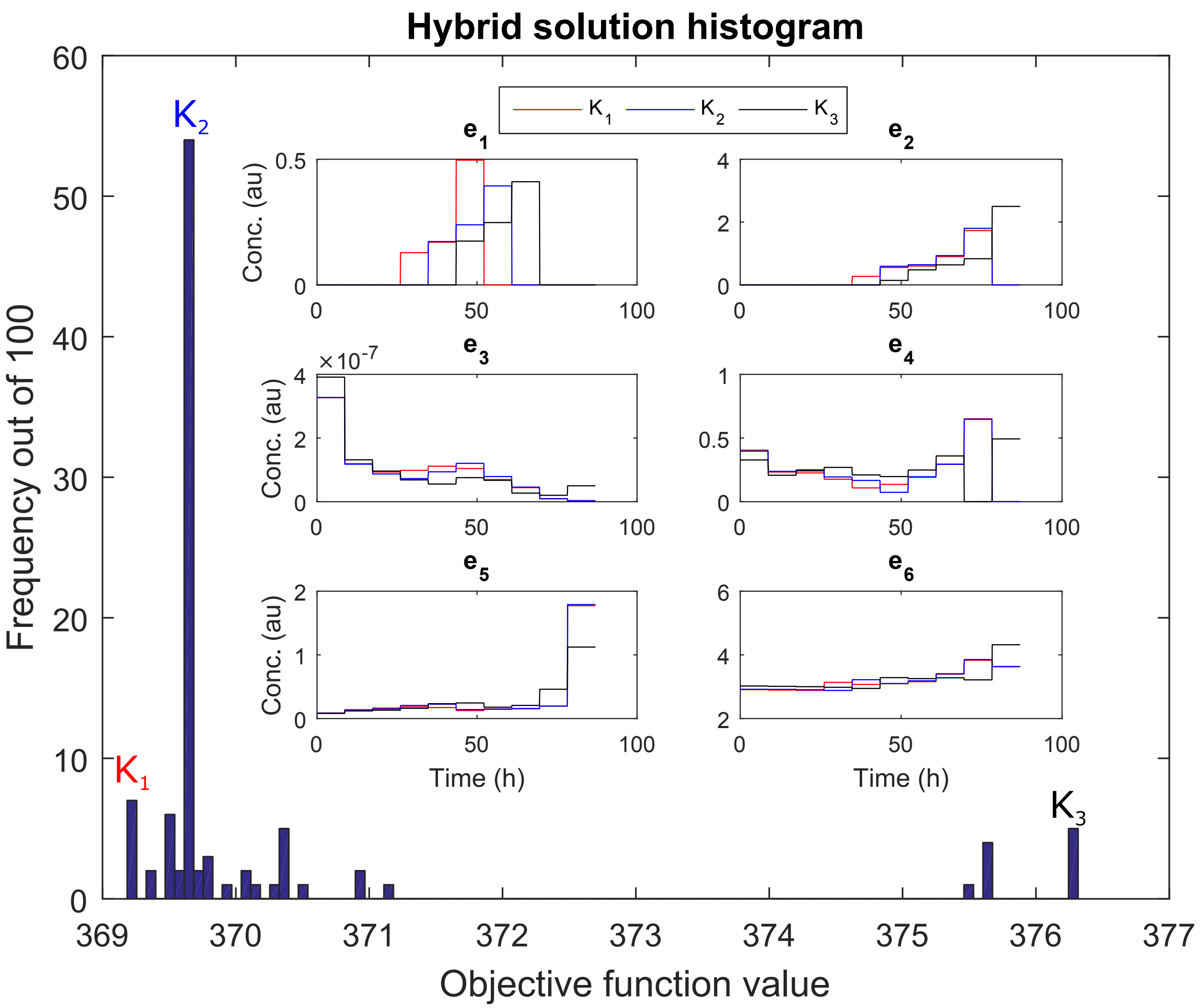


Figure S24: SC: A comparison of the optimal controls that correspond to three solutions taken with `msICLOCS` 10PWC for point O on the Pareto front, that correspond to Figures S19. The trajectories in pink correspond to the global optimal solution with objective function value around 369.2 while the ones in cyan and blue correspond to two local solutions with objective function values around 369.7 and 376.3 respectively.

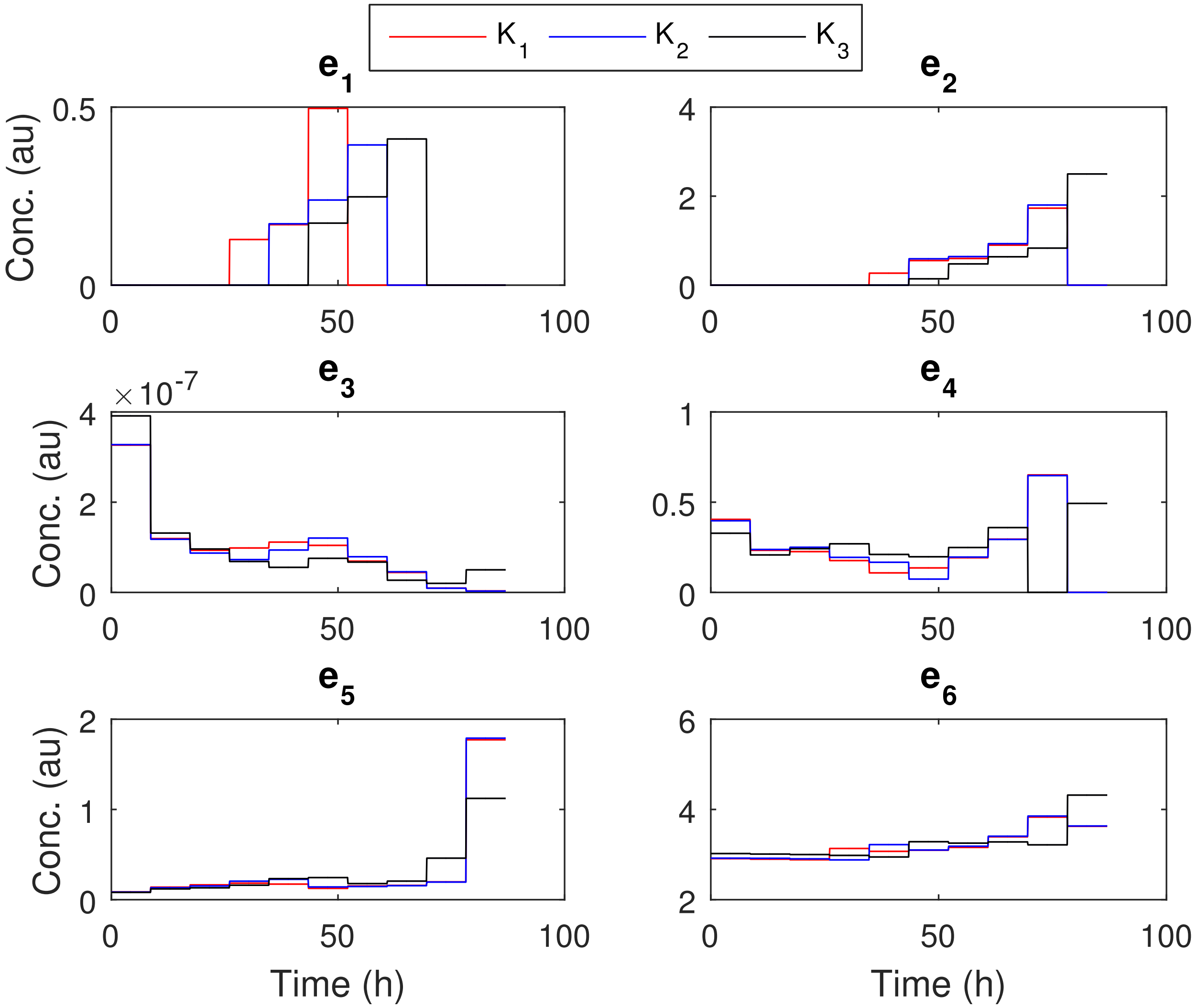


Figure S25: SC: A comparison of the optimal controls that correspond to three solutions taken with `msICLOCS` 10PWC for point O on the Pareto front, that correspond to Figures S19. The trajectories in pink correspond to the global optimal solution with objective function value around 369.2 while the ones in cyan and blue correspond to two local solutions with objective function values around 369.7 and 376.3 respectively.

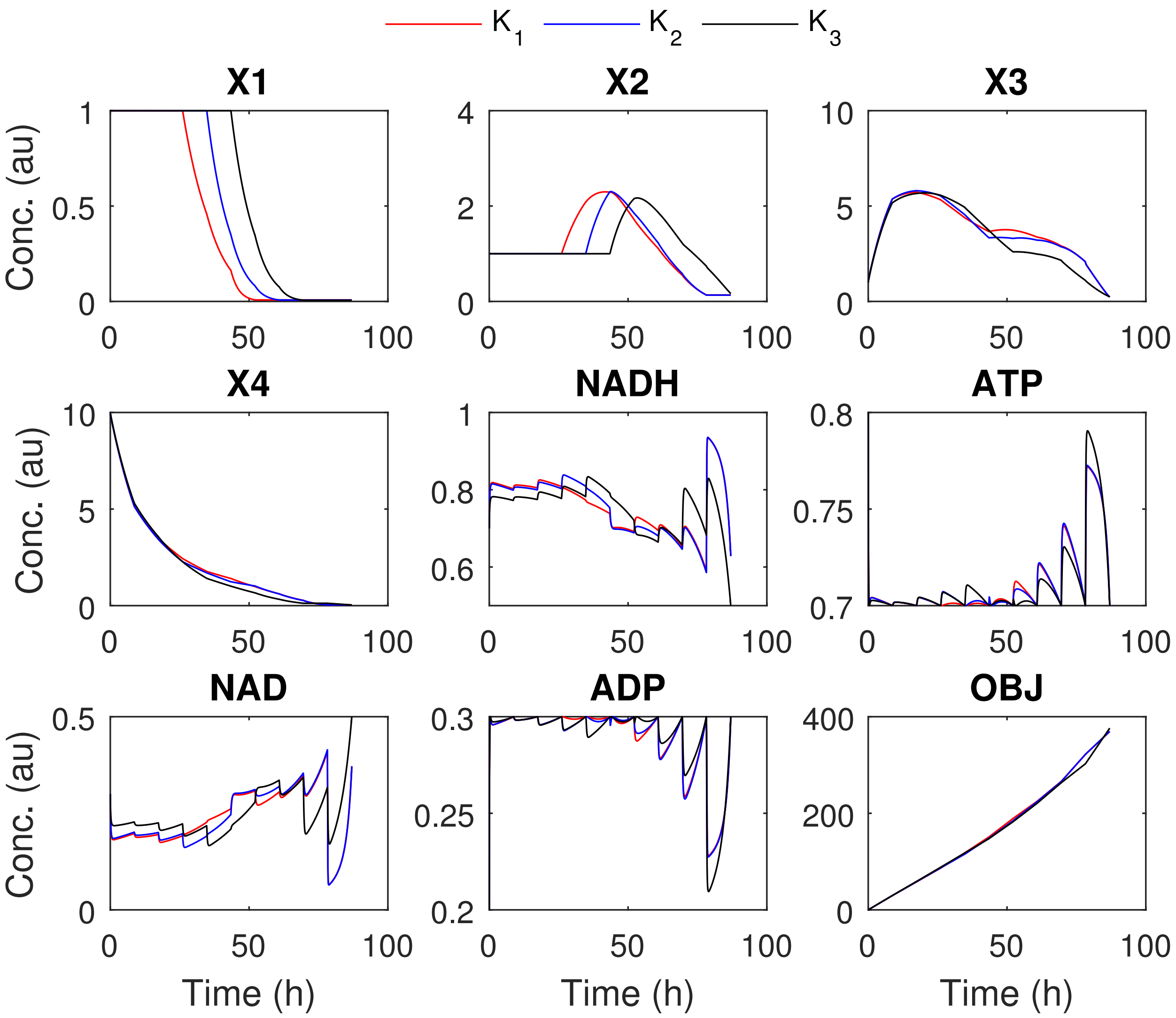


Figure S26: SC: A comparison of the optimal state trajectories that correspond to three solutions taken with msICLOCS 10PWC for point O on the Pareto front, that correspond to Figures S19. The trajectories in pink correspond to the global optimal solution with objective function value around 369.2 while the ones in cyan and blue correspond to two local solutions with objective function values around 369.7 and 376.3 respectively.

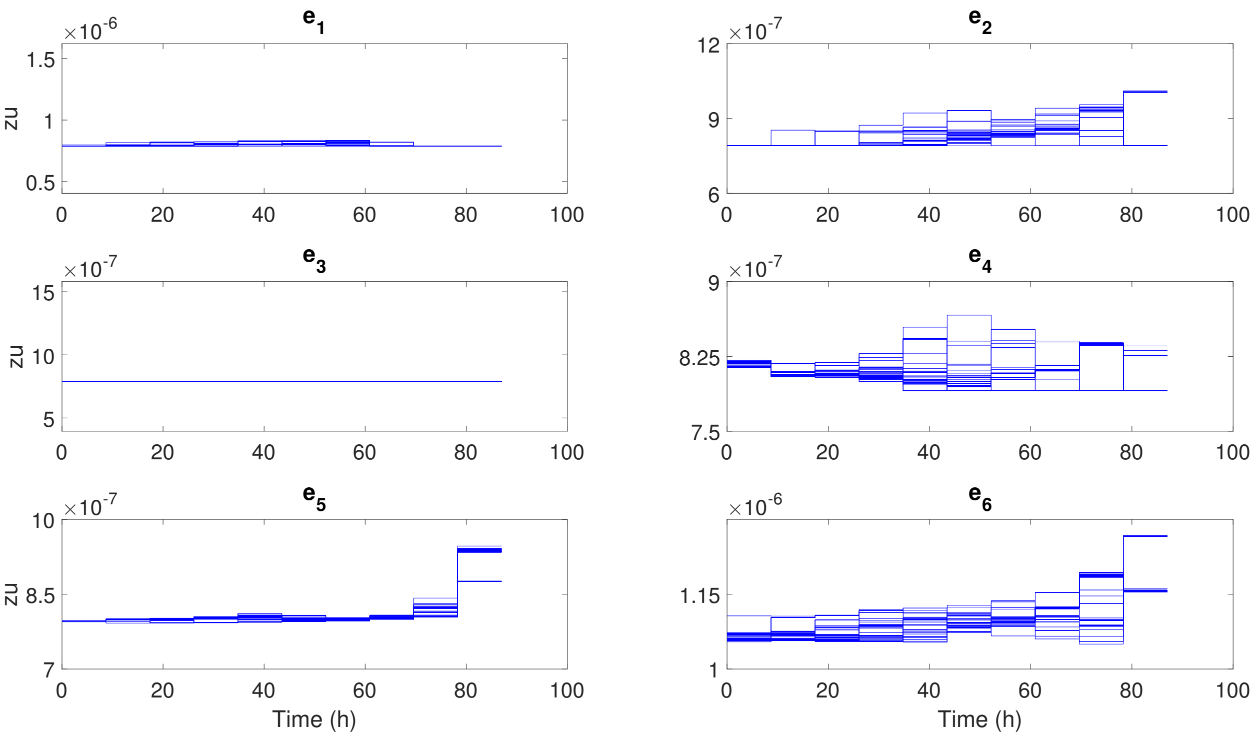


Figure S27: SC: All Lagrange multipliers on the controls' upper bounds obtained from the 100 `AMIGO2_DO+ICLOCS` runs for point O on the Pareto front that are shown in Figure S19.

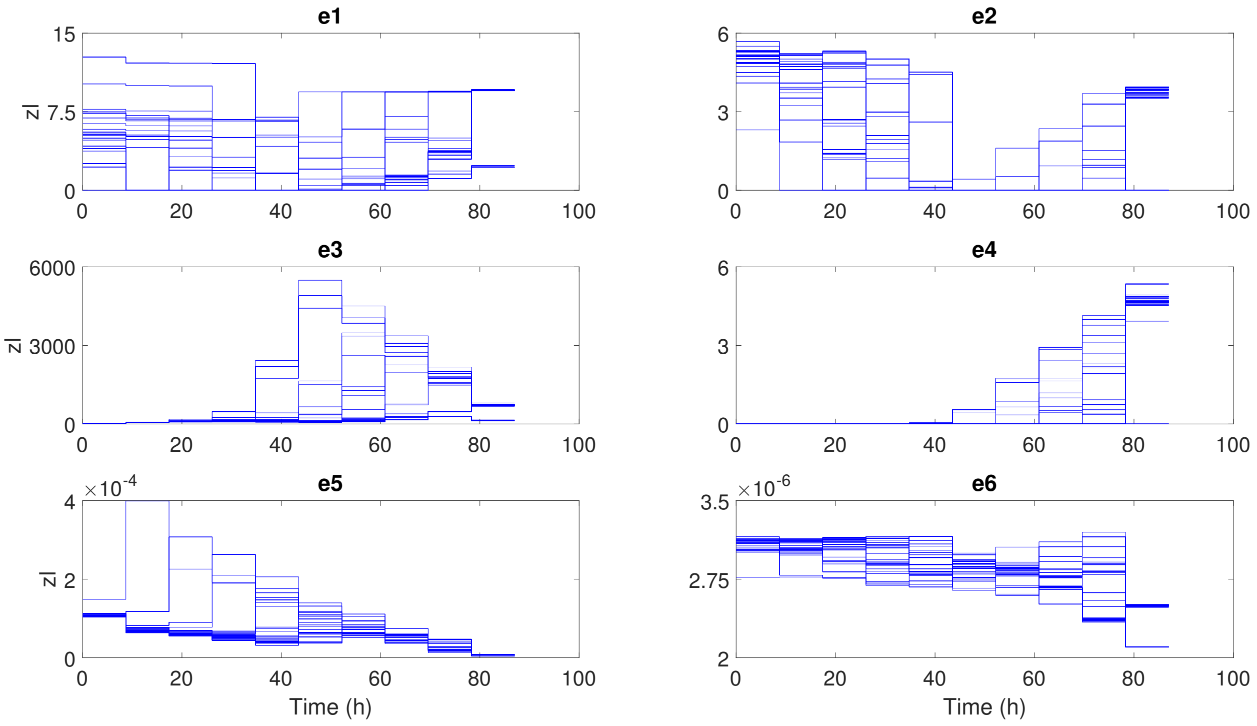


Figure S28: SC: All Lagrange multipliers on the controls' lower bounds obtained from the 100 `AMIGO2_DO+ICLOCS` runs for point O on the Pareto front that are shown in Figure S19.

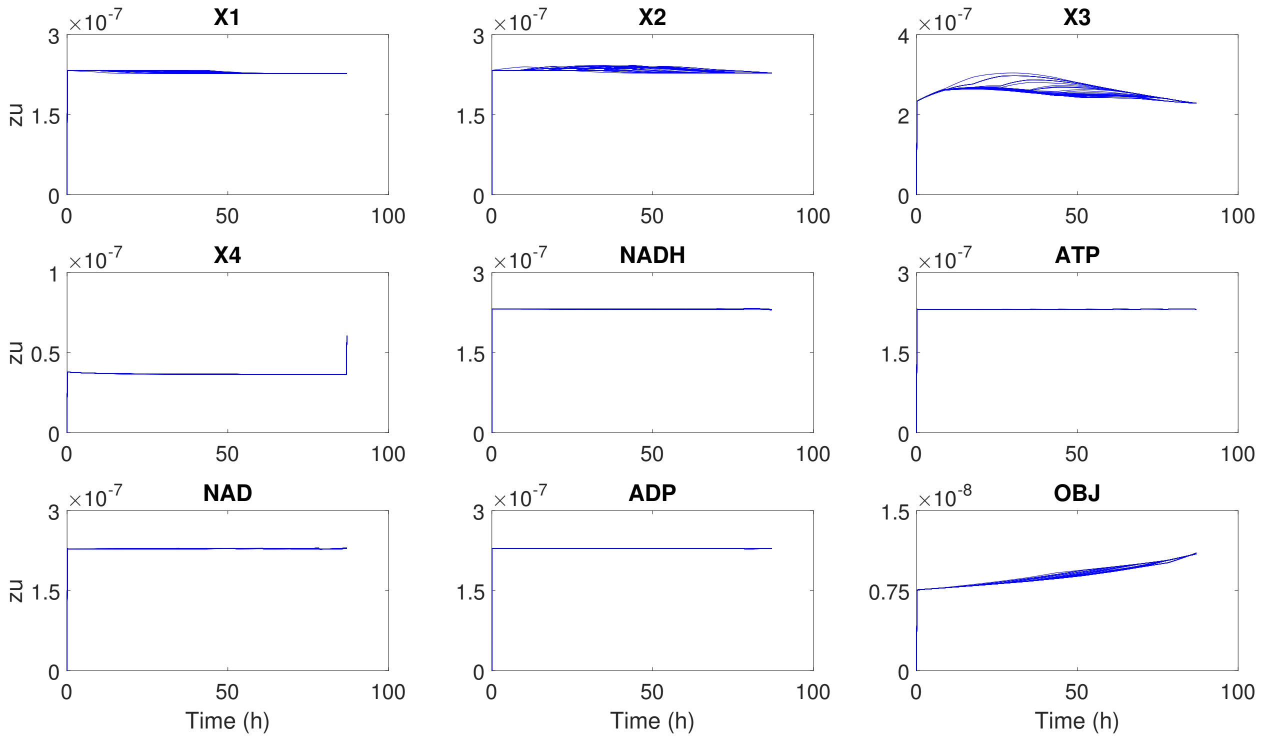


Figure S29: SC: All Lagrange multipliers on the states' upper bounds obtained from the multiple AMIGO2_DO+ICLOCS runs for point O on the Pareto front in Figure S19.

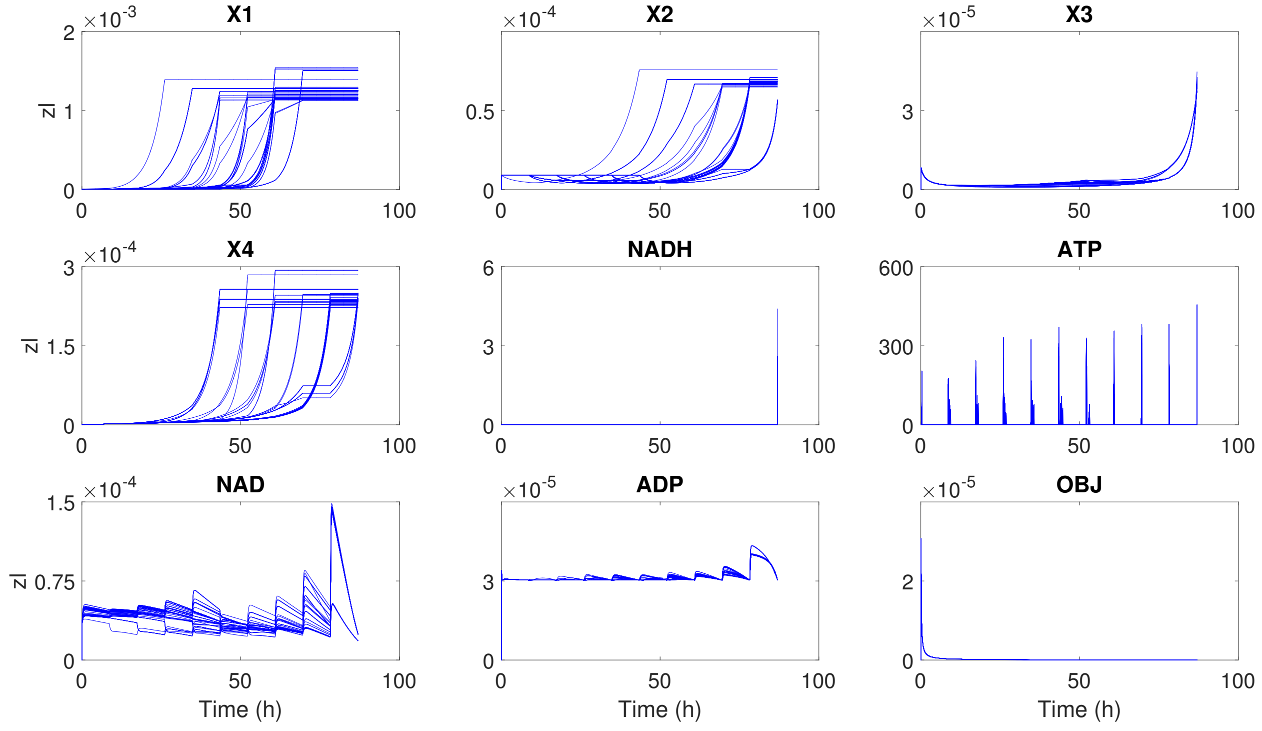


Figure S30: SC: All Lagrange multipliers on the states' lower bounds obtained from the 100 AMIGO2_DO+ICLOCS runs for point O on the Pareto front that are shown in Figure S19.

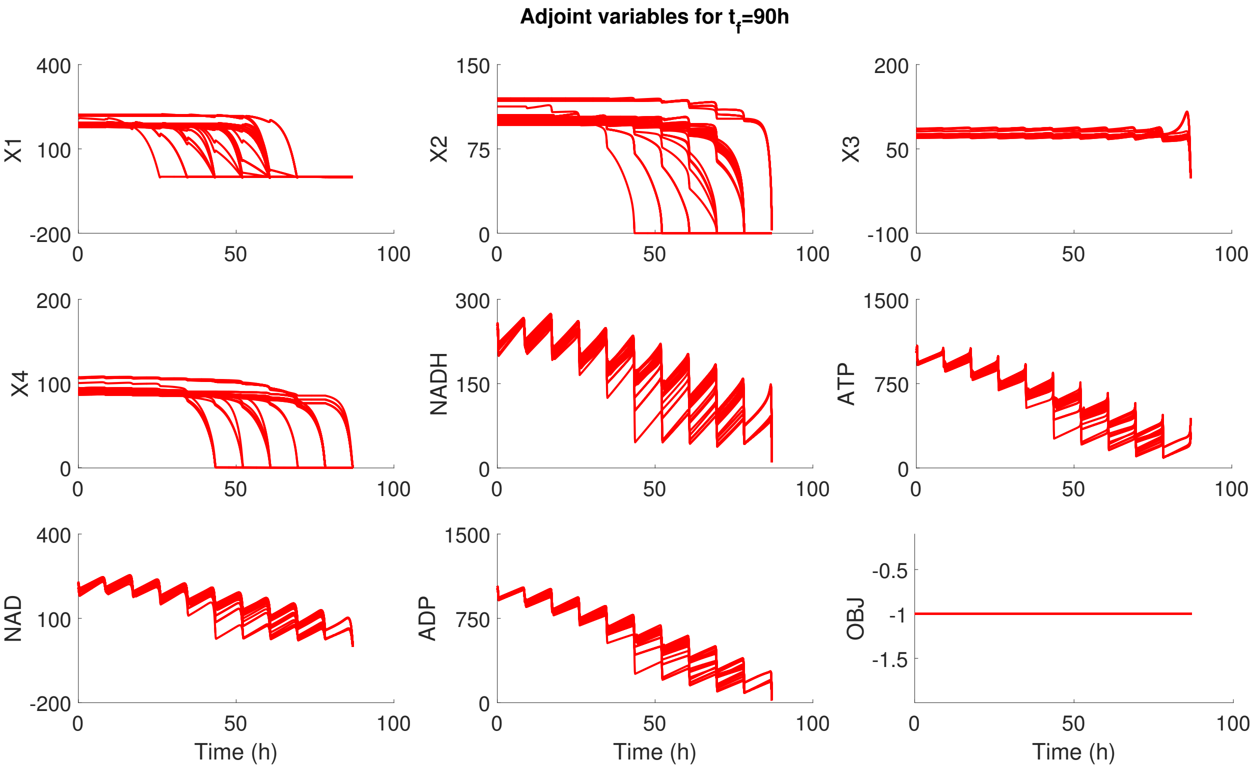


Figure S31: SC: All Lagrange multipliers on the states obtained from the 100 `AMIGO2_DO+ICLOCS` runs for point O on the Pareto front that are shown in Figure S19.

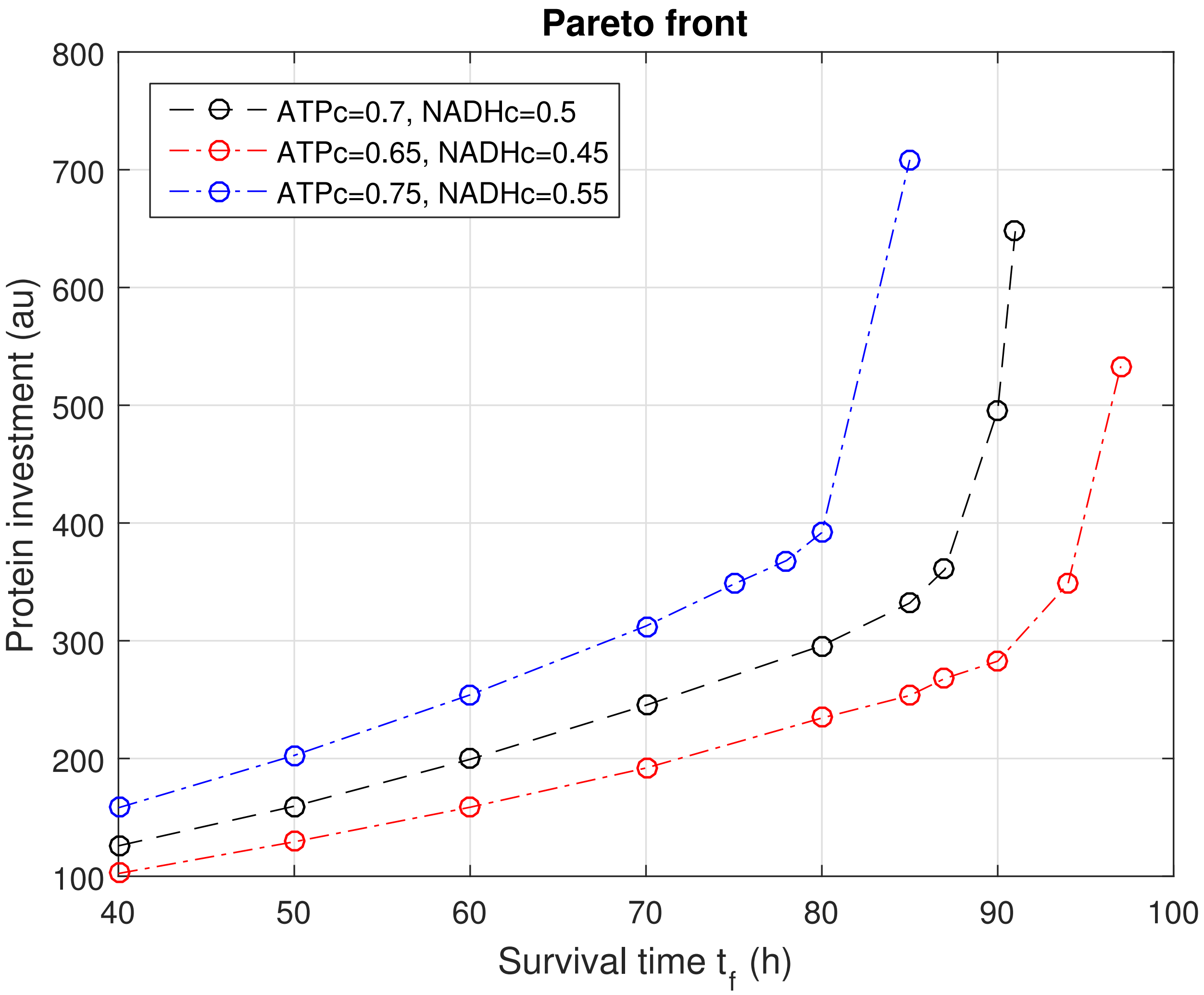


Figure S32: Comparison of the resulting Pareto front in SC case study for various co-factor critical values.

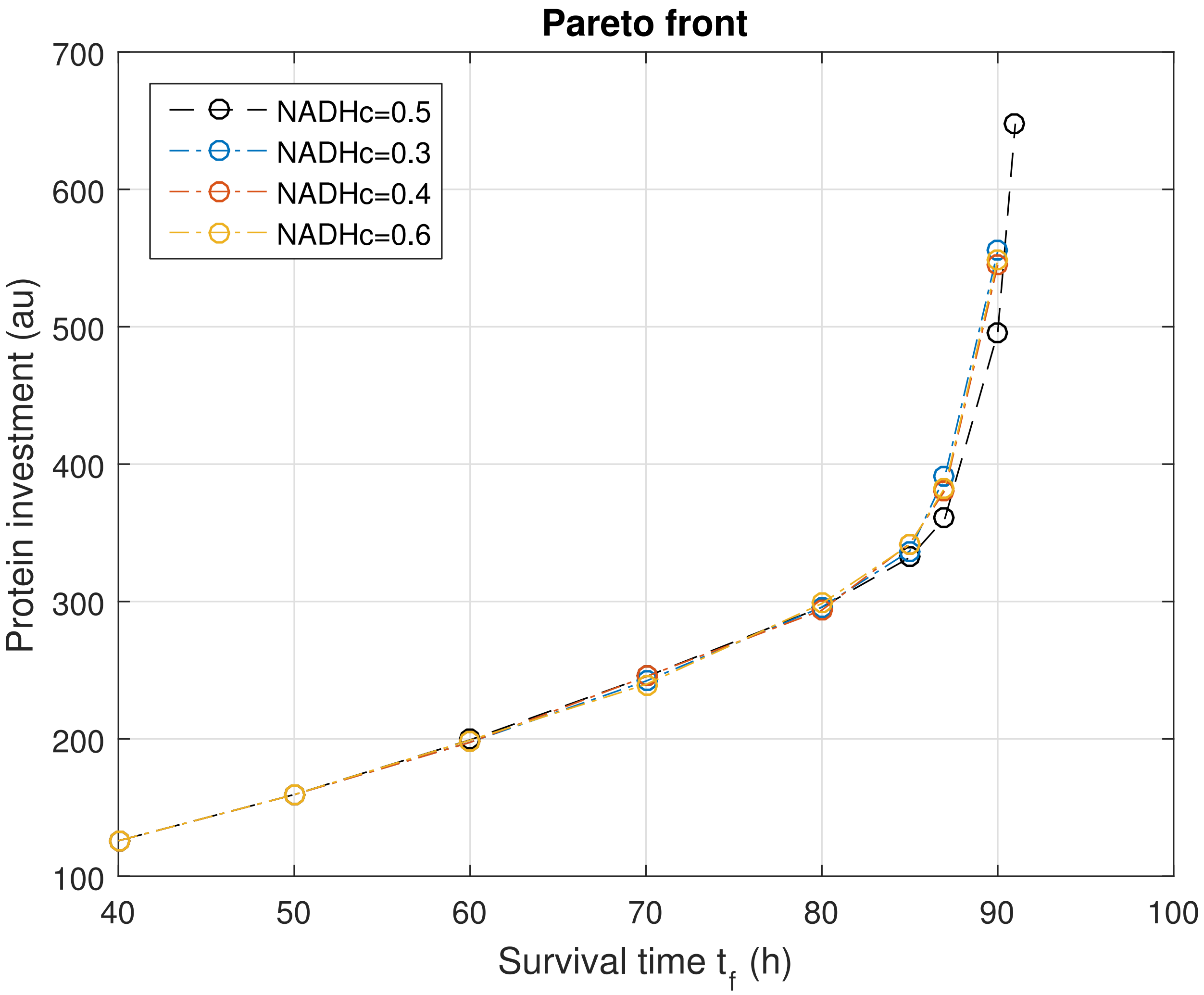


Figure S33: Comparison of the resulting Pareto front in SC case study for various NADH critical values.

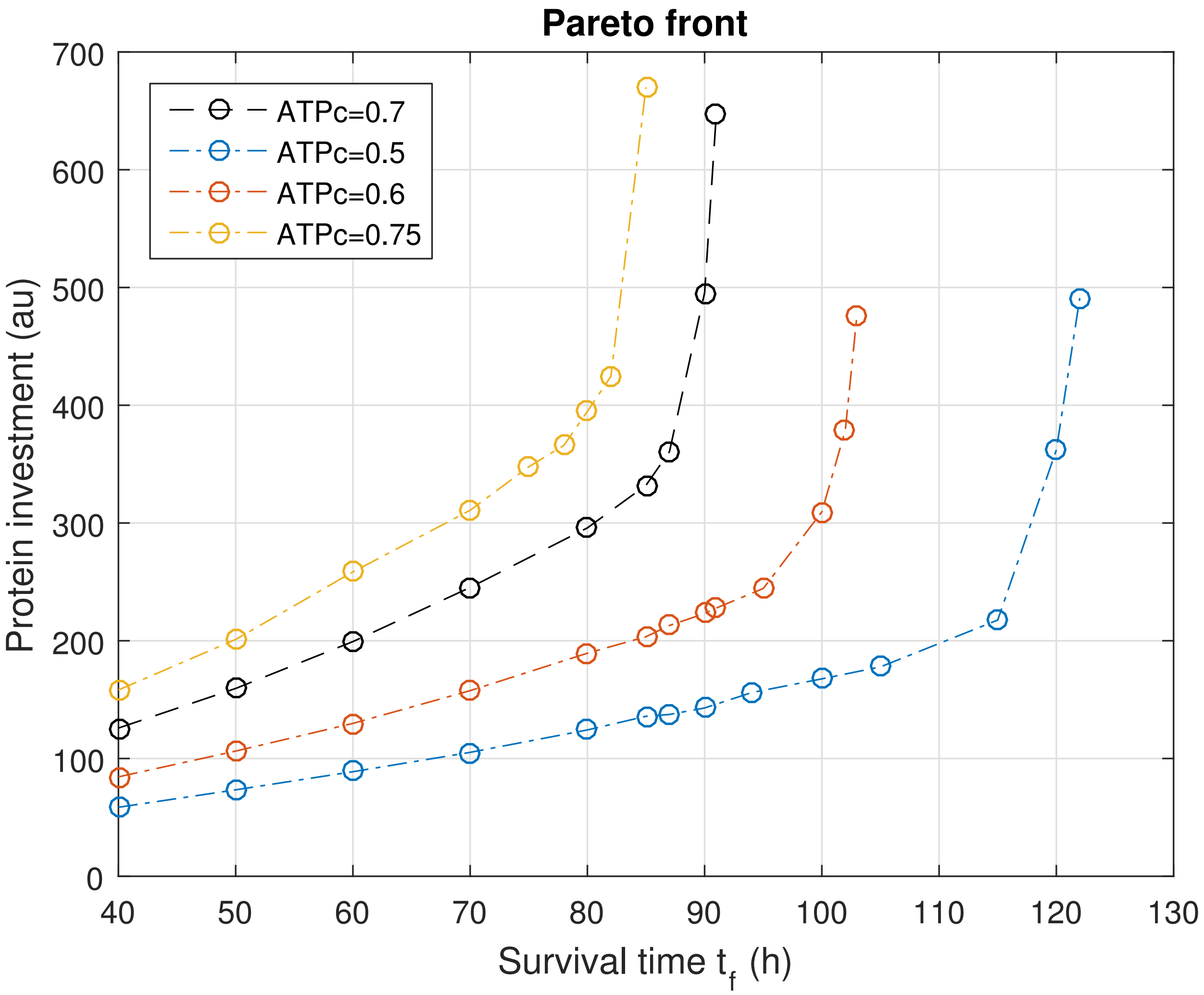


Figure S34: Comparison of the resulting Pareto front in SC case study for various ATP critical values.

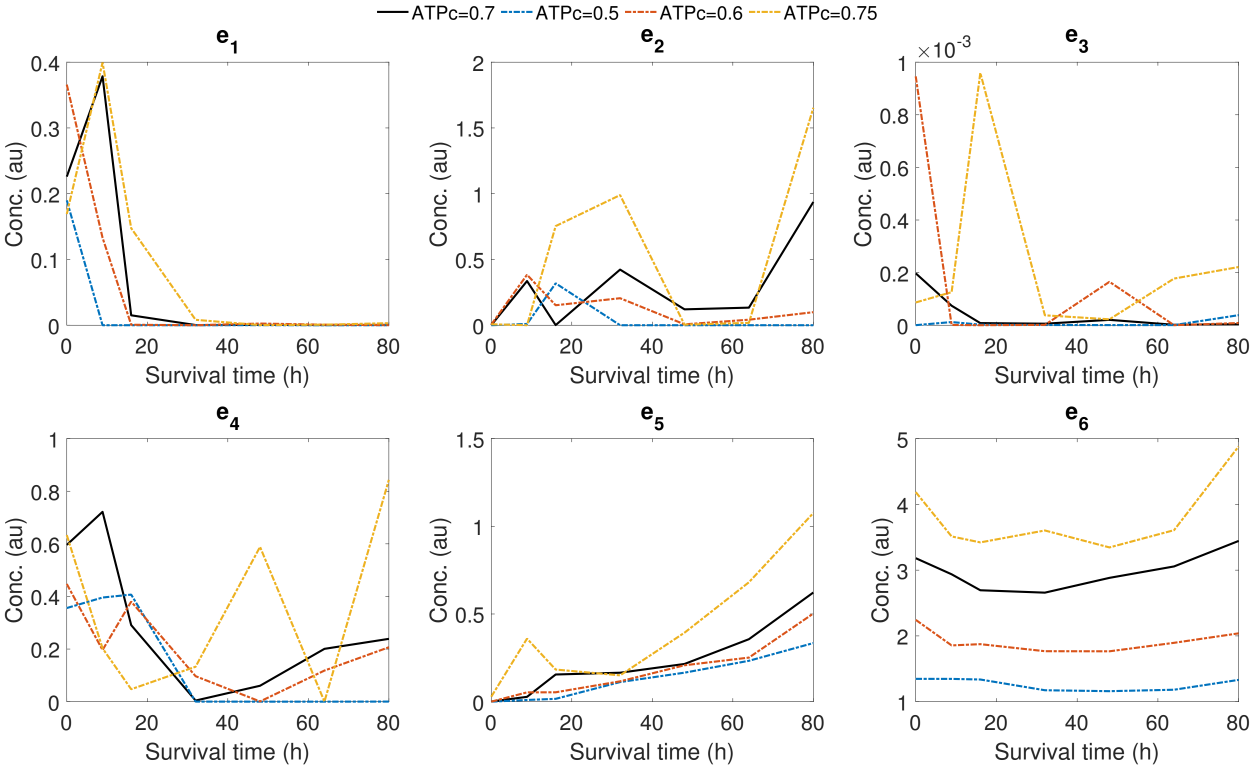


Figure S35: Comparison of the optimal controls for point $t_f = 80$, in SC case study, for various ATP critical values.

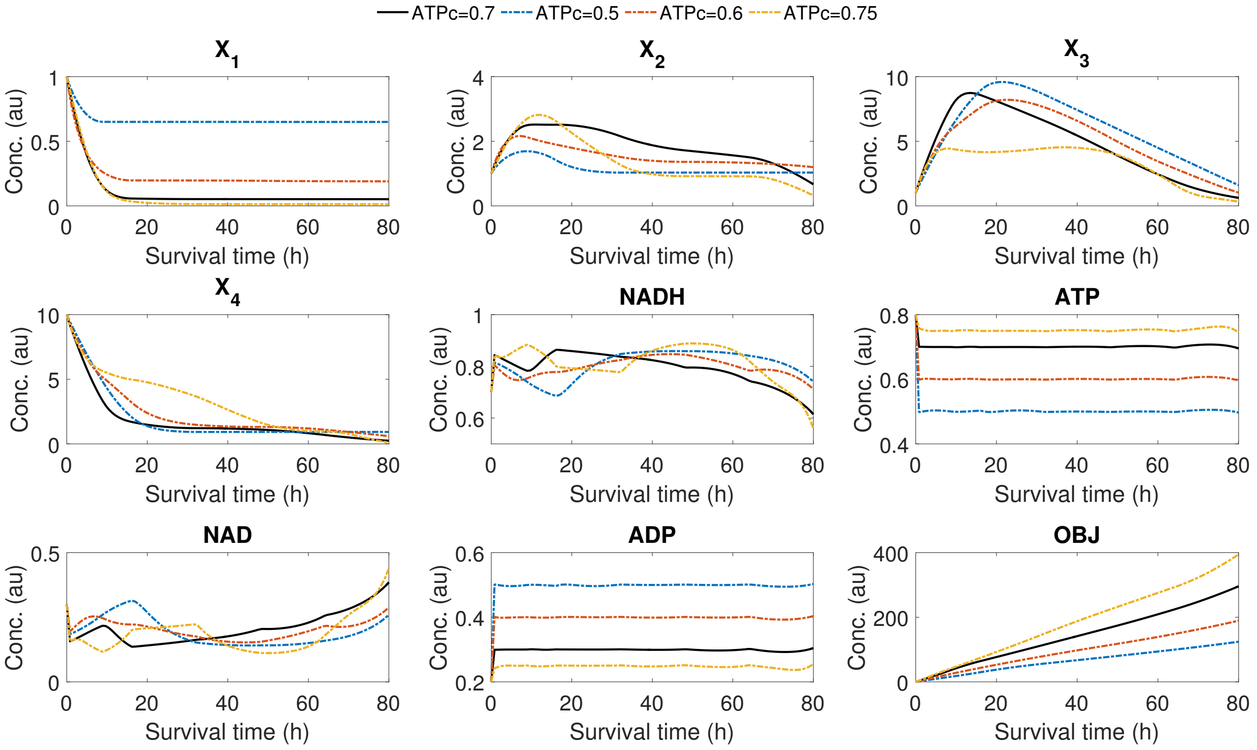


Figure S36: Comparison of the optimal state trajectories for point $t_f = 80$, in SC case study, for various ATP critical values.

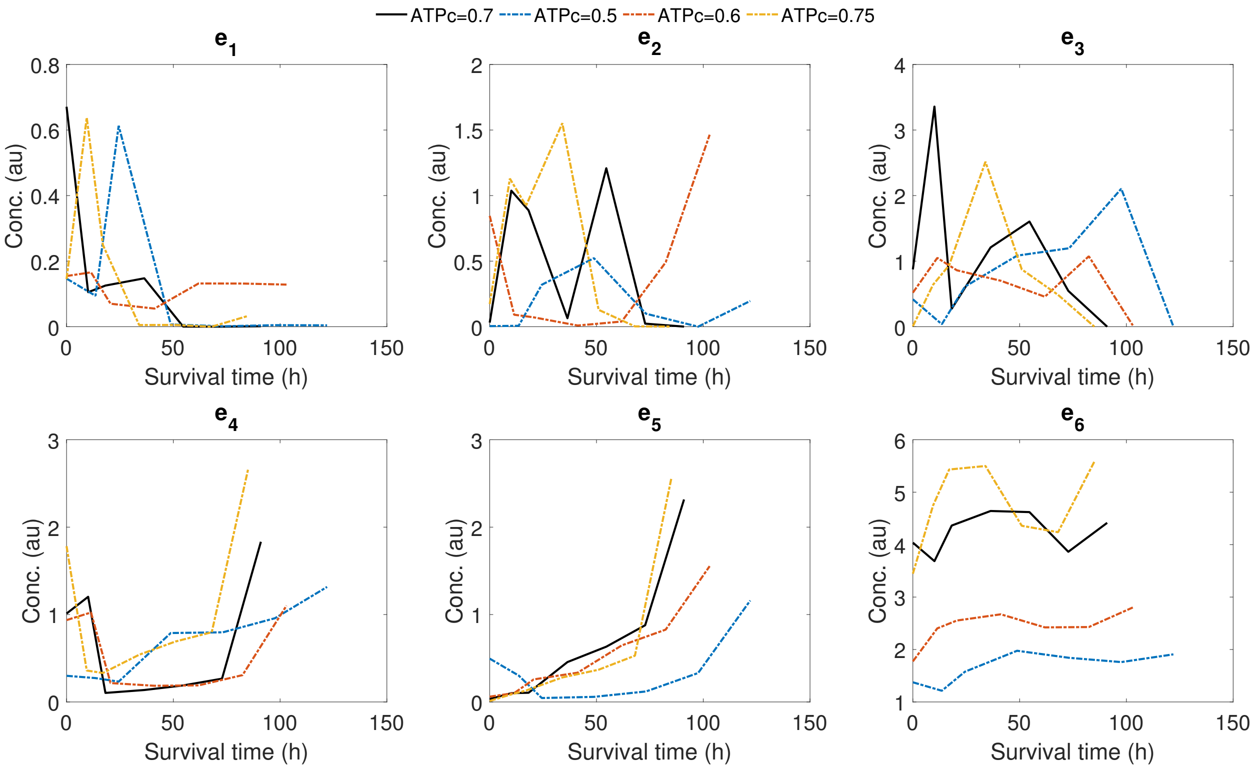


Figure S37: Comparison of the optimal controls for the extreme point with maximum survival time in the Pareto front, in SC case study, for various ATP critical values.

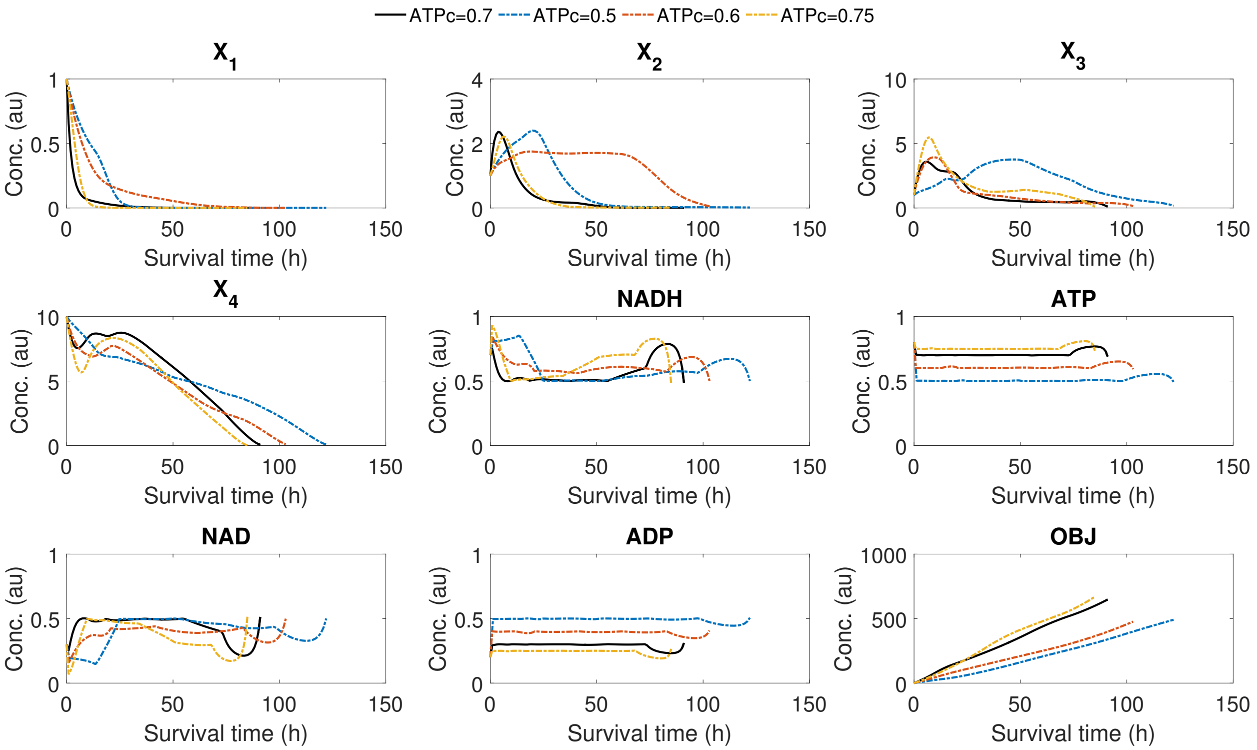


Figure S38: Comparison of the optimal state trajectories for the extreme point with maximum survival time in the Pareto front, in SC case study, for various ATP critical values.
