## Additional file 3 - case BSUB for "Using optimal control to understand complex metabolic pathways"

### Additional file 3: Solution details for case study BSUB

### Supplementary information

This case study considers two dynamic (two-phase) scenarios. The first phase of both scenarios (time 0-80 au) is the computation of a steady state while considering only one substrate present (either glucose or malate) in abundance. The second phase (time 80-160 au) is the introduction of the other substrate and the nutrient depletion scenario which is then computed. In this work, for simplicity we only present results for the second phase, with time scale 80-160 arbitrary units.

In Figure S1 we show that all three different approaches produced the same Pareto front when considering the first scenario (from glucose to malate). In Figures S2-S7 we present the optimal controls and state trajectories for several specific points of the Pareto shown. For the second scenario (from malate to glucose), results are presented in Figures S8-S14.

We also present a computational analysis for one of the points of the Pareto front. First we show how the set-up of the sequential hybrid `AMIGO2_DO+ICLOCS` affects the convergence in both scenarios in Figures S15,S16.

We also illustrate the performance of `AMIGO2_DO` in Figures S32-S34.

The performance of the multi-start strategy `msICLOCS` is shown in Figures S17-S19. Those results were obtained after fine-tuning the solver, providing by symbolic manipulation, analytical, while in Figures S20-S22 we present the results obtained with a numerical approximation, clearly showin worse convergence. Fine-tuning `msICLOCS` was essential to ensure good results and avoid numerical crashes.

We also present results obtained with `msICLOCS` for different points in the Pareto front as we observed significant differences in the CPU requirements amongst them (Figures S23-S31).

Finally, we performed an approximate analysis of sensitivity of the controls in order to evaluate the possible non-uniqueness of the solutions. This was done by performig 100 runs of `AMIGO2_DO+ICLOCS`. Results are given In Figures S35-S42.

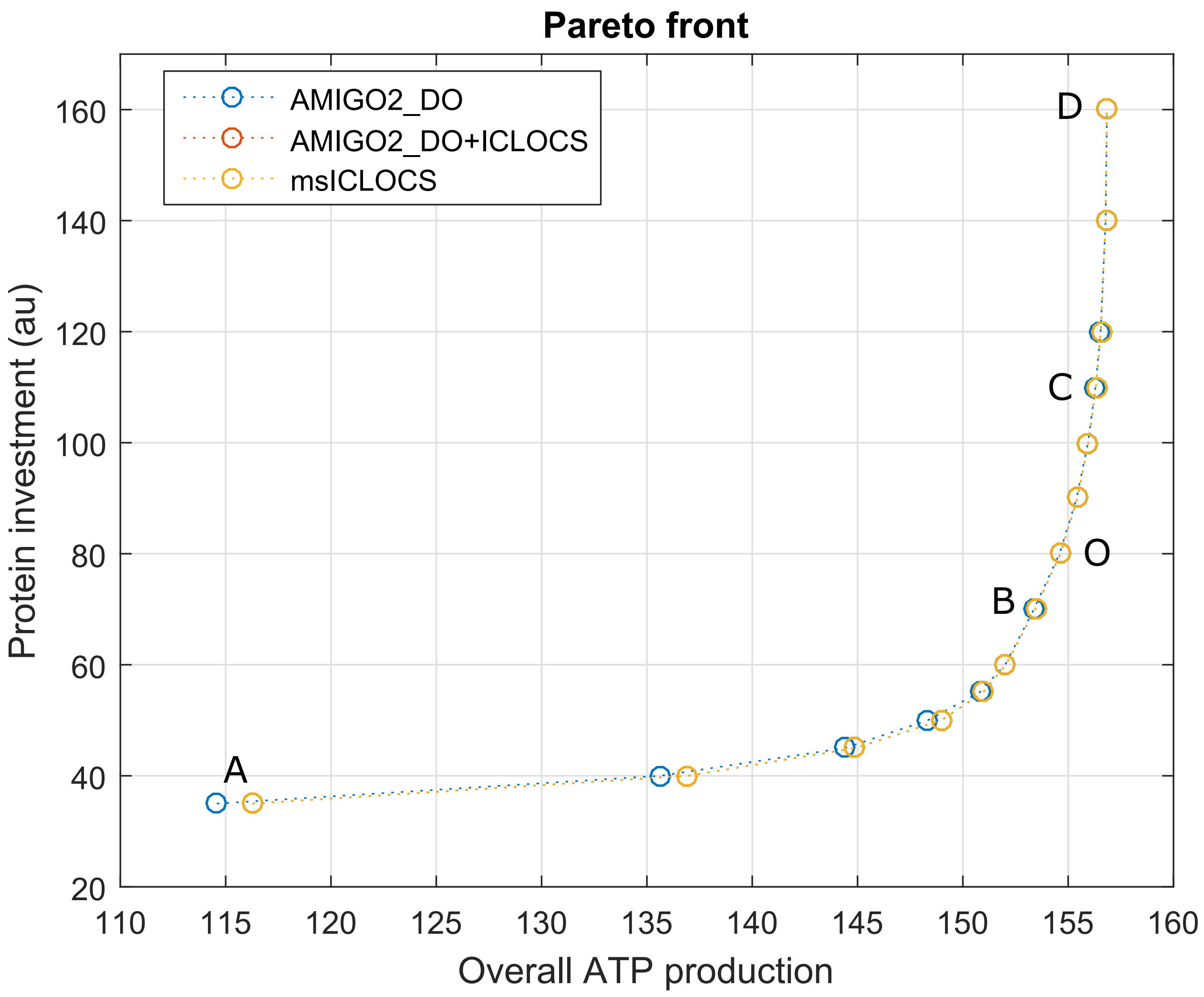


Figure S1: Pareto front comparison between `AMIGO2_DO` with 10PWC, `AMIGO2_DO+ICLOCS` with 10PWC and `msICLOCS` with 10PWC for the G-M scenario.

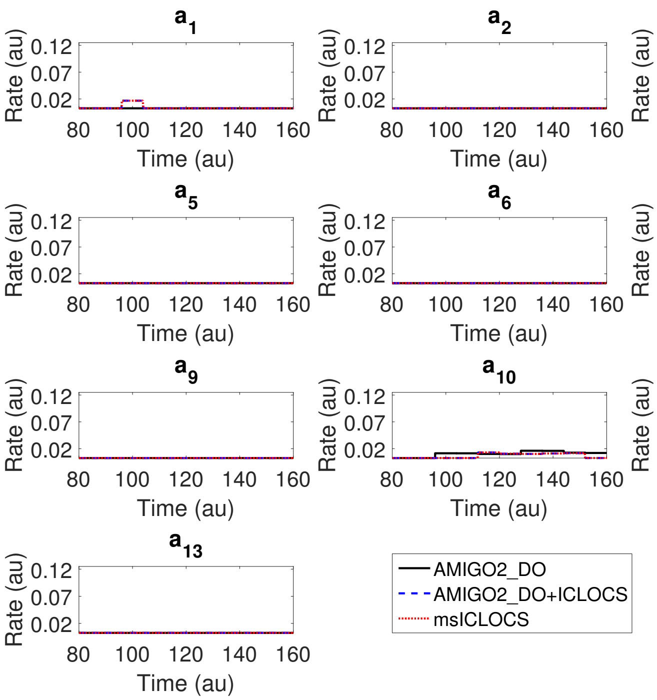


Figure S2: The optimal controls computed for point A of the Pareto front corresponding to the G-M scenario, computed with `AMIGO2_DO` with 10PWC (black solid lines) versus the ones computed with `AMIGO2_DO+ICLOCS` 10PWC (blue dash lines) and with `msICLOCS` 10PWC (red dotted line).

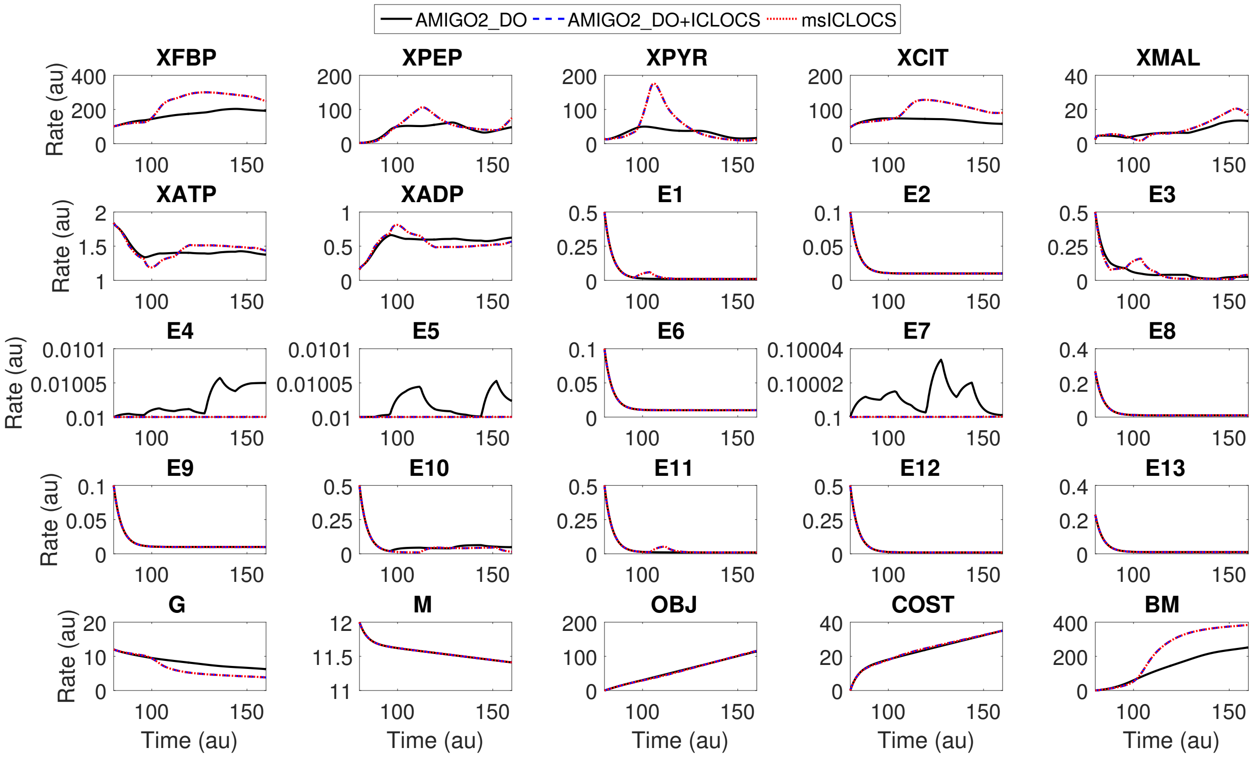


Figure S3: The optimal state trajectories computed for point A of the Pareto front corresponding to the G-M scenario, computed with `AMIGO2_DO` with 10PWC (black solid lines) versus the ones computed with `AMIGO2_DO+ICLOCS` 10PWC (blue dash lines) and with `msICLOCS` 10PWC (red dotted line).

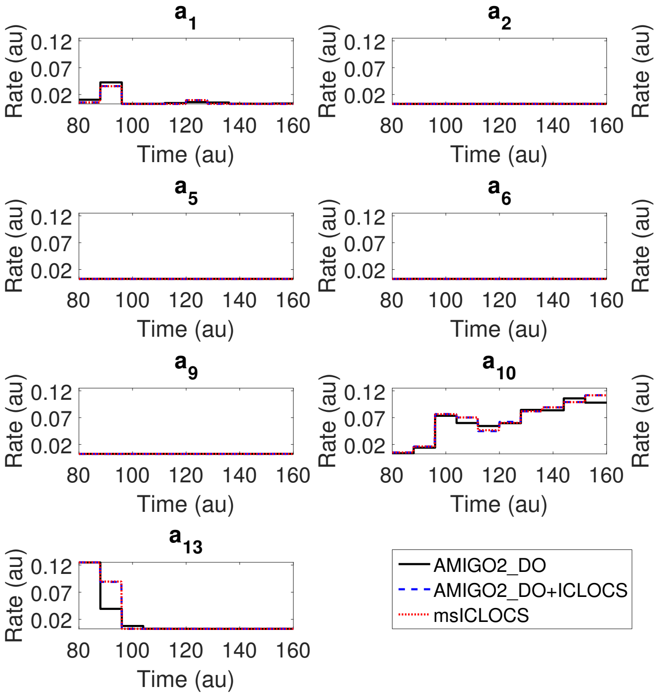


Figure S4: The optimal controls computed for point B of the Pareto front corresponding to the G-M scenario, computed with `AMIGO2_DO` with 10PWC (black solid lines) versus the ones computed with `AMIGO2_DO+ICLOCS` 10PWC (blue dash lines) and with `msICLOCS` 10PWC (red dotted line).

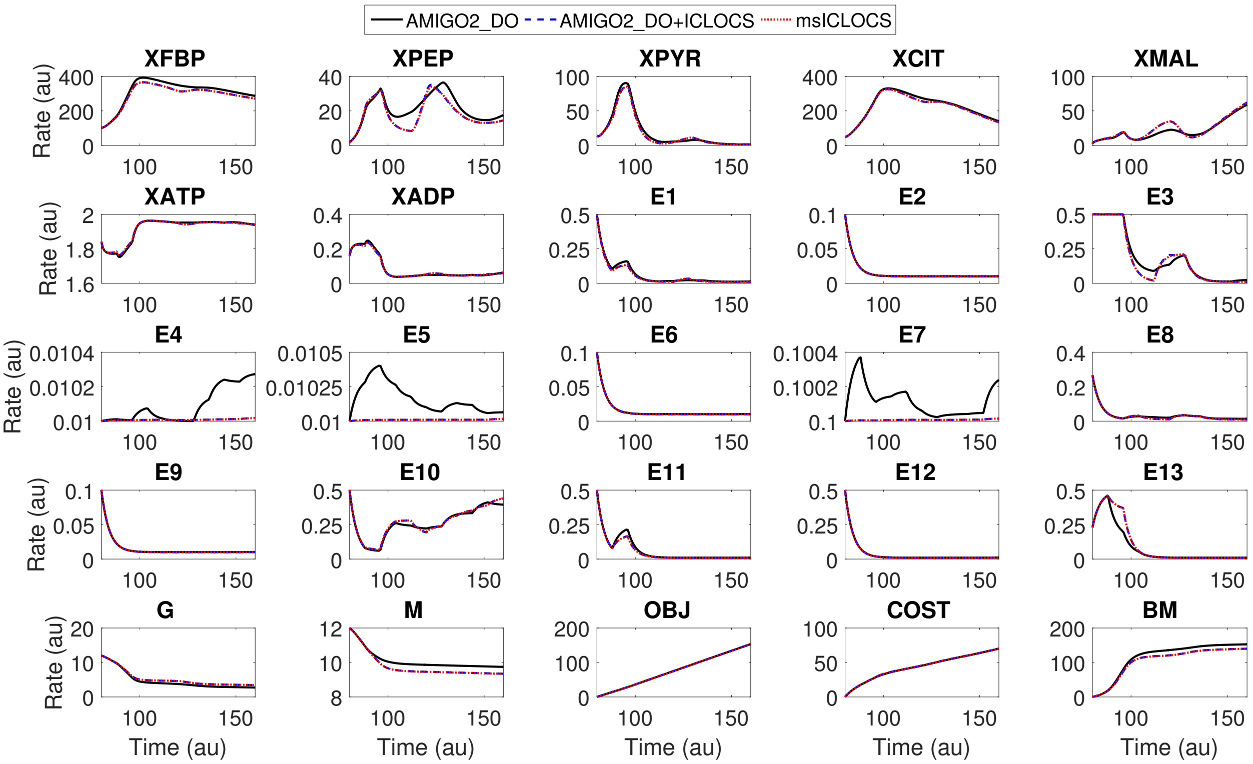


Figure S5: The optimal state trajectories computed for point B of the Pareto frontt corresponding to the G-M scenario, computed with `AMIGO2_DO` with 10PWC (black solid lines) versus the ones computed with `AMIGO2_DO+ICLOCS` 10PWC (blue dash lines) and with `msICLOCS` 10PWC (red dotted line).

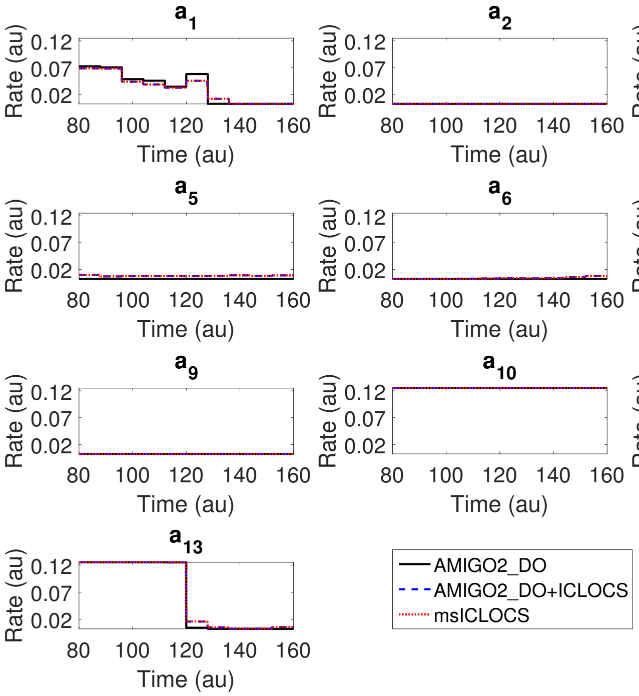


Figure S6: The optimal controls computed for point D of the Pareto front corresponding to the G-M scenario, computed with `AMIGO2_DO` with 10PWC (black solid lines) versus the ones computed with `AMIGO2_DO+ICLOCS` 10PWC (blue dash lines) and with `msICLOCS` 10PWC (red dotted line).

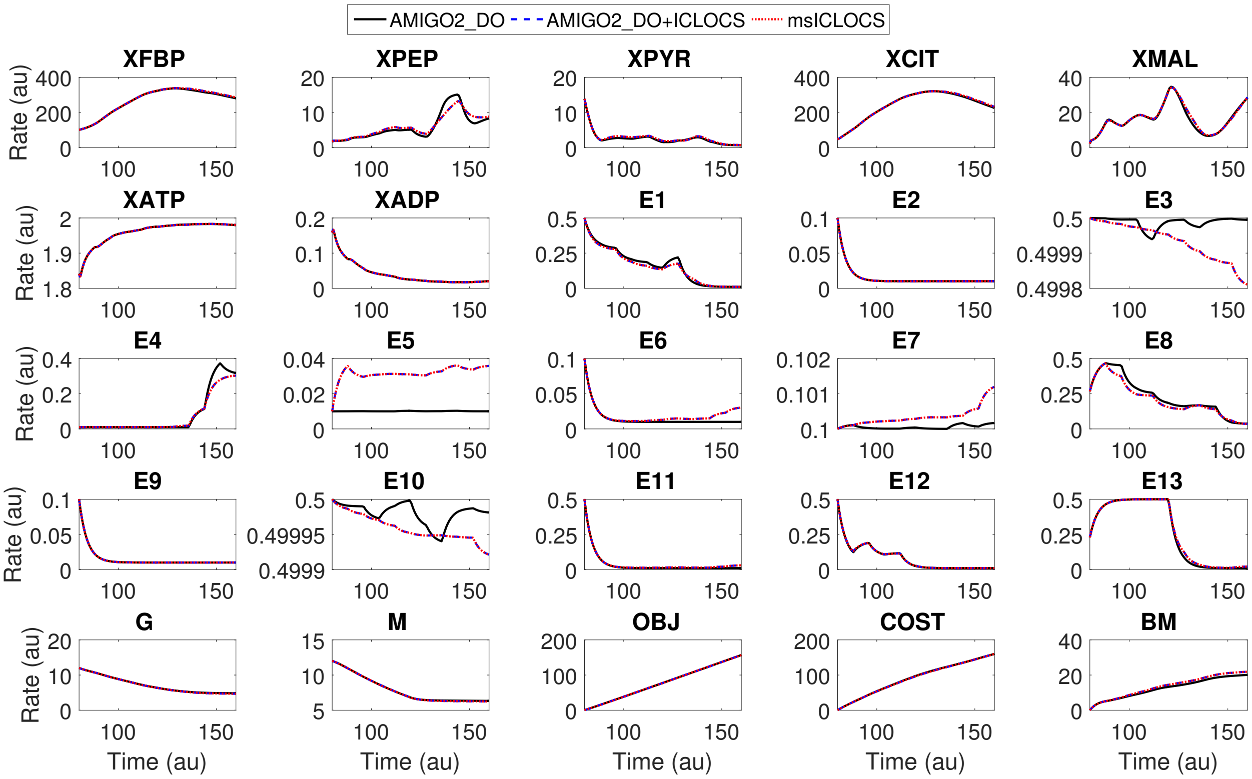


Figure S7: The optimal state trajectories computed for point D of the Pareto front corresponding to the G-M scenario, computed with `AMIGO2_DO` with 10PWC (black solid lines) versus the ones computed with `AMIGO2_DO+ICLOCS` 10PWC (blue dash lines) and with `msICLOCS` 10PWC (red dotted line).

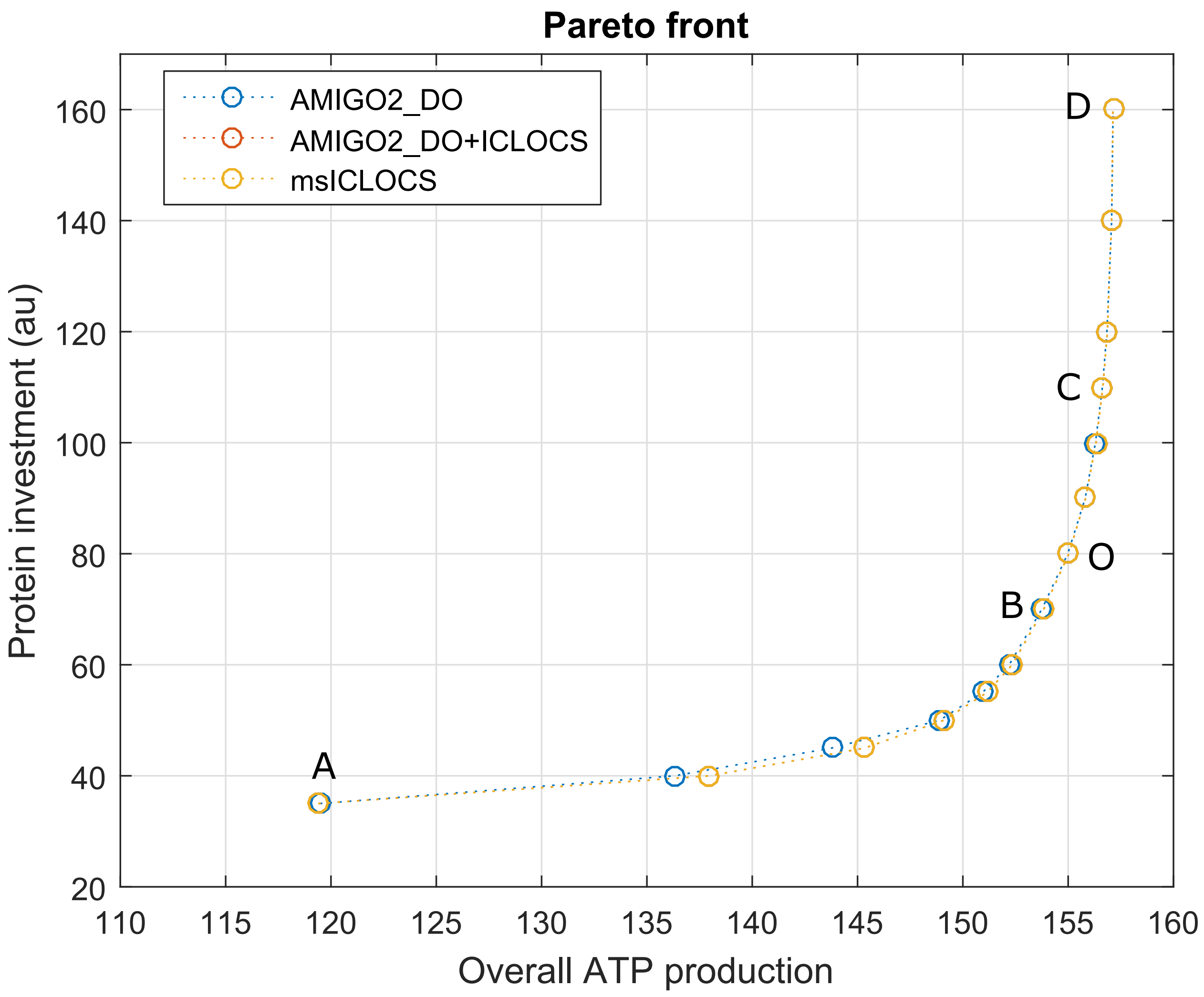


Figure S8: Pareto front comparison between `AMIGO2_DO` with 10PWC, `AMIGO2_DO+ICLOCS` with 10PWC and `msICLOCS` with 10PWC for the M-G scenario.

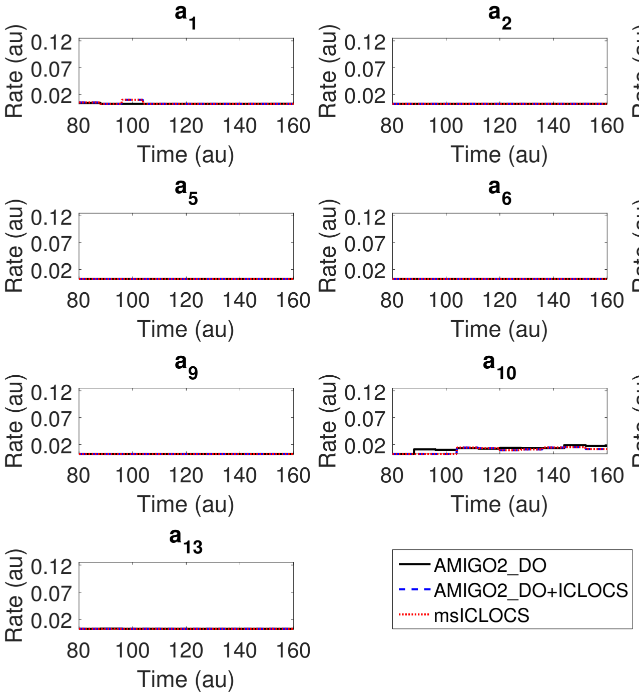


Figure S9: The optimal controls computed for point A of the Pareto front corresponding to the M-G scenario, computed with `AMIGO2_DO` with 10PWC (black solid lines) versus the ones computed with `AMIGO2_DO+ICLOCS` 10PWC (blue dash lines) and with `msICLOCS` 10PWC (red dotted line).

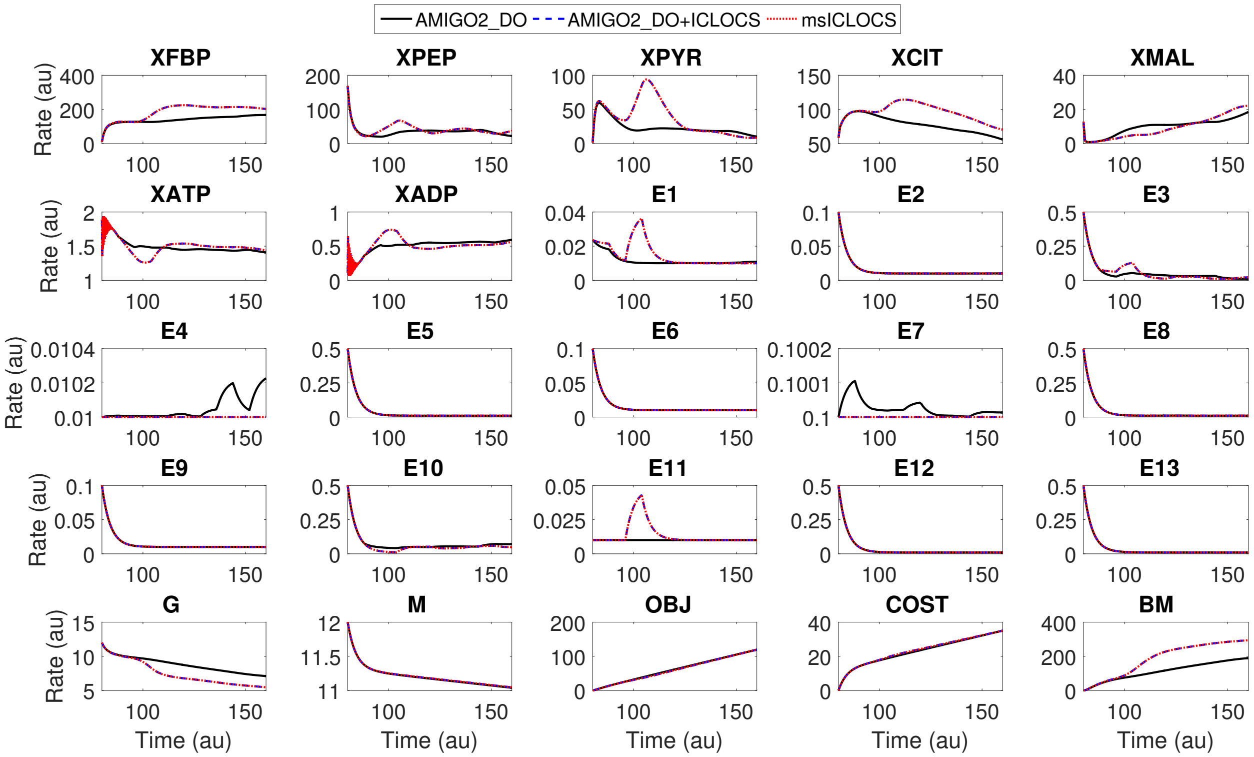


Figure S10: The optimal state trajectories computed for point A of the Pareto front corresponding to the M-G scenario, computed with `AMIGO2_DO` with 10PWC (black solid lines) versus the ones computed with `AMIGO2_DO+ICLOCS` 10PWC (blue dash lines) and with `msICLOCS` 10PWC (red dotted line).

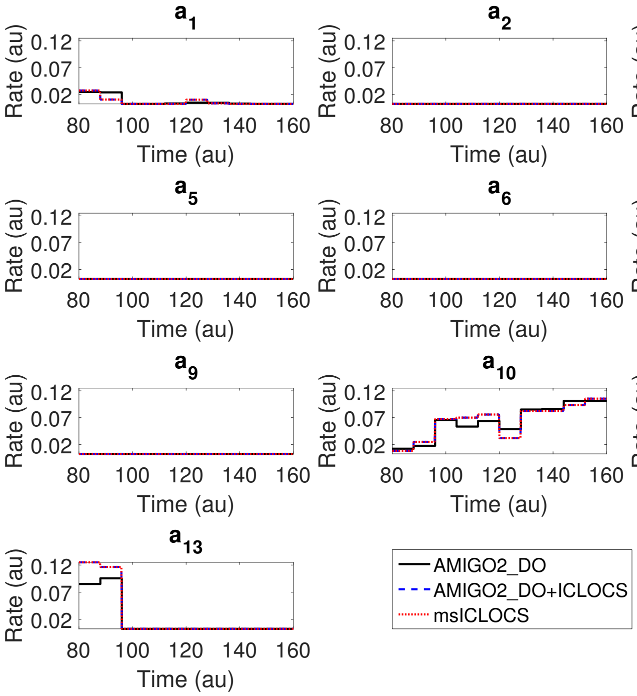


Figure S11: The optimal controls computed for point B of the Pareto front corresponding to the M-G scenario, computed with `AMIGO2_DO` with 10PWC (black solid lines) versus the ones computed with `AMIGO2_DO+ICLOCS` 10PWC (blue dash lines) and with `msICLOCS` 10PWC (red dotted line).

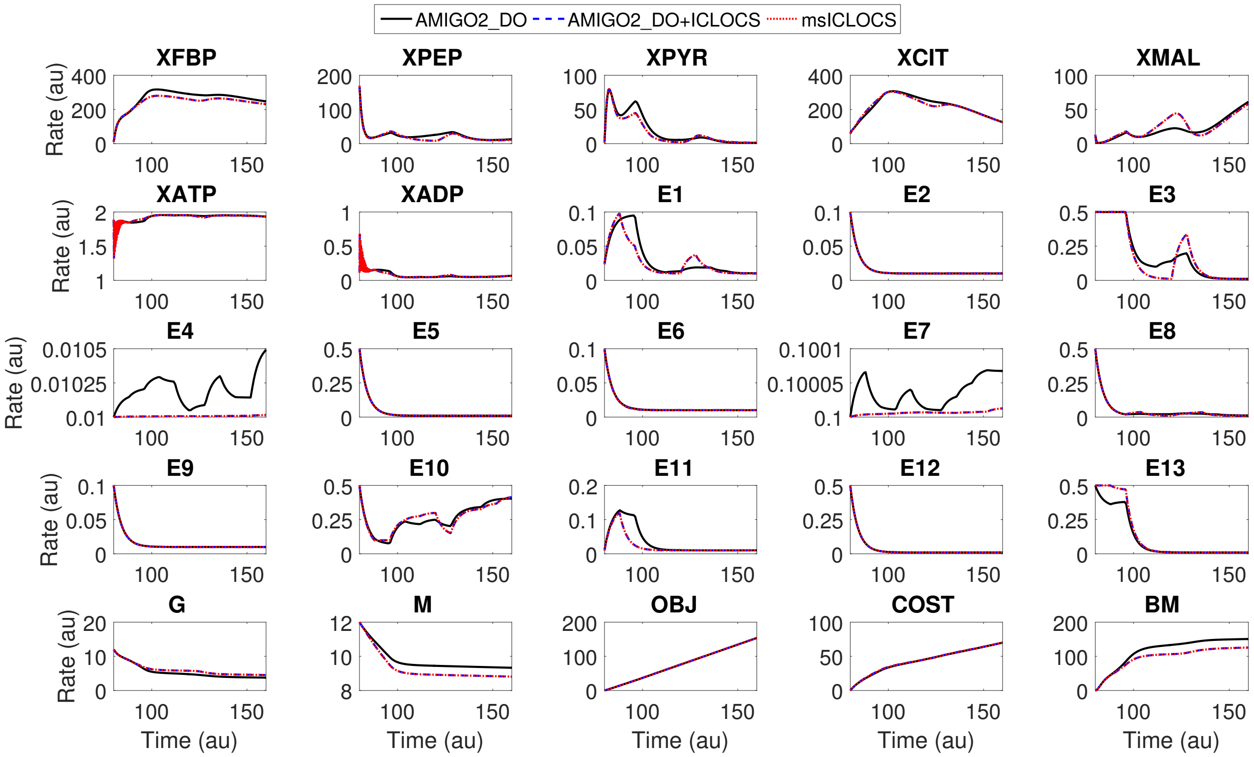


Figure S12: The optimal state trajectories computed for point B of the Pareto front corresponding to the M-G scenario, computed with `AMIGO2_DO` with 10PWC (black solid lines) versus the ones computed with `AMIGO2_DO+ICLOCS` 10PWC (blue dash lines) and with `msICLOCS` 10PWC (red dotted line).

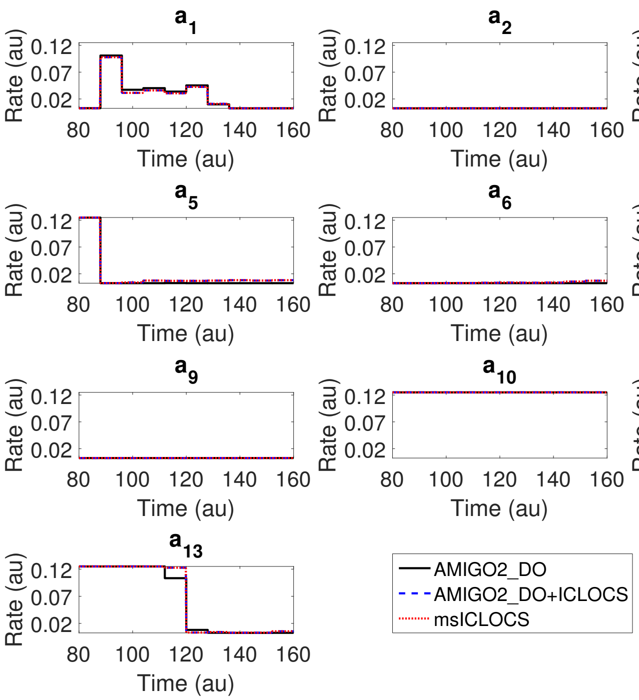


Figure S13: The optimal controls computed for point D of the Pareto front corresponding to the M-G scenario, computed with `AMIGO2_DO` with 10PWC (black solid lines) versus the ones computed with `AMIGO2_DO+ICLOCS` 10PWC (blue dash lines) and with `msICLOCS` 10PWC (red dotted line).

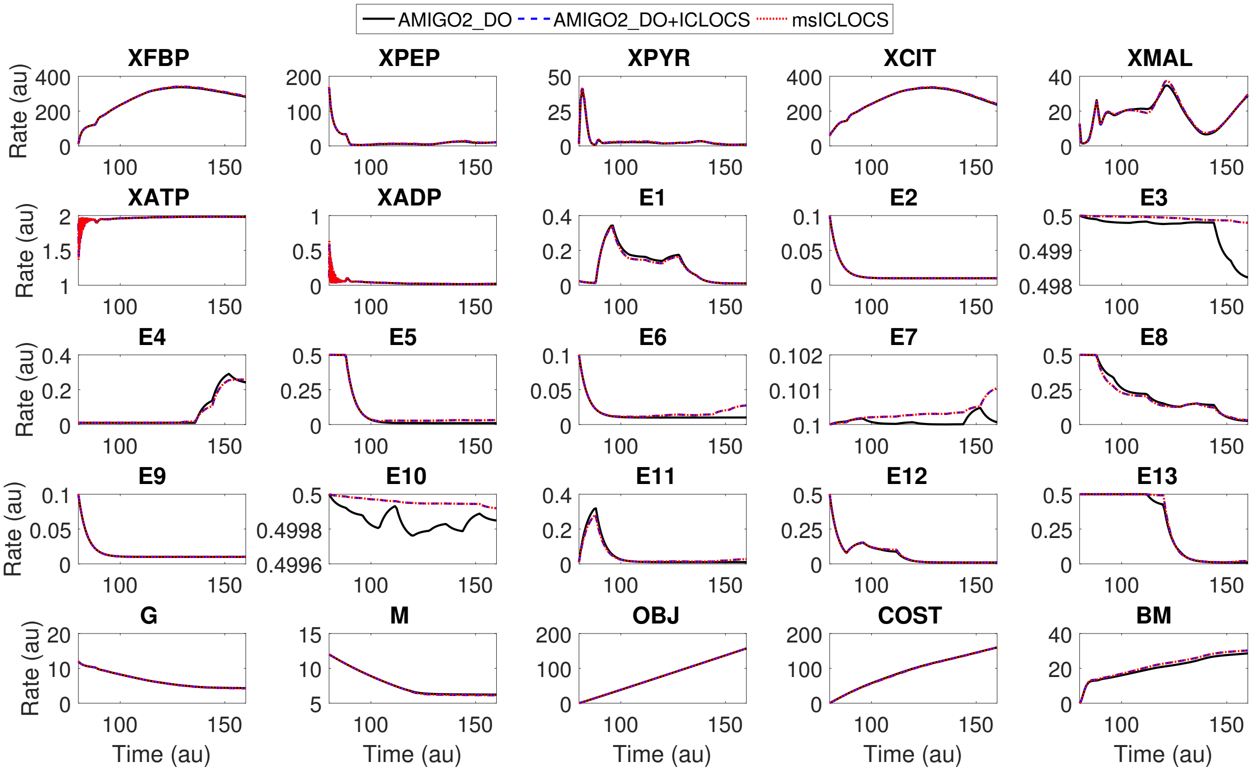


Figure S14: The optimal state trajectories computed for point D of the Pareto front corresponding to the M-G scenario, computed with AMIGO2_DO with 10PWC (black solid lines) versus the ones computed with AMIGO2_DO+ICLOCS 10PWC (blue dash lines) and with msICLOCS 10PWC (red dotted line).

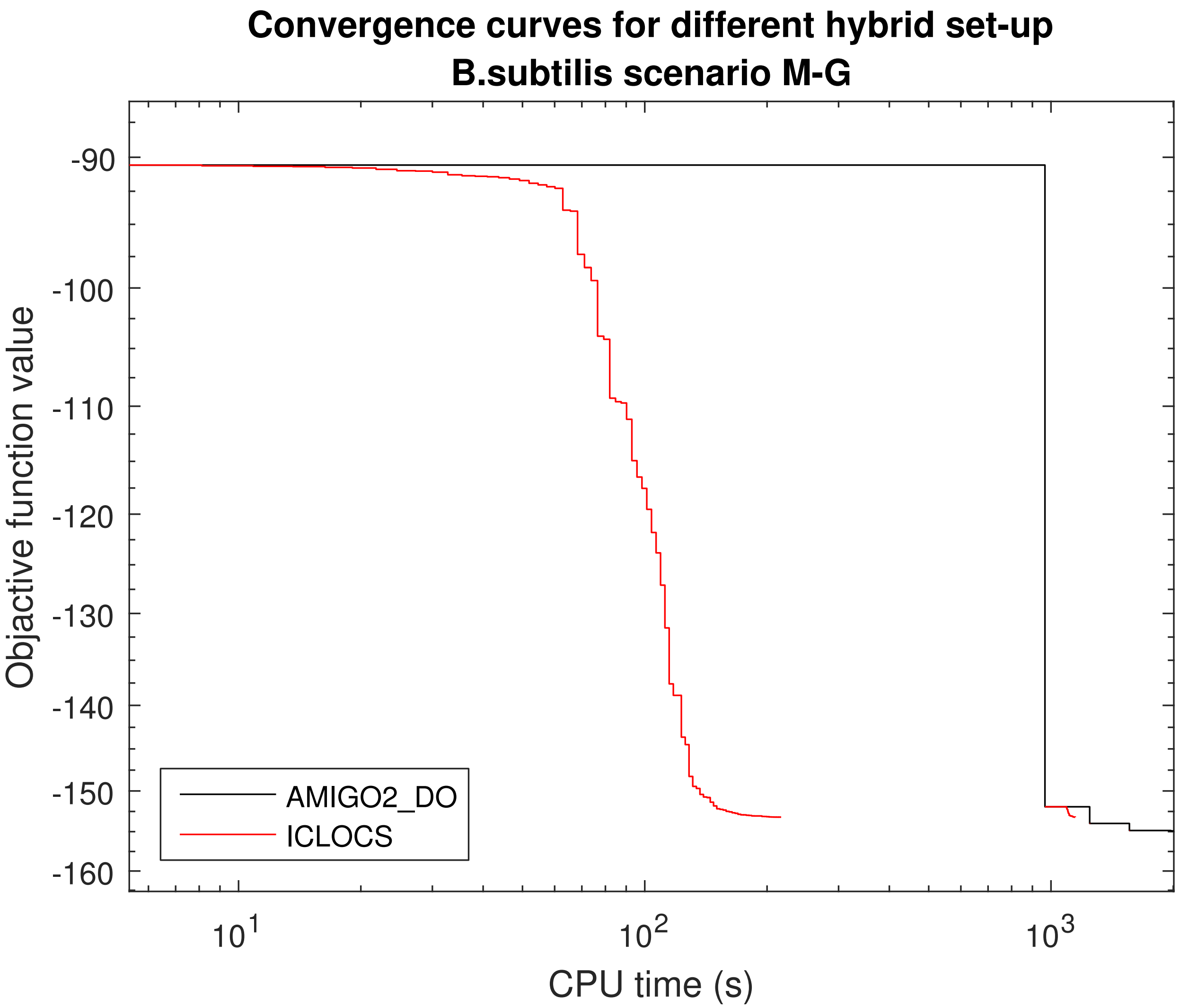


Figure S15: BSUB: Convergence curves of `AMIGO2_DO+ICLOCS` for point O of the Pareto front and the M-G scenario. `ICLOCS` (10PWC) is initialized from different point of `AMIGO2_DO` (5PWC) convergence.

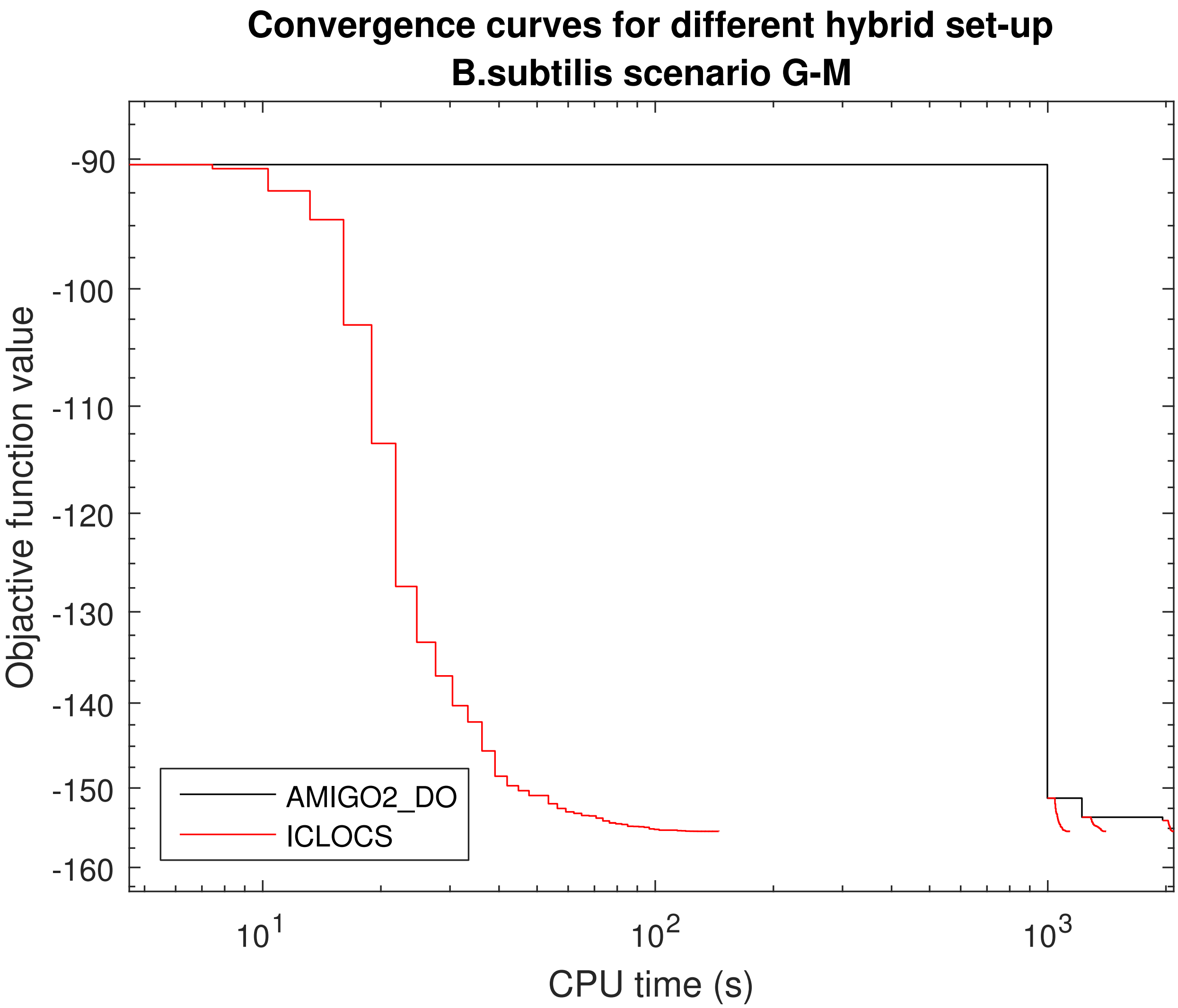


Figure S16: BSUB: Convergence curves of `AMIGO2_DO+ICLOCS` for point O of the Pareto front and the G-M scenario. `ICLOCS` (10PWC) is initialized from different point of `AMIGO2_DO` (5PWC) convergence.

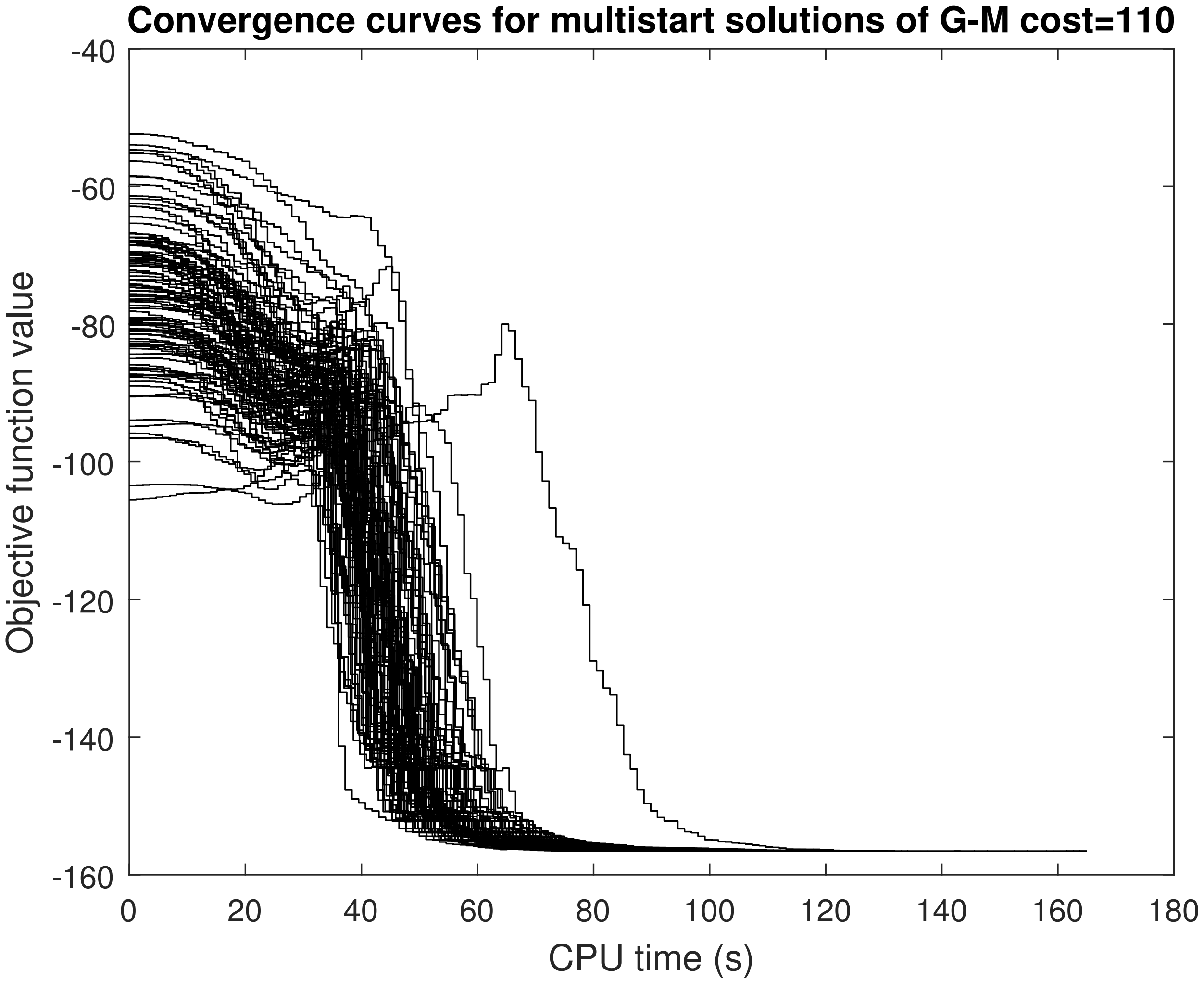


Figure S17: BSUB: Convergence curves of the multistart OCP solutions of a total of 100 `ICLOCS` runs initialized randomly in the search space for scenario G-M and for point C of the Pareto front.

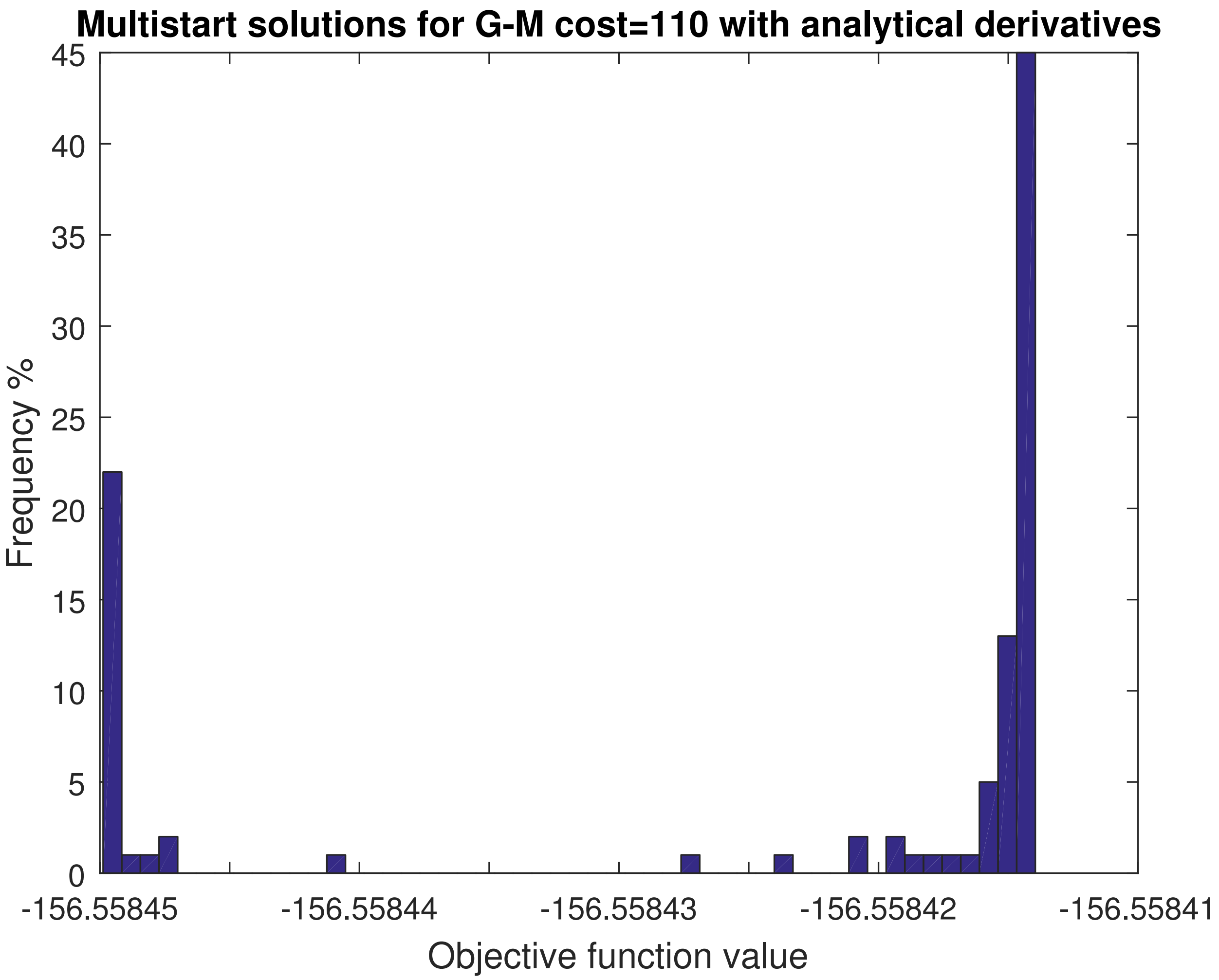


Figure S18: BSUB: Histogram of the multistart OCP solutions of a total of 100 `ICLOCS` runs initialized randomly in the search space for scenario G-M and for point C of the Pareto front.

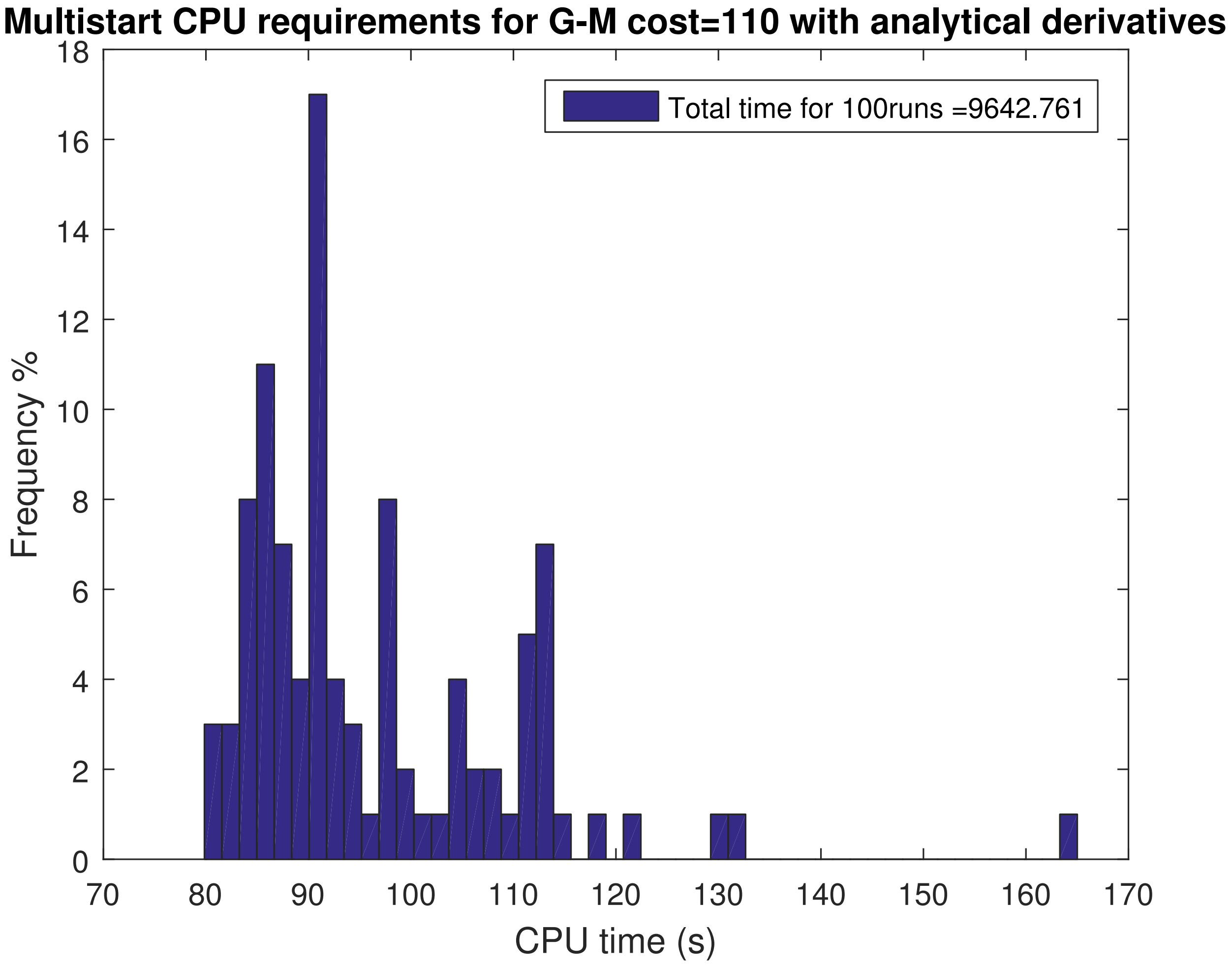


Figure S19: BSUB: Histogram of the computational time required obtaining the multistart OCP solutions of a total of 100 ICLOCS runs initialized randomly in the search space for scenario G-M and for point C of the Pareto front.

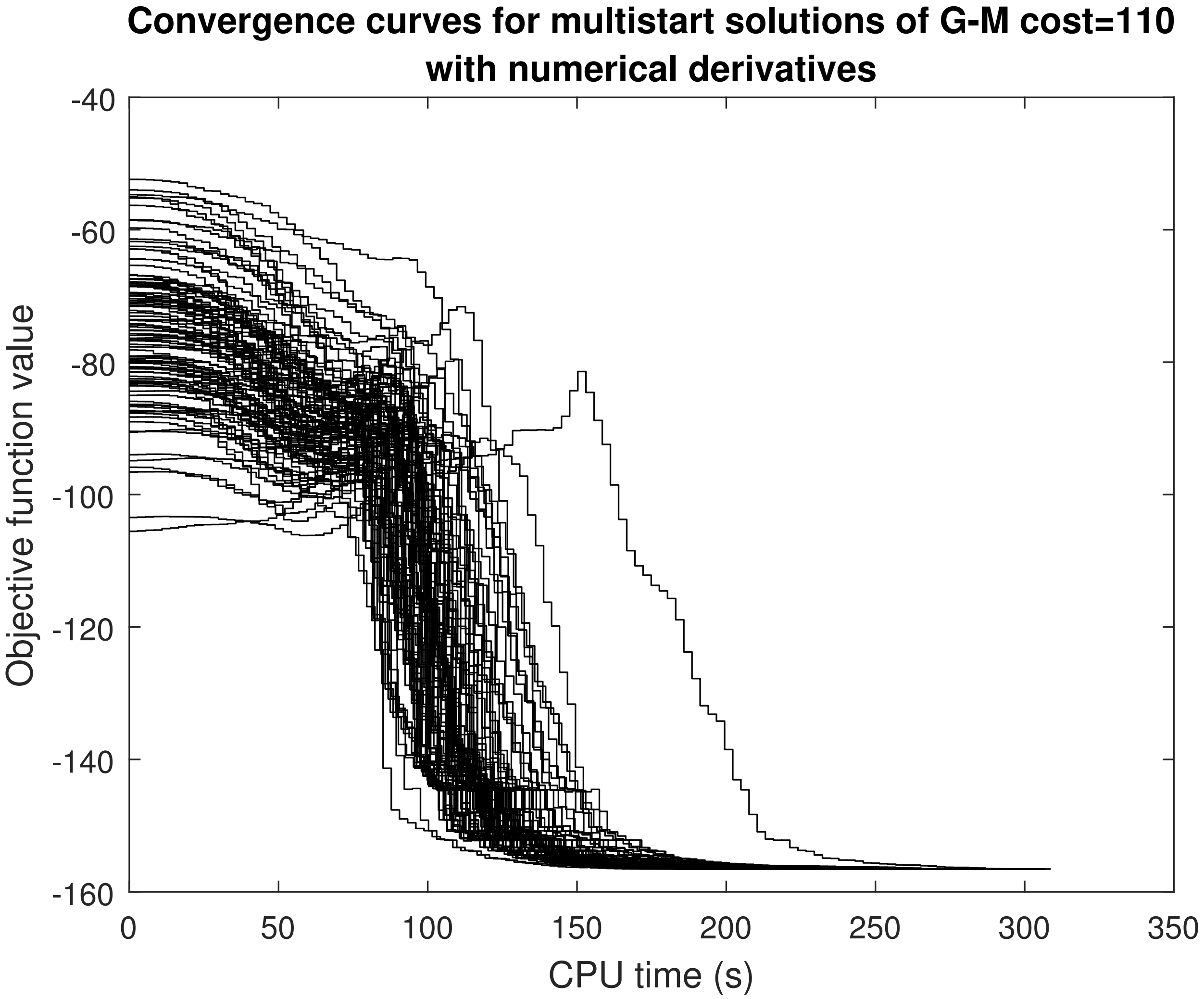


Figure S20: BSUB: Convergence curves of the multistart OCP solutions of a total of 100 `ICLOCS` runs initialized randomly in the search space for scenario G-M and for point C of the Pareto front. Here the system's derivatives were computed numerically.

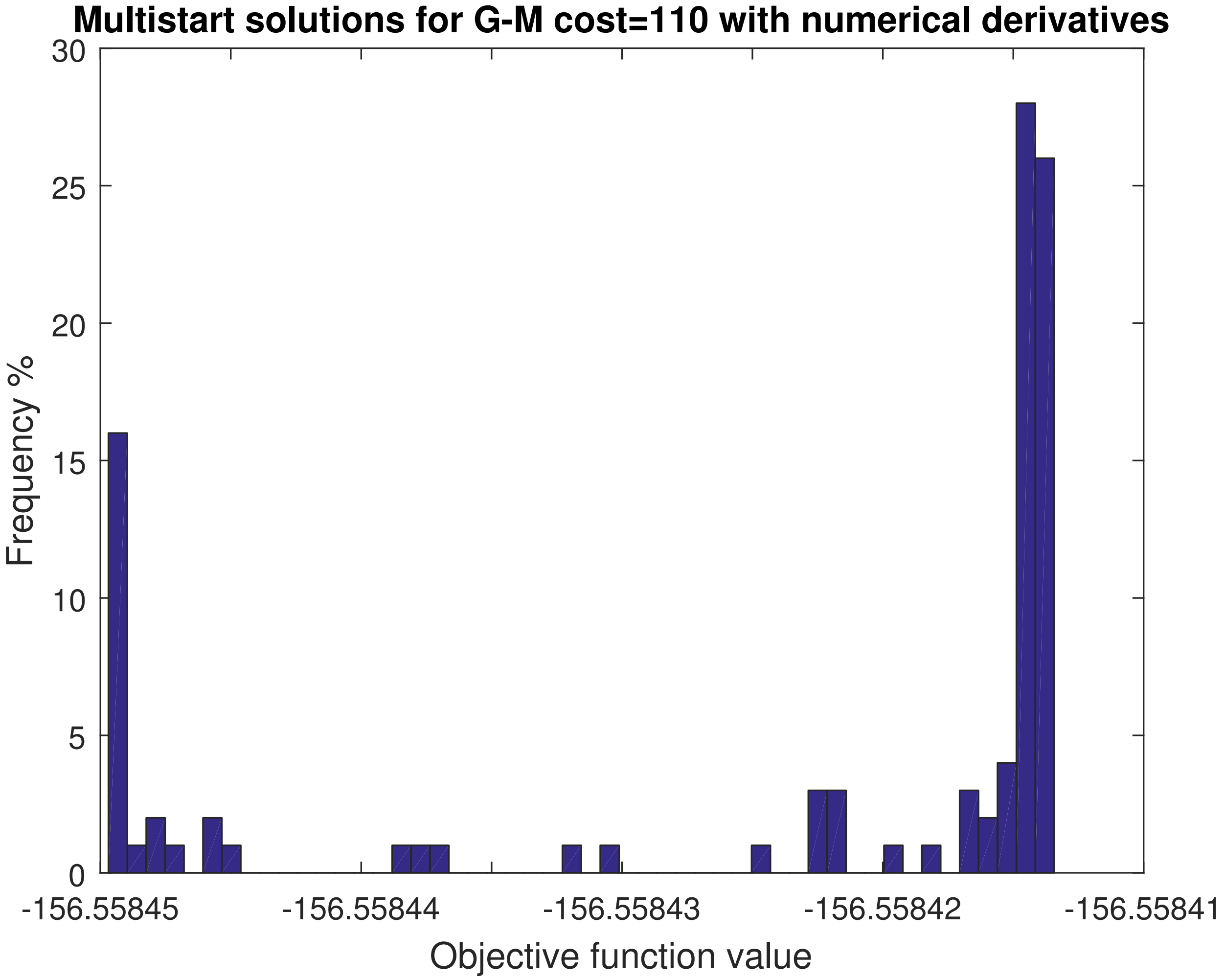


Figure S21: BSUB: Histogram of the multistart OCP solutions of a total of 100 `ICLOCS` runs initialized randomly in the search space for scenario G-M and for point C of the Pareto front. Here the system's derivatives were computed numerically.

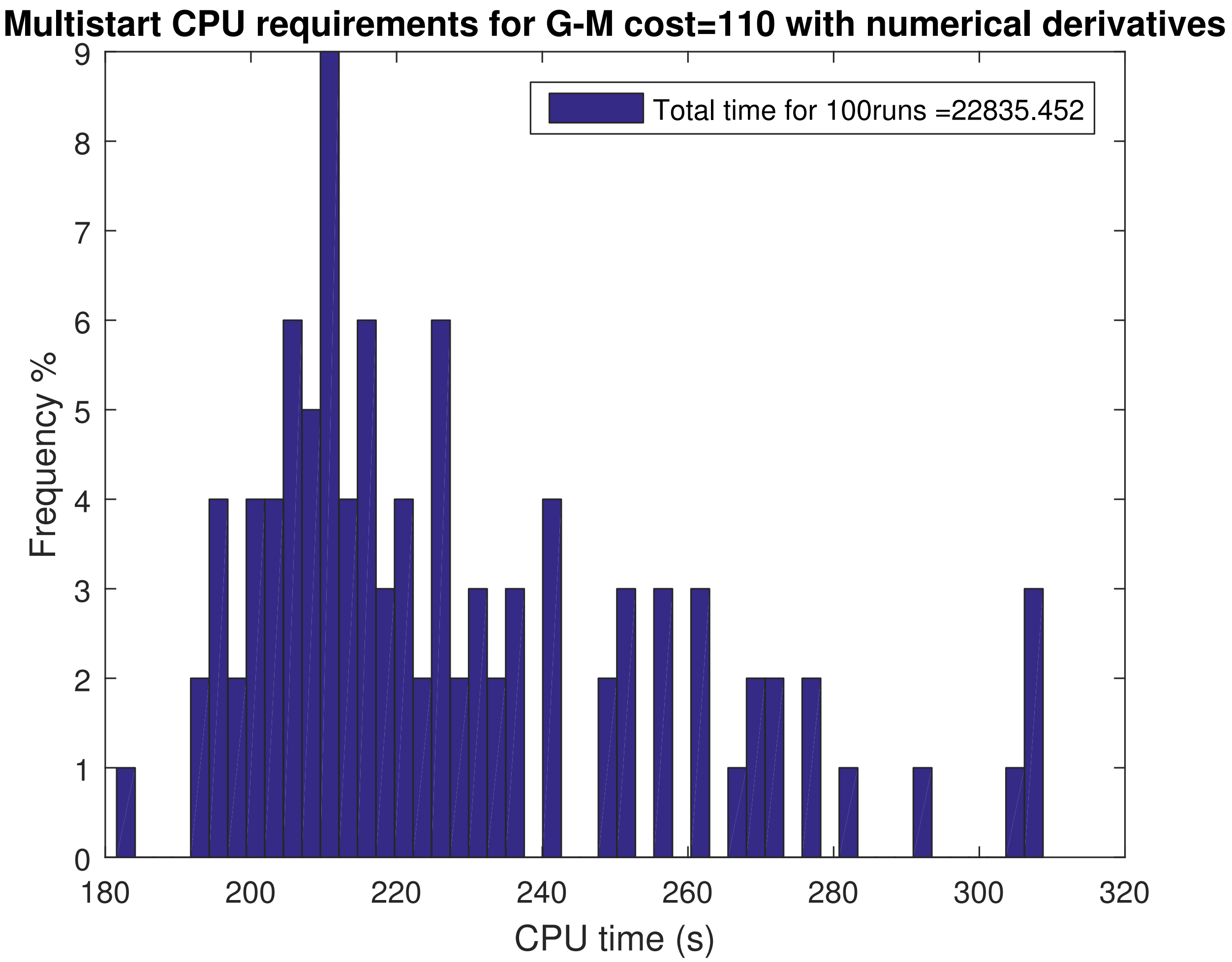


Figure S22: BSUB: Histogram of the computational time required obtaining the multistart OCP solutions of a total of 100 ICLOCS runs initialized randomly in the search space for scenario G-M and for point C of the Pareto front. Here the system's derivatives were computed numerically.

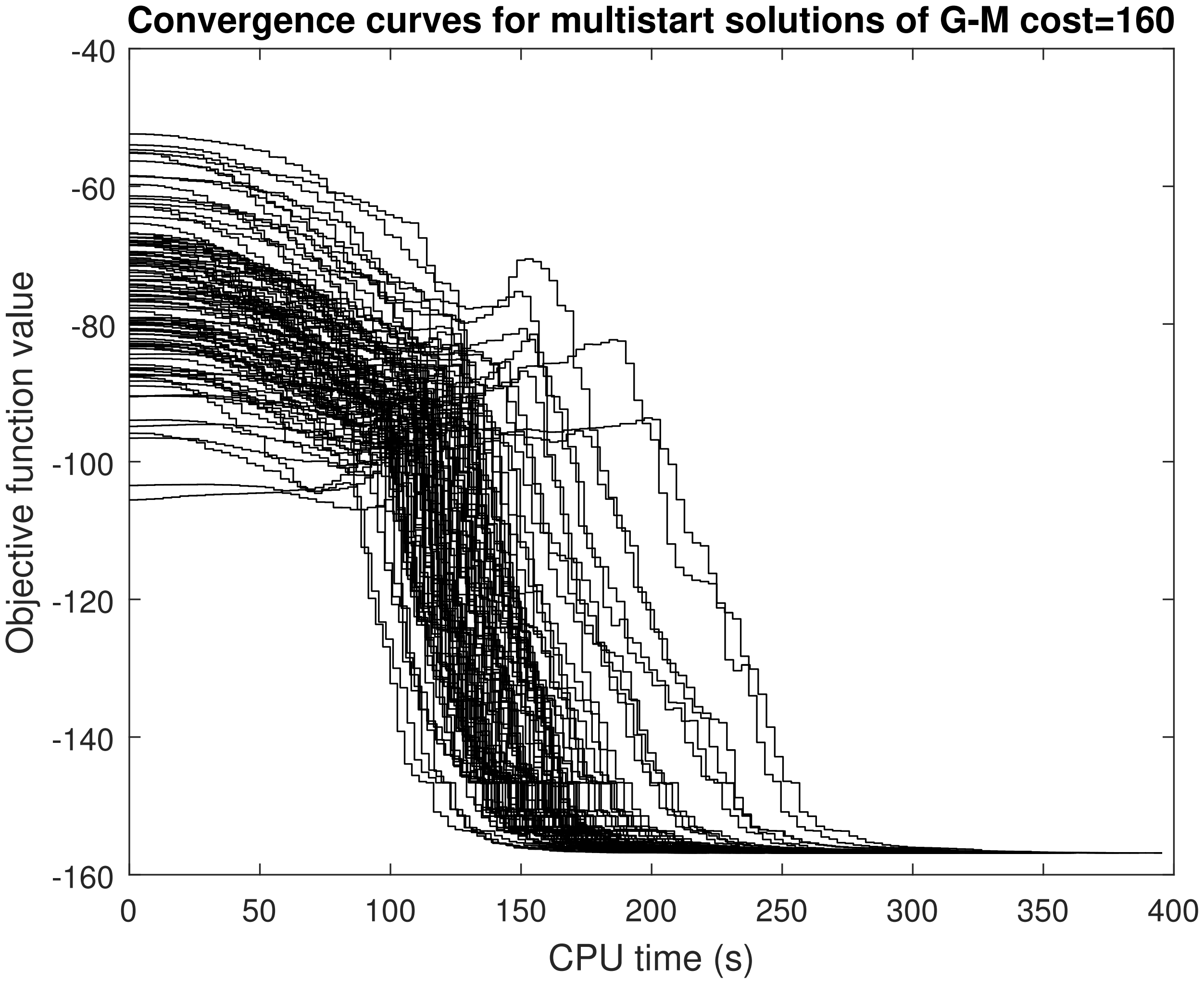


Figure S23: BSUB: Convergence curves of the multistart OCP solutions of a total of 100 ICLOCS runs initialized randomly in the search space for scenario G-M and for point D of the Pareto front.

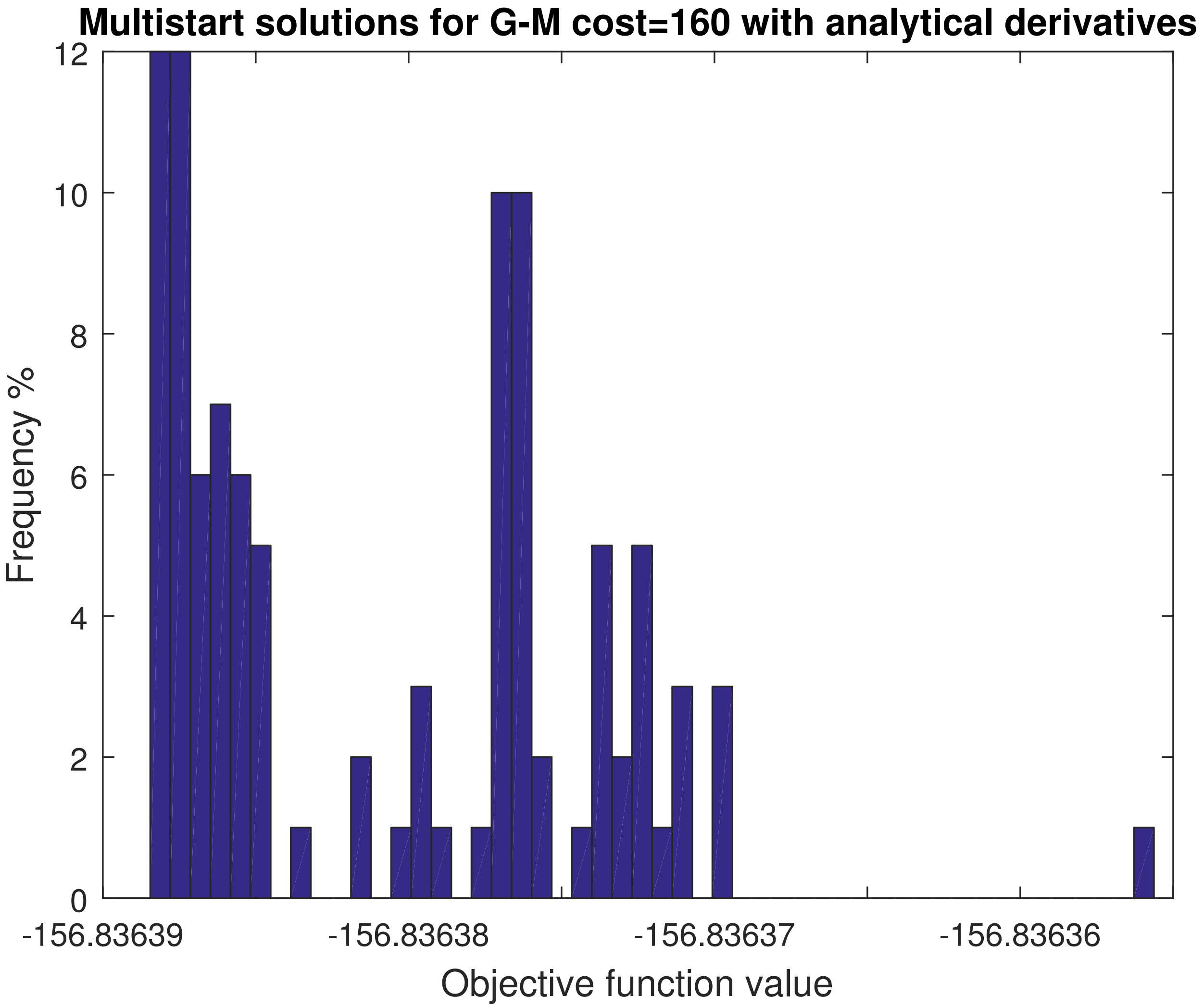


Figure S24: BSUB: Histogram of the multistart OCP solutions of a total of 100 `ICLOCS` runs initialized randomly in the search space for scenario G-M and for point D of the Pareto front.

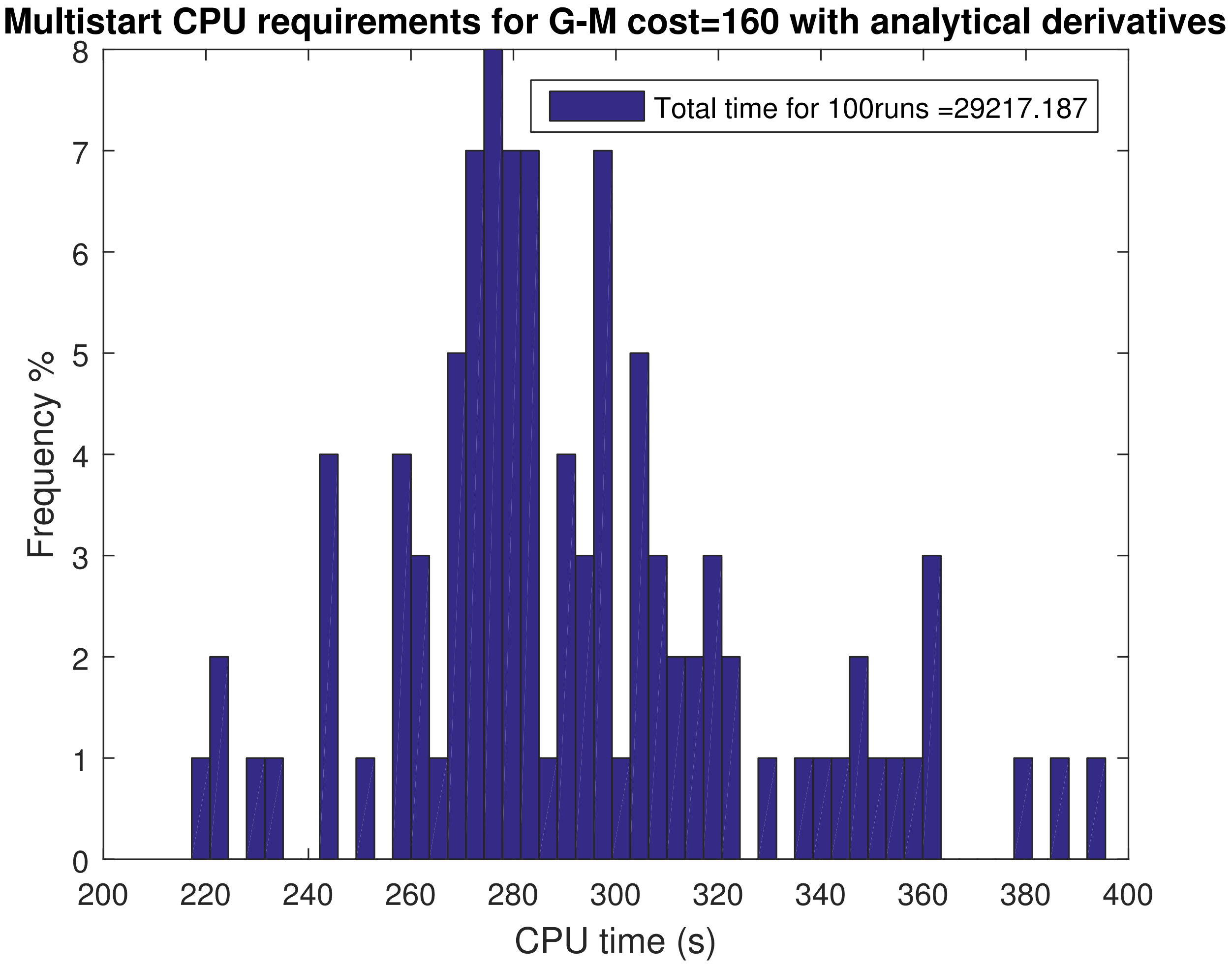


Figure S25: BSUB: Histogram of the computational time required obtaining the multistart OCP solutions of a total of 100 `ICLOCS` runs initialized randomly in the search space for scenario G-M and for point D of the Pareto front.

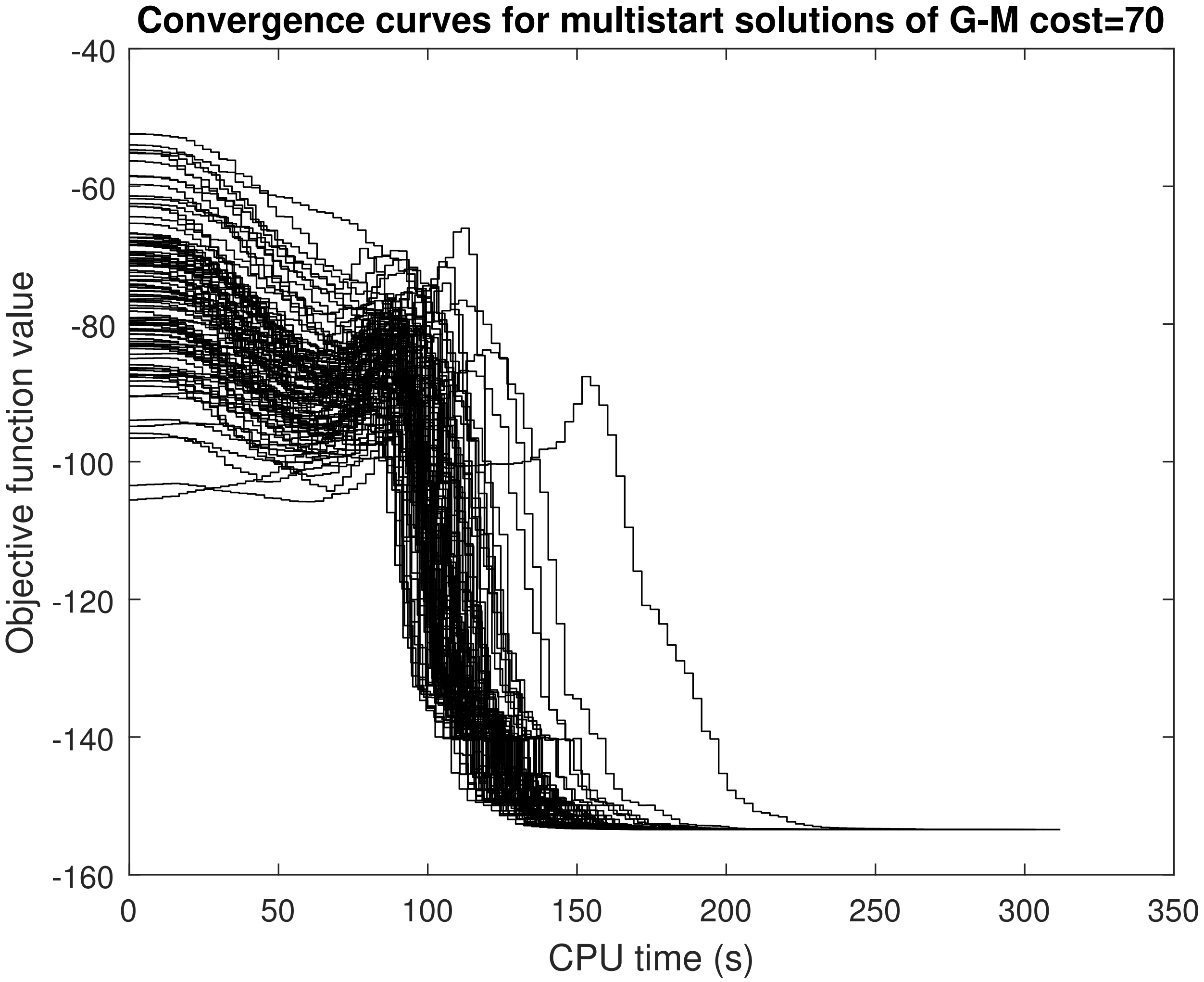


Figure S26: BSUB: Convergence curves of the multistart OCP solutions of a total of 100 ICLOCS runs initialized randomly in the search space for scenario G-M and for point B of the Pareto front.

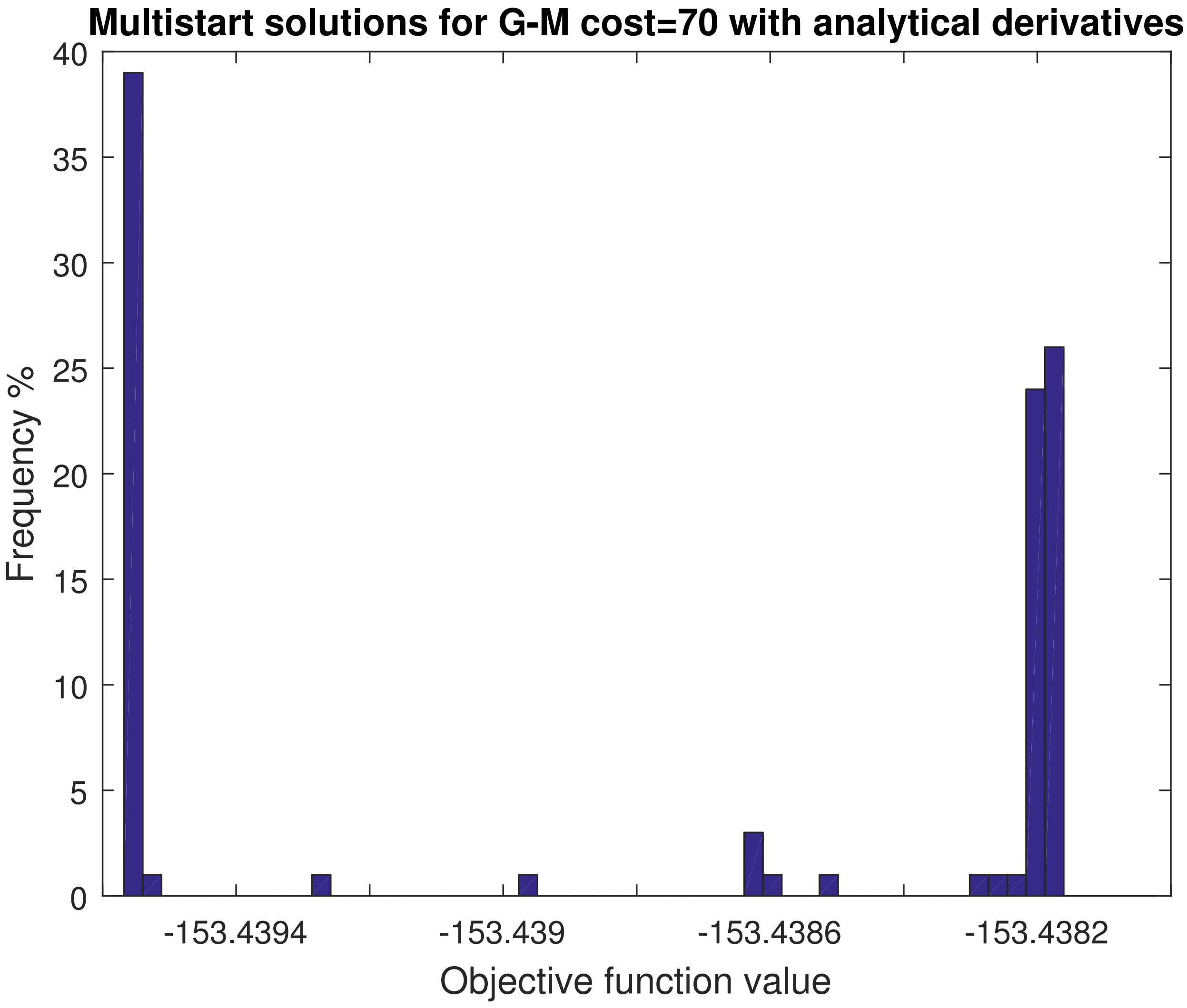


Figure S27: BSUB: Histogram of the multistart OCP solutions of a total of 100 ICLOCS runs initialized randomly in the search space for scenario G-M and for point B of the Pareto front.

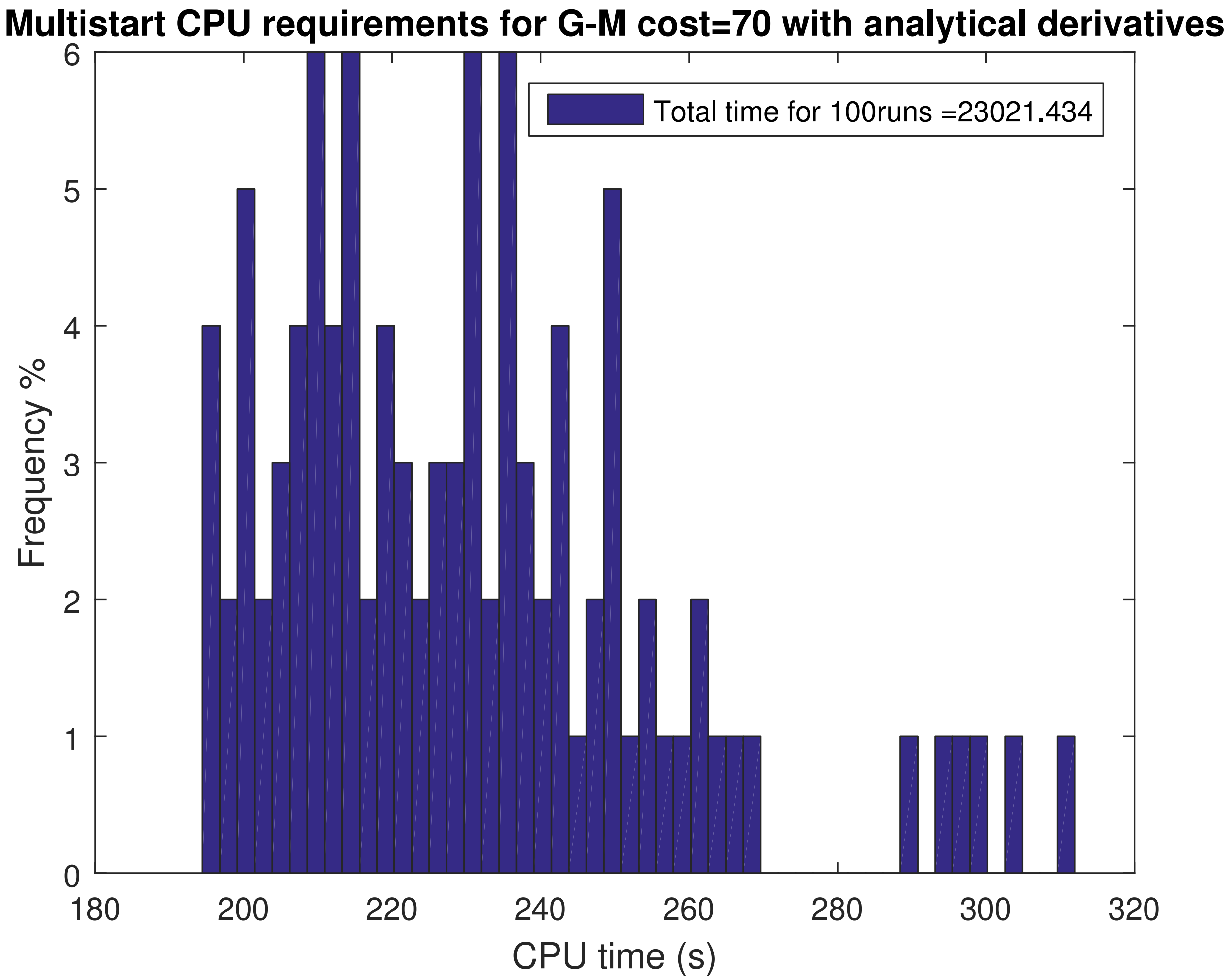


Figure S28: BSUB: Histogram of the computational time required obtaining the multistart OCP solutions of a total of 100 `ICLOCS` runs initialized randomly in the search space for scenario G-M and for point B of the Pareto front.

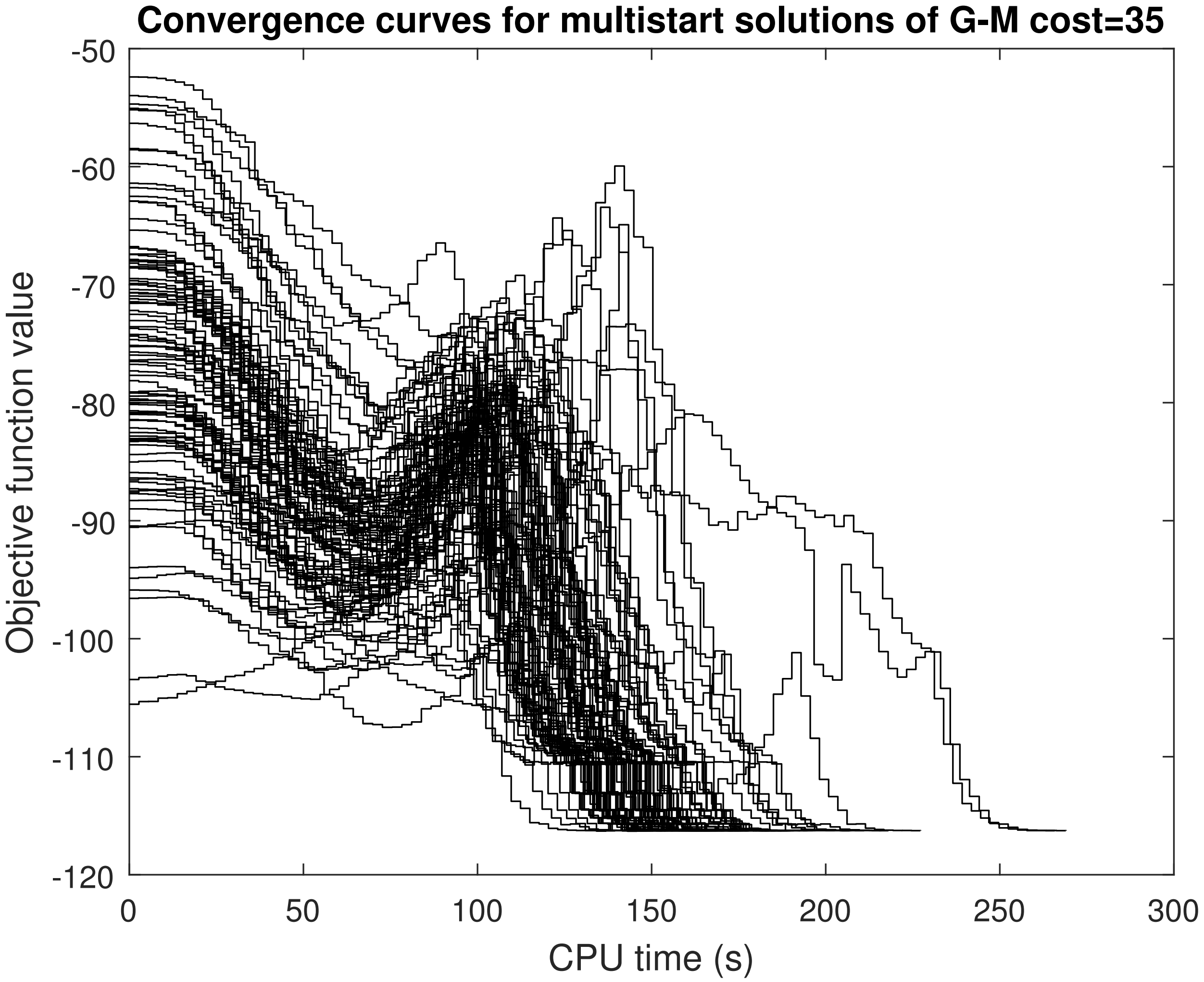


Figure S29: BSUB: Convergence curves of the multistart OCP solutions of a total of 100 `ICLOCS` runs initialized randomly in the search space for scenario G-M and for point A of the Pareto front.

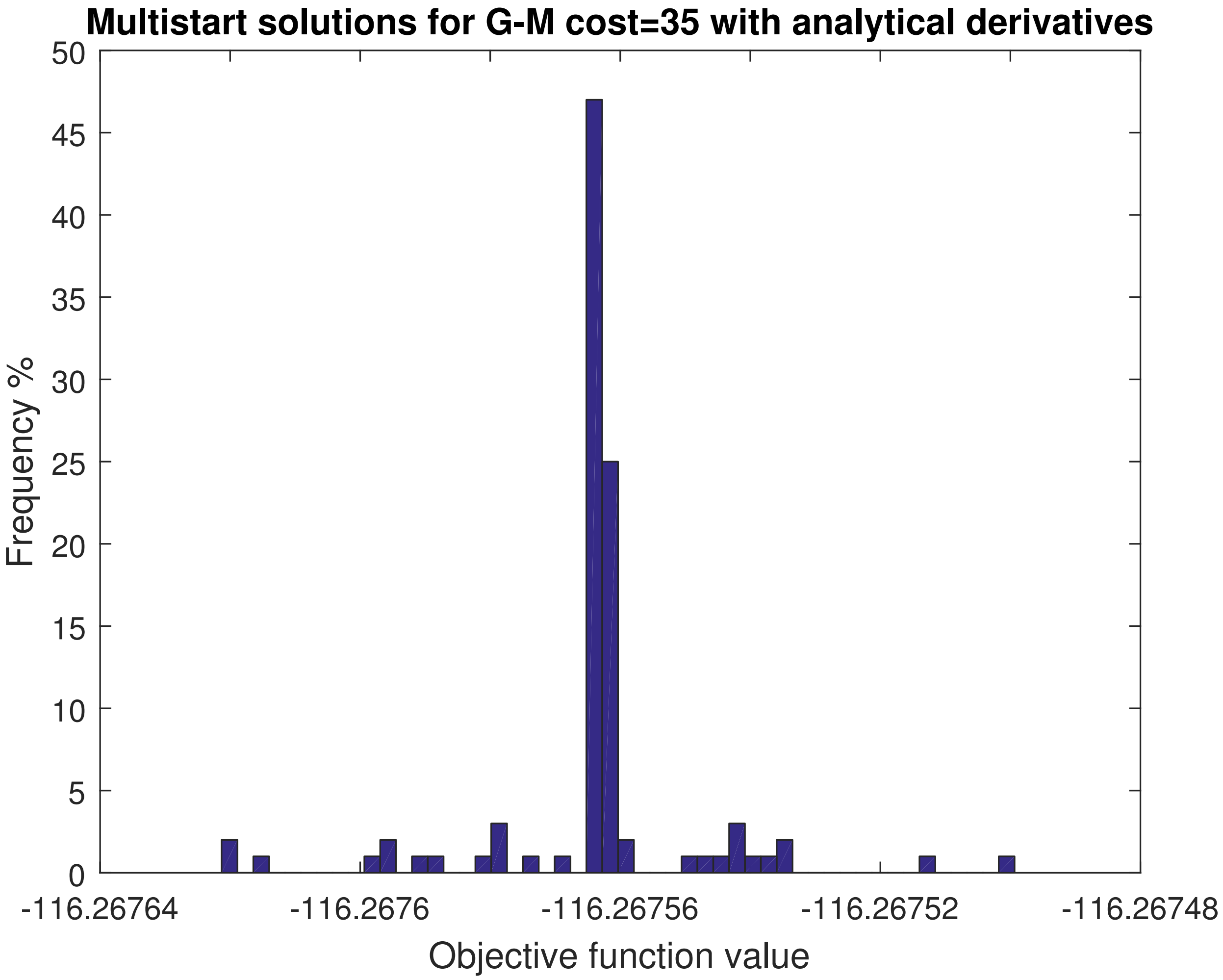


Figure S30: BSUB: Histogram of the multistart OCP solutions of a total of 100 `ICLOCS` runs initialized randomly in the search space for scenario G-M and for point A of the Pareto front.

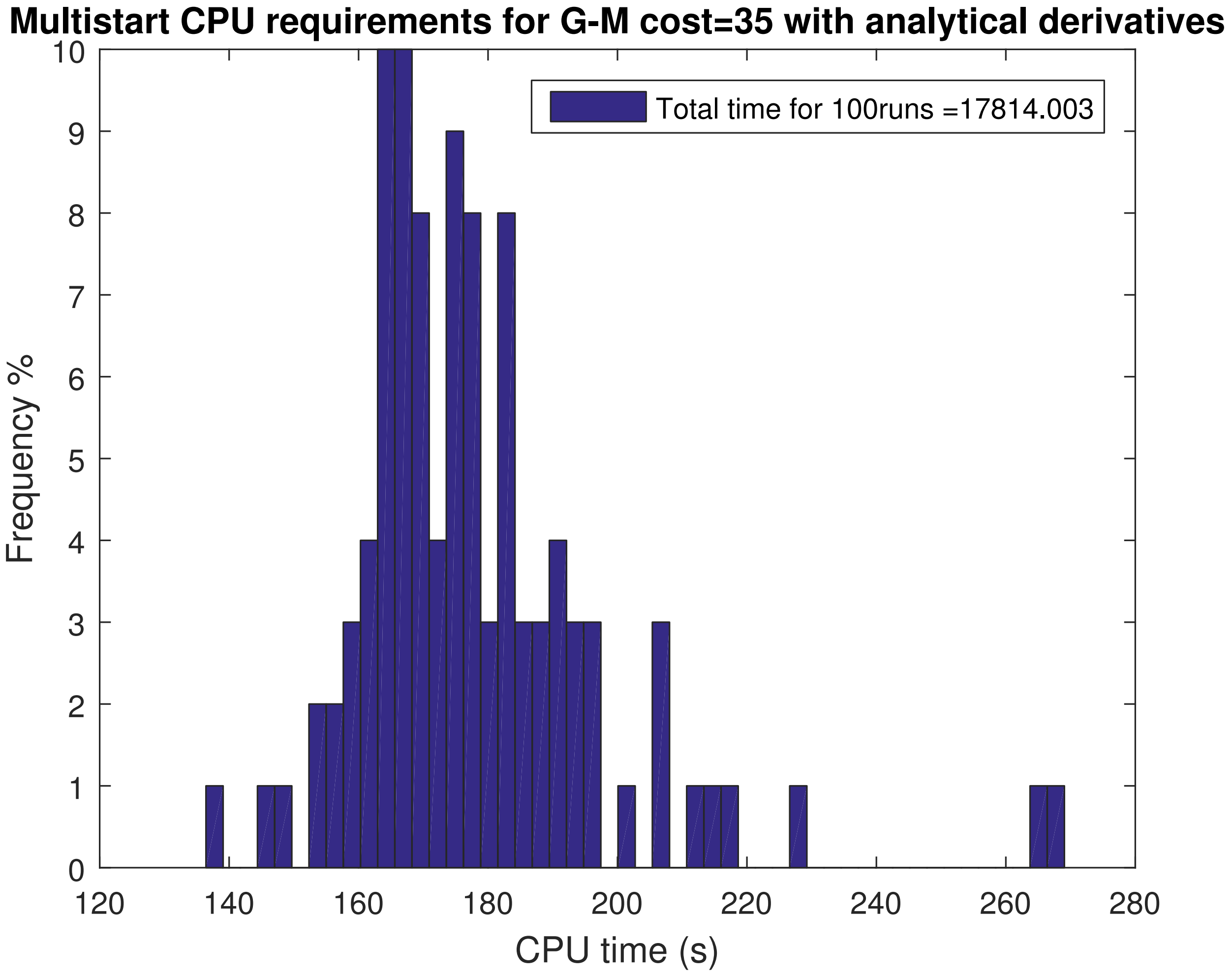


Figure S31: BSUB: Histogram of the computational time required obtaining the multistart OCP solutions of a total of 100 `ICLOCS` runs initialized randomly in the search space for scenario G-M and for point A of the Pareto front.

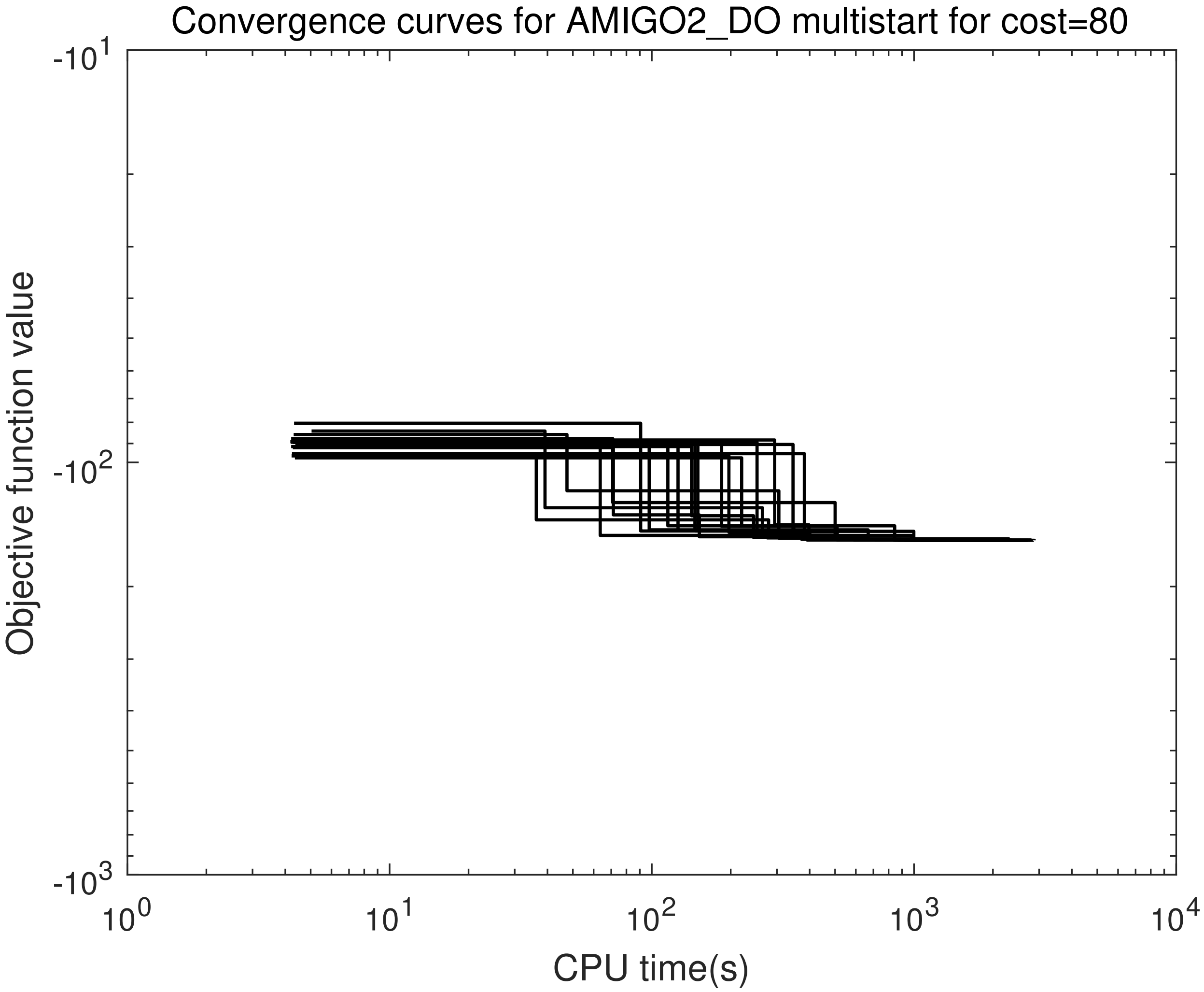


Figure S32: BSUB: Convergence curves of a multistart of 20 runs with AMIGO2_DO (5PWC) for point O of the Pareto front and the G-M scenario.

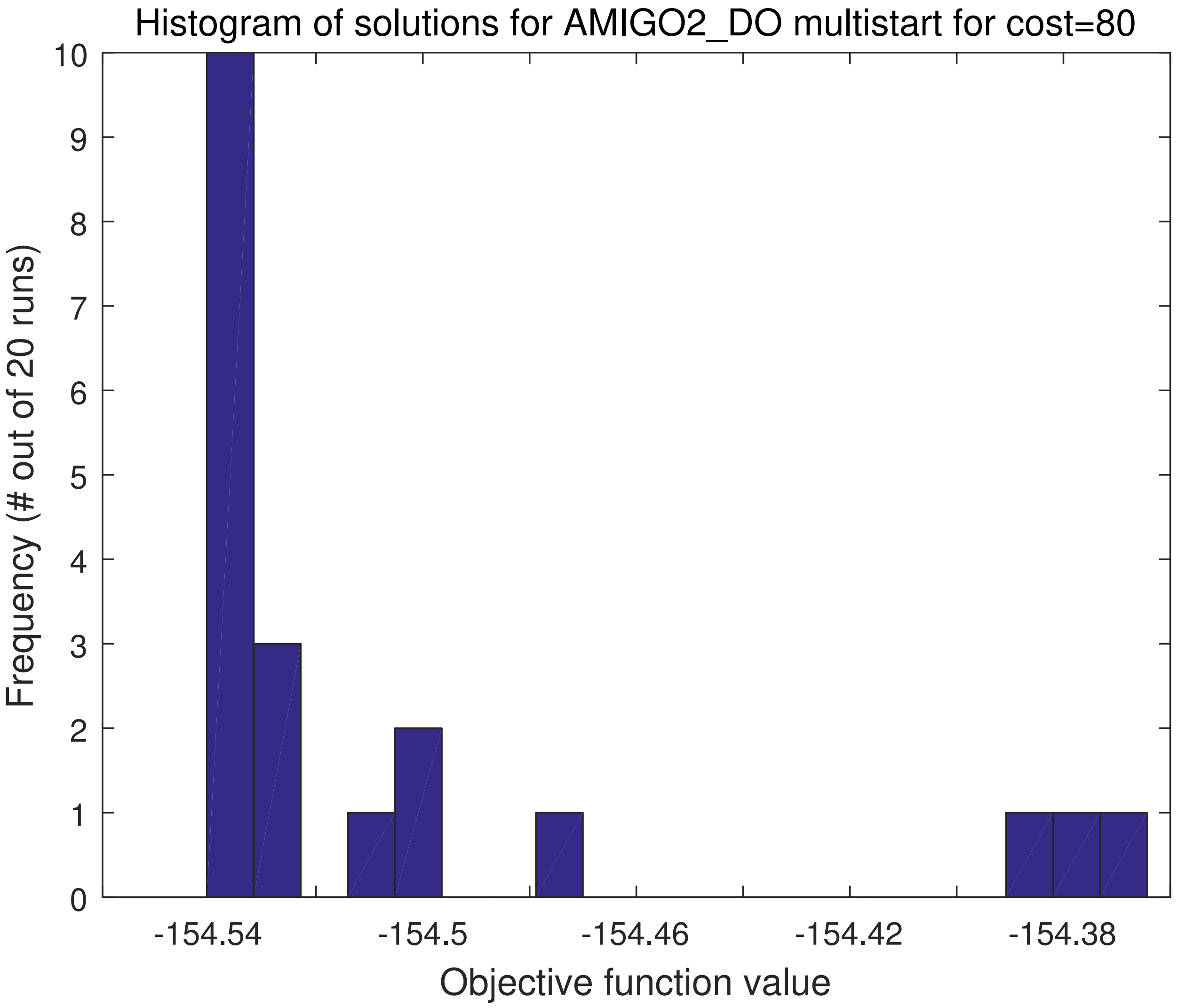


Figure S33: BSUB: Solution histogram of the multistart of 20 runs with AMIGO2_DO (5PWC) for point O of the Pareto front and the G-M scenario.

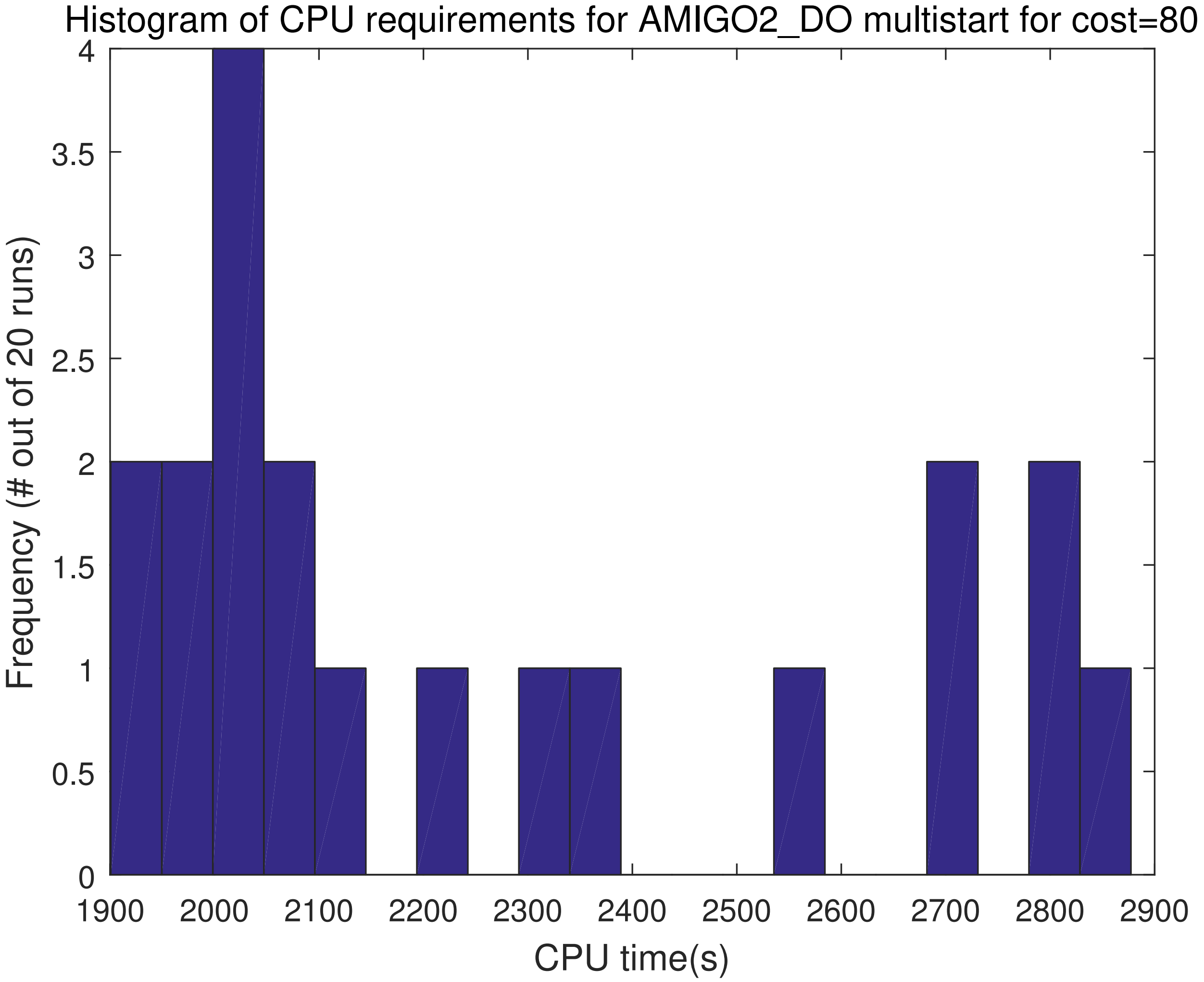


Figure S34: BSUB: Computational requirements histogram of the multistart of 20 runs with AMIGO2_DO (5PWC) for point O of the Pareto front and the G-M scenario.

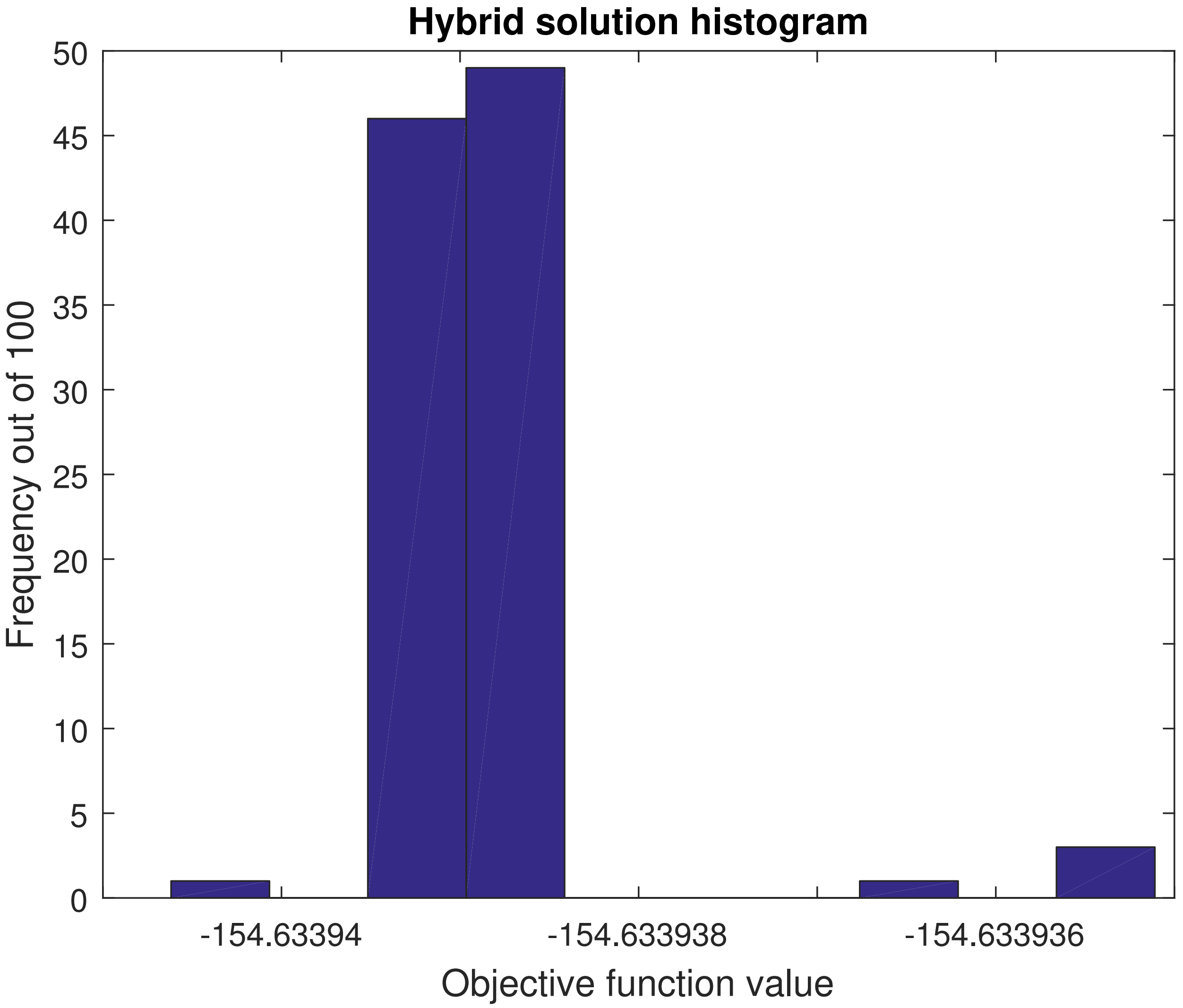


Figure S35: BSUB: Histogram of solution from the multiple AMIGO2_DO+ICLOCS runs for point O of the Pareto front. Starting from different random initial points and performing a fast hybrid run, we attempt to illustrate the possible practical multiplicity of solutions in this scenario. As presented, all runs converge to the same solution.

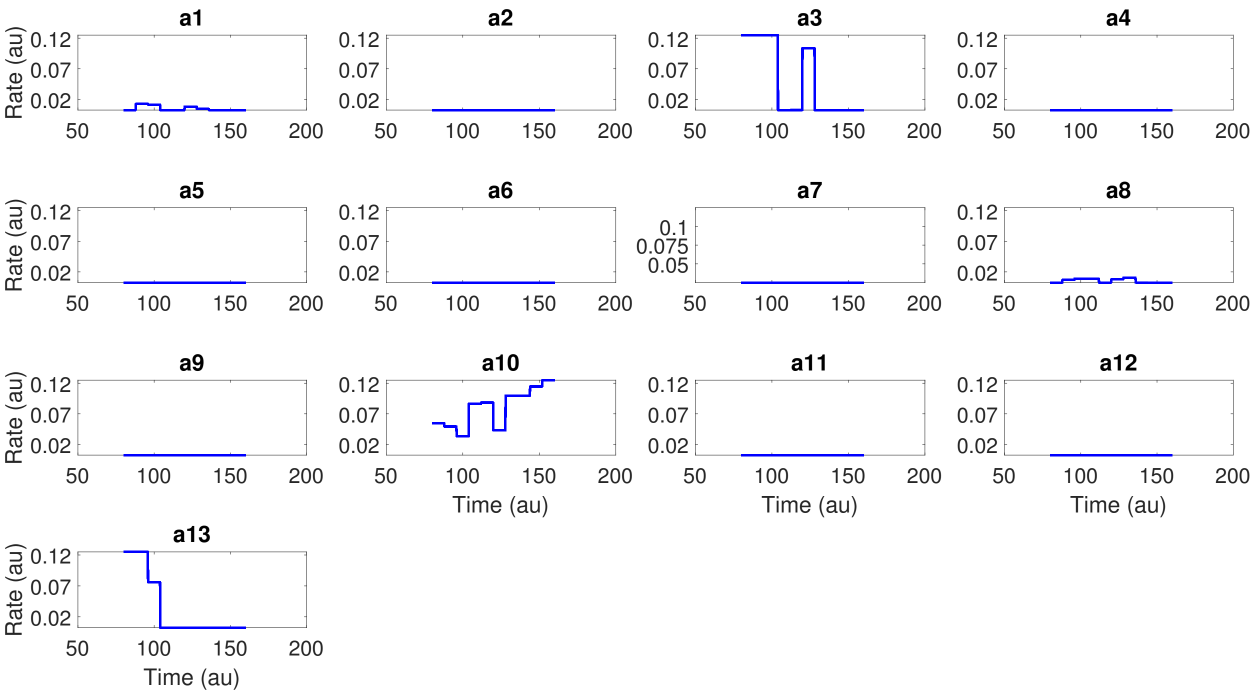


Figure S36: BSUB: Optimal controls from the multiple AMIGO2_DO+ICLOCS runs for point O of the Pareto front, corresponding to Fig S35. Evidently, all runs converge to the same optimal solution.

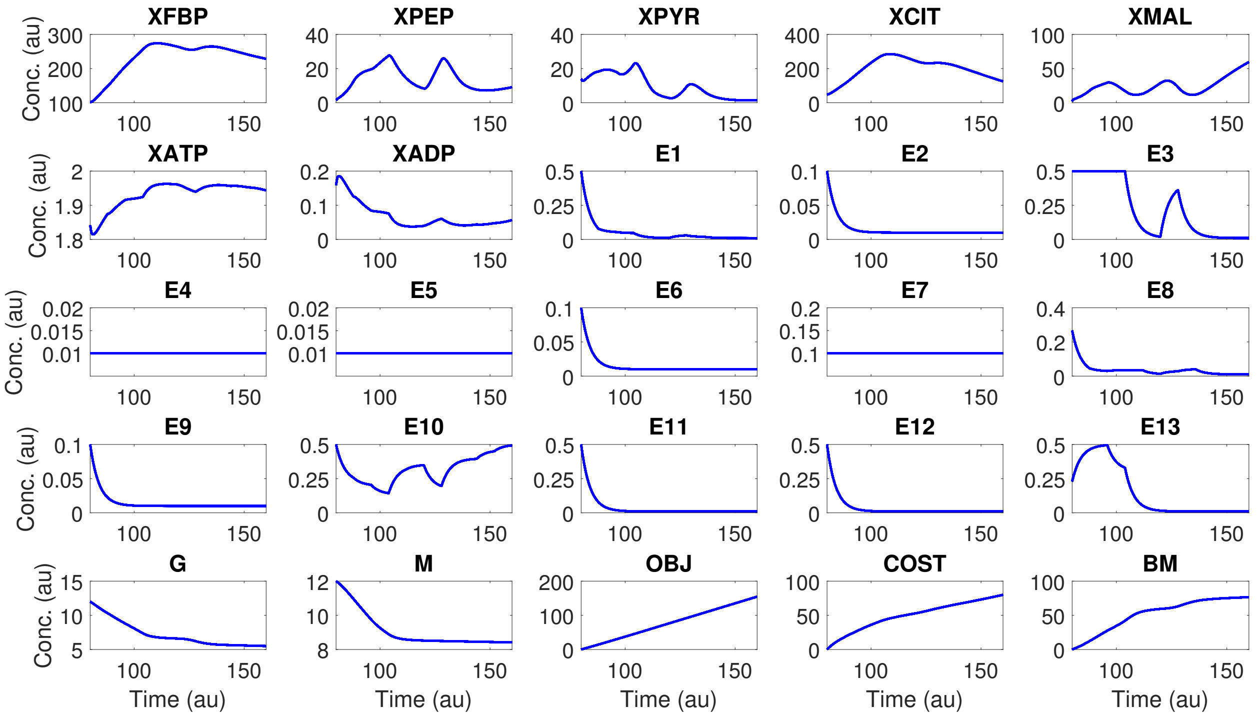


Figure S37: BSUB: Optimal state trajectories from the multiple AMIGO2_DO+ICLOCS runs for point O of the Pareto front, corresponding to Fig S35. Evidently, all runs converge to the same optimal solution.

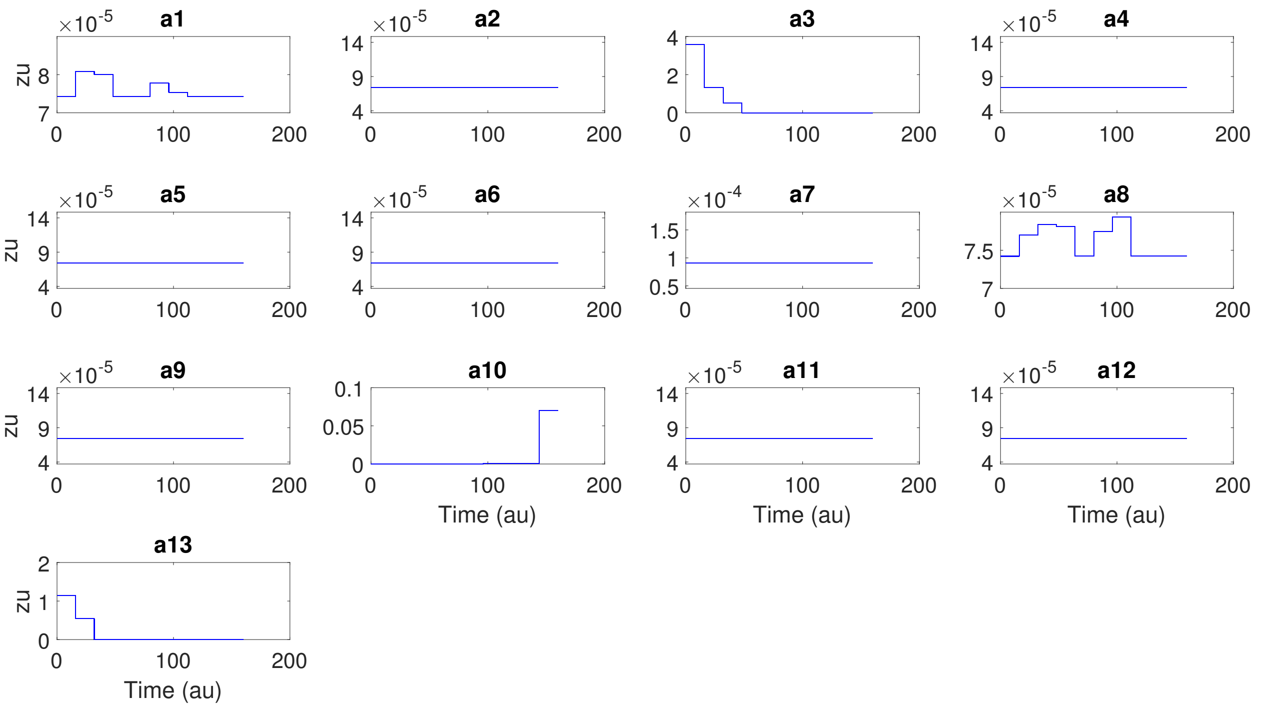

Figure S38: BSUB: All Lagrange multipliers on the controls' upper bounds obtained from the 100 AMIGO2_DO+ICLOCS runs for point O of the Pareto front, corresponding to Fig S35. Evidently, all runs converge to the same optimal solution.

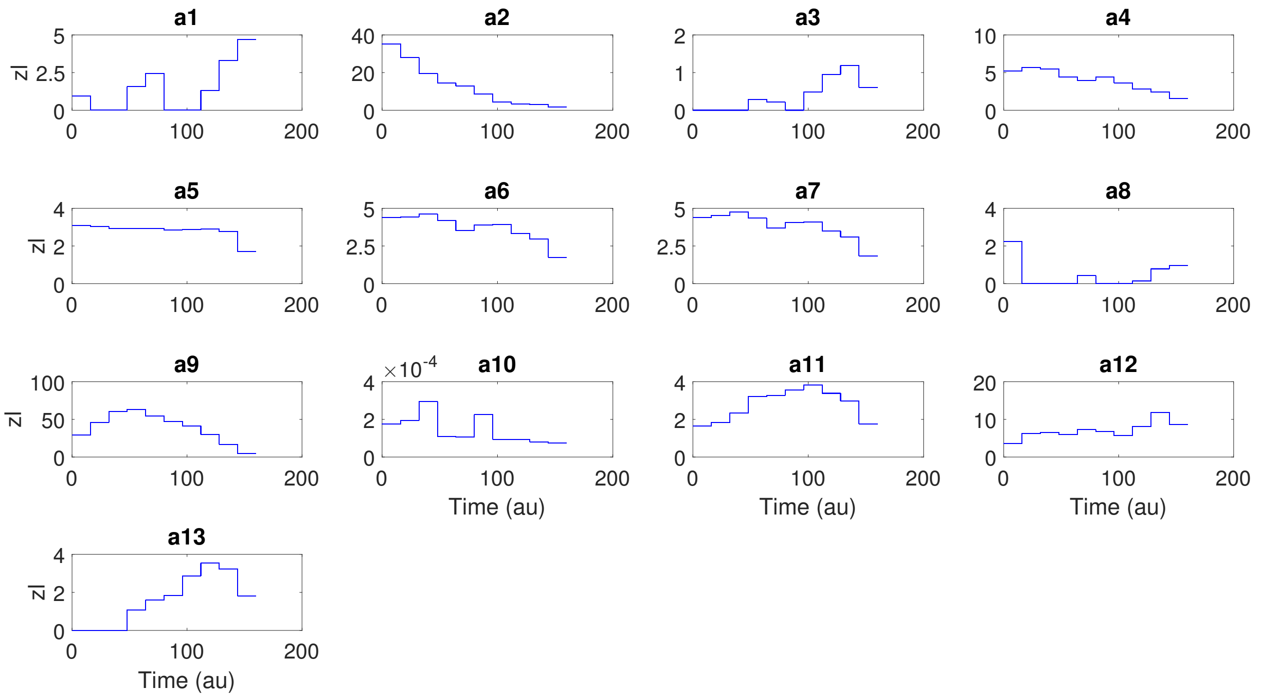

Figure S39: BSUB: All Lagrange multipliers on the controls' lower bounds obtained from the 100 AMIGO2_DO+ICLOCS runs for point O of the Pareto front, corresponding to Fig S35. Evidently, all runs converge to the same optimal solution.

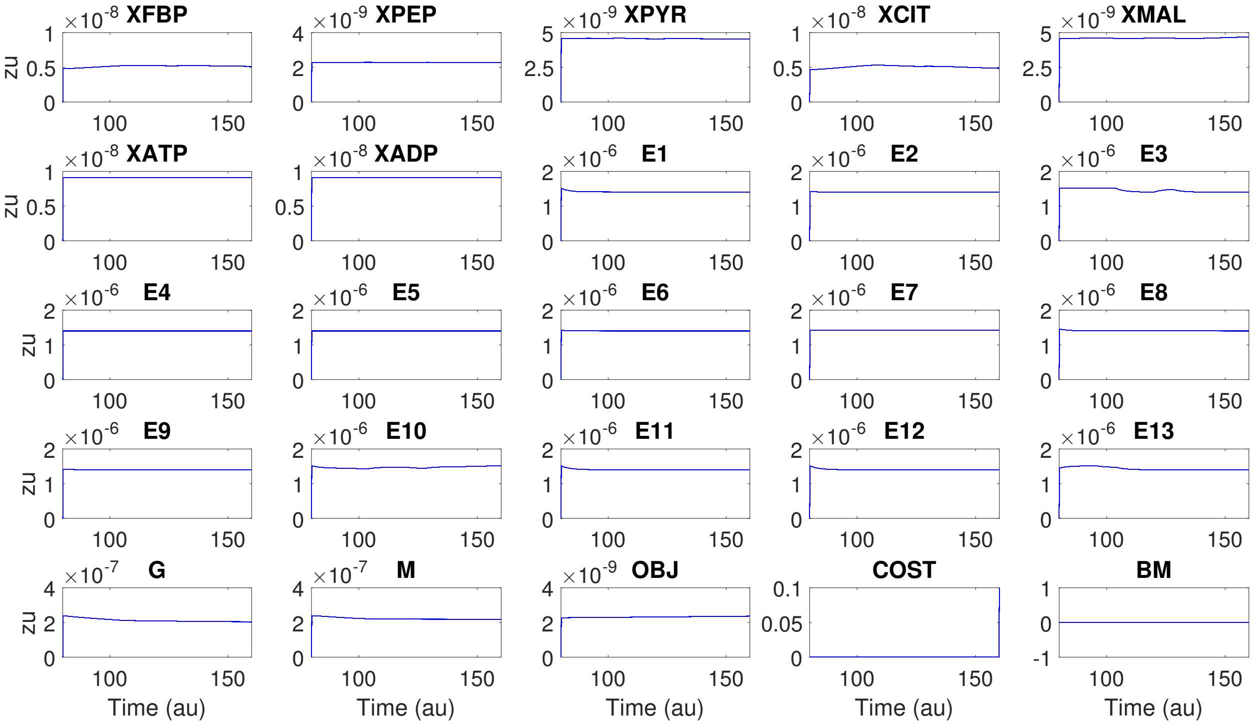


Figure S40: BSUB: All Lagrange multipliers on the states' upper bounds obtained from the 100 AMIGO2_DO+ICLOCS runs for point O of the Pareto front, corresponding to Fig S35. Evidently, all runs converge to the same optimal solution.

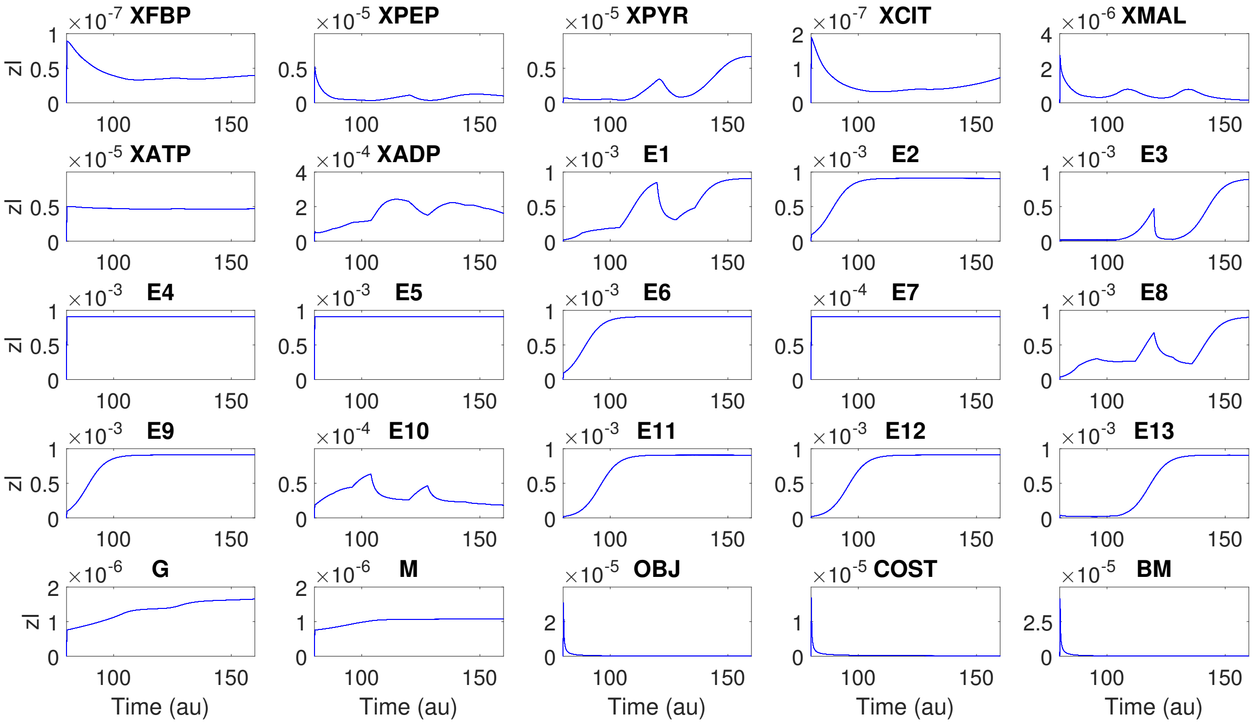


Figure S41: BSUB: All Lagrange multipliers on the states' lower bounds obtained from the 100 AMIGO2_DO+ICLOCS runs for point O of the Pareto front, corresponding to Fig S35. Evidently, all runs converge to the same optimal solution.

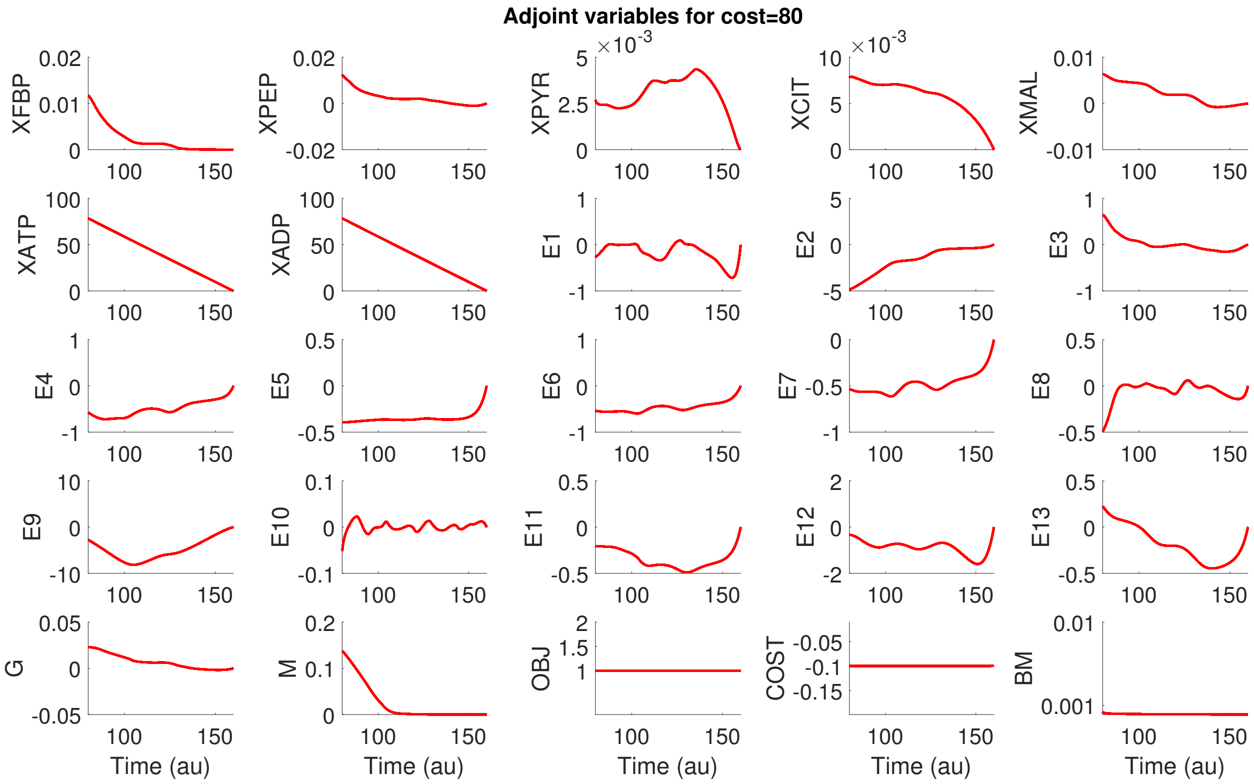


Figure S42: BSUB: All Lagrange multipliers on the states (approximating the adjoint variables) obtained from the 100 AMIGO2_DO+ICLOCS runs for point O of the Pareto front, corresponding to Fig S35. Evidently, all runs converge to the same optimal solution.
